# Dynamics of immune memory and learning in bacterial communities

**DOI:** 10.1101/2022.07.07.498272

**Authors:** Madeleine Bonsma-Fisher, Sidhartha Goyal

## Abstract

From bacteria to humans, adaptive immune systems provide learned memories of past infections. Despite their vast biological differences, adaptive immunity shares features from microbes to vertebrates such as emergent immune diversity, long-term coexistence of hosts and pathogens, and fitness pressures from evolving pathogens and adapting hosts, yet there is no conceptual model that addresses all of these together. To address these questions, we propose and solve a simple phenomenological model of CRISPR-based adaptive immunity in microbes. We show that in coexisting phage and bacteria populations, immune diversity in both populations emerges spontaneously and in tandem, that bacteria track phage evolution with a context-dependent lag, and that high levels of diversity are paradoxically linked to low overall CRISPR immunity. We define average immunity, an important summary parameter predicted by our model, and use it to perform synthetic time-shift analyses on available experimental data to reveal different modalities of coevolution. Finally, immune cross-reactivity in our model leads to qualitatively different states of evolutionary dynamics, including an influenza-like traveling wave regime that resembles a similar state in models of vertebrate adaptive immunity. Our results show that CRISPR immunity provides a tractable model, both theoretically and experimentally, to understand general features of adaptive immunity.

## 2 Introduction

Adaptive immunity equips organisms to survive changing pathogen attacks across their lifetime. Many diverse organisms from bacteria to humans possess adaptive immune systems, and their presence shapes the survival, diversity, and evolution of both hosts and pathogens. How adaptive immunity changes the landscape of host-pathogen coexistence, how immune diversity emerges and evolves, and how the pressures of evolving pathogens and adaptive immunity are coupled to produce unique evolutionary outcomes: all of these factors are of fundamental importance to understanding the role of adaptive immunity in populations.

These questions have naturally been explored in the vertebrate adaptive immune system, which protects humans and other vertebrates from evolving pathogens. In these organsisms, a diverse repertoire of T cell and B cell receptors can rapidly recognize and respond to a wide range of threats. Immune specificity is determined by the unique genetic sequence of each cell’s receptor, and individuals may harbour millions to billions of unique sequences distributed across four or more orders of magnitude of abundance [1, 2, 3]. Quantitative frameworks to model immune diversity and clone abundance have revealed that simple low-level interactions can give rise to complex outcomes including broad distributions of clone abundance [1, 2, 3, 4, 5, 6], long-lived biologically realistic transient states [7, 5], and clonal restructuring following immune challenges [8, 9, 10, 11, 5]. Phenomenological models of pathogen co-evolution with the immune system have accelerated our understanding of how the fitness landscape generated by the immune system constrains pathogen evolution [12, 13, 7, 14, 15], how the adaptive immune system responds to rapid pathogen evolution [16, 17, 14, 15], and what drives pathogen extinction [7, 13] or the extinction of particular clonal cell lineages [17, 10]. These models have also explored trade-offs such as between immune receptor specificity and cross-reactivity [4, 18], between the specifity of host-pathogen discrimination and sensitivity to pathogens [8, 19, 20], between the speed of an immune response and the efficiency of that response [14], or between metabolic resource use and immune coverage [15]. All of these models have shown rich dynamics and qualitatively different states of diversity and evolution arising from simple rules. However, experiments in vertebrates are difficult: vertebrate immunity depends on a complex interplay of many protein components and experiments are time-consuming because of long generation times [21].

Adaptive immunity in microbes is realized through the CRISPR system, conceptually similar to the vertebrate adaptive immune system. The CRISPR system is functionally simple, yet it is incredibly powerful, as indicated by its widespread presence in many diverse bacteria and archaea [22] and its experimentally-demonstrated ability to provide strong immunity against phages [23, 24, 25, 26]. Attacking phages expose their DNA to bacteria, and bacteria with a CRISPR immune system acquire small segments of phage DNA, called spacers. They store spacers in their genome and use them to recognize and destroy matching phage sequences in future infections: spacers are transcribed into RNA and guide DNA-cleaving CRISPR-associated proteins to recognize and cut re-infecting phages. Spacers provide a highly specific immune memory of infecting phages, preventing recognized phages from reproducing. In turn, phages can acquire mutations in the protospacer regions of their genome that are targeted by spacers. These features of the CRISPR immune system mean that (a) phage genetic evolution occurs by selection for escape mutants, and (b) the network of CRISPR immune interactions between bacteria and phages can be inferred by sequencing the genomes of co-living bacteria and phages. Spacer acquisition and phage mutation are rare random events, and many such events must be observed in order to understand their impact on populations. Bacteria and phages have short life cycles and can reach large population size, making it possible to build a statistical picture of the impacts of adaptive immunity.

The kinetics and interactions of phages and bacteria with CRISPR systems have been the subject of numerous experiments [25, 27, 28, 29, 30, 31]. Some themes have emerged from experimental studies of CRISPR immunity: (a) high spacer diversity relative to phage diversity increases the likelihood of phage extinction [25, 28, 31], (b) bacteria become more immune to phages over time [32, 33, 27, 34], and (c) phages readily gain mutations [35, 23, 36, 37, 38, 34, 31, 39] and sometimes genome rearrangements [24] to escape CRISPR targeting. Explorations of CRISPR immunity in natural environments have also documented ongoing spacer acquisition and phage escape mutations [35, 39]. Likewise, previous theoretical work has addressed the impact of parameters such as spacer acquisition rate and phage mutation rate on spacer diversity [40, 41, 42] and population survival and extinction [43], how costs of CRISPR immunity impact bacteria-phage coexistence [44] and the maintenance of CRISPR immunity [45, 43, 46, 47], how spacer diversity impacts population outcomes [48, 35, 40, 49, 41, 50, 51, 42], and how stochasticity and initial conditions impact population survival [52, 53]. Notably, foundational work by Childs *et al*. [40, 50] and Weinberger et al. [35, 43] found through simulations that spacer diversity readily emerges in a population of CRISPR-competent bacteria interacting with mutating phages.

However, the majority of both experiments and theory are based on observations and models of transient phenomena and short-term dynamics, while it is at long timescales that natural microbial communities experience bacteria-phage coexistence. Some notable experiments have measured long-term coexistence [24, 54], and long-term sequential sequencing data from natural populations is becoming more available [55, 56], but appropriate theories to understand steady-state coexistence, sequence evolution and turnover, and immune memory in microbial populations remain rare. Because the processes of growth, death, and immune interaction are inherently random, understanding population establishment and extinction requires a fully stochastic analysis, and theoretical models that explore long-term coexistence have been partially deterministic to date [43, 35, 40, 36, 50, 57, 58, 47]. These models do not accurately capture rare stochastic events, in particular mutation, establishment, and extinction. Notable fully stochastic simulations of CRISPR immunity, on the other hand, have lacked rigorous analytic results [41, 42].

To understand the emergent properties of immune memory and diversity in microbial populations and how phages and bacteria coexist long-term, we developed a simple theoretical model of bacteria and phages interacting with adaptive immunity. We model a population of bacteria with CRISPR immune systems interacting with phages that can mutate to escape CRISPR targeting, building on our previous work [59]. We stochastically simulate thousands of bacteria-phage populations across a range of population sizes, spacer acquisition rates, spacer effectiveness rates, and phage mutation rates, and derive analytic expressions for the probability of establishment for new phage mutants, the time to extinction for phage and bacterial clones, and the dependence of bacterial spacer diversity on spacer acquisition rate, effectiveness, and phage mutation rate. We show that even with the simplest assumptions of uniform spacer acquisition and effectiveness, complex dynamics and a wide range of outcomes of diversity and population structure are possible. We link diversity to the dynamical quantities of establishment and extinction, and show that the length of immune memory is limited by these same quantities: bacteria can either follow phages quickly or remember for a long time, but not both. We compute bacterial average immunity in our simulations and in available experimental data and show that our model predicts qualitative trends that are visible in data. Finally, we show that adding immune cross-reactivity leads to qualitatively different states of evolutionary dynamics: (1) a traveling wave regime that resembles a similar state in models of vertebrate adaptive immunity [7, 13, 60] emerges when high cross-reactivity creates a fitness gradient for phage evolution, and (2) a regime of low turnover protected from new establishment by the reduced fitness of new phage mutants.

## 3 Results

### 3.1 Bacteria and phages dynamically coexist and coevolve

We model bacteria and phage interacting and coevolving in a well-mixed system (Figure 1A, Methods, and SI Section 1). Bacteria divide by consuming nutrients and phages reproduce by creating a burst of *B* new phages after successfully infecting a bacterium. Bacteria can contain a single CRISPR spacer that confers immunity against phages with a matching protospacer; phages are labeled with a single protospacer type that can mutate to a new type during a burst with probability *µL*, where *µ* is the per-base mutation rate per generation and *L* = 30 is the protospacer length.

Coexistence occurs across a wide range of parameters but is not guaranteed: below a certain success probability 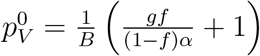, phages are not able to reproduce often enough to overcome their base death rate due to outflow and adsorption and are driven extinct (Figure 1D) [59]. In this expression, *g* is the bacterial growth rate, *B* is the phage burst size, *α* is the phage adsorption rate, *f* = *F*/(*gC*_0_) is a normalized outflow rate, and *C*_0_ is the inflow nutrient concentration. This is the same extinction threshold reported by Payne *et al*. [61] as the cutoff for achieving herd immunity in a well-mixed bacterial population. To a first approximation, phages must successfully infect every 1/*B* bacteria they encounter, but if bacteria are growing quickly, then phages must do better to overcome bacterial growth, leading to the extra terms in this expression (see SI section 2.1.1). We write this extinction threshold as *A* = 1, where 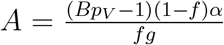. Above the phage extinction threshold (*A >* 1), the phage population size increases with increasing *p*_*V*_ but eventually decreases again as bacterial numbers are driven too low to support a large phage population [59]. A similar non-monotonicity as a function of the probability of naive bacterial resistance (1 − *p*_*V*_) was described in theoretical work by Weinberger *et al*. [43]. The position of the peak in phage population size as a function of infection success probability is determined by *e*, the effectiveness of CRISPR spacers against phage; increasing *e* pushes the peak to higher *p*_*V*_. While *e* is a constant parameter that determines the outcome of pairwise interactions between bacteria and phages, the bacterial population as a whole possesses an average immunity to phages that is a weighted average of all the possible pairwise interactions (Figure 1A inset). It is the overall average immunity that determines population outcomes, which we describe in detail in section 3.7.

**Figure 1:**
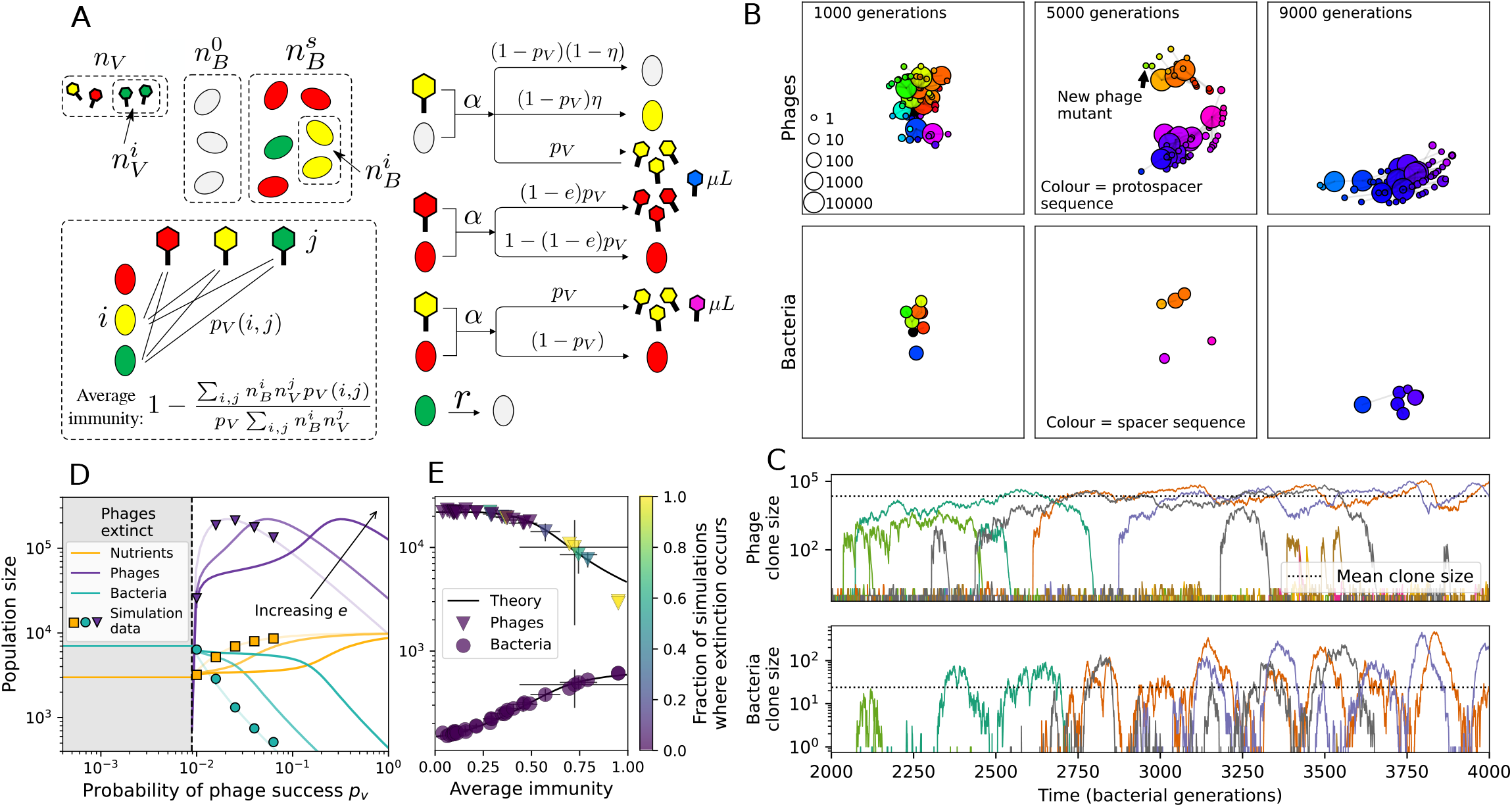
(A) We model bacteria and phages interacting in a well-mixed vessel. We track nutrient concentration (*C*), phage population size (*n*_*V*_), and bacteria population size (*n*_*B*_). Bacteria can either have no spacer 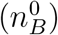 or a spacer of type 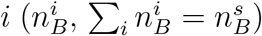, and phages can have a single protospacer of type 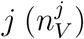. With rate *α*, a phage interacts with a bacterium. If the bacterium does not have a matching spacer, the phage kills with probability *p*_*V*_ and produces a burst of *B* new phages, while for bacteria with a matching spacer that probability is reduced to 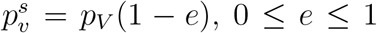. Bacteria without spacers that survive an attack have a chance to acquire a spacer with probability *η*, and bacteria with spacers lose them at rate *r*. Lower inset: average immunity is the weighted average pairwise immunity between spacer-containing bacteria and phages, given by 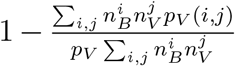. The probability of a phage with protospacer *j* successfully infecting a bacterium with spacer *i* is *p*_*V*_ (*i, j*). (B) Three time points in a typical simulation with *C*_0_ = 10^4^, *e* = 0.95, *η* = 10^−4^, and *µ* = 10^−5^. Coloured circles represent unique protospacer or spacer sequences; shared sequences are shown with the same colour. The size of each circle is proportional to clone size, and new mutants are shown radially more distant from the centre. (C) Ten individual clone trajectories vs. simulation time for phages (top) and bacteria (bottom). The mean clone size is shown with a horizontal dashed line. (D) Total phage, bacteria, and nutrient concentration as a function of phage success probability *p*_*V*_. Markers show an average over 5 independent simulations for different values of *p*_*V*_ with *C*_0_ = 10^4^, *η* = 10^−3^, *e* = 0.95, and *µ* = 10^−7^. Solid lines show theoretical predictions for different constant values of effective *e*. As *p*_*V*_ decreases, phages go extinct at a critical value given by *A* = 1, where 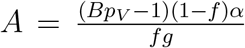. (E) Total phage and bacteria population size as a function of average bacterial immunity to phages. Colours indicate the fraction of simulations that end in phage or bacteria extinction before a set endpoint. Solid lines show the mean-field prediction.

To focus on regimes where bacteria and phages coexist, we select parameters within the deterministic coexistence regime to explore bacteria-phage coevolution. Even in this regime, stochastic extinction will eventually come for one or both populations in simulations (Figure 1E), though the timescale of extinction may be extremely long for large population sizes [62]. Phages are more susceptible to stochastic extinction than bacteria because of their large burst size *B* which increases their overall population fluctuations (SI section 5.3). The length of coexistence before stochastic extinction depends on population size as well as simulation parameters and initial conditions: phage populations can be rescued from extinction by high mutation rate or high initial protospacer diversity, but are more likely to go extinct if spacer effectiveness is high. Conversely, bacteria are more likely to go extinct if spacer effectiveness or spacer acquisition rate are low (see SI Section 3.2). Population survival and persistence in natural populations is impacted by additional factors we do not address in our model, including immigration [63, 64], niche partitioning [65, 66, 67, 62, 68], environmental fluctuations [69, 68], and spatial structure [70, 49, 71, 72, 73, 44].

Across a wide range of coexistence parameters, our simulations show continual phage evolution and bacterial CRISPR adaptation in response (Figure 1B). New phage protospacer clones arise by mutation, and a small fraction of new mutants grow to a large size and become established. Once phage clones become large, bacteria acquire matching spacers and an immune bacterial subpopulation becomes established. The specific protospacer and spacer types present in the population continually change as old types go extinct and new types are created by phage mutation, but the average total diversity and average overlap between bacteria and phage remains constant at steady-state (SI Figure 1). Both bacteria and phage clones stochastically go extinct, completing the life cycle of a clonal population (Figure 1C).

### 3.2 Phages drive stable emergent sequence diversity

Over time in our simulations, new phage protospacer clones continually arise and go extinct, generating turnover in clone identity in the population. Despite constant turnover, however, the total number of clones remains fixed at steady state. We use the mean number of bacterial clones at steady state, designated *m*, as a measure of system diversity. This choice of diversity measurement is equivalent to the Hartley entropy of the clone size distribution, a special case of the Rényi entropy [74, 21]. This definition weights all clones equally regardless of their abundance; such a measurement is not appropriate when clone size distributions are very broad and small clones may be unsampled, but is reasonable when clone size distributions are relatively narrow and all clones are sampled [74]. In our simulation results, both bacteria and phage populations exhibit relatively narrow clone size distributions across a range of parameters, with the exception of low values of spacer acquisition *η* (Figure 2A). Even at low *η*, however, clone size distributions are approximately exponential, indicating that they are not scale-invariant and that the mean clone size still captures important information about the full clone size distribution.

**Figure 2:**
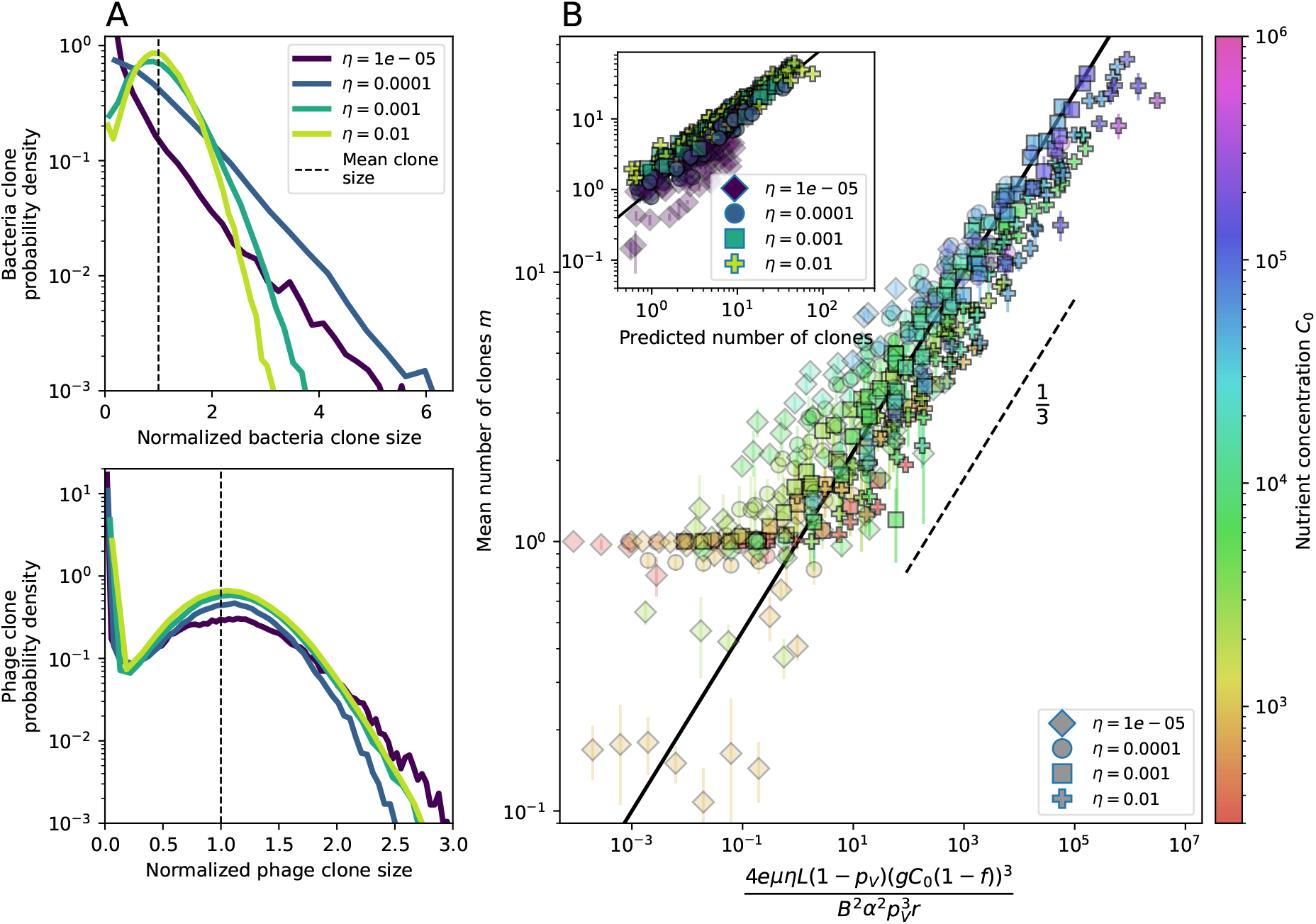
Diversity depends sub-linearly on parameters. (A) Bacteria and phage clone size distributions normalized to the measured mean clone size for *C*_0_ = 10^5^, *µ* = 3 × 10^−7^, and *e* = 0.95. As *η* increases, both clone size distributions become more sharply peaked. (B) The mean number of bacterial clones depends only on a combined parameter in the limit of small average immunity (generally coinciding with high *C*_0_). (Inset) the mean number of bacterial clones can be predicted by numerically solving equation 1 for *m*. The two lowest values of *η* are shown with lighter shading.

What determines clonal diversity? Many factors that correlate with transient diversity have been experimentally identified, such as phage extinction and slower phage evolution at high bacterial spacer diversity [25, 28] and maintenance of a diverse bacterial population when exposed to diverse phages [24, 27, 31, 75], but a conceptual framework to understand emergent diversity has remained elusive. For instance, while initial high spacer diversity puts low-diversity phage populations under intense pressure to the point of driving them extinct [25, 28], is the same true for emergent bacterial diversity after an extended period of coexistence? Is observed high bacterial spacer diversity indicative of successful bacterial escape from phage predation, or an indicator of increased phage pressure? In our model, phage and bacterial diversity is tightly coupled: the number of large phage clones is approximately the same as the number of bacterial clones (SI Figure 24). This is also the case in experimental coevolution data from ref. [24]: the number of phage protospacer types is on the same order of magnitude as the number of bacterial spacer types across most similarity thresholds (SI Figure 95). There is evidence that this coupling of diversity may also occur in the wild: a recent longitudinal study of *Gordonia* bacteria interacting with phage in a wastewater treatment plant identified 14 high-coverage phage genotypes and 11 high-coverage bacterial variants based on CRISPR spacer sequence [39].

Using the tight correspondence between bacterial diversity and phage diversity, we calculate the overall steady-state diversity by balancing the effective phage clone mutation rate 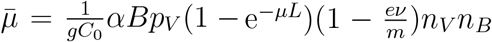, phage clone establishment probability *P*_*est*_, and the time to extinction for large phage clones *T*_*ext*_ (details in SI Section 4.1):

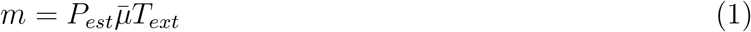

Equation 1 arises from the simple statement that the number of large clones must be equal to their establishment rate 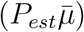 multiplied by their average time to extinction (*T*_*ext*_). This relationship successfully predicts the number of bacterial clones at steady state across a wide range of parameters and a wide range of diversity values (Figure 2B). Through approximations (SI Section 4.2), we find that diversity depends on a single combined parameter to the power 1/3 (Figure 2B, equation 2), and this parameter is proportional to spacer effectiveness *e*, the probability of bacterial survival followed by spacer acquisition (1 − *p*_*V*_)*η*, and the phage mutation rate *µL*.

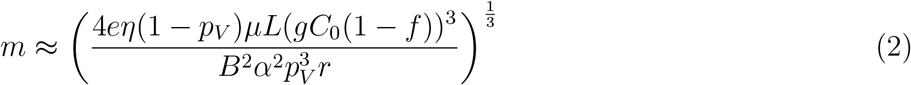

Each of these parameters intuitively increases diversity (for example, a higher phage mutation rate means that phage diversity increases and bacterial diversity follows suit). What is surprising is that their combined effect on diversity is to a power much less than 1: this 1/3 exponent means that if the mutation rate increased ten-fold, the diversity would only increase by about a factor of two (SI Figure 88). In contrast, a simple neutral model of cell division with mutations gives a linearly proportional increase in diversity for the same increase in mutation rate (SI Section 5.4 and SI Figure 89).

To understand where the dependence of diversity on these key parameters come from, we look more closely at each component expression. The effective phage mutation rate 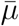 depends linearly on the parameter 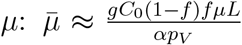, while both the probability of establishment and the time to extinction depend inversely on diversity 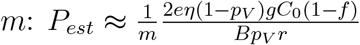 (equation 5) and 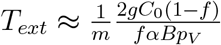 (equation 9). By comparison with equation 1, we find that *m*^3^ depends approximately linearly on mutation rate, resulting in the weak 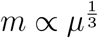 dependence on mutation rate.

The dependence of diversity on both *e* and *η* comes from the probability of phage establishment, since 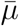 depends only very weakly on these parameters through its dependence on total population sizes and *T*_*ext*_ depends explicitly on *m* alone, not *e* or *η*. The phage probability of establishment is proportional to 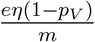 (equation 5), and as before, this gives *m*^3^ ∝ *eη*(1 − *p*_*V*_). Bacteria are more successful at high *η*(1 − *p*_*V*_)*e*, which increases the phage establishment probability. Previous theoretical work has predicted that diversity increases as spacer acquisition rate increases [40]; here we provide a quantitative prediction for this dependence. In the following sections we explore phage establishment and extinction in more detail.

### 3.3 What determines the fitness and establishment of new mutants?

We find that diversity emerges in our model from the balance of phage clone establishment and extinction. However, only some phage mutants escape initial stochastic extinction and survive long enough to become established. What determines the fate of a new phage mutant? In our model, a single phage mutation in a protospacer can completely overcome CRISPR targeting, which means that new phage mutants can infect all bacteria equally well and their initial growth rate *s*_0_ is independent of CRISPR: *s*_0_ ≈ *αn*_*B*_(*B*_*pV*_ − 1) − *F*, where *F* is the chemostat flow rate (a shared death rate for phages and bacteria). Surprisingly, however, even once bacteria start to acquire matching spacers, the probability of establishment for new phage mutants is still well-described by theory in which CRISPR targeting only influences total average population sizes (Figure 3C); that is, the specific interaction between a phage and its matching clone can be ignored. Intuitively, this is because phage clones must grow to a certain size before bacteria encounter them enough to begin to acquire spacers, and this size turns out to be large enough to avoid stochastic extinction (SI Figure 17). The probability of phage establishment is 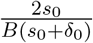, where *δ*_0_ = *F* + *αn*_*B*_(1 − *p*_*V*_) is the initial phage mutant death rate. Importantly, these rates are independent of the population size of *matching* CRISPR bacteria clones; the only dependence on bacteria is on the total bacterial population size *n*_*B*_ (SI Figure 78), which is fairly stable at steady state. Even though CRISPR targeting does not explicitly affect the establishment probability for new phage mutants, the probability of phage establishment increases as the average bacterial immunity increases (Figure 3C). Average immunity is a measure of the overall effectiveness of CRISPR immunity for the entire bacterial population; it is the average of all pairwise immunities between phage clones and bacterial clones weighted by their population sizes (Figure 1A inset). Because higher average immunity is beneficial for bacteria and leads to larger bacterial population sizes (Figure 5C), higher average immunity also means there is stronger selective advantage for new phage mutants that can escape CRISPR targeting.

**Figure 3:**
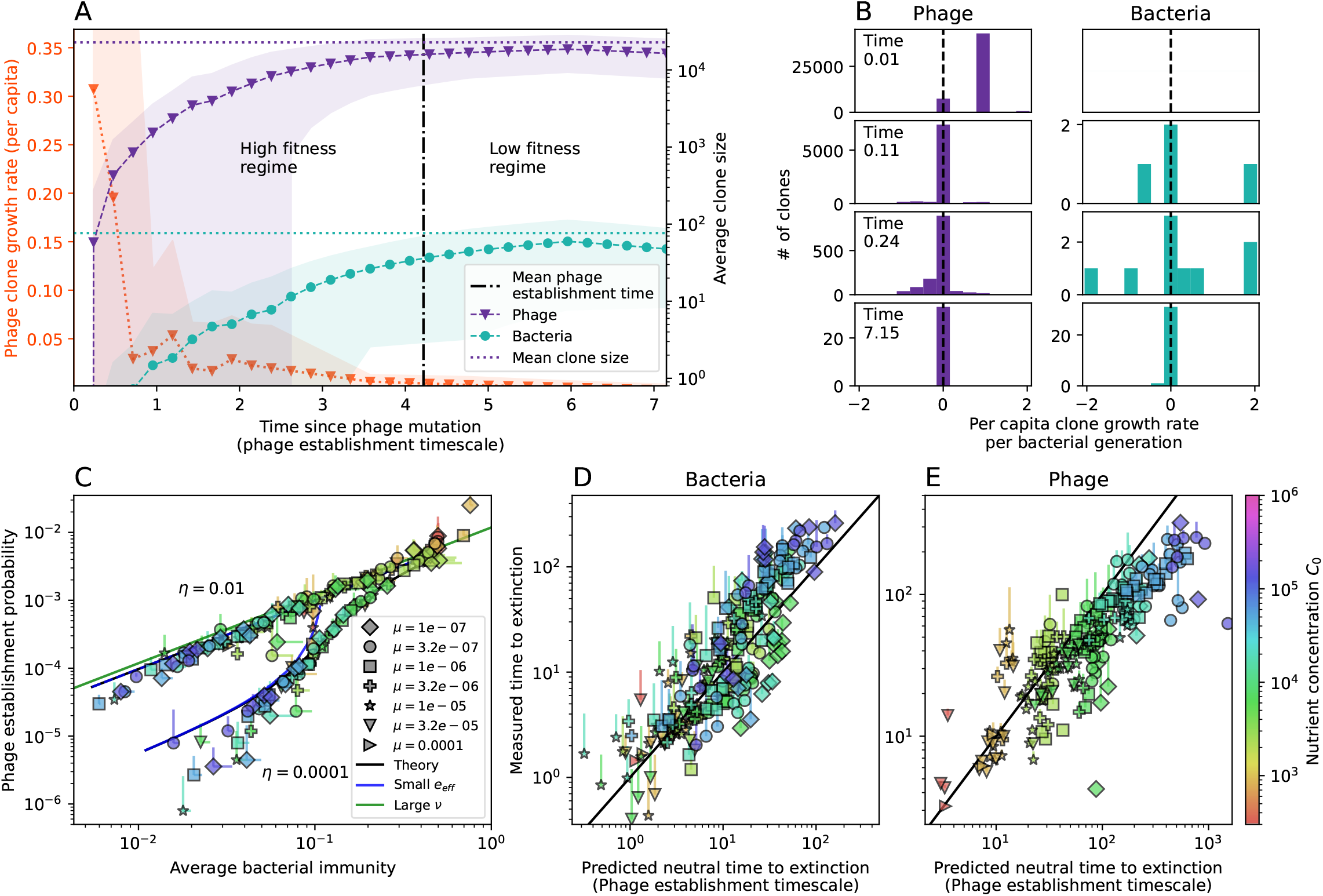
The fate of individual clones. (A) Phage and bacteria coevolve in two timescale-separated regimes characterized by phage clone fitness. Average phage and bacteria clone size vs. time since phage mutation (right axis), and average clone growth rate vs. time since phage mutation (left axis). Markers show the average over all clone trajectories after steady-state from six simulations with the same parameters. (B) Histograms of individual clone fitness grouped by time since phage mutation. Phage clones initially have fitness *>* 0, but rapidly most clones reach neutral growth (fitness 0). Bacteria clones also follow suit, initially having fitness *>* 0 and rapidly reaching 0 fitness on average. Individual clone trajectories are highly variable. (C) Probability of phage clone establishment vs. average immunity. Clones are considered established in simulations when they reach the mean clone size. Equation 3 with *ν* = 1 is shown in green and with *ν* given by equation 4 in blue. (D,E) Mean time to extinction for large bacteria (D) and phage (E) clones as a function of the neutral fitness mean time to extinction prediction. The solid black lines describe approximate analytic expressions for bacteria and phage time to extinction using a neutral fitness assumption. In (A,B), *C*_0_ = 10^4^, *e* = 0.95, *η* = 10^−3^, and *µ* = 3 *×* 10^−6^.

For insight into the nature of this dependence, we approximated the probability of establishment at two extremes of average immunity (see SI Section 4.2.1). The probability of establishment can also be written

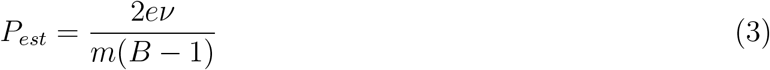

where *ν* is the fraction of bacteria that have spacers and *m* is the average number of bacterial clones at steady-state. Low average immunity occurs when *e*/*m* is small: in this limit, the fraction of bacteria with spacers is

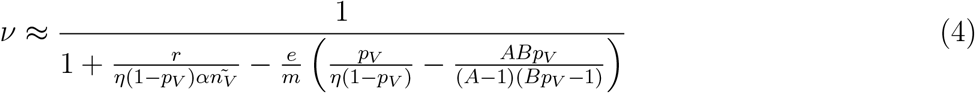

where 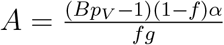 is the extinction threshold for phages (*A >* 1 for phage survival), and 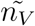 is the phage population size calculated with *e* = 0 (SI equation 49); this is the extreme limit of low average immunity where total population sizes approach their values in the absence of CRISPR immunity (SI Figure 30). The quantity 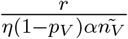 can be understood as the ratio of the rate of spacer loss to the rate of spacer acquisition. If spacer loss is high and acquisition is low, the fraction of bacteria with spacers (*ν*) decreases. This in turn correlates with a lower probability of phage establishment (equation 3), since phages are less likely to establish when there is less pressure from bacteria defending against them with CRISPR. Equation 4 is plotted in Figure 3C in blue. At very high average immunity, on the other hand, *ν* becomes close to 1 regardless of *η* (SI Figure 37), which is why the curves in Figure 3C overlap at high average immunity (*ν* = 1 is plotted in green).

For more insight into the parameter dependence of *P*_*est*_, we apply the same approximation as for *m*, expanding *P*_*est*_ in 1/*B* and *η* (SI Section 4.2.5):

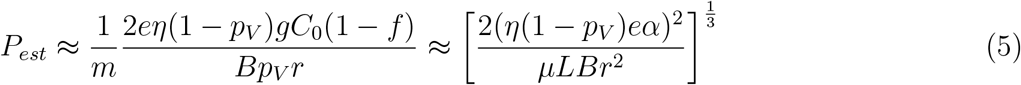

The probability of establishment increases with the probability of bacterial escape and spacer acquisition *η*(1 − *p*_*V*_) and with spacer effectiveness *e*, but decreases with increasing mutation rate *µL*. This is consistent with the intuition that more successful bacteria increase the strength of selection for phage mutants (higher *η* and *e*), but that higher mutation rate reduces the probability of establishment for any particular mutant.

### 3.4 Large bacteria and phage clones undergo neutral extinction dynamics

Once phage clones become large, bacteria acquire matching spacers and each large phage clone has a matching immune bacterial clone. We expected to find that clone dynamics would be strongly coupled due to the CRISPR immune response, but instead we find that at steady state both bacteria and phage clones that grow large behave with neutral dynamics on average. This is because there is a certain average clone size dictated by birth and death rates; once clones reach the average clone size, their birth and death rates are matched and their fitness is zero on average (see SI Sections 5.1.1 and 5.3.4).

A consequence of this neutral behaviour is that a neutral prediction for the mean time to extinction matches our simulation results well for both bacteria and phage clones (Figure 3D-E). As in the case of establishment, the constant fitness assumption breaks down once clones deviate from the mean clone size, but it turns out that this “restoring force” does not significantly affect the extinction times for large clones (SI Figure 82). Approximating the neutral extinction time for phage and bacteria at low average immunity gives an extinction time proportional to 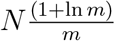 (equations 6 and 7, details in SI Sections 5.1.2 and 4.2.2), where *m* is the mean number of bacterial clones at steady state and *N* is the total number of phage or bacteria. For both bacteria and phage, the dominant dependence is on the mean clone size *N*/*m*: larger clones take longer to go extinct (SI Figures 84 and 87).

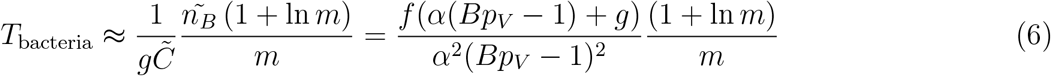

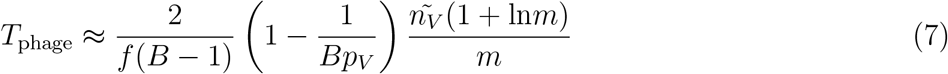

The tilde in equations 6 and 7 (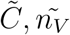 and 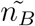) represents a population quantity calculated with *e* = 0, low average immunity where total population sizes approach their values in the absence of CRISPR immunity.

These simplified expressions for time to extinction give insight into the main drivers of extinction. The phage time to extinction decreases as the outflow rate *f* increases — if phages die at a higher rate, they go extinct more quickly. Interestingly, a larger burst size *B* decreases the time to extinction for large phage clones, consistent with an overall increase in the size of fluctuations as the burst size increases. Similarly but perhaps counterintuitively, the bacteria clone time to extinction decreases as the growth rate 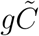 increases — a higher growth rate means faster dynamics overall and lower time to extinction. Note however that a higher growth rate (for instance with larger *C*_0_) also leads to larger *n*_*B*_.

These expressions still depend on *m*, which is an emergent quantity dependent on all other parameters as well. We apply the approximation for *m* from SI Section 4.2.5 for more insight.

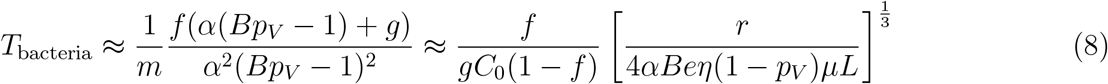

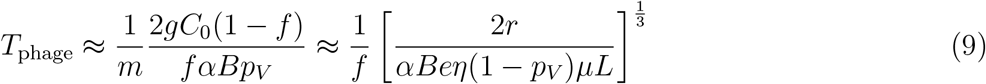

Both phage and bacteria extinction depend inversely on *α, B, e, µL*, and *η*(1 − *p*_*V*_), with the time to extinction decreasing with all parameters except the spacer loss rate *r*. The bacteria time to extinction increases with increasing flow rate *f*, while the phage time to extinction decreases with *f*.

### 3.5 Cross-reactivity leads to dynamically unique evolutionary states

We next asked how cross-reactivity between spacer and protospacer types impacts population dynamics and outcomes. Adding cross-reactivity was motivated by several experimental observations in CRISPR immunity: (1) In Type I and Type II CRISPR systems, single mutations in the PAM or protospacer seed regions (approximately 8 nucleotides at the start of a protospacer) can facilitate phage escape, whereas mutations elsewhere in the protospacer are tolerated by the CRISPR system [76, 34]. Even when a phage manages to escape direct targeting, in Type I and II systems an imperfect spacer match can facilitate priming: when Cas machinery binds to a protospacer match, even if unable to cleave the target, the likelihood of acquiring a nearby spacer is increased [46, 77, 34, 78]. (2) Type III CRISPR targeting in *Staphylococcus epidermidis* has been shown to be completely tolerant to all single and even double mutations, meaning that these systems are naturally cross-reactive [37].

We simulated two types of cross-reactivity: one in which phages experience an exponential decrease in CRISPR effectiveness with mutational distance, and a step-function cross-reactivity where phages require an additional 1 ≤ *θ* ≤ 3 mutations to perfectly escape CRISPR targeting. Exponential crossreactivity has been modeled extensively in vertebrate adaptive immunity [79, 4, 13, 7, 14, 15], and our definition of exponential cross-reactivity as a function of mutational distance is the same as in ref. [7]. Step-function cross-reactivity, on the other hand, is reminiscent of the Type III CRISPR system in which multiple point mutations are required to escape CRISPR targeting [37] and was also modelled theoretically by Han *et al*. [41].

We find that cross-reactivity results in strikingly different dynamics of clone establishment and persistence (Figure 4 and SI section 4.4), including a traveling wave regime in which genetically neighbouring clones “pull” each other along (Figure 4B-D), giving way to a regime in which new mutants have very low establishment rates in the case of step-function cross-reactivity (Figure 4C). When cross-reactivity is high and phages require multiple mutations to escape targeting, we expect new phage mutants to have a lower fitness on average because they may already be within the immunity range of existing bacterial clones. However, unlike without cross-reactivity, not all new mutant fitnesses are the same because fitness now depends on the distribution of matching spacers in the bacterial population. Cross-reactivity adds a ‘direction’ in the genetic landscape and a fitness gradient for new phage mutants, leading to a series of rapid establishments and a traveling-wave regime. This is true for both types of cross-reactivity we studied, but we also observed striking qualitative differences between exponential and step-function cross-reactivity. In the former, all phage mutants are guaranteed a slight fitness advantage, and the traveling wave appears right at the start of simulations, with occasional lineage-splitting events (Figure 4B). In contrast, step-function cross-reactivity means that mutants within the cross-reactivity radius have no fitness advantage at all. In this case, there is some variable initial length of time required to establish the traveling wave pattern: at the start of a simulation, all mutants are within the crossreactivity radius and evolve purely neutrally. At least *n* = *θ* new mutants must establish through neutral dynamics before the next mutant can escape CRISPR targeting and grow with high fitness; this leads to the traveling-wave regime appearing later for *θ* = 2 than for *θ* = 1, and sometimes never appearing at all. Once the traveling wave appears, the dominant phage and bacteria clone types are exactly offset by *θ*: the most abundant phage clone will in general be *θ* mutations away from the most abundant concurrent bacteria clone. This is directly visible in the clone trajectories in Figure 4C-D inset and SI Figure 63: matching-colour bacteria and phage clones are offset in time. Another regime is also possible in the case of step-function cross-reactivity: if multiple large clones establish that are each outside of all the others’ cross-reactivity radii (SI Figure 72), then new mutants all have zero fitness and the traveling wave comes to an abrupt halt (around 7000 generations in Figure 4C). Establishment of new clones is once again extremely rare, and the existing large clones persist for a long time (SI Figures 62 and 71).

**Figure 4:**
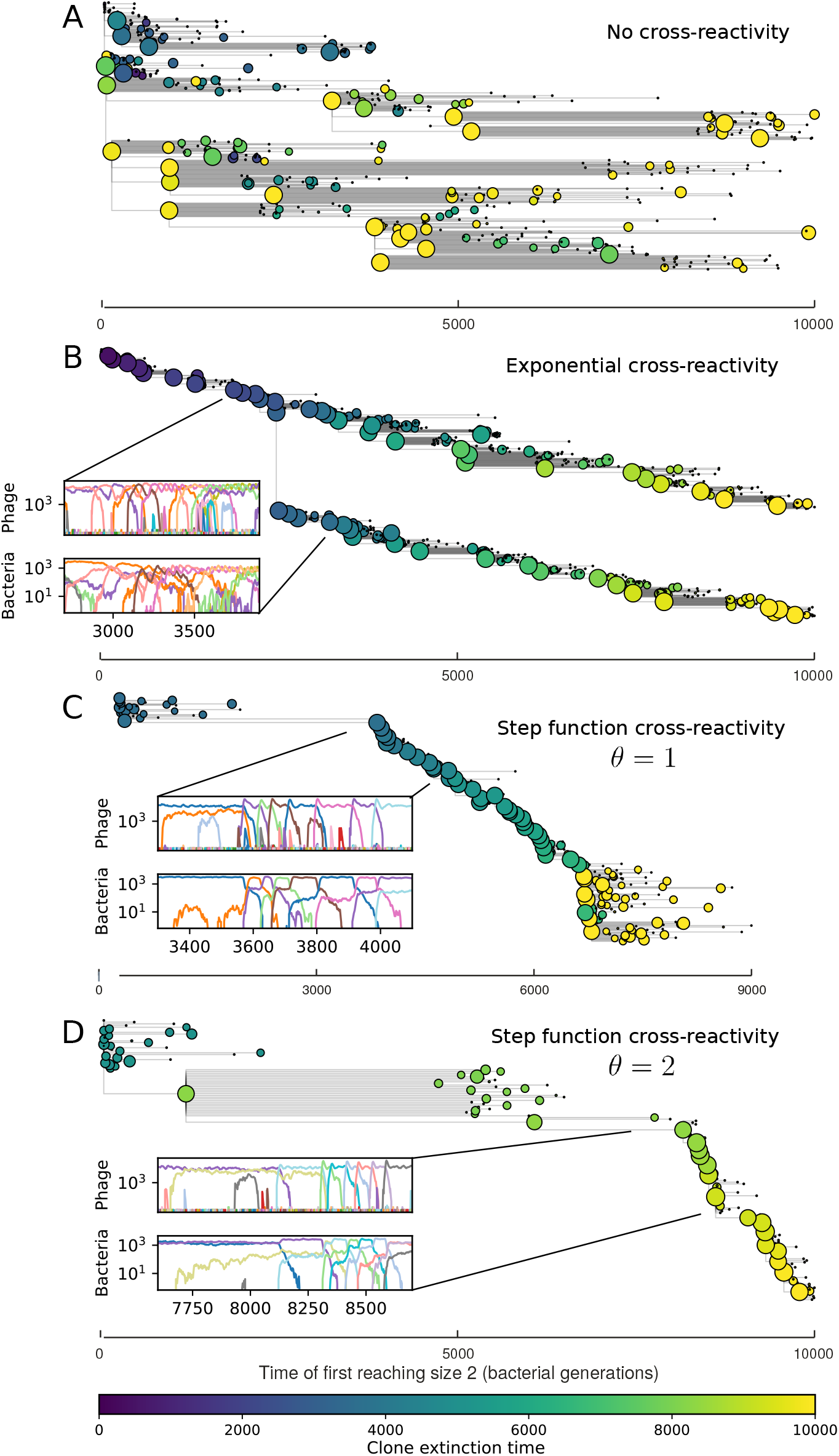
Cross-reactivity leads to ‘spindly’ phylogenies and regime switching. Phage clone phylogenies for four simulations with different cross-reactivities: no cross-reactivity (A), exponential cross-reactivity with *d* = 4 (B), and step-function cross-reactivity with *θ* = 1 (C) and *θ* = 2 (D). All simulations share all other parameters: *C*_0_ = 10^4^, *η* = 10^−4^, *µ* = 10^−6^, *e* = 0.95. Phage clones are plotted at the first time they pass a population size of 2 (to remove clutter from many new mutations destined for extinction), and the size of each circle is logarithmically proportional to the maximum size reached by that clone. Colours indicate the time of extinction of each clone. For each simulation with cross-reactivity, the left inset shows phage (top) and bacteria (bottom) clone sizes over time; colours indicate unique clone identities.

Changes to the fitness landscape of individual clones caused by cross-reactivity also influence population-level outcomes such as diversity. In general we find that for high cross-reactivity, average diversity is lower than predicted for simulations without cross-reactivity because the probability of phage establishment per mutant decreases (Figure 5A). A decrease in phage mutant fitness as the strength of cross-reactivity increases is in line with the dependence of fitness on cross-reactivity reported by refs. [7] and [80]. Even with more complicated underlying dynamics, though, measuring average immunity alone is enough to reproduce total population sizes using our simple deterministic equations developed in the limit of no cross-reactivity (Figure 5B-C). This is because average immunity completely captures the population-level impact of CRISPR immunity, and we can replace the CRISPR effectiveness parameter *e* with measured average immunity to get very good agreement between measured and predicted population sizes even away from steady-state (SI Figure 61).

**Figure 5:**
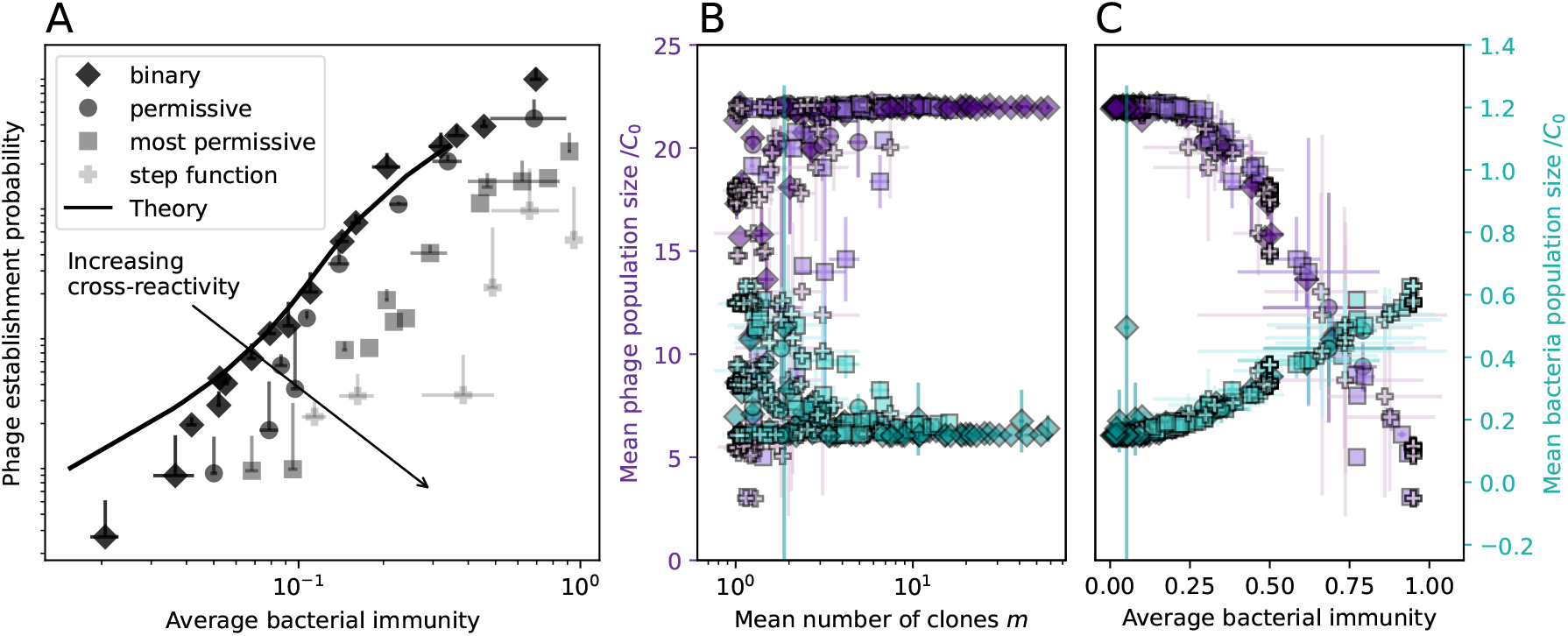
Average immunity underlies population outcomes. (A) Probability of phage clone establishment vs. average immunity for different amounts and types of cross-reactivity. Markers and shading indicate type and level of cross-reactivity. Simulation averages are shown for *η* = 10^−4^ and *µ* = 10^−6^. (B,C) Total phage (purple, left axis) and total bacteria (teal, right axis) average population sizes vs. the mean number of bacterial clones *m* (B) and vs. average bacterial immunity (C). Each point is an average at steady-state over three or more independent simulations with the same parameters. Total sizes are scaled by the initial nutrient concentration *C*_0_. Lighter colours indicate stronger cross-reactivity, marker shapes match legend in (A).

### 3.6 Dynamics are determined by diversity

How quickly does the phage population evolve? We model protospacers as length 30 binary sequences (a DNA alphabet with two letters) and mutations as random bit flips in these sequences. With these explicit spacer and protospacer sequences, we quantify the speed of evolution as the ‘mutational distance’ per generation [80]: between two time points, how far in genome space has the phage population traveled, or how many mutations have occurred on average (SI section 4.3)?

The speed of evolution is both stable and repeatable between simulations at steady-state and highly correlated between bacteria and phage (SI Figure 44). We find that the speed of evolution is inversely proportional to the time to extinction for large phage clones (Figure 6B, SI section 4.3.1). Intuitively, if phage clones turn over more quickly (small time to extinction), the population is able to move more quickly to a different genetic state, while if the time to extinction is large, new mutants are seeded close to parent populations that persist for a long time, limiting the mutational distance. Since the phage time to extinction has a simple relationship to bacterial diversity, we can also relate speed to diversity, and we find that the speed of evolution is proportional to diversity and proportional to the same parameter combination to the power 1/3 (Figure 6C). As in the case of diversity, increasing phage mutation rate *µ* increases the speed of evolution sub-linearly.

**Figure 6:**
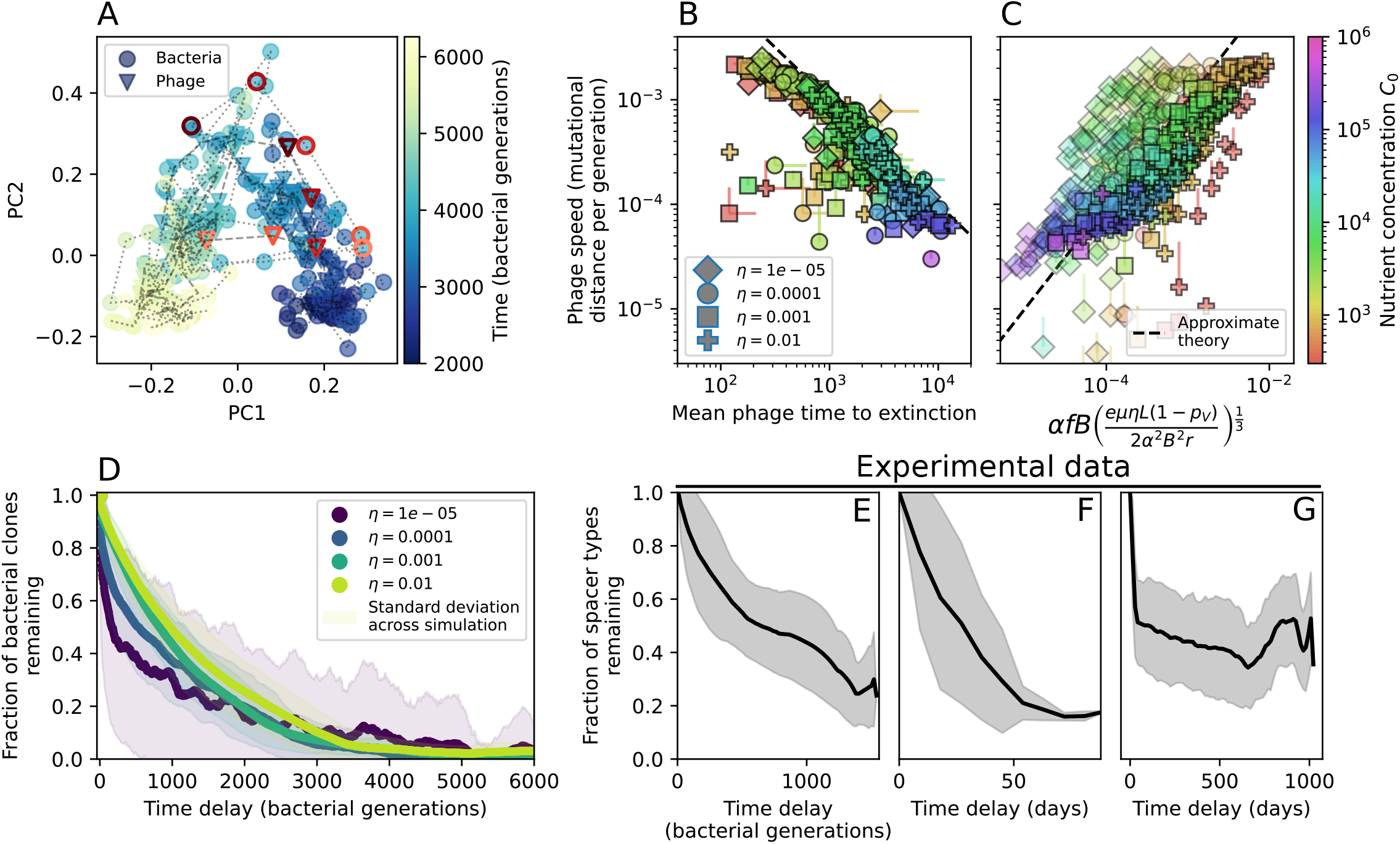
Phage evolution and spacer turnover. (A) PCA decomposition of phage and bacteria clone abundances for a simulation with *C*_0_ = 10^4^, *e* = 0.95, *η* = 10^−4^, and *µ* = 10^−5^. Clone abundances are normalized at each time point, then PCA is performed for the entire phage time series over*≈* 4000 generations (4 times the mean extinction time for phage clones). Bacteria and phage clone abundances are transformed into the PCA coordinates; colours indicate simulation time. Five time points are highlighted in progressively lighter shades of red for emphasis. (B) Phage genomic speed of evolution vs. mean large phage clone time to extinction. The phage speed is the weighted average genomic distance between the phage population at the end of the simulation and the phage population at an earlier time, divided by the time interval. The dashed line is 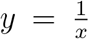. (C) The speed of evolution increases as spacer effectiveness *e*, spacer acquisition probability *η*, and phage mutation rate *µ* increase. The dashed line shows an approximate theoretical calculation (assuming speed = 1/time to extinction) which captures the trend across a wide range of parameters. (D) Spacer turnover as a function of time delay for four simulations with *C*_0_ = 10^4^, *e* = 0.95, and *µ* = 10^−5^. The fraction of bacterial clones remaining is the fraction of clones that were present at time *t* that are still present at time *t*+ delay. Solid lines are an average across steady-state for each value of the time delay; shaded regions are the standard deviation. (E-G) Spacer type turnover calculated as in (D) using experimental data from [24] (E), metagenomic data sampled from groundwater from [56] (F), and metagenomic data sampled from a wastewater treatment plant from [39] (G). Experimental time points are interpolated to the minimum sampling interval to allow averaging across the experiment.

Our simulations show continual spacer turnover at steady-state (Figure 6D), a feature that we might also expect to see in actively evolving laboratory or natural populations of bacteria and phages. We analyzed data from a long-term in vitro coevolution experiment with *Streptococcus thermophilus* bacteria and phage [24], data from a time-series sampling of a natural aquifer community of bacteria [56], and data from a time-series sampling of a wastewater treatment plant [39] (see Methods) and calculated spacer turnover over time (Figure 6E-G). We found that spacer sequences experienced large amounts of turnover in both cases, indicating ongoing change in the spacer content in these populations. Even if spacer loss is happening for non-selective reasons (for example mixing or flow in the aquifer system), turnover indicates that new spacers are being acquired as well and that CRISPR systems are active. Moreover, the timescale of spacer turnover in the *S. thermophilus* experiment is similar to the timescale in our simulations – we modeled all simulation parameters on known parameters for *S. thermophilus* where possible.

### 3.7 Time-shifted average immunity calculated from data reveals distinctive patterns of turnover

Bacteria acquire spacers in response to phage clones becoming large. The spacer composition of the bacterial population tracks the phage protospacer composition with a lag: Figure 6A shows the first two components of a PCA decomposition of bacteria and phage abundances over time in a simulation, a visual illustration of bacterial tracking. Without cross-reactivity, trajectories in this lower-dimensional space do not travel in a straight line; they are reminiscent of the diffusive coevolutionary trajectories in antigenic space described in a theoretical model of vertebrate virus-host coevolution [13]. In the traveling wave regime with cross-reactivity, trajectories are much more ballistic (SI Figure 47). Can we quantify how well and how quickly can bacteria track the evolving phage population with CRISPR?

Average immunity is a simple metric that quantifies the overlap between bacteria and phages. In experiments with both microbes and vertebrates, time-shift infectivity analyses between host and pathogen populations typically show that hosts are more immune to pathogens from the past and less immune to pathogens from the future [81, 82, 83, 84, 85, 86, 27, 87]. We conduct a time-shift analysis on our simulation data (Figure 7A-B) and find that the same pattern holds true, but only for a limited time window: bacteria are indeed more immune to phages from the past, but this past immunity has a peak and then decays as we look further into the past, eventually reaching 0 immunity (Figure 7B). The presence of a peak in past immunity reflects the timescale of spacer turnover: once the bacterial population has lost all spacers from a previous time point, it is no longer immune to contemporaneous phages and time-shifted average immunity falls to 0. In simulations, the position of this peak is dependent on parameters such as *η* and *µ* (Figure 7E-F). As *µ* increases, the peak occurs further in the past since phage are now moving more quickly away from bacteria tracking, while as *η* increases, the peak moves closer to the present since bacteria are responding more quickly to changes in the phage population. This suggests that there is a tradeoff between immune memory durability and responding quickly to immune threats: bacteria must choose between tracking the phage population closely or keeping past immunity for a long time.

**Figure 7:**
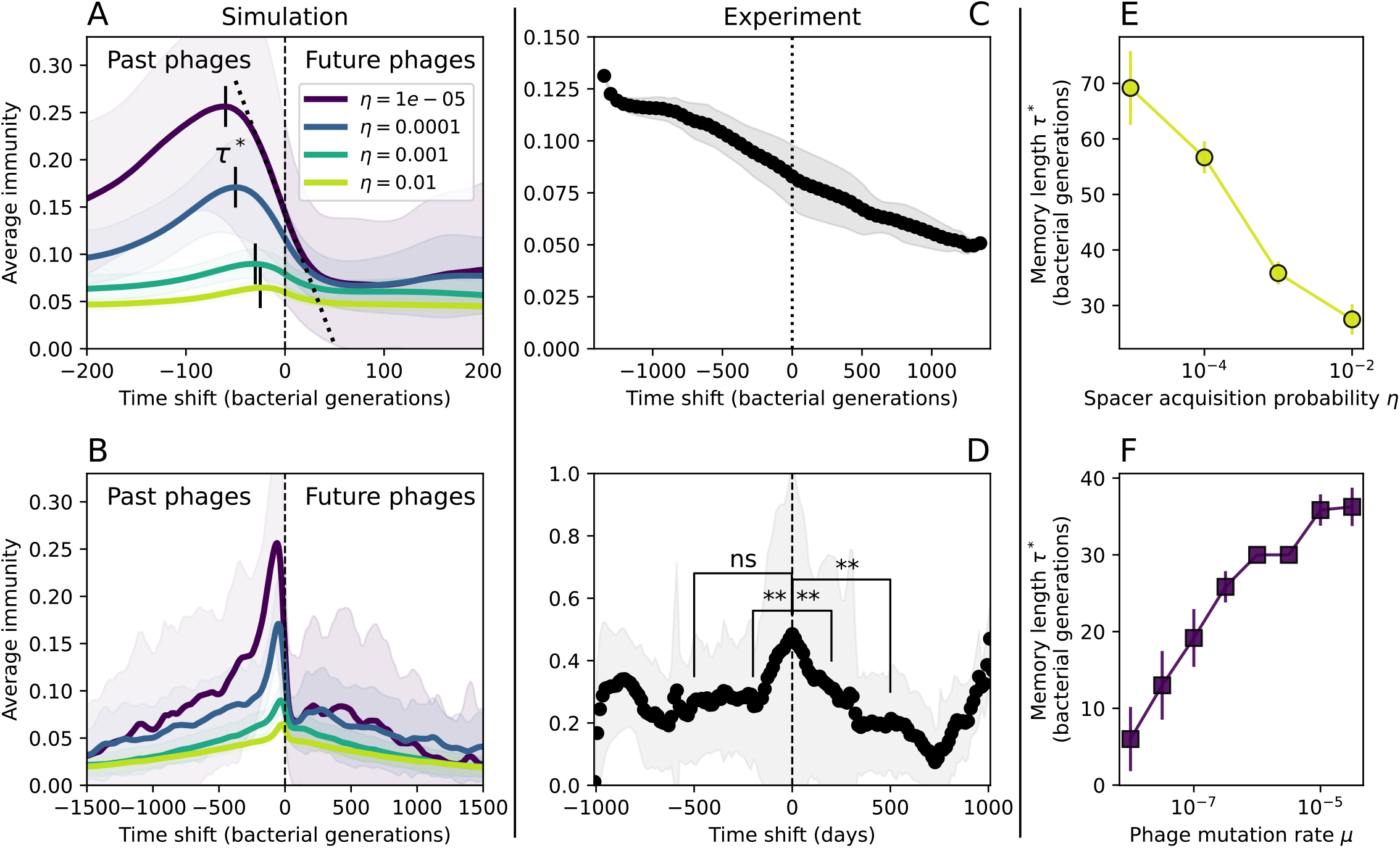
Quantifying immune memory in data. (A,B) Average immunity of bacteria against phage for four simulations with different values of *η* as a function of time shift. Solid lines are an average across steady-state for each value of the time shift; shaded regions are the standard deviation. Average immunity peaks in the recent past (A, indicated by *τ* ^*∗*^) with a negative slope through zero delay (A, black dashed line) and decays to zero at long delays in the past or future (B). For all simulations *C*_0_ = 10^4^, *µ* = 10^−5^, and *e* = 0.95. (C,D) Average overlap between bacterial spacer and phage protospacer types using data from a lab experiment with *S. thermophilus* and phage from ref. [24] (C) and data from a wastewater treatment plant sampled over three years from ref. [39] (D). Spacer types are grouped by 85% similarity, and shaded region is standard deviation across averaged data. Base average immunity values were multiplied by the average number of protospacers corresponding to the *S. thermophilus* CRISPR system (C) and the *Gordonia* CRISPR systems (D) to account for multiple potential protospacer targets per phage. In (D), we compared two time shifts with zero delay average immunity using a Wilcoxon signed-rank test: *p* = 0.27 for lower past immunity at 500 days, *p* = 0.008 for lower past immunity at 200 days, *p* = 0.001 for lower future immunity at 500 days, and *p* = 0.003 for lower future immunity at 200 days. (E,F) The position of the peak in past immunity for simulated data vs. spacer acquisition probability *η* (E) and phage mutation rate *µ* (F). The peak position is the time shift value for which the curves in (A) are largest, indicated by *τ*\*.

Time shift analyses are usually performed explicitly by directly combining stored samples from different time points [86, 27, 87]. However, by sequencing CRISPR spacers and phage genomes, a pseudo-analysis can be done without any time-shifted competition experiments by calculating the overlap between bacterial spacers and phage protospacers at different times delays. We performed such an analysis for two published datasets: a long-term laboratory co-evoluation experiment with *S. thermophilus* and phage [24] and time series of metagenomic samples over three years from a wastewater treatment plant [39]. In both experiments, whole-genome shotgun sequencing was performed on bacteria and phage DNA, and CRISPR spacers and protospacers were recovered and reported. We re-analyzed these datasets to detect CRISPR spacers by finding sequences adjacent to known CRISPR repeats in raw reads. We also detected protospacers by finding matches to our detected CRISPR spacers in reads that did not match the CRISPR repeats or the bacterial reference genome(s) (see Methods and SI Section 6.2 and 6.3).

We grouped spacers with an 85% similarity threshold and interpolated counts between sequenced time points. We calculated the average overlap as a function of time delay, averaging over all combinations of interpolated data with the same time delay (Figure 7C-D). We found two qualitatively different trends as a function of time delay. In the laboratory coevolution experiment, we found that bacteria are more immune to past phages and less immune to future phages (Figure 7C), consistent with what we see at short time delay in our model (Figure 7A, black dashed line). In contrast, in the wastewater treatment plant data, we found a peak in average immunity that is roughly centred at zero time delay: bacteria are most immune to phages from their same time point, and immunity rapidly decays in both the past and the future (Figure 7D), qualitatively similar to our simulation results on very long timescales (Figure 7B). These different trends may reflect different regimes of CRISPR immune memory: in the laboratory experiment data we see no evidence of the decay of immune memory, while in the wastewater treatment data we see a suggestion of decay on the timescale of weeks. Bacteria appear to track the phage population closely in the wastewater data, while in the laboratory data bacteria lag behind the phage population.

Because the variability in average immunity between time points was very high in the wastewater treatment plant data, we performed a Wilcoxon signed-rank test between the average immunity at zero time delay and the average immunity at a time delay ±200 and ±500 (see Methods and SI Section 6.3.1 for details). We found that past immunity after 200 days is significantly lower than present (*Z* = 1241, *p* = 0.008), but that immunity after 500 days is not significantly lower than present (*Z* = 392, *p* = 0.27). For both time delays, we found that future immunity is significantly lower than present (*Z* = 580, *p* = 0.0012 for 500 days and *Z* = 1293, *p* = 0.003 for 200 days). Interestingly, these significance values are not symmetric if we pose the question from the perspective of phage: the overlap between phage and future bacteria at 500 days is lower than the overlap for present phages (*Z* = 507, *p* = 0.024), and the overlap between phage and past bacteria is lower than the overlap at present (*Z* = 388, *p* = 0.027, SI Figure 118). This asymmetry, that bacteria are generally more immune to all past phages while phage are not more infective against all past bacteria, is qualitatively the same as that reported by Dewald-Wang *et al*. in an explicit time shift study of bacteria and phage immunity and infectivity in chestnut trees [87].

## 4 Discussion

Many bacteria and archaea possess CRISPR systems, and a significant fraction of these systems are likely to provide immunity against phages [88, 89, 90]. Given that bacteria and phages coexist in natural environments over extremely long timescales, the impact of CRISPR immunity in these steady-state conditions has remained under-explored. We constructed a phenomenological model of CRISPR immunity in a bacterial population interacting with phages to explore the impacts of adaptive immunity on population survival, fitness, and diversity. We found that both phage and bacterial genetic diversity emerged spontaneously with a minimal set of interactions, and we derived approximate analytic predictions for population outcomes. These rigorous analyses of our simple model lay a foundation for theoretical analysis of adaptive immunity in host-pathogen systems.

Our model is mechanistically simple, and we left out many known biological features and interactions by choice in order to gain a deep understanding of the factors that influenced our results. We modeled uniform spacer effectiveness, uniform spacer acquisition probability, and a constant phage mutation rate, all in a well-mixed system. This constitutes a null model of CRISPR immunity that provides a useful comparison point for both more accurate mechanistic models and experimental data. CRISPR is one of many other bacterial antiviral defense systems [91], and within the CRISPR world, CRISPR systems are highly evolutionarily diverse and there are many known differences in function and effect between CRISPR systems of different bacterial species [92, 93, 94, 95]. Experiments typically study a particular CRISPR system, and it is unclear which revealed mechanisms are specific to that system or are a more general property of CRISPR systems. For example, the spacer acquisition rate has been demonstrated to vary across particular phage sequences in several experiments [96, 97, 23] and is the source of differences in spacer abundance in some experiments [96], but whether this is a general principle that causes broad abundance distributions is not known. Many theoretical works also include mechanistic details such as a lag between infection and burst [36, 57], multiple protospacers and spacers (typically with a fixed upper bound) [43, 98, 59, 40, 50, 58], spatial structure [61, 70], autoimmunity [58, 29], and fitness costs of immunity and/or escape mutations [43, 99]. Many phages also contain anti-CRISPRs which impact phage evolution and CRISPR immunity [26, 100]. These details are all biologically important, but stripping them away as we do provides great insight into which population features depend on these details and which may be more general properties of adaptive immunity.

### Pathogen and host diversity cannot be separated

We showed that immune interactions without cross-reactivity produce the counterintuitive result that more diversity is beneficial for phages but not for bacteria, in contrast to several experimental results that show that more bacterial diversity increases the likelihood of phage extinction [25, 28] and decreases the ability of phages to adapt [33]. This discrepancy occurs because bacteria and phage diversity is decoupled in these experiments. Experimental manipulations of CRISPR diversity have focused on changing the number of bacterial spacers present in the population [25, 33, 28] while keeping phage diversity fixed (and low), with the exception of Guillemet *et al*. [31] who also explored high vs. low phage diversity. In contrast, emergent phage and bacterial diversity were tightly coupled in our model: if bacterial diversity was high, phage diversity was also high. At high diversity, bacteria must “choose” which of many phage clones to gain immunity against, meaning that they are then immune to a smaller fraction of the total phage population compared to when diversity is low. This appears at first glance to be a result of limiting bacteria and phage to a single spacer or protospacer, but the trend also holds when there are multiple spacers or protospacers provided phage and bacterial diversity are correlated (see SI Section 6.1.1). The intuition is this: for a fixed number of protospacers and fixed CRISPR array length, if diversity of both bacteria and phage increase beyond the length of the CRISPR array, bacteria can’t be immune to all phage strains at once and average immunity goes down as diversity increases. The actual benefit of bacterial diversity is thus context-dependent, and manipulations of diversity should be understood in terms of their impact on average immunity. This has implications for understanding the so-called “dilution effect” which is typically described as a decrease in the fraction of susceptible hosts as host diversity increases [53, 28]. Our model and analysis show that a dilution effect can only occur if host diversity increases out of proportion to pathogen diversity, whereas if both host and pathogen diversity increase in tandem, the dilution effect actually changes direction, decreasing the fraction of pathogens that a host may be immune to as pathogen diversity increases. In the same vein, previous theoretical work has found that CRISPR with a fitness cost would be selected against when viral diversity is high for this reason [43, 98]. Our results, building on existing theory, show that phage and bacterial diversity must be explored in tandem and that CRISPR immunity may provide less benefit when phage diversity is high.

The discrepancy in the relationship between average immunity and diversity between our model and experiments suggests that diversity itself may not be the most informative observable feature, but that average immunity is what actually determines population outcomes. Indeed, when we introduced cross-reactivity between spacer sequences, we found that diversity decreased and that other processes such as establishment were impacted, but that average immunity still correctly predicted population sizes. Average immunity completely summarizes the effect of CRISPR immunity and accurately predicts population outcomes. Experiments that manipulate bacterial diversity only change average immunity if they include a changing proportion of sensitive bacteria as in Common *et al*. [28].

Our results showed complex outcomes and a wide range of overall spacer diversity despite limiting bacteria and phage to a single protospacer and spacer. However, we also observed rapid loss of previous spacer types as bacteria returned to the naive state before acquiring new spacers. In reality, bacteria can store multiple spacers in their CRISPR arrays [101, 90] and phages can have several hundred to several thousand protospacer sequences which are determined by a particular protospacer-adjacent motif (PAM) in type I and II CRISPR systems [102]. In *S. thermophilus* phage 2972 for example, there are about 230 protospacers that can be acquired into the CRISPR1 locus of *S. thermophilus* [23] and 465 that can be acquired into the CRISPR3 locus. Similarly, many bacteria can store tens to hundreds of spacers in their CRISPR arrays [101, 103, 104, 90]. Our single-spacer assumption is reasonable for the experimental systems we consider. In liquid culture experiments with staphylococci, most bacteria acquire a single spacer [105, 96, 34], and in short-term experiments with *S. thermophilus*, most bacteria acquired just one CRISPR spacer over two weeks [23] or one to three spacers over 9 days [27]. In the experimental data we analyzed here (a 200+ day experiment), the average number of newly acquired spacers hovered around 5 throughout the experiment [24]. In a long-term study of metagenomic data, most spacer matches to extant viral sequences were located near the leader end of CRISPR loci; conserved trailer-end spacers had very few matches to sampled viruses [35]. All these results suggest that dynamics may be dominated by just a handful of spacers per bacterium. In addition, theoretical work has found that the most recently-acquired spacer provides more immune benefit than older spacers [40, 41]. Our single-protospacer assumption for phages had a more extreme impact, however: in our model, we assumed in the non-cross-reactive case that a single point mutation allowed phages to escape from all CRISPR targeting, when in reality a phage with hundreds of protospacers may need mutations in many different protospacers to fully escape from the bacterial population. Previous work on the impact of CRISPR diversity takes place in this context: that having more diverse bacterial spacers means phages need mutations in multiple protospacers to escape, and that a single mutation only escapes a single bacterial spacer at most, leaving other bacteria still resistant [53, 25, 28].

We explored the impact of phages containing multiple protospacers on our qualitative diversity results and found that we can account for multiple protospacers in the framework of average immunity. At one theoretical extreme, the phage population is clonal, there are no mutations, and all protospacers are present in the same genome. In this limit, average immunity reaches its maximum value of 1: all bacteria with spacers will have a spacer that matches the entire phage population. A more realistic scenario is somewhere between this limit and the single-protospacer limit we explore in our model: phages contain multiple protospacers, but mutations mean that multiple phage clones are present in the population. We explored this middle ground with a toy model. For a given synthetic set of observed spacers and protospacers, we grouped protospacers randomly together into phage clones, then asked how average immunity changed as the number of protospacers per phage increased. If bacteria are restricted to a single spacer, the solution is simple: multiplying average immunity by the average number of protospacers per phage accurately reproduces the true average immunity. We extended this to multiple spacers as well: if bacteria have *n* spacers on average, average immunity can be calculated using a simple relationship between the average spacer array length *a*_*n*_ and the base average immunity assuming one spacer per bacterium (*a*_1_): *a*_*n*_ = 1 − (1 − *a*_1_)^*n*^ (SI Section 6.1). An interesting result made clear by this analysis is that the relative immune benefit from gaining a spacer decreases as immunity increases: a CRISPR array twice as long does not provide double the immunity. The same effect was observed by Iranzo *et al*. [98] in a model of CRISPR immunity with unconstrained array lengths. A similar diminishing-returns result has also been observed in theoretical models of vertebrate immunity where the relative decrease in immune susceptibility and increase in immune memory both decrease as the number of infections increases over an organism’s lifetime [106]. Long CRISPR arrays are also subject to a dilution effect: spacers compete to form complexes with limited Cas protein machinery, and if the number of spacers is high, there is a higher chance that the needed spacer is not available in high enough numbers during an infection [107, 104]. The simple relationship between average immunity and longer spacer and protospacer arrays means that extending our population-level results to more physiological regimes of spacer and protospacer number is straightforward.

We have showed that we can relax our single spacer and protospacer assumptions and understand the impact on average immunity under simple assumptions about array arrangement. However, it remains unknown how the linkage of different spacers and protospacers within genomes would affect individual clone dynamics, and this is an interesting area for further study. Our work effectively assumes that all spacers and protospacers evolve independently and are uncoupled, but in real genomes evolving with CRISPR, spacers and protospacers may hitchhike to prominence through selection acting on a different sequence in the same genome. Models of CRISPR locus spacer addition have shown that trailer-end clonality emerges from selective sweeps of newly-added spacers [35, 41], and hitchhiking will affect interpretations that can be drawn from the abundance of individual spacer clones. Exploring correlations between spacer abundances at different locus positions can be done in experimental data (such as ref. [23]), and extending our simulation to explicitly include multiple protospacers and spacers per genome will provide further data.

Our model makes quantitative predictions about the relationship of population size and population outcomes to average immunity that can be tested in data. We directly calculated average immunity in experimental data without the need to assemble phage genomes (Figure 7) and we found that average immunity is anticorrelated with phage population size in data in one experimental dataset, as predicted by our model (SI Figure 104). However, we did not find a correlation between average immunity and read count (a proxy for population size) in wastewater treatment plant data (SI Figure 124). When calculating average immunity using experimental data, one must make assumptions about the immune benefit of spacers. In our work, we assumed that spacer and protospacer sequences that belong to the same similarity group provide perfect immunity. Other more complex assumptions are possible, and there has been a great deal of work quantifying the efficiency of spacers based on the position of SNPs in the protospacer in vitro (see [108] for an example) and in vivo [109]. Additionally, phages regularly acquire mutations in protospacer-adjacent motifs (PAMs), and these mutations strongly decrease the targeting efficiency of bacterial spacers [102, 110].

In certain limits, average immunity can be related to the Morisita-Horn similarity index Ψ (cite Wolda 1981 from [40], see SI Section 4.1.1), where instead of comparing two populations from different timepoints or different regions, we compared bacteria and phage populations scaled by the immune benefit of CRISPR. By comparing our definition of average immunity to the Morisita-Horn index, we find that the Morisita overlap is constant across all parameters we study, which implies that the population diversity evolves to attain the highest possible overlap. Average immunity is also theoretically similar to the concept of distributed immunity introduced by Childs *et al*. [40, 50], the ‘immunity’ quantity calculated by Han *et al*. [41], and the probability of an immune encounter used by Iranzo *et al*. [98] to derive mean-field population predictions. The mathematical definition of average immunity is slightly different than these quantities, but the effect on populations is strikingly similar, and it is significant that several different models and different metrics find that a quantification of immunity is the important predictor for population-level outcomes.

### In silico time shift experiments yield qualitative trends

We calculated average immunity in two experimental datasets and performed synthetic time-shift experiments by calculating average immunity between time-shifted populations of bacteria and phage. This effectively replicates explicit time-shift experiments [81, 82, 83, 111, 85, 64, 34, 32, 27, 87]. A similar time-shift calculation using sequence data was performed by Guillemet *et al*. [31]; they approached the question from the phage perspective and found that phages are most infectious against bacteria from the past and least infectious against bacteria from the future. Our results show that powerful insights into the dynamical immune state of a population can be obtained from sequencing data without explicitly performing time shift experiments. We applied the same simple method to two very different experimental populations, suggesting that this analysis may be feasible in other time-series datasets, including from other natural microbial populations.

Our model predicts that an immune-evolving population in true steady-state must eventually experience declining past immunity. CRISPR arrays aren’t infinite, and spacers must eventually be lost, even though this may happen on very long timescales. This pattern of declining past immunity is also in line with the expectations of fluctuating selection dynamics in which more rare genotypes are more fit [112, 84, 111, 87]. Previous theoretical work applied to vertebrate adaptive immunity also supports the existance of a peak in adaptive benefit in the recent past [111, 18], and some experimental time-shift studies have reported a peak in adaptation for bacteria-phage interactions [84, 85, 87] and for HIV-immune system interactions [111, 18]. We saw two qualitatively different time shift curves in the experimental data we analyzed: in the long-term laboratory coevolution data, bacteria were more immune to all past phages and less immune to all future phages, while in the wastewater treatment plant data, bacteria were most immune to phages in their present context and less immune to phages in both the past and future. The time shift pattern of the laboratory coevolution data was consistent with experimental time-shift data that is typically said to be indicative of arms-race dynamics [81, 82, 83, 111, 86, 27], while the wastewater data may be consistent with a “zoomed-out” view of the eventual decline in immunity in both the distant past and distant future [18, 31]. Indeed, the distinction between arms race dynamics and fluctuating selection dynamics in time shift experiments may be a question of the timescale being investigated [112, 111, 87]. Our simulation results may qualitatively match both of these regimes depending on the timescale observed, which suggests that measuring very long-term coexistence is an area for further work in host-parasite coexistence experiments. In general, we expect that the timescale of memory length is related to the size of CRISPR arrays: if overall rates of spacer gain and loss are balanced, then we expect an individual spacer’s lifetime in the population to be proportional to array length. The factors that affect the timescale of memory length is an interesting area for further work.

One reason why past immunity didn’t unambiguously decline in our analysis of the laboratory coevolution data may be that spacer loss was not observed in the original experiment [24]. This is also true of several other similar experiments with *S. thermophilus* [23, 58], though spacer loss was directly observed coincident with spacer acquisition in ref. [76]. Spacer loss has also been observed in other experimental systems [113, 77, 114, 115], and indirect evidence of spacer loss through sequence comparisons has been reported for *S. thermophilus* [116] and metagenomic data [117, 35]. Spacers may also be functionally lost despite remaining in the genome, either from stochastic decoupling of RNA polymerase [118, 107, 109] or from a dilution effect of competition for limited Cas protein complexes [107, 104, 115]. The ‘loss’ rate in our model could also be taken to be gain and loss of the entire CRISPR system, which is known to happen [119, 120], although this would likely happen on much slower timescales than the acquisition and loss we model. The qualitative shape and timescale of time shift curves like the ones we presented may indirectly contain information about the timescale of spacer loss.

We found in our simulations that memory length decreased as spacer acquisition probability increased and as phage mutation rate decreased: if bacteria followed phages more closely (higher *η*) or if phages evolved more slowly (lower *µ*), the peak in past immunity was closer to the present and memory was more quickly lost (Figure 7A,E-F). A similar correspondence was derived in a theoretical model of optimal immune memory formation in vertebrates: more frequent sampling of the pathogen landscape (corresponding to higher spacer acquisition *η* in our model) led to faster memory loss [106]. Notably, that result is the optimal update scheme for an immune system wishing to minimize the costs of infection; there is no explicit optimization in our model, yet we find a pattern consistent with a potentially optimal solution.

### Vertebrate adaptive immunity

CRISPR adaptive immunity is mechanistically very different from the vertebrate adaptive immune system, yet there are several striking conceptual similarities between them and in the way past theoretical works have modeled vertebrate immunity. For example, phenomenological models of binding affinity between antigens and the immune system have used the degree of similarity between strings as a metric for interaction strength [121, 122, 79, 16, 18, 10], and more abstract models of the change in immunity after a virus mutation have also used string similarity to track the strength of immune coverage [12, 80, 7]. This is a simplification of a complex protein-protein interaction in the case of the vertebrate immune system [21], but it is nearly identical to the actual mechanism of CRISPR immunity, suggesting that lessons from models of CRISPR immunity may in turn be applicable to other forms of adaptive immunity.

We found that the speed of viral evolution was proportional to the fitness benefit of CRISPR escape mutations in our model. Theoretical models of vertebrate immunity have also studied the speed and direction of viral evolution and found several timescales of interest, ranging from the rapid timescales of population turnover (comparable to phage time to extinction in our model) and immune memory escape (comparable to immune memory length in our model) to the timescale of changes to the direction of motion in phenotype space [60]. We found a sublinear dependence of the speed of evolution on phage population size in our model (Figure 6C), analogous to the weak positive dependence on population size described by Marchi *et al*. [60]. Adding cross-reactivity between spacer types in our model changed overall population outcomes as well as population dynamics. We observed a traveling wave regime in which new phage mutants “pull” existing bacterial clones along, even if their matching clone is small or extinct (SI Figure 63). This traveling wave regime is qualitatively similar to that observed in theoretical models of “ballistic” pathogen evolution in effective low-dimensional antigenic space [7, 13, 60]. Similar clone population dynamics were observed in a model of B cell activation and memory generation in which existing memory B cells are reactivated in response to infection with a mutated virus that remains within its cross-reactivity radius [15]. We also observed splitting of lineages into divergent subtypes in our model, a phenomenon that has been studied in vertebrate pathogen evolution as well [60, 13]. We found that the average number of distinct clans was proportional to overall diversity but that increasing cross-reactivity decreased the average clan size (SI Section 4.3.3). These qualitative similarities highlight the usefulness of CRISPR adaptive immunity to understand population dynamics that also occur in the vertebrate immune system.

In our model, cross-reactivity was uniformly beneficial for bacteria, increasing bacterial average immunity. We did not explicitly model a tradeoff between affinity and cross-reactivity: in our model, increasing cross-reactivity did not decrease the maximum affinity between perfectly-matched spacers and protospacers. Such a tradeoff has been modeled in the vertebrate adaptive immune system where it was predicted that organisms should store a mixture of cross-reactive and highly specific memories to balance strong immunity against evolved threats with high specificity [14]. There may be a role for such a mixture in CRISPR immunity as well, for instance in primed adaptation [123].

Selection in the vertebrate immune system happens at many distinct stages within individuals, including T cell receptor selection in the thymus [124] and affinity maturation of B cells following an immune challenge [15, 18, 17, 11]. Selection also happens at the population level as pathogens and individuals coevolve over time [13, 7], for instance in the evolution of influenza [12, 7]. In contrast, within-host dynamics are much simpler in bacteria, and the possibilities for within-host coevolution are especially limited in the case of virulent phages that kill their hosts after a single infection. Similarly, in the vertebrate immune system, extremely high immune diversity is possible in a single individual, whereas in CRISPR immunity, total diversity per individual is many orders of magnitude lower: most bacteria contain tens of spacers with a very few examples of a few hundred spacers [101]. This reintroduces the question of how to think about immunity in microbial populations, whether as playing out on the level of individuals or on the level of the entire population. Finally, unlike the vertebrate adaptive immune system, CRISPR immunity is heritable, and so the optimal immune strategy for bacteria takes place on a timescale potentially much longer than an individual’s lifetime as well as on the level of the entire population [125]. We have highlighted several analogies between CRISPR adaptive immunity and vertebrate adaptive immunity. Overall, the mechanistic simplicity and experimental tractability of CRISPR immunity mean that abstract ideas from vertebrate immunity can be linked to measurable features of CRISPR immunity, and insights from CRISPR immunity may be useful in understanding vertebrate immunity as well.

### Selection patterns in CRISPR immunity

The mechanism of selection for immune variants in CRISPR adaptive immunity is a matter of debate. We observed several features that suggest the presence of negative frequency-dependent selection in our model: 1) decreased clone growth rates as clone size increased, 2) cyclic clone size dynamics as immune pairs oscillate in size, and 3) continual turnover in spacer types over time. This is consistent with recent experimental observations of negative frequency-dependent selection in host-pathogen systems [86, 31]. Our model also contains the base assumption that phages that can successfully infect dominant bacterial strains will experience positive selection, generating ‘kill-the-winner’ dynamics that lead to negative frequency-dependent selection for bacteria [35]. Our model does not capture true host-pathogen coevolution because bacteria are not able to evolve beyond “following” the mutational landscape of phages by acquiring spacers. In true coevolution, bacterial genomic mutations apart from the ability to acquire spacers are also drivers of evolution and sources of selection pressure on phages.

### Phase transitions in immunity

We observed large population size fluctuations when average immunity reached a particular value of approximately 10% for the parameters we used (SI Figure 35). This critical value is determined by a balance of the rate of spacer acquisition and spacer loss that leads to exactly equal fitness for both bacteria with spacers and bacteria without spacers (see SI Section 4.2.1). Despite the presence of spacers with non-zero effectiveness, near this critical point the population appears to behave as if there is no CRISPR immunity at all. This has two interesting results: first, the probability of phage establishment drops to zero for some simulations near this critical point (SI Figure 37), since if CRISPR has no influence then mutant phages do not experience strong positive selection. Second, population sizes become unstable near the transition: outcomes with quite different total bacteria population size and fraction of bacteria with spacers are possible (SI Figures 34 and 35). This critical point will be interesting to explore further as a possible phase transition.

### Experimentally accessible parameters

We found that diversity in our model depended on spacer acquisition rate, spacer effectiveness, and spacer mutation rate. These parameters are all known to vary in natural systems, and some may also be experimentally manipulated. CRISPR systems display a range of spacer acquisition rates even within the same system [96], and both spacer acquisition rate and spacer effectiveness may be controlled by the level of expression of Cas proteins, which has been found to be connected to the quorum sensing pathway [126]. Additionally, recent work has found that spacer acquisition is more efficient at slow bacterial growth rate because phages also grow more slowly, giving bacteria more time to acquire spacers [127, 30]. This change in bacterial growth rate can be controlled by nutrient concentration [30, 61], temperature [127], or by bacteriostatic antibiotics which slow bacterial growth without killing [30]. Phage mutations rates also vary and may be experimentally modified [128]. These findings all represent possible experimental manipulations of parameters relevant for CRISPR immunity; for example, given these findings we predict that a bacteria-phage population exposed to bacteriostatic antibiotics would evolve higher CRISPR spacer diversity than one without antibiotics.

## 5 Methods

### 5.1 Model

Model details can be found in SI Sections 1 and 2.1.

### 5.2 Simulation methods

Simulation details can be found in SI Section 3. Simulations were written in Python and performed on the Béluga and Niagara supercomputers at the SciNet HPC Consortium [129] and on a local server in our group. We used the tau leaping method, a fast approximation for Gillespie simulation [130].

We model spacers and protospacers as a binary sequence of length *L* as in refs. [41, 42]. When a phage reproduces and creates a burst of *B* phages, we draw *BL* numbers from a binomial distribution with probability *µ*. Successes in this draw designate bits that will mutate (flip) in the newly created phages.

Parameter descriptions and values can be found in SI Table 3 and SI Section 3.1. Our simulation code can be found on GitHub.

### 5.3 Data analysis

#### 5.3.1 Bacteria and phage coevolution sequence data

We analyzed data published by Paez-Epsino *et al*. (2015) [24] from a long-term coevolutionary experiment with *S. thermophilus* and phage 2972. We analyzed data from the MOI-2B experimental series, the longest coevolutionary series in the experiment. Full details of our methods can be found in SI Section 6.2.

We detected spacers in the CRISPR1 and CRISPR3 loci by searching raw reads for matches to the *S. thermophilus* CRISPR repeats using blast and counting adjacent sequences as spacers. We grouped all extracted spacers using agglomerative clustering with an 85% similarity threshold to assign a type label to each spacer sequence.

We detected protospacers by blasting all unique spacer sequences against all reads, then removing reads that matched to the bacterial genome or CRISPR repeat. We extracted ten nucleotides downstream from each match to check for a PAM sequence (all spacers were stored oriented in the same direction relative to the repeat to facilitate comparison and PAM detection).

We pooled all possible protospacer and spacer sequences together (separated by CRISPR locus) and performed agglomerative clustering with several different similarity thresholds between 85% and 99.5% to assign type labels based on each similarity threshold. There were no protospacer matches to the reference genome wild-type spacers for CRISPR1 and only a single protospacer match to CRISPR3, even with a lenient 85% similarity threshold. This means either that the phage has effectively escaped all the wild-type spacers, or that existing wild-type spacers target a different phage or phages altogether. We removed any protospacer sequences that did not have a perfect match to the appropriate PAM. Since targeting is highly sensitive to PAM mutations [131], we assumed that any deviation from the perfect PAM meant that the protospacer would not be successfully targeted. Relaxing this assumption did not significantly affect the qualitative trends in our results (SI Figure 101).

To calculate average immunity, we interpolated counts between sequenced time points, then calculated the time-shifted average immunity between bacteria and phage types by comparing types separated by a time delay and averaging over all points with the same time delay. We multiplied the raw average immunity values by the total number of possible protospacers on the phage genome - this accounts for bacterial immunity to the same phage via multiple possible protospacers. A theoretical justification for this procedure can be found in SI Section 6.1.

We calculated average immunity at several different spacer similarity thresholds and found that the qualitative results did not depend on the similarity threshold.

#### 5.3.2 Metagenomic CRISPR spacer analysis

We detected spacer sequences in metagenomic data from Burstein *et al*. [56]. We analyzed the 0.1 *µm* filter dataset which contains metagenomic reads at six time points, found under accession PRJNA268031. We searched for matches to the CRISPR repeats identified in the study using blast; we accessed a list of repeats from the study’s Supplementary Data 2. There were 144 unique repeat sequences. Reads are 150 bp long, and we kept only blast results where a repeat had two or more matches to a read in order to accurately detect spacers and to reduce spurious matches to repeats. To further ensure match quality, we kept only alignments that were at least 85% similar to repeat queries, unless another match to the same repeat was present on the read.

It is possible that multiple unique repeats match the same read. For each matched read, we kept the repeat match that had the highest alignment identity (fractional alignment length times percent identity). We detected spacers between repeats, then used the detected spacer length to extract spacers on either side of matched repeats. Because we required at least two repeat matches, at most three spacers could be detected from a read, and at minimum two spacers were detected. We detected a total of 37963 spacers of which there were 17491 unique sequences. The most abundant sequence was present 572 times. After grouping within each unique repeat by spacer sequence similarity with an 85% average similarity threshold, there were 12022 unique types with the largest type having an abundance of 616.

We also analyzed metagenomic data from Guerrero *et al*. [39]. We calculated average immunity and time-shifted average immunity using the same method as for data from ref. [24] (see SI section 6.3.1).

Because individual average immunity values are not normally-distributed and can also be paired with a corresponding time-shifted value, we performed a Wilcoxon signed-rank test between the average immunity at zero time shift and the average immunity at a time shift ±500 (arbitrarily chosen). There are 38 interpolated time points with a delay of ±500, and we paired each of these with their corresponding average immunity at zero delay for each bacterial abundance. Using a paired test answers the following question for each time point: is the overlap between bacteria and past phages significantly lower than the overlap in the present, and is the overlap between bacteria and future phages significantly lower than the overlap in the present? We found a Wilcoxon statistic of 392 (*p* = 0.27) for the difference between present and past immunity and a Wilcoxon statistic of 580 (*p* = 0.0012) for the difference between present and future immunity. Interestingly, these significance values are not symmetric if we pose the question from the perspective of phage: is the overlap between phage and future bacteria lower than the overlap for present phages? (Yes - Wilcoxon statistic = 507, *p* = 0.024.) And is the overlap between phage and past bacteria lower than the overlap at present? (Yes - Wilcoxon statistic = 388, *p* = 0.027.)

#### 5.3.3 Spacer turnover

For both sets of experimentally observed spacers ([24] and [56]), we calculated spacer turnover over time. We interpolated spacer abundances grouped by 85% similarity over time using the smallest time delay between samples, then calculated the average fraction of spacer types remaining as a function of time delay across the entire dataset.

## Supporting information

Figure 1B animation

Figure 4A animation

Figure 4B animation

Figure 4C animation

Figure 4D animation

Figure 6A animation

## Notes

### Competing Interest Statement

The authors have declared no competing interest.

