## Supplementary Information for "Dynamics of immune memory and learning in bacterial communities"

### Dynamics of immune memory and learning in bacterial communities - supporting information

### 1 Model description

We model bacteria and phages interacting in a chemostat. The populations we track are nutrient concentration ($C$), phages ($n_V$), and bacteria ($n_B$). Bacteria can either have no spacer ($n_B^0$) or a spacer of type $i$ ($n_B^i$), and phages can have a single protospacer of type $j$ ($n_V^j$). Nutrients flow in at concentration $C_0$ with rate $F$, and all species flow out with rate $F$. The total number of bacteria with a spacer is $n_B^s$ and the total number of bacteria is $n_B$. Bacteria grow at rate $gC$. With rate $\alpha$, a phage interacts with a bacterium. With probability $p_V$, the phage will kill bacteria without a matching spacer and produce a burst of new phages with size $B$, while for bacteria with a matching spacer that probability is reduced to $p_v^s = (1-e)p_V$ ($0 \leq e \leq 1$). Bacteria without spacers that survive an attack have a chance to acquire a spacer with probability $\eta$. Bacteria with a spacer lose their spacer at rate $r$. Phages are limited to a single protospacer, and phages can mutate to a new protospacer type with probability $\mu L$, where $\mu$ is the per-base mutation rate and $L$ is the protospacer length ($L = 30$ in all simulations). Parameter descriptions and default values are shown in SI Table 3. See refs [1, 2] for recent examples of other chemostat-based models.

#### 1.1 Reactions

Table 1: Model reactions

| | |
|---|---|
| $b^{0,i} + C \xrightarrow{g} 2b^{0,i}$ | bacterium divides |
| $b^{0,i} \xrightarrow{F} \emptyset$ | bacterium flows out |
| $V^j \xrightarrow{F} \emptyset$ | phage flows out |
| $\emptyset \xrightarrow{FC_0} C$ | nutrients flow in |
| $C \xrightarrow{F} \emptyset$ | nutrients flow out |
| $\sum_{n=0}^{B} \left( b^0 + V^j \xrightarrow{\alpha p_V P_n} (B-n)V^j + \sum_{k=1}^{n} V^{m+k} \right)$ | interaction, phage wins, $P_n$ probability of $n$ mutant phages |
| $b^0 + V^j \xrightarrow{\alpha(1-p_V)(1-\eta)} b^0$ | interaction, bacterium survives |
| $b^0 + V^j \xrightarrow{\alpha(1-p_V)\eta} b^i$ | interaction, bacterium survives and acquires a spacer |
| $\sum_{n=0}^{B} \left( b^i + V^j \xrightarrow{\alpha p_V(i,j) P_n} (B-n)V^j + \sum_{k=1}^{n} V^{m+k} \right)$ | interaction, phage wins, $P_n$ probability of $n$ mutant phages |
| $b^i + V^j \xrightarrow{\alpha(1-p_V(i,j))} b^i$ | interaction, bacterium survives |
| $b^i \xrightarrow{r} b^0$ | bacterium loses spacer |

Table 1 lists all the interactions present in our model between individual bacteria ($b$), phages ($V$), and nutrients ($C$).

In Table 1, $p_V(i,j)$ is the probability of a phage with protospacer type $j$ killing a bacteria with spacer type $i$.

$P_n$ in Table 1 is the probability of $n$ mutant phages in a burst, given by a binomial distribution:

$$P_n = P(n; B, 1-P_0) = \binom{B}{n}(1-P_0)^n (P_0)^{B-n} \tag{1}$$

where $P_0 = \mathrm{e}^{-\mu L}$ is the probability of no mutations for an individual phage. $L$ is the protospacer length and $\mu$ is the mutation probability per base per generation.

In this formulation of the mutation term, new mutants are assumed to be a new, not previously existent, phage type which increases $m$ by $n$, the number of mutations $(+\sum_{k=1}^{n} V^{m+k})$. In our simulations, phage mutations can happen to existing types, but these are rare events and we neglect them in our theoretical analysis.

#### 1.2 Master equation

The reactions in Table 1 can be formulated as a master equation describing the probability of observing $n_B^0$ bacteria without spacers, the set of $n_B^i$ bacteria with spacers of type $i$, the set of $n_V^j$ phages with protospacers of type $j$, and a nutrient concentration of $C$ at time $t$ (equation 2).

$$
\begin{aligned}
\frac{dP(n_B^0,\{n_B^i\},\{n_V^j\},C,t)}{dt} &= g(C+1)(n_B^0-1)P(n_B^0-1,\{n_B^i\},\{n_V^j\},C+1,t)\\
&+\sum_{k=1}^{m} g(C+1)(n_B^k-1)P(n_B^0,\{n_B^{i\neq k}\},n_B^k-1,\{n_V^j\},C+1,t)\\
&+F(n_B^0+1)P(n_B^0+1,\{n_B^i\},\{n_V^j\},C,t)\\
&+\sum_{k=1}^{m} F(n_B^k+1)P(n_B^0,\{n_B^{i\neq k}\},n_B^k+1,\{n_V^j\},C,t)\\
&+\sum_{\ell=1}^{m} F(n_V^\ell+1)P(n_B^0,\{n_B^i\},\{n_V^{j\neq \ell}\},n_V^\ell+1,C,t)\\
&+F(C+1)P(n_B^0,\{n_B^i\},n_V,C+1,t)\\
&+FC_0P(n_B^0,\{n_B^i\},n_V,C-1,t)\\
&+\sum_{\ell=1}^{m}\alpha(1-p_V)(1-\frac{\eta}{m})n_B^0(n_V^\ell+1)P(n_B^0,\{n_B^i\},\{n_V^{j\neq\ell}\},n_V^\ell+1,C,t)\\
&+\sum_{k=1}^{m}\sum_{\ell=1}^{m}\frac{\alpha(1-p_V)\eta}{m}(n_B^0+1)(n_V^\ell+1)P(n_B^0+1,\{n_B^{i\neq k}\},n_B^k-1,\{n_V^{j\neq\ell}\},n_V^\ell+1,C,t)\\
&+\sum_{k=1}^{m}\sum_{\ell=1}^{m}\alpha(1-p_V(k,\ell))n_B^k(n_V^\ell+1)P(n_B^0,\{n_B^{i\neq k}\},n_B^k,\{n_V^{j\neq\ell}\},n_V^\ell+1,C,t)\\
&+\sum_{k=1}^{m}\sum_{\ell=1}^{m}\sum_{n=0}^{B}\alpha p_V(k,\ell)P_n(n_V^\ell-(B-n)+1)(n_B^k+1)\\
&P(n_B^0,\{n_B^{i\neq k}\},n_B^k+1,\{n_V^{j\neq\ell}\},n_V^\ell-(B-n)+1,C,t)\\
&+\sum_{k=1}^{m}\sum_{n=0}^{B}\alpha p_V P_n(n_V^\ell-(B-n)+1)(n_B^0+1)\\
&P(n_B^0+1,\{n_B^i\},\{n_V^{j\neq\ell}\},n_V^\ell-(B-n)+1,C,t)\\
&+\sum_{k=1}^{m} r(n_B^k+1)P(n_B^0-1,\{n_B^{i\neq k}\},n_B^k+1,\{n_V^j\},C,t)\\
&-\Bigg(F(n_B^0+\sum_{k=1}^{m}n_B^k+\sum_{\ell=1}^{m}n_V^\ell+C+C_0)+gC(n_B^0+\sum_{k=1}^{m}n_B^k)\\
&+\alpha\sum_{\ell=1}^{m}n_V^\ell(n_B^0+\sum_{k=1}^{m}n_B^k)+r\sum_{k=1}^{m}n_B^k\Bigg)P(n_B^0,\{n_B^i\},\{n_V^j\},C,t)
\end{aligned}
\tag{2}
$$

Each term is included only if all respective population quantities are $\geq 0$; for instance, the first term is only included if $n_B^0 > 0$, since if $n_B^0 = 0$ then $n_B^0 - 1 < 0$ and the entire term is negative and non-physical. This is important to note especially in terms containing $n_V - (B - n) + 1$, which are only included if $n_V > B - n - 1$.

#### 1.3 Clone size master equation

We can also write master equations for the total number of clones of size $k$. These equations describe the population-level distribution of clone sizes instead of the size of a single typical clone as in equations 99 and 122.

The master equation for $b_k$, the number of bacteria clones of size $k$, is:

$$\begin{aligned}\partial_t b_k = \underbrace{gC[(k-1)b_{k-1} - kb_k]}_{\text{Bacteria growth}} + \underbrace{(F+r)[(k+1)b_{k+1} - kb_k]}_{\text{Removal of spacer-containing bacteria by chemostat outflow or spacer loss}} \\ + \underbrace{\alpha \sum_\ell \ell v_\ell \left[(k+1)b_{k+1}p_V(k+1,\ell) - kb_k p_V(k,\ell)\right]}_{\text{Phage predation - reduces clone size by 1 for each successful infection}} \\ + \underbrace{\alpha\eta n_B^0 n_v^{k*}(1-p_V)\left[b_{k-1} - b_k\right]}_{\text{New bacteria clones from spacer acquisition}}\end{aligned} \tag{3}$$

The last term representing spacer acquisition is complicated: $n_V^{k*}$ is the total number of phages which contain protospacers that match bacteria clones of size $k-1$, multiplied by the probability of getting that protospacer if infected by that phage. In other words, these are the phages that bacteria clones of size $k-1$ could acquire spacers from to increase to size $k$, multiplied by the probability of acquiring a particular spacer from those phages. In a model where phages are all identical with a large number of protospacers $m$, $n_V^{k*} = n_V/m$, where $n_V$ is the total number of phages and $1/m$ is the probability of acquiring a particular spacer. If instead phages can only have a single protospacer, $n_V^{k*}$ is not known in general, but approximations may be found; for instance, if there are $m$ phage types present and all phage clones are the same size, $n_V^{k*}$ is again equal to $n_V/m$.

The corresponding master equation for $v_\ell$, the number of phage clones of size $\ell$, is:

$$\begin{aligned}\partial_t v_\ell = \underbrace{(F + \alpha(n_B^0 + n_B^s))[(\ell+1)v_{\ell+1} - \ell v_\ell]}_{\text{Phage death from chemostat flow and adsorption}} \\ + \underbrace{\alpha \sum_k k b_k \left[\sum_{n=0}^{B} P_n(\ell-(B-n))v_{\ell-(B-n)}p_V(k,\ell-(B-n)) - \ell v_\ell p_V(k,\ell)\right]}_{\text{Phage burst after infecting spacer-containing bacteria (subtracting mutant phages which become new clones)}} \\ \underbrace{+\alpha n_B^0 p_V \left[\sum_{n=0}^{B} P_n(\ell-(B-n))v_{\ell-(B-n)} - \ell v_\ell\right]}_{\text{Phage burst after infecting bacteria without spacers}} \\ + \underbrace{\delta_{\ell,1}\alpha B(1-e^{-\mu L_p})\left[\sum_{k',\ell'} p_V(k',\ell')k'b_{k'}\ell' v_{\ell'} + p_V n_B^0 \sum_{\ell'} \ell' v_{\ell'}\right]}_{\text{New phage clones created by mutation from both spacer-containing and naive bacteria}}\end{aligned} \tag{4}$$

The phage master equation assumes that all new mutants are unique and that they are a type not currently present in the population.

$P_n$ is a binomial distribution giving the probability of $n$ mutant phages per burst (equation 1).

$p_V$ is the probability of phage successfully infecting bacteria, and in general is an unknown function of both $k$ and $\ell$. All details of immunity are contained in $p_V$; later we discuss certain choices of $p_V$ and their implications for the model and population dynamics. $p_V$ with no functional arguments is the probability of phage success against bacteria with no spacers, a constant.

### 2 Mean-field model results

#### 2.1 Total population size

The total number of bacteria with spacers, $n_B^s$, is equal to $\sum kb_k$, and the total number of phages, $n_V$, is $\sum \ell v_\ell$. $\partial_t n_B^s = \sum k \partial_t b_k$ and $\partial_t n_V = \sum \ell \partial_t v_\ell$, and summing over $k$ and $\ell$ in equations 3 and 4 gives mean-field equations:

$$\dot{n_B^s} = (gC - F - r)n_B^s + \alpha(1-p_V)\eta n_B^0 n_V - \alpha n_B^s n_V p_V^a \tag{5}$$

$$\dot{n_V} = -(F + \alpha(n_B^0 + n_B^s))n_V + \alpha B(p_V^a n_B^s n_V + p_V n_B^0 n_V) \tag{6}$$

We can also write mean-field equations for the total number of bacteria without spacers, $n_B^0$, and the total nutrients, $C$. The total number of bacteria $n_B$ is $n_B^s + n_B^0$.

$$\dot{n_B^0} = (gC - F)n_B^0 - \alpha p_V n_B^0 n_V - \alpha(1-p_V)\eta n_B^0 n_V + r n_B^s \tag{7}$$

$$\dot{C} = F(C_0 - C) - gCn_B \tag{8}$$

$$\dot{n_B} = (gC - F)n_B - \alpha(p_V^a n_B^s n_V + p_V n_B^0 n_V) \tag{9}$$

The probability of phage success against bacteria with spacers, averaged across the whole population, is given by $p_V^a = \frac{\sum_{k,\ell} kb_k \ell v_\ell p_V(k,\ell)}{n_B^s n_V}$.

$p_V^a$ is not known in general, but it can be simplified if certain assumptions are made about the population. In particular, we begin by assuming that immunity is all-or-nothing: if a bacterium has a matching spacer to a phage, the phage success probability is reduced to $p_V(1-e)$ (where $e$ is the "spacer effectiveness"), but if a bacterium has a spacer that is not exactly matching, that spacer confers no immunity and the phage probability of success is $p_V$, the same as against naive bacteria. This amounts to defining $p_V$ in terms of the spacer type $i$ and protospacer type $j$ as follows:

$$p_V(i,j) = \begin{cases} p_V(1-e) & \text{if } i = j \\ p_V & \text{if } i \neq j \end{cases} \tag{10}$$

If each bacterium and phage can have only one spacer or protospacer, then the number of bacteria or phage with a particular spacer type $i$ is $n_B^i$ and $n_V^i$ respectively, and

$$p_V^a = p_V\left(1 - \frac{e}{n_B^s n_V}\sum_i n_V^i n_B^i\right)$$

Equations 5 to 9 can be solved analytically at steady-state if we further assume $p_V^a = p_V(1-e/m)$, where $m$ is the average number of bacterial clones at steady state. This assumption is described in detail in section 4.1. The solution is exactly the mean-field analytic solution described in [3] with the simple replacement of the parameter $e$ with $e/m$.

##### 2.1.1 Phage extinction threshold

We reported in our previous work [3] that equations 5 to 9 experience a change in fixed point at a critical threshold of the phage infection success probability $p_V$: below $p_V^0 = \frac{1}{B}\left(\frac{gf}{(1-f)\alpha} + 1\right)$, phages are driven extinct (main text figure 1E). This is the same extinction threshold reported by Payne *et al.* [4] as the cutoff for achieving herd immunity in a well-mixed bacterial population. To a first approximation, phages must successfully infect every $1/B$ bacteria they encounter, but if bacteria are growing quickly, then phage must do better to overcome bacterial growth. The correction gives $\frac{1}{B}\left(1+\frac{\text{bacterial birth rate}}{\text{phage birth rate}}\right)=\frac{1}{B}\left(1+\frac{\text{phage generation time}}{\text{bacterial generation time}}\right)$ as reported in [4]. To see this, we note that the bacterial birth rate in our model is $gC \approx gC_0 f$ when phage population sizes are small, and the phage birth rate in our model is $\alpha B p_v n_B \approx \alpha n_B$ since $Bp_V \approx 1$. When phage population sizes are small, the total number of bacteria is $n_B \approx C_0(1-f)$. Combining these expressions, we find $\frac{\text{bacterial birth rate}}{\text{phage birth rate}} \approx \frac{fgC_0}{\alpha(1-f)C_0} = \frac{fg}{\alpha(1-f)}$ as in our original expression.

#### 2.2 Steady-state

Simulation results are calculated and presented at steady-state, unless otherwise specified. We define steady-state to be the simulation run-time divided by 5, choosing simulation run-times so that mean-field quantities have equilibrated by the steady-state time. We run large population simulations longer than small population simulations: these simulations have less frequent interactions on average because of decreased $\alpha$ and take longer to equilibrate (SI Figure 1).

We set the total bacterial generations to be $10000(\log C_0 - 3)$, rounded to the nearest thousand, with a minimum of 10000 generations (see Table 2). The following table lists simulation length and assumed steady-state time $t_{ss}$ for different values of $C_0$.

Table 2: Simulation length

| $C_0$ | Simulation length (bacterial generations) | Steady-state start time ($t_{ss}$) |
|---|---|---|
| 300 | 10000 | 2000 |
| 1000 | 10000 | 2000 |
| 3000 | 10000 | 2000 |
| 10000 | 10000 | 2000 |
| 30000 | 15000 | 3000 |
| 100000 | 20000 | 4000 |
| 300000 | 25000 | 5000 |
| 1000000 | 30000 | 6000 |

SI Figure 2 shows the mean number of bacterial clones ($m$) at steady-state as a function of the initial number of phage clones. For most parameters, the steady-state $m$ is independent of the initial m. There is a slight dependence on initial $m$ at high $C_0$.

#### 2.3 Single-clone mean-field dynamics

We can write mean-field equations for individual clones $n_B^i$ and $n_V^j$ as well:

$$\dot{n_V^j} = -(F + \alpha(n_B^0 + n_B^s))n_V^j + \alpha B P_0 p_V n_B^0 n_V^j + \alpha B P_0 \sum_i p_V(i,j) n_B^i n_V^j + \alpha B \sum_j \sum_{k \neq i} p_V(j,k) \mu^{\Delta_{ik}} n_B^j n_V^k \tag{11}$$

$$\dot{n_B^i} = (gC - F - r)n_B^i - \alpha \sum_j p_V(i,j) n_B^i n_V^j + \alpha \eta n_B^0 n_V^i (1 - p_V) \tag{12}$$

If $p_V(i,j)$ is binary and all spacers are equally effective (equation 10), $\dot{n_V^j}$ and $\dot{n_B^i}$ can be simplified:

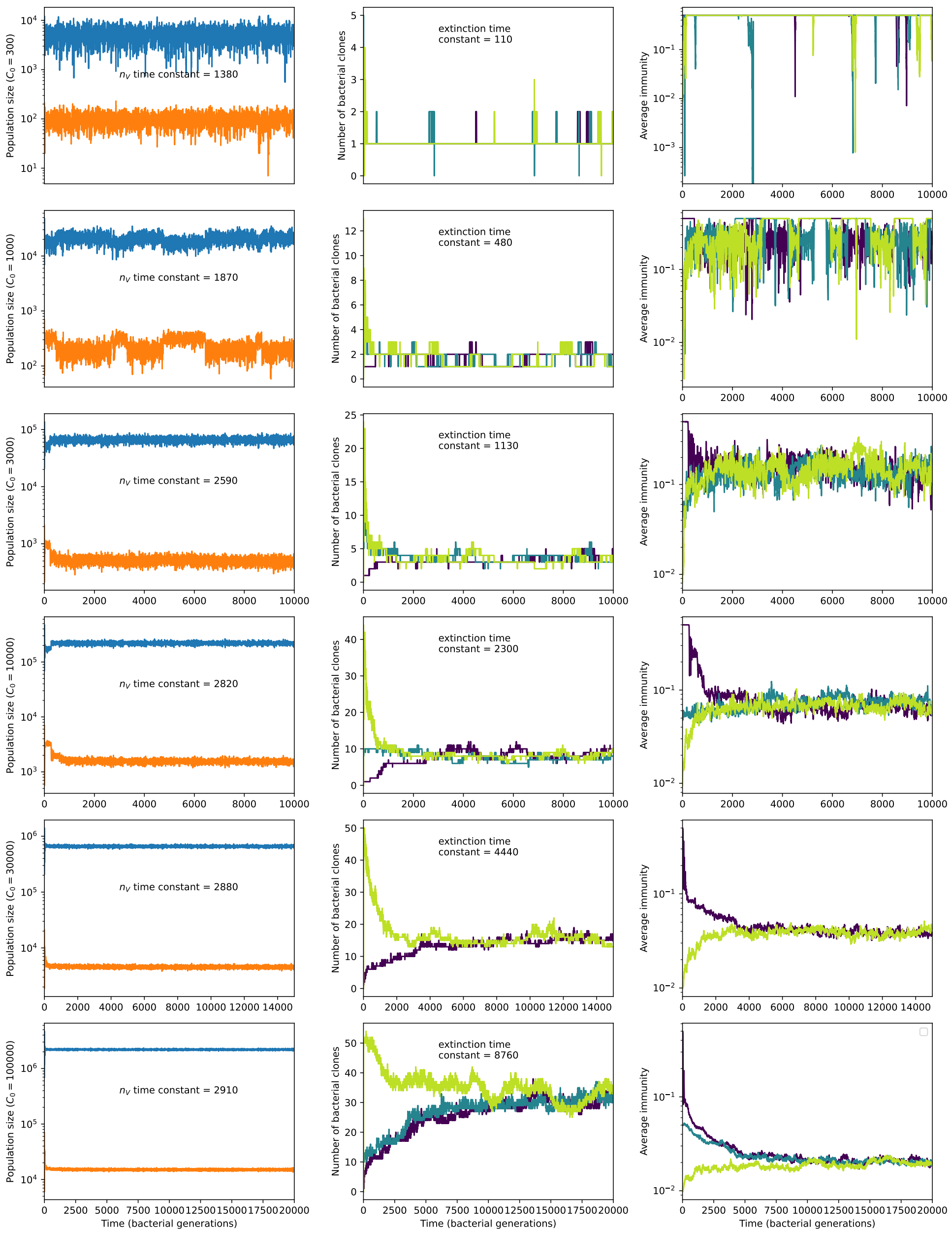

Figure 1: Total phage, total bacteria (left), and mean number of bacterial clones (right) vs. simulation time for five simulations with $\mu = 10^{-6}$, $e = 0.5$, $\eta = 0.001$, and $C_0$ ranging from 300 (top row) to 30000 (bottom row). Total population sizes equilibrate very quickly, but the total number of clones can take longer at large population sizes (high $C_0$). The time constants inset are a measure of how quickly we expect each mean-field quantity to equilibrate: $n_V$ time constant is the inverse growth rate of the total phage population ($1/(-F - \alpha n_B(Bp_V - 1) - \alpha B p_V n_B^s e/m)$) and the extinction time constant is the mean time to extinction for large phage clones (equation 146), a measure of the rate of turnover of the number of clones.

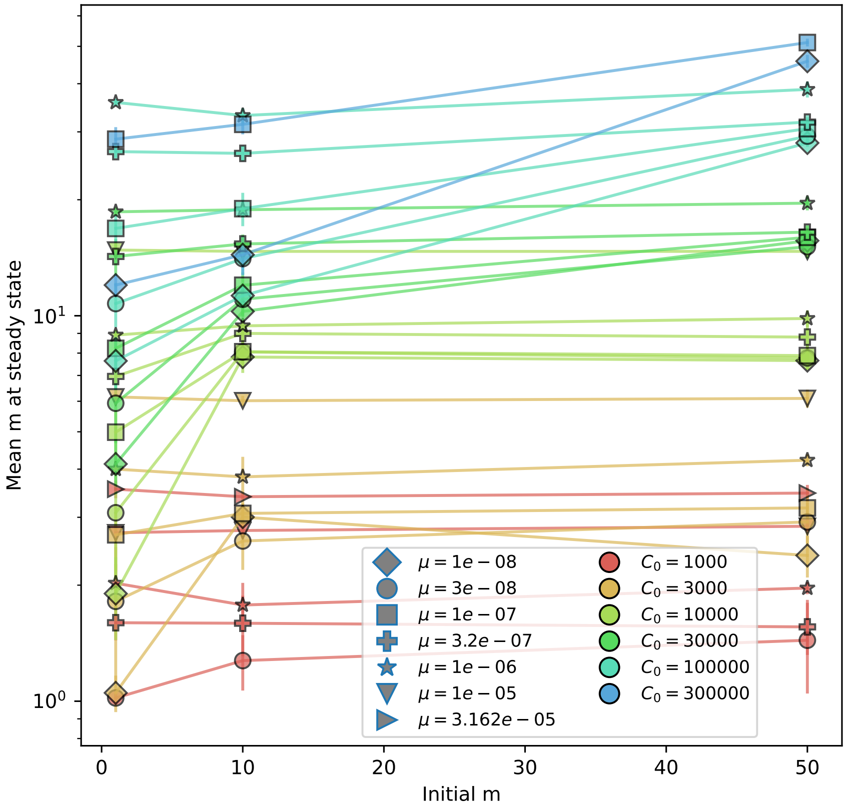

Figure 2: Mean number of bacterial clones after $t = t_{ss}$ bacterial generations vs. initial number of phage clones for $e = 0.8$, $\eta = 10^{-3}$. The mean is an average of 15 evenly-spaced points from $t = t_{ss}$ to $t = 5t_{ss}$ bacterial generations. Error bars are an average across three or more independent simulations.

$$\dot{n_V^j} = -(F + \alpha n_B)n_V^j + \alpha B P_0 p_V n_V^j(n_B - e n_B^j) + \alpha B \sum_j \sum_{k \neq i} p_V(j,k)\mu^{\Delta_{ik}} n_B^j n_V^k \tag{13}$$

$$\dot{n_B^i} = (gC - F - r)n_B^i - \alpha p_V n_B^i(n_V - e n_V^i) + \alpha\eta n_B^0 n_V^i(1 - p_V) \tag{14}$$

In equations 11 and 13, mutations that arise from phage $i$ decrease $n_V^i$, which is captured in that $B$ becomes $BP_0 = B\mathrm{e}^{-\mu L}$. $B$ for type $i$ is effectively lower, since mutations go to different phage types.

The last term in equations 11 and 13 is for mutations that happen in all other phages besides type $i$ that convert type $k$ to type $i$. Any particular phage $k$ has a certain mutational distance from phage $i$, $\Delta_{ik}$, which means $\Delta_{ik}$ specific mutations must happen to get from type $k$ to type $i$. This happens with probability $\mu^{\Delta_{ik}}$. Going forward we assume that this term is small, which is true if the overall mutation rate $\alpha B p_V^a \mu n_V n_B$ is sufficiently low and/or the space of protospacer types is sufficiently large so that mutations are almost always to new types not present in the population.

We solve the coupled time-dependent system given by equations 13 and 14 numerically, assuming all other populations ($n_B$, $n_B^0$, $n_V$, and $C$) are at their deterministic steady-state value and ignoring the last term of equation 13. These solutions are plotted alongside mean clone sizes from a simulation in Figure 3 ("Numerical deterministic prediction"). The deterministic solution matches the mean clone size well at early times but does not capture the effects of clone extinction at later times. By including only surviving clones in the simulation mean and normalizing the deterministic solution by the predicted fraction of surviving clones ("Numerical deterministic prediction normalized to survival probability"), the theoretical prediction matches well at early times and can be piecewise-combined with the steady-state mean clone size to give good agreement at all times (see "Predicted clone size" dashed lines in Figure 77). We estimate the predicted fraction of surviving phage clones by numerically solving equation 128 which gives the phage clone probability of extinction in the absence of matching bacterial clones. The fraction of surviving clones is $1 - P_0(t)$ where $P_0(t)$ is the probability of extinction. Normalizing by $1 - P_0(t)$ does not give good agreement at long times because $P_0(t)$ goes to 1 as $t$ goes to infinity, causing the solution to diverge at long times. While individual trajectories do eventually go extinct, the mean clone size conditioned on survival reaches the deterministic steady state at long times.

We consider bacteria clones in the background of phage clones; that is, time 0 for a bacterial clone starts when the matching phage clone arises by mutation. This is the case in the numerical solution for the coupled bacteria-phage clone system given by equations 13 and 14. For this reason, normalizing the bacterial clone size prediction by the phage clone probability of extinction also gives good agreement to the simulation mean conditioned on bacterial clone survival. (If a bacterial clone size is 0 but its corresponding phage clone has not yet gone extinct, that 0 will count in the mean clone size, but after the phage goes extinct, a 0 in the bacterial clone size will not be included.)

We can also solve equations 13 and 14 analytically by assuming $n_B^i$ and $n_V^i$ are constant respectively; this amounts to removing the dependence of phage clone dynamics on bacteria clone dynamics and vice versa.

If we assume $n_B^i = 0$, we get the following solution for $n_V^i(t)$:

$$n_V^i(t) = n_V^i(0)\mathrm{e}^{s_0 t} \tag{15}$$

where $s_0 = \alpha B p_V n_B \mathrm{e}^{-\mu L} - F - \alpha n_B$ is the average growth rate of phage clones and $t$ is in minutes. This is the same growth rate as is in equation 131 with a small correction: $B \to B\mathrm{e}^{-\mu L}$; the burst of an individual phage clone is reduced by phages that mutate to a new type.

Likewise if we assume that $n_V^i$ is constant, we get the following solution for $n_B^i(t)$:

$$n_B^i(t) = n_B^i(0)\mathrm{e}^{s_B t} + \frac{\delta}{s_B}\left(\mathrm{e}^{s_B t} - 1\right) \tag{16}$$

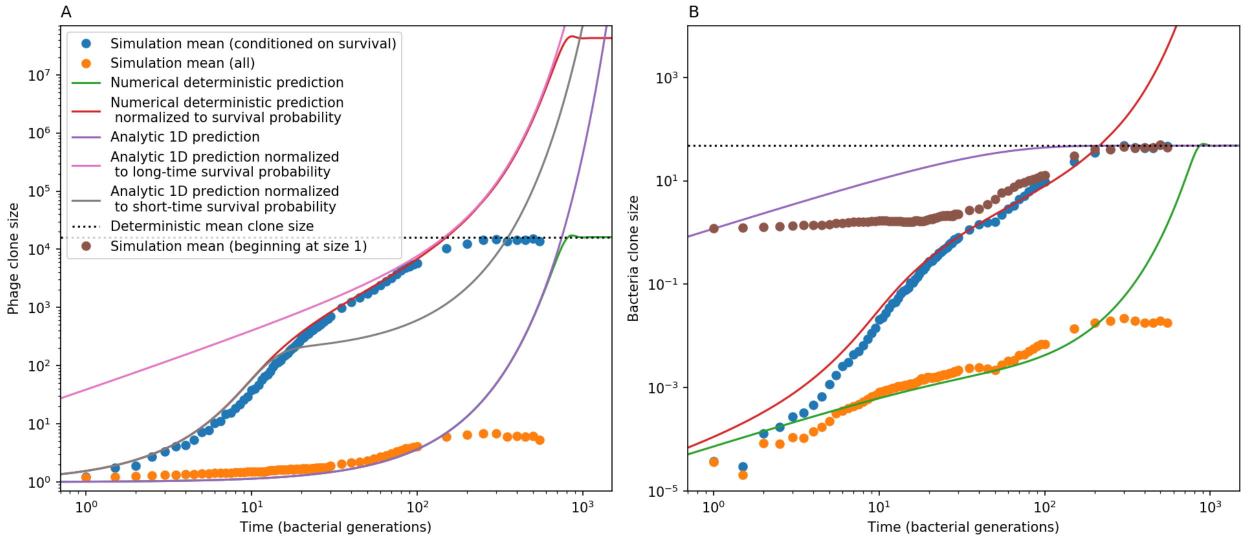

Figure 3: Phage (A) and bacteria (B) mean clone size in a simulation, either conditioned on survival (blue circles) or including extinct clones (orange circles). Theoretical predictions are plotted as solid lines: the time-dependent numerical solution to equations 13 and 14 in green, the same solution divided by the phage clone probability of survival in red, and a one-dimensional solution to equation 13 in (A) and 14 in (B). Equations 17 and 18 are black dashed lines. An alternate simulation mean clone size is plotted for bacteria (brown circles) in which each clone trajectory is stacked based on the bacterial acquisition time and averaged across trajectories, conditioned on survival. Simulation parameters are $C_0 = 10000.0$, $\mu = 10^{-5}$, $\eta = 0.001$, and $e = 0.95$.

where $s_B = gC - F - r - \alpha p_V(n_V - n_V^i)$ is the average growth rate for bacterial clone $i$ and $\delta = \alpha\eta n_B^0 n_V^i(1 - p_V)$ is the rate of spacer acquisition for that clone.

Equations 15 and 16 are plotted in Figure 3 ("Analytic 1D prediction"). Equation 15 matches the phage clone size not conditioned on survival at early times, but continues to grow past the steady-state phage clone size without growing bacterial clones to reign it in. Equation 16 is not comparable to the simulation mean with time zero as the time of phage clone birth; rather it should be compared to bacterial clone growth independent of what the phages are doing. Using the steady-state mean phage clone size in equation 16 (purple line), we obtain a rough correspondence with the simulation mean (brown circles). The prediction does not match at intermediate times, likely because this solution assumes that phage clones are at their mean size when in fact they are likely still smaller.

Equations 13 and 14 can be solved at steady state in terms of the total mean-field variables (ignoring the last term of equation 13). These solutions (equations 17 and 18) are indicated by horizontal dashed lines in Figure 3.

$$n_B^{i\,*} = \frac{1}{e}\left(n_B - \frac{F + \alpha n_B}{\alpha B P_0 p_V}\right) \tag{17}$$

$$n_V^{i\,*} = \frac{n_B^{i\,*}(\alpha p_V n_V - (gC - F - r))}{\alpha\eta n_B^0(1 - p_V) + \alpha p_V e n_B^{i\,*}} \tag{18}$$

##### 2.3.1 Quasi-steady-state clone size solution

We can find a quasi-steady-state solution for the clone sizes $n_B^i$ and $n_V^i$: we assume that $n_B^i$ equilibrates much faster than $n_V^i$ so that $\dot{n}_B^i = 0$. We first solve equation 14 leaving $n_V^i$:

$$n_B^{i\,*} = \frac{\alpha\eta n_B^0(1-p_V)n_V^i}{F+r-gC+\alpha p_V(n_V-en_V^i)} \tag{19}$$

We substitute this solution into equation 13 giving a one-dimensional equation for $n_V^i$ (neglecting the third term in the equation as before):

$$\dot{n_V^i} \approx -(F+\alpha n_B)n_V^i + \alpha BP_0p_Vn_V^i\left(n_B - e\frac{\alpha\eta n_B^0(1-p_V)n_V^i}{F+r-gC+\alpha p_V(n_V-en_V^i)}\right) \tag{20}$$

We numerically solve for $n_V^i$, then substitute the solution for $n_V^i$ back into the solution for $n_B^i$. An example of one such quasi-steady-state solution is shown in Figure 5.

##### 2.3.2 Clone size stability analysis

With the above deterministic clone size solutions, we can investigate the stability of this fixed point. The Jacobian of this fixed point is

$$J = \begin{pmatrix} gC-F-r-\alpha p_V(n_V-en_V^i) & \alpha p_Ven_B^i + \alpha\eta n_B^0(1-p_V) \\ -\alpha BP_0p_Ven_V^i & -(F+\alpha n_B)+\alpha BP_0p_V(n_B-en_B^i) \end{pmatrix} \tag{21}$$

The lower right term is zero because we have set $\dot{n_V^i}=0$. Substituting the steady-state solutions for $n_B^i$ and $n_V^i$ (equations 17 and 18):

$$J = \begin{pmatrix} \frac{\alpha B\eta n_B^0P_0(1-p_V)(F-gC+\alpha n_Vp_V+r)}{F+\alpha(n_B-B\eta n_B^0P_0(1-p_V)-Bn_BP_0p_V)} & -\frac{F+\alpha(n_B-B\eta n_B^0P_0(1-p_V)-Bn_BP_0p_V)}{BP_0} \\ -\frac{(BP_0(-F+\alpha n_B(-1+BP_0p_V))(F-gC+\alpha n_Vp_V+r)}{F+\alpha(n_B-B\eta n_B^0P_0(1-p_V)-Bn_BP_0p_V)} & 0 \end{pmatrix} \tag{22}$$

The eigenvalues of a Jacobian $J = \begin{pmatrix} a & b \\ c & d \end{pmatrix}$ are given by $\lambda = \frac{\tau\pm\sqrt{\tau^2-4\Delta}}{2}$, with $\tau = a+d$ and $\Delta = ad-bc$. Here we have $\tau = \frac{\alpha B\eta n_B^0P_0(1-p_V)(F-gC+\alpha n_Vp_V+r)}{F+\alpha(n_B-B\eta n_B^0P_0(1-p_V)-Bn_BP_0p_V)}$ and $\Delta = -(F+\alpha n_B(1-BP_0p_V))(F-gC+\alpha n_Vp_V+r)$. If the real part of the eigenvalues is negative, the fixed point is stable, and if $\tau^2-4\Delta$ is negative (i.e. nonzero imaginary part), the fixed point is oscillatory.

For $\tau$, we have $\alpha B\eta n_B^0P_0(1-p_V) > 0$ (since $0 < p_V < 1$), and we can use $\dot{n_B^s} = 0$ to show that $F-gC+\alpha n_Vp_V+r > 0$ at steady-state and $\dot{n_V} = 0$ to show that $F+\alpha(n_B - B\eta n_B^0P_0(1-p_V) - Bn_BP_0p_V) < 0$ at steady-state, which means $\tau < 0$ for all physical parameters.

Here we show this in more detail. From $\dot{n_V} = 0$: $F+\alpha n_B = \alpha B(p_V^an_B^s + p_Vn_B^0)$. Assuming $P_0 \approx 1$, we can substitute this into the denominator of $\tau$:

$$\alpha B(p_V^an_B^s + p_Vn_B^0) - \alpha B\eta n_B^0(1-p_V) - \alpha Bn_Bp_V$$

$$\alpha Bp_V\left(\frac{p_V^a}{p_V}n_B - \frac{p_V^a}{p_V}n_B^0 + p_Vn_B^0 - p_Vn_B - \frac{\eta n_B^0(1-p_V)}{p_V}\right)$$

$$\alpha Bp_V\left((p_V - \frac{p_V^a}{p_V})(n_B^0-n_B) - \frac{\eta n_B^0(1-p_V)}{p_V}\right)$$

Now $p_V < 1$, $p_V^a \leq p_V$ by definition, and $n_B^0 < n_B$, so this term is strictly negative. Putting it all together, $\tau$ is strictly negative.

Turning to the square root: we calculate $\tau^2 - 4\Delta$:

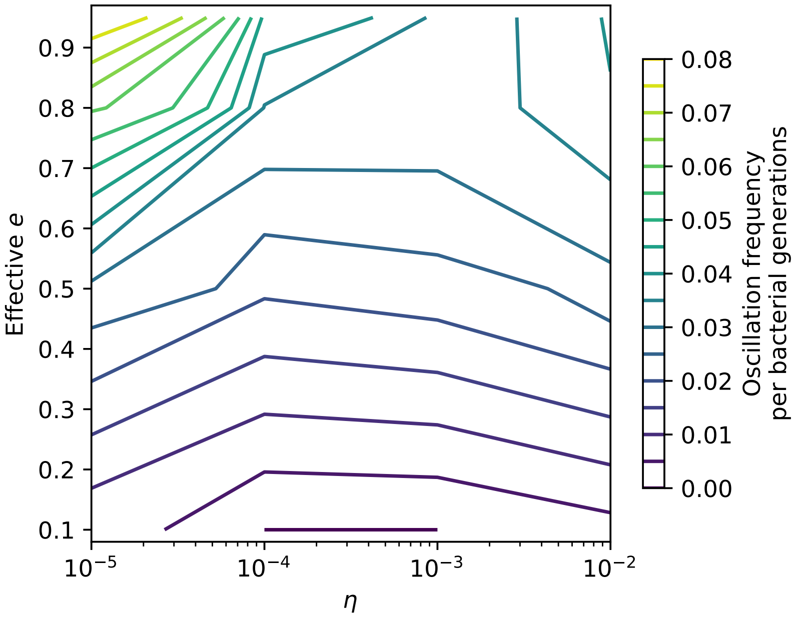

Figure 4: Theoretical oscillation frequency $\omega$ for clone trajectories oscillating about their clone size fixed point for different values of effective $e$ and $\eta$. $C_0 = 10^4$, and $\mu = 10^{-6}$.

$$\tau^2 - 4\Delta = \frac{\alpha\eta^2 {n_B^0}^2 n_V (1-p_V)^2 (\eta n_B^0 (1-p_V) + (n_B - n_B^0)(p_V - p_V^a)}{(n_B - n_B^0)\left[\eta n_B^0 (1-p_V) + (n_B - n_B^0)(p_V^2 - p_V^a)\right]^2} - 4\alpha B (n_B - n_B^0)(p_V - p_V^a)$$

All of these terms except the denominator are strictly positive, but the denominator is difficult to evaluate because $p_V^2 - p_V^a < 0$. Numerically, most parameters give $\tau^2 - 4\Delta < 0$, except at low $e$, which means the fixed point is stable and usually oscillatory except at low $e$. For a two-dimensional system, trajectories can be written $x(t) = c_1 e^{\lambda_1 t}\vec{v_1} + c_2 e^{\lambda_2 t}\vec{v_2}$, where $\lambda$ are the eigenvalues and $\vec{v}$ are the eigenvectors. If the eigenvalues are complex, i.e. $\lambda = a \pm i\omega$, then the trajectories are combinations of $e^{at}\cos(\omega t)$ and $e^{at}\sin(\omega t)$, and so $\omega = \frac{1}{2}\sqrt{4\Delta - \tau^2}$ gives the frequency of oscillation about the fixed point [5].

Figure 4 shows the predicted clone size fixed point oscillation frequency for different values of effective $e$ and $\eta$. The oscillation frequency is larger at high $e$ and low $\eta$, and the oscillation frequency is zero for $e = 0.1$ and moderate to high $\eta$.

We find that clone trajectories are oscillatory in simulations; Figure 5 shows theoretical and simulated clone trajectories for a simulation. The details of each trajectory's oscillation are different from the deterministic mean, but the average over all simulation trajectories is qualitatively similar to the deterministic prediction (Figure 6) and the quasi-steady-state solution effectively describes trajectories if oscillations were removed.

### 3 Description of simulations

Simulations were written in Python and performed on the Béluga and Niagara supercomputers at the SciNet HPC Consortium [6] and on a local server in our group. We used the tau leaping method, a fast approximation for Gillespie simulation [7] and compared with full Gillespie simulations for some cases. Gillespie and tau-leaping methods showed similar dynamics and good agreement for total population

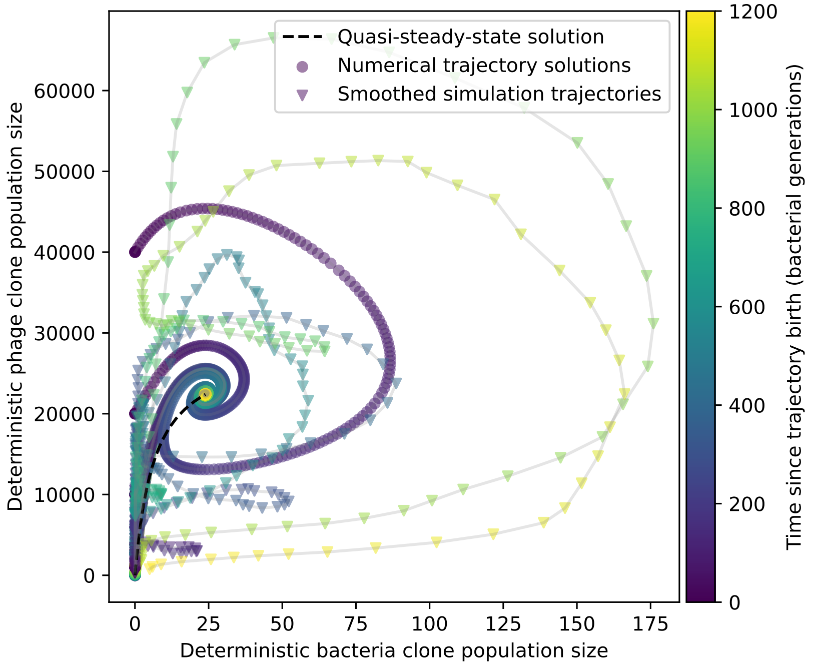

Figure 5: Predicted and observed trajectories in a simulation with $C_0 = 10^4, \eta = 10^{-4}, \mu = 10^{-5}$, and $e = 0.95$. Circles show the numerical solution to the two-dimensional system described by equations 13 and 14. The dashed line is a quasi-steady-state solution reached by solving equation 20, then substituting the solution into equation 19 for $n_B^i$. Triangles show smoothed simulation trajectories; each point is a running mean of the previous 100 points. Oscillations have larger amplitude in the simulated trajectories than in the deterministic trajectories.

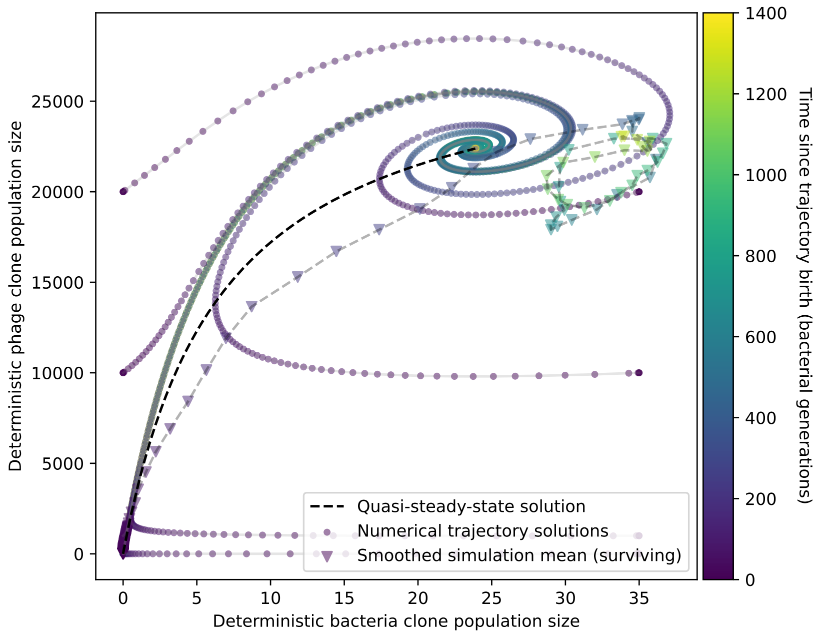

Figure 6: Predicted and observed mean trajectories in a simulation with $C_0 = 10^4, \eta = 10^{-4}, \mu = 10^{-5}$, and $e = 0.95$. Circles show the numerical solution to the two-dimensional system described by equations 13 and 14. The dashed line is a quasi-steady-state solution reached by solving equation 20, then substituting the solution into equation 19 for $n_B^i$. Triangles show the smoothed mean simulation trajectory conditioned on survival; each point is a running mean of the previous 10 points.

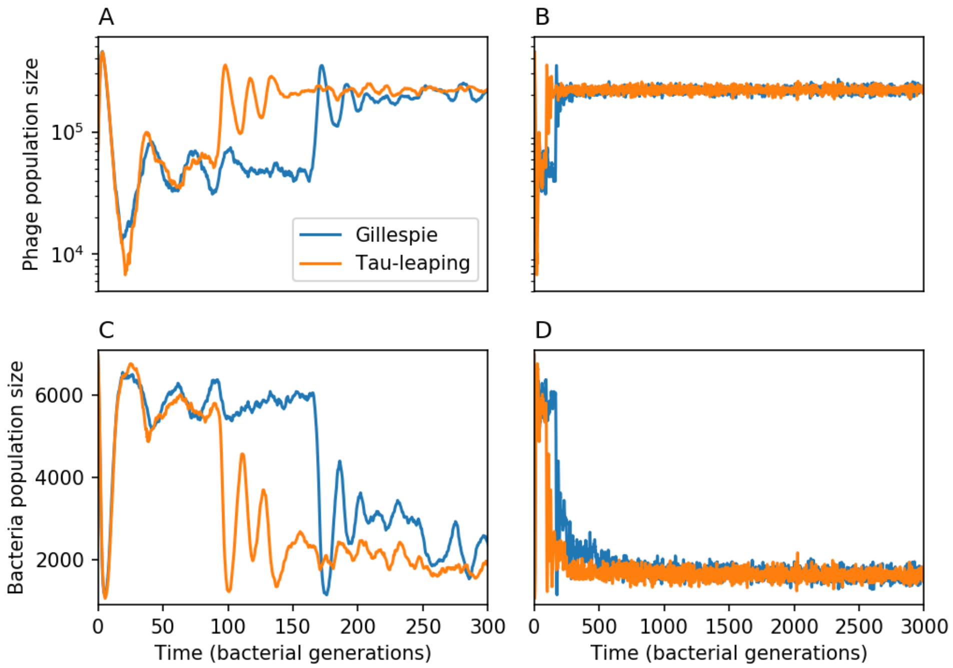

Figure 7: Total phage ($n_V$) and total bacteria ($n_B$) as a function of time for a Gillespie simulation and a tau leaping simulation with the same parameters. Total phage is shown at early times (A) and late times (B), and total bacteria at early times (C) and late times (D). The simulation parameters are $C_0 = 10^4$, $\mu = 10^{-6}$, $\eta = 0.001$, and $e = 0.95$. Early time dynamics differ slightly in this example, but the long-time behaviour and steady-state values are similar.

sizes (Figure 7), total spacer diversity (Figure 8), mean phage and bacteria clone sizes (Figure 9) and produced the same qualitative behaviour for individual spacer types (Figure 10).

We initialized five simulations for each parameter combination for a total of approximately 19800 simulations, not including simulations with cross-reactivity. A small subset of simulation parameters were run six times instead of five. Not all simulations were successfully completed, either due to errors while running or because their running time was exceedingly long. Of the initialized simulations, approximately 10000 were completed and included in analysis. Simulations with very low $\mu$ tended to either have no phage establishments (at low $C_0$) or very long running times (at high $C_0$), and simulations with very high $\mu$ tended to have higher diversity and longer running times in general. Unless otherwise noted, plots with simulation averages are an average across three to six simulations with the same parameters. We set simulations to run for a fixed number of bacterial generations intended to be long enough to allow the system to reach steady-state and remain there for a long time to generate statistics (SI Section 2.2).

We model spacers and protospacers as a binary sequence of length $L$ as in refs [8, 9]. When a phage reproduces and creates a burst of $B$ phages, we draw $BL$ numbers from a binomial distribution with probability $\mu$. Successes in this draw designate bits that will mutate (flip) in the newly created phages. It is possible but very rare for more than one mutation to happen in the same new phage: this occurs with probability $1 - \mathrm{e}^{-\mu L} - \mu L \mathrm{e}^{-\mu L} \approx (\mu L)^2$ whereas a single mutation occurs with probability $\mu L \mathrm{e}^{-\mu L} \approx \mu L$. Multiple mutations occur approximately $\mu L$ times as often as single mutations; for the largest value of $\mu$ we use, they represent 0.3% of events.

Our simulation code can be found on GitHub.

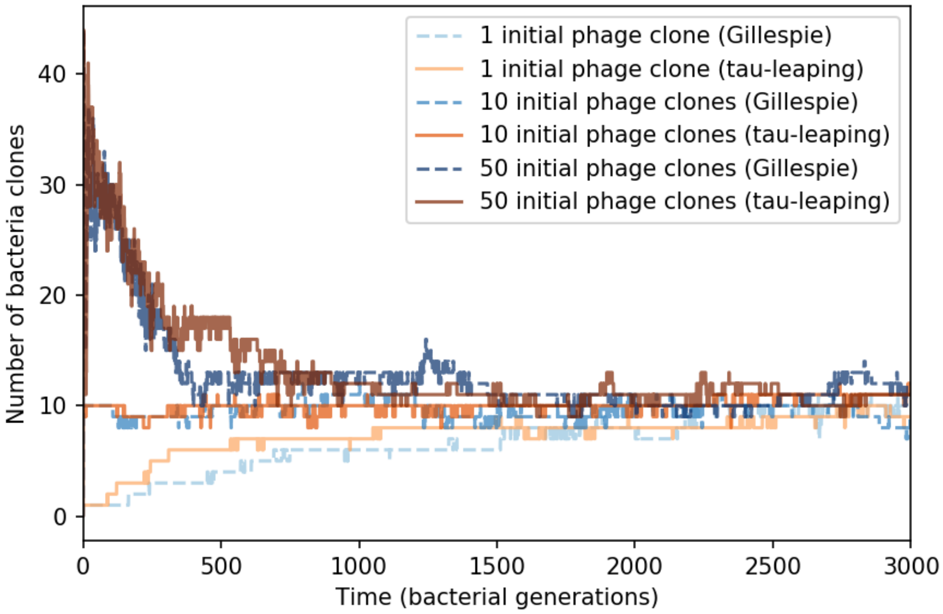

Figure 8: Number of bacterial clones ($m$) vs. simulation time for three sets of simulations, each beginning with 1, 10, or 50 phage clones. Gillespie simulations are dashed blue lines and tau leaping simulations are solid orange and red lines. The simulation parameters are $C_0 = 10^4$, $\mu = 10^{-6}$, $\eta = 0.001$, and $e = 0.95$.

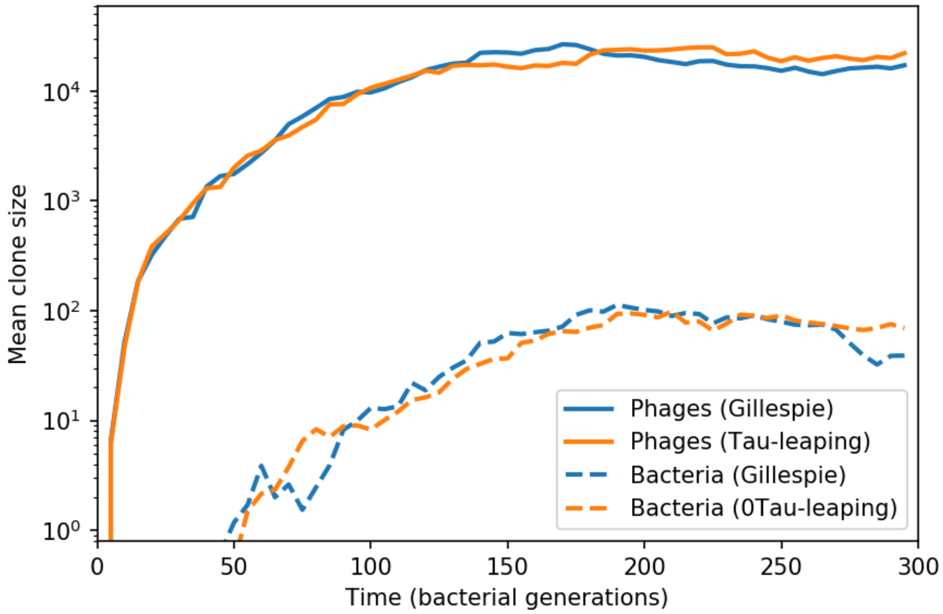

Figure 9: Average population size of phage clones (solid lines) and bacterial clones (dashed lines) in a Gillespie and tau-leaping simulation with the same parameters. The simulation parameters are $C_0 = 10^4$, $\mu = 10^{-6}$, $\eta = 0.001$, and $e = 0.95$.

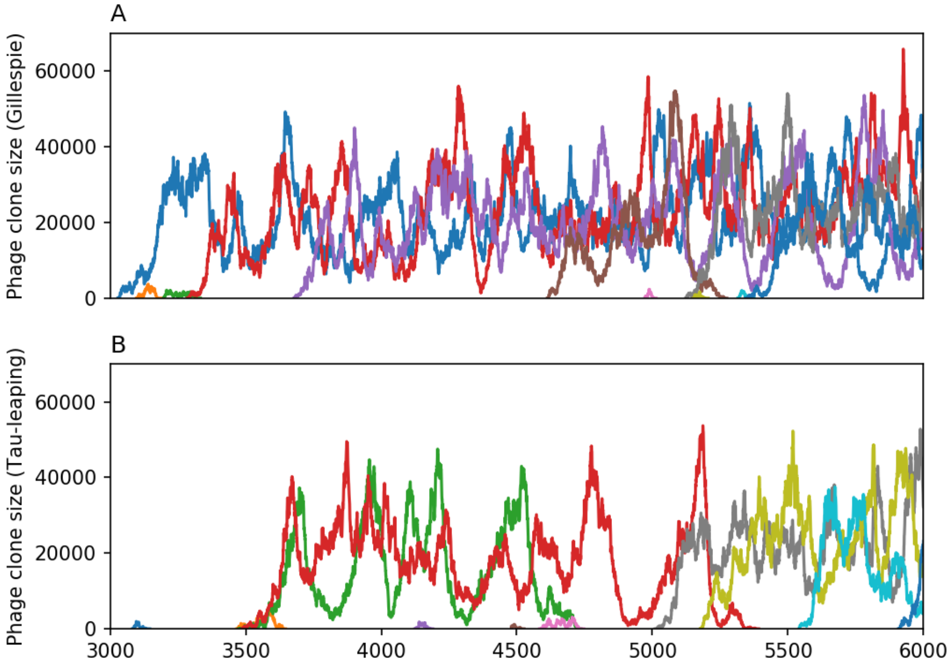

Figure 10: 10 spacer trajectories for a Gillespie simulation (A) and tau-leaping simulation (B). The first 10 trajectories that surpass $n_V^i = 1000$ are shown. The simulation parameters are $C_0 = 10^4$, $\mu = 10^{-6}$, $\eta = 0.001$, and $e = 0.95$.

#### 3.1 Parameter values

Parameter descriptions and values are listed in Table 3. The burst size for phage that target *S. thermophilus* has been measured at between 140-200 [10] and 80 for phage 2972 [11]. We use a burst size of 170. Ref. [12] measured the maximum growth rate of *S. thermophilus* in milk at 42°C to be $2.4 \times 10^{-2}$ min$^{-1}$. This corresponds to $gC_0$ in our model; we choose $g$ based on $C_0$ so that $gC_0 = 2.4 \times 10^{-2}$ min$^{-1}$.

The parameter $\alpha$ in our model is the phage adsorption rate constant divided by the culture volume. The rate of adsorption for phage is of the order of $10^{-8}$ min$^{-1}$ ml [13]; this is similar to values used in [11] and [14]. In order to explore regimes of different total population sizes, we decrease $\alpha$ as we increase $C_0$ in order to maintain stable coexistence between bacteria and phage; this is equivalent to decreasing the culture volume as $C_0$ decreases. For example, if $C_0 = 10^5$, we set $\alpha = 2 \times 10^{-2}/C_0 = 2 \times 10^{-7}$, implying a culture volume of approximately 50$\mu$l.

Levin *et al.* [11] estimated the frequency of phage mutants that escape CRISPR targeting by *S. thermophilus* to be between $5 \times 10^{-7}$ to $5 \times 10^{-5}$. These measurements are the fractions of phages from lysate that can evade CRISPR targeting of different unique spacers. This is analogous to $\mu L$ in our model, which is the probability of a mutation occurring in a newly burst phage. We use values of $\mu$ between $10^{-8}$ and $10^{-4}$ (corresponding to $\mu L$ between $3 \times 10^{-7}$ and $3 \times 10^{-3}$ to encompass the physiological range of phage mutation rates as well as unusually high mutation rates.

We generally fix the probability of successful phage infection against naive bacteria at $p_V = 0.02$ (though $p_V$ is varied in Figure 1E and in [3]). This is consistent with the value of $10^{-2}$ used in [15] and [16]. Several other models of CRISPR immunity have assumed phage success probability to be much higher, typically close to 1 [17, 18, 14]. Increasing $p_V$ does not qualitatively alter our results beyond changing the relative population sizes of phage and bacteria; at high values of $p_V$ bacteria population sizes are small and they become more likely to experience stochastic extinction (see Figure 1E). A low value of $p_V$ can be taken to reflect the presence or effectiveness of other anti-phage defense systems.

Table 3: Model parameters

| Parameter | Description | Value |
|---|---|---|
| $\frac{1}{gC_0}$ | Bacterial doubling time | 41.7 min |
| $C_0$ | Inflow nutrient concentration in units of bacterial cell density | $10^2$ to $10^6$ |
| $\alpha$ | Phage adsorption rate | $2 \times 10^{-2}/C_0$ |
| $B$ | Phage burst size | 170 |
| $F$ | Chemostat flow rate | $0.3gC_0$ |
| $p_V$ | Probability of phage success for bacteria without spacers | 0.02 |
| $e$ | Spacer effectiveness | 0.1 to 0.95 |
| $r$ | Rate of spacer loss | $0.04gC_0$ |
| $\eta$ | Probability of spacer acquisition | $10^{-5}$ to $10^{-2}$ |
| $\mu$ | Phage mutation rate per base per generation | $10^{-8}$ to $10^{-4}$ |
| $L$ | Phage protospacer length in nucleotides | 30 |

Parameter values are as above unless otherwise indicated. Representative values estimated for Streptococcus thermophilus bacteria in lab conditions.

The rate of spacer loss in our model, $r$, can be thought of as a phenomenological parameter, since the true rate of spacer loss is not well understood [14]. For comparison, Jiang *et al.* estimate the rate of loss of function of the entire CRISPR system in S. *epidermis* at $10^{-4}$ to $10^{-3}$ per individiual per generation [19]. We rescale the parameter $r$ as a function of bacterial growth rate, which means that the rescaled parameter $R = r/(gC_0) = 0.04$ is constant per bacterial generation. Our loss rate is an order of magnitude higher than the rate in [19].

We vary the parameters $\eta$ and $e$ to explore different "strengths" of CRISPR immunity. The parameter $e$ is spacer effectiveness: when $e = 0$, spacers provide no immunity and bacteria with spacers are functionally no different from bacteria without spacers. When $e = 1$, a spacer-containing bacterium that encounters a phage with a matching protospacer is guaranteed to survive. We vary $e$ between 0.1 and 0.95 to explore different regimes of CRISPR effectiveness. The parameter $\eta$ is the probability that a naive bacterium will acquire a spacer if it is infected but not killed by a phage. Rates of naive spacer acquisition vary widely, and we vary $\eta$ between $10^{-5}$ and $10^{-2}$, with the value of $\eta$ constant within a simulation. For comparison with measured acquisition rates, Pyenson *et al.* measured naive acquisition in *Staphylococcus aureus* to be approximately $10^{-6}$ to $10^{-7}$ [20]. Acquisition rates may hundreds of times higher in primed acquisition [21], and Heler *et al.* measured four orders of magnitude difference in spacer abundances shortly after infection, likely a result of differences in acquisition rate [22].

The flow rate $F$ was picked in order to get a stable fixed point where phage and bacteria coexist. Stability conditions approximately correspond to those derived in our previous work [3].

#### 3.2 Stochastic population extinction

Even when bacteria and phage are within the deterministic coexistence regime, populations may experience stochastic extinction (main text Figure 1F). Simulations are run for a fixed number of generations (see section 2.2) or until bacteria or phage go extinct. Population extinction on the timescale of our simulation run-time is restricted to low values of $C_0$ (no phage population extinctions happen for $C_0 > 3000$, and no bacteria extinctions happen for $C_0 > 300$). Phages are prone to population extinction at higher population sizes than bacteria because of their large burst size which makes their dynamics more noisy (see section 5.3.1).

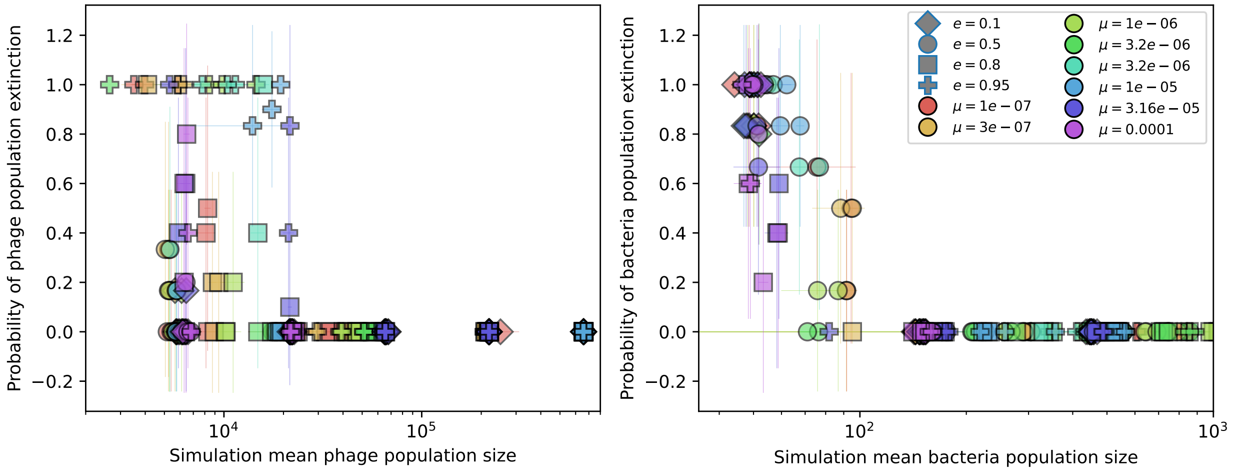

Figure 11: Probability of extinction in 4 or more simulations with the same parameters vs. mean phage population size (left) and mean bacteria population size (right) for the lowest value of spacer acquisition probability ($\eta = 10^{-5}$). Colours indicate mutation rate $\mu$ and shapes indicate CRISPR effectiveness $e$.

Extinction probability is strongly related to total population size, with smaller populations being more likely to go extinct across all parameters (Figures 11 and 12). However, there are differences that depend on parameters and initial conditions. First, high spacer effectiveness increases the likelihood of phage extinction and decreases the likelihood of bacterial extinction (Figure 11). High spacer effectiveness correspondingly increases the time to extinction for bacteria (Figure 14) and decreases the time to extinction for phages (Figure 13). Bacteria also appear to be less likely to go extinct at high values of $\eta$ (Figure 14). Phages have longer times to extinction and lower extinction probability at high values of $\mu$ (Figure 13). And finally, phages have a longer time to extinction if the initial number of phage clones ($m_{init}$) is high (Figure 16), but this effect disappears at low $\eta$ (Figure 15), perhaps because bacteria do not acquire spacers quickly enough for the initial phage diversity to make a difference before the number of clones equilibrates, which happens rapidly at low total population size (see upper rows of Figure 1).

### 4 Simulation results

#### 4.1 Measuring diversity

A measure for the overall sequence diversity in the population is the total number of unique clones. In general the bacterial diversity and phage diversity need not be the same, but it turns out that the number of bacterial clones (which we call $m$) closely tracks the number of "large" phage clones across a wide range of parameters.

To measure the number of large phage clones in a simulation, we scale the observed phage clone size distribution by the probability of extinction for each clone size below a size cutoff given by equation 91. We multiply each observed number of clones of size $n$ by the probability of extinction (equation 134) to the power of the clone size $n$ (equation 23): this gives a scaling factor for each clone size that predicts how many clones of that size will survive at long times (equation 24). $P_n^{\text{large}}$ is plotted alongside the full normalized histogram of clone sizes for one simulation in Figure 23.

$$1 - P_0(n) = 1 - \left(1 - \frac{2s_0}{B(s_0 + \delta_0)}\right)^n \tag{23}$$

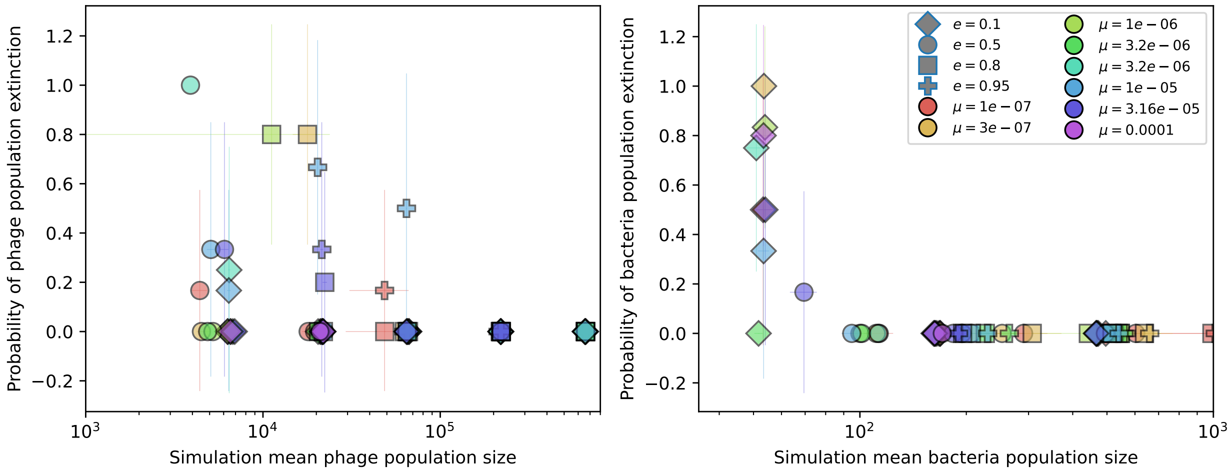

Figure 12: Probability of extinction in 4 or more simulations with the same parameters vs. mean phage population size (left) and mean bacteria population size (right) for the highest value of spacer acquisition probability ($\eta = 10^{-2}$). Colours indicate mutation rate $\mu$ and shapes indicate CRISPR effectiveness $e$.

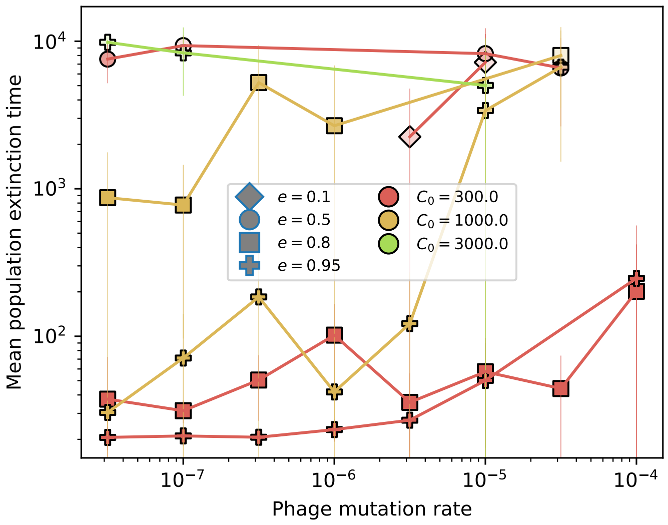

Figure 13: Mean time for the phage population to go extinct vs. phage mutation rate $\mu$ across simulations where at least one simulation experienced phage extinction. The darkness of each point indicates the fraction of simulations that went extinct with darkest colours representing all simulations extinct. Simulations are shown for $\eta = 10^{-2}$.

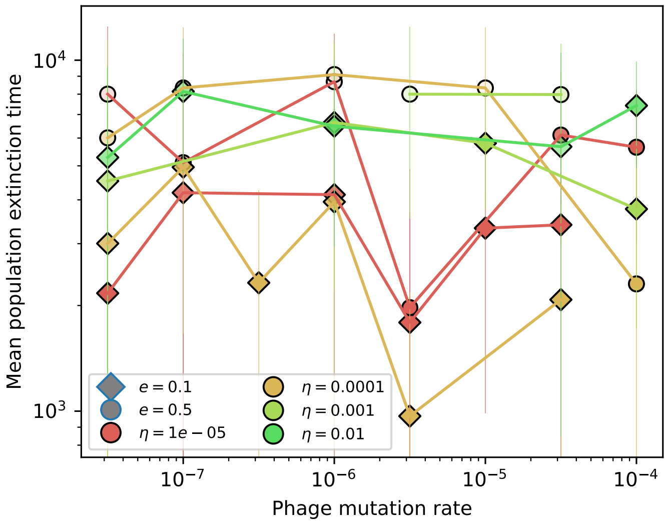

Figure 14: Mean time for the bacteria population to go extinct vs. phage mutation rate $\mu$ across simulations where at least one simulation experienced bacteria extinction. The darkness of each point indicates the fraction of simulations that went extinct with darkest colours representing all simulations extinct.

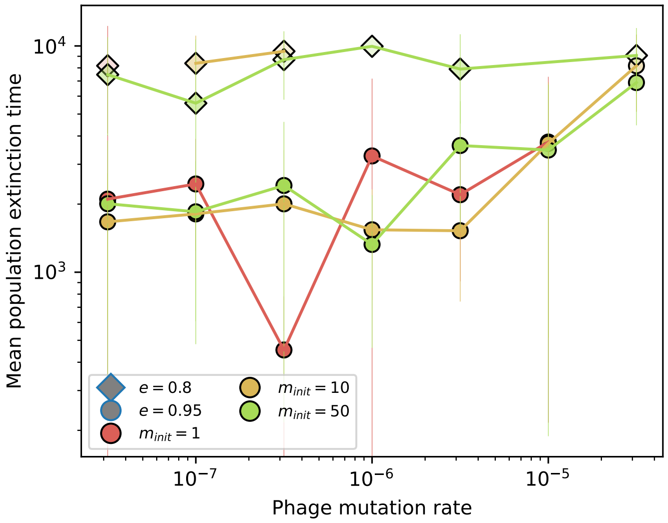

Figure 15: Mean time for the phage population to go extinct vs. phage mutation rate $\mu$ across simulations where at least one simulation experienced phage extinction. The darkness of each point indicates the fraction of simulations that went extinct with darkest colours representing all simulations extinct. Colours represent number of initial phage clones $m_{init}$. Simulations are shown for $\eta = 10^{-5}$ and $C_0 = 1000$.

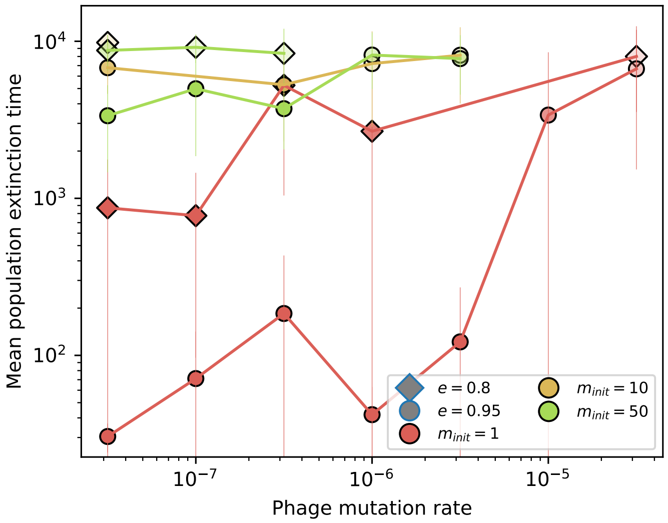

Figure 16: Mean time for the phage population to go extinct vs. phage mutation rate $\mu$ across simulations where at least one simulation experienced phage extinction. The darkness of each point indicates the fraction of simulations that went extinct with darkest colours representing all simulations extinct. Colours represent number of initial phage clones $m_{init}$. Simulations are shown for $\eta = 10^{-2}$ and $C_0 = 1000$.

$$P_n^{\text{large}} = \begin{cases} P_n(1 - P_0(n)) & \text{if } n < N_{\text{est}} \\ P_n & \text{if } n \geq N_{\text{est}} \end{cases} \tag{24}$$

We then calculate the number of large phage clones at steady-state as $N_{\text{total}} \times \sum_n P_n^{\text{large}}$, the total number of phage clones multiplied by the fraction of clones that survive for long times. Note that $P_n^{\text{large}}$ is not normalized; it is a quantity $\leq 1$ representing the proportion of the clone size distribution that corresponds to large clones. The predicted number of large phage clones is plotted against the observed mean number of bacterial clones in Figure 24. There is extremely good agreement at all parameters except the lowest value of $\eta$, $\eta = 10^{-5}$, where the estimated number of large phage clones tends to be larger than $m$. In this regime, phage clones go extinct because of clonal interference before bacteria are able to acquire spacers (Figure 18). At low $\eta$ and high $\mu$, phage clone fitness declines more rapidly with less influence from bacterial spacer acquisition than at high $\eta$ and moderate $\mu$: Figure 19 shows the ratio of phage clone initial fitness to phage clone fitness at the mean bacteria spacer acquisition time, measured from simulations, vs $\eta/\mu$. At most parameters, however, phage clone fitness has not changed much by the time bacteria are acquiring spacers.

There is an $\eta$-dependent "floor" on the minimum size a phage clone must be before bacteria begin to acquire spacers (SI Figure 17). We can estimate the size of a phage clone at the time of first spacer acquisition by calculating the mean time of acquisition from an exponentially growing population. Let $n(t)$ be the size of a phage clone relative to the time of mutation. We assume phage clones grow exponentially with rate $s_0$ at early times, where $s_0 = \alpha n_B(Bp_V - 1) - F$ is the average initial growth rate of phage clones: $n(t) = \mathrm{e}^{s_0 t}$.

The probability that no acquisitions have happened by time $t$ is given by $P_0$ in a Poisson process. Let $a = \alpha\eta(1 - p_V)n_B^0$ be the rate of spacer acquisition (see equation 7), then the probability of no acquisitions by time $t$ is $P_0(t) = \mathrm{e}^{-a\int_0^t n(t')dt'} = \mathrm{e}^{-\frac{a}{s_0}(\mathrm{e}^{s_0 t} - 1)}$. Now, the probability that an acquisition happens between time $t$ and $t + dt$ is then

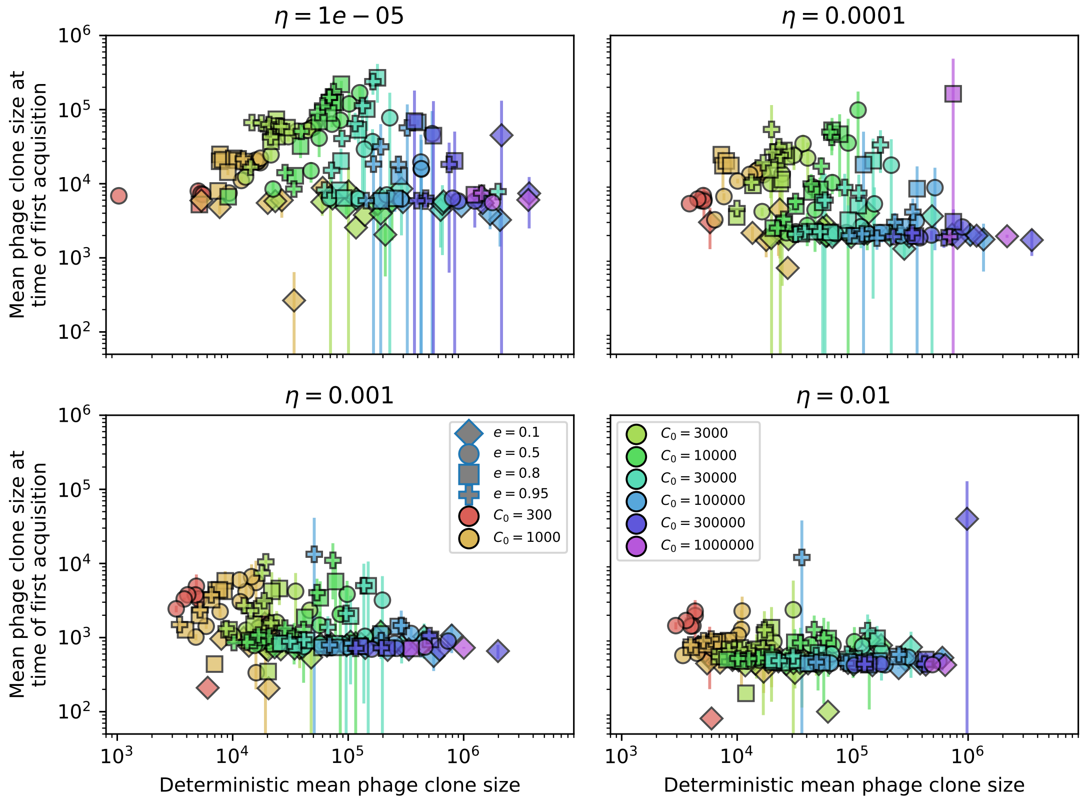

Figure 17: Mean phage clone size at the time of first spacer acquisition vs. the deterministic mean phage clone size. For each phage clone trajectory, the clone size at the time of first spacer acquisition is recorded and these are averaged across each simulation.

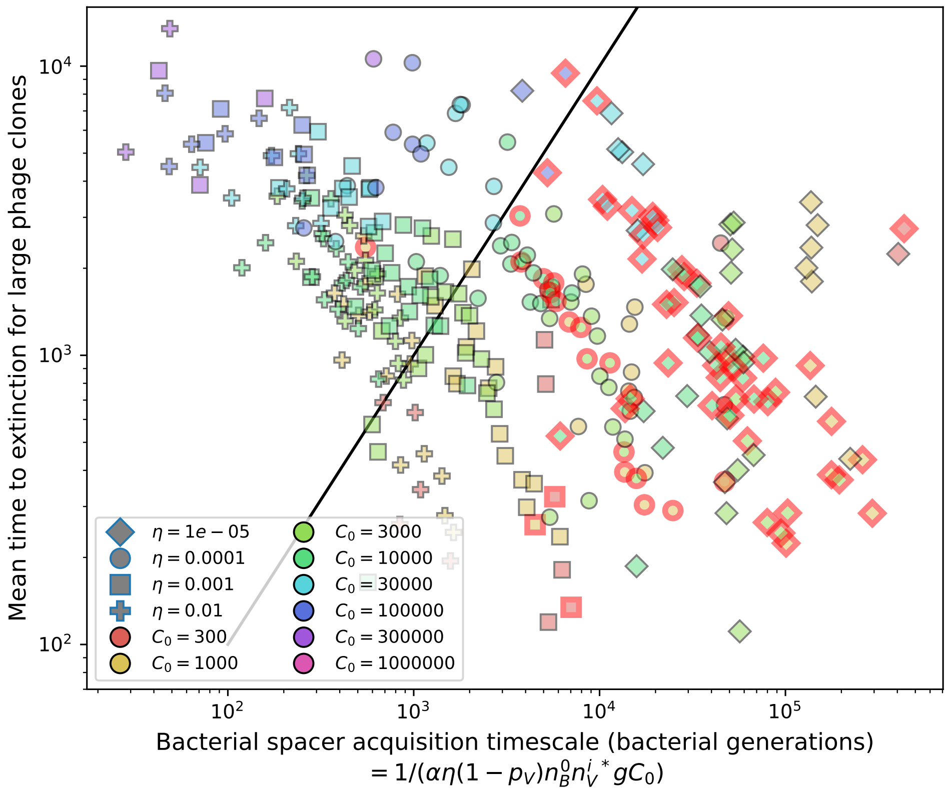

Figure 18: Mean time to extinction for phage clones vs. the timescale of bacteria spacer acquisition given by $1/D$ where $D = \alpha\eta(1-p_V){n_V^i}^* n_B^0 g C_0$. Points outlined in red are simulations where the ratio of large phage clones to bacterial clones exceeds 1.2. Phage clones experience clonal inteference at low $\eta$: they go extinct faster than bacteria acquire spacers.

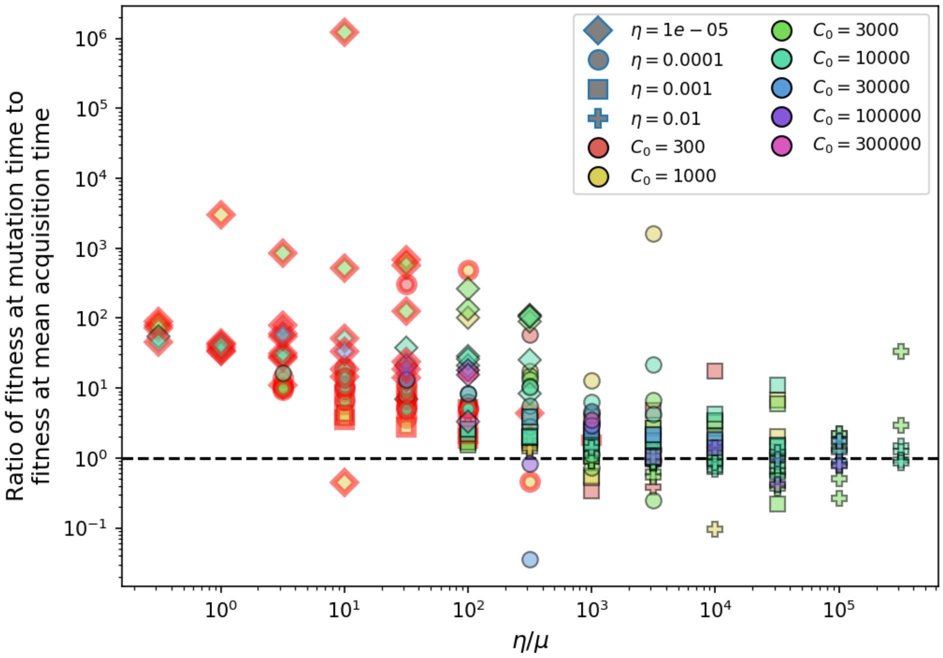

Figure 19: Ratio of average phage clone initial fitness to the average phage clone fitness at the mean bacterial spacer acquisition time vs. $\eta/\mu$. Points outlined in red are simulations where the ratio of large phage clones to bacterial clones exceeds 1.2. The phage fitness is the average per capita growth rate of phage clones conditioned on survival. Phage clones experience clonal inteference at low $\eta$ and high $\mu$.

$$P(t) = P_0(t)an(t) = ae^{s_0 t}e^{-\frac{a}{s_0}(e^{s_0 t}-1)} \tag{25}$$

Equation 25 is shown as a function of $t$ for four simulations with different values of $\eta$ in Figure 20. The probability has a sharp peak: there is a particular time related to phage clone growth at which the first acquisition is most likely — intuitively, this is what leads tot he appearance of a sharp floor in phage clone size at first acquisition. Interestingly, the mean time of first acquisition is non-monotonic in $\eta$: the mean time is smallest for the lowest and highest values of $\eta$.

We calculate the mean time from equation 25:

$$\langle t \rangle = \int_0^\infty ae^{s_0 t}e^{-\frac{a}{s_0}(e^{s_0 t}-1)}t dt \tag{26}$$

$$\langle t \rangle = \frac{1}{s_0}e^{\frac{a}{s_0}}\Gamma(0, \frac{a}{s_0}) \tag{27}$$

where $\Gamma(0, \frac{a}{s_0}) = \int_{\frac{a}{s_0}}^\infty \frac{e^{-t}}{t}dt$ is the incomplete gamma function. We can plug this time into $n(t) = e^{s_0 t}$ to estimate the mean phage clone size at first acquisition; the measured clone size is plotted as a function of this prediction in Figure 22. The clone size depends on the phage growth rate and bacterial acquisition rate, which are themselves primarily dependent on the total bacteria population size and the acquisition probability parameter $\eta$. If $a << s_0$ (true for our parameters), then $e^{\frac{a}{s_0}} \approx 1$ and $\Gamma(0, \frac{a}{s_0}) > 1$, meaning that the time of first acquisition is larger than $1/s_0$. ($\Gamma(0, \frac{a}{s_0}) > 1$ for $\frac{a}{s_0} \lessapprox 0.25$.) Since $1/s_0$ is the approximate time at which phages are safe from stochastic extinction due to drift (see section 5.0.1), this means that phage clones are out of the stochastic extinction regime by the time bacteria begin to acquire spacers.

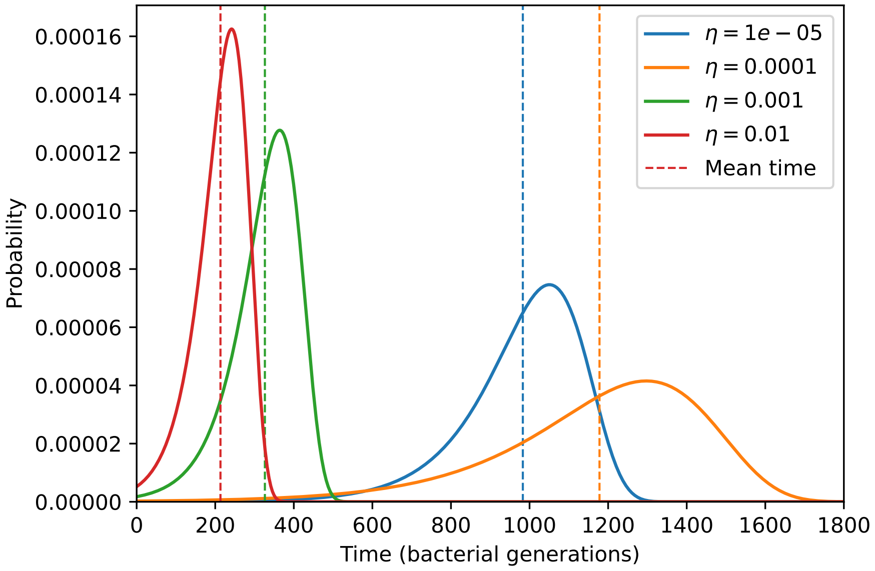

Figure 20: Probability of the first spacer acquisition happening at time $t$ for four simulations with different values of $\eta$ and $C_0 = 10^4, \mu = 10^{-5}$, and $e = 0.95$. The mean of each distribution is shown as a vertical dashed line.

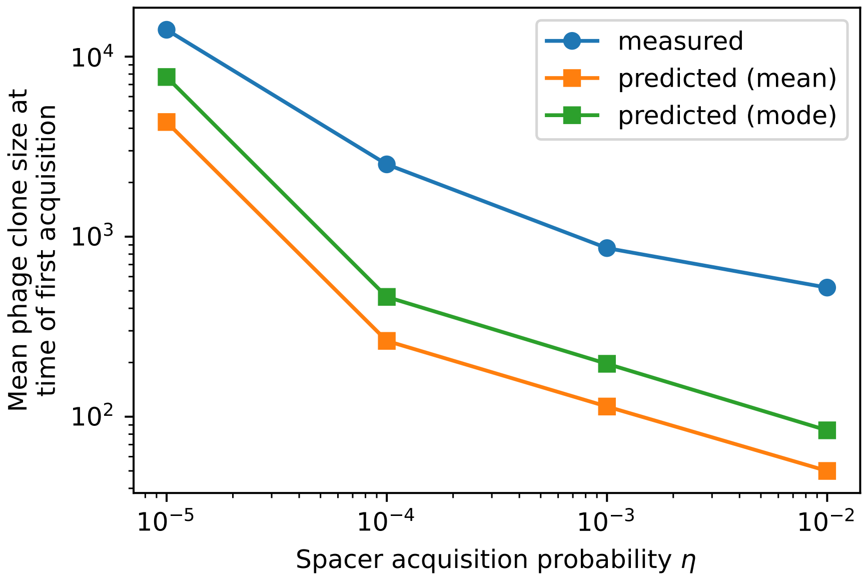

Figure 21: Mean phage clone size at time of first spacer acquisition for simulation data, the predicted with equation 27, and the prediction with the mode of the distribution given by equation 25 for four simulations with different values of $\eta$ and $C_0 = 10^4, \mu = 10^{-5}$, and $e = 0.95$.

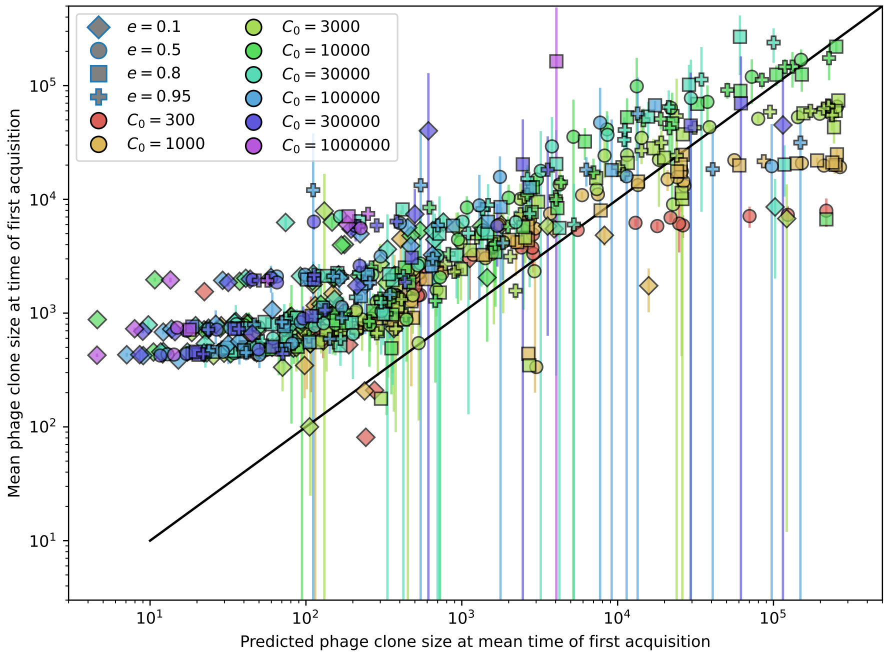

Figure 22: Measured mean phage clone size at the time of first spacer acquisition vs. the prediction given by $e^{s_0\langle t\rangle}$ of equation 27.

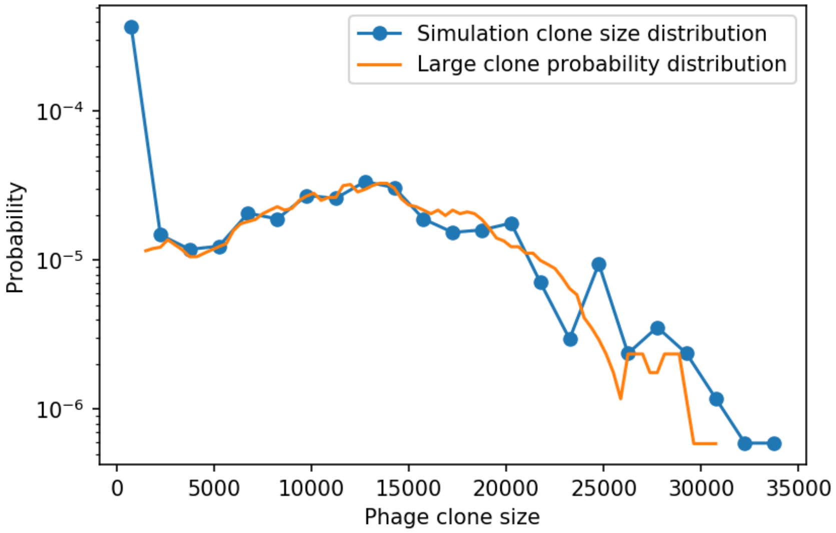

Figure 23: Phage clone size distribution from 15 combined time points for a simulation with the parameters $C_0 = 10^4$, $e = 0.95$, $\eta = 0.01$, and $\mu = 3 \times 10^{-6}$. The blue points are the values of the full normalized phage clone size histogram with a bin width of 1500. The orange line is given by $P_n^{\text{large}}$ in equation 24 smoothed with a running average of window size 3000. Both distributions are scaled by the total number of phage clones.

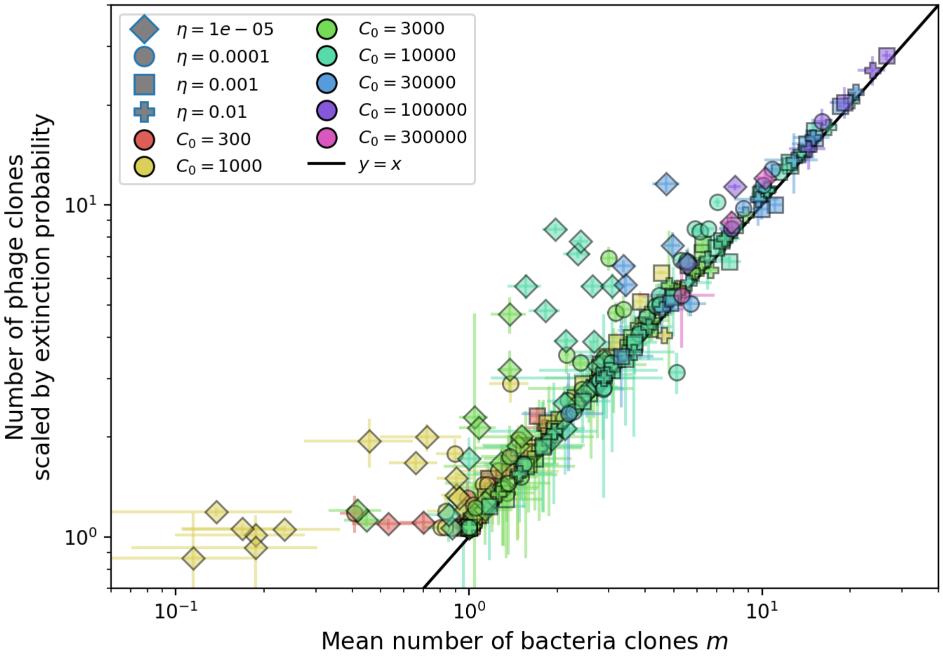

Figure 24: Mean number of large phage clones vs. mean number of bacterial clones in simulations. For each simulation, we take a subset of 15 evenly-spaced timepoints at steady-state and calculate the size and number of phage clones present. We scale the observed clone sized distribution with equation 24 and calculate the mean number of large phage clones by multiplying the total number of clones with the fraction of large phage clones given by $\sum_n P_n^{\text{large}}$. We use the simulation mean total population sizes to calculate $s_0$ and $\delta_0$ in equation 23. We obtain the mean number of bacterial clones by averaging the number of clones present at 15 evenly-spaced timepoints at steady-state. Error bars are the standard deviation across three or more independent simulations.

The definition of large phage clones we just described depends on the measured simulation distribution of phage clone sizes. We can instead approximate the number of large phage clones as the product of the rate at which phage clones become established and the rate at which large phage clones go extinct. This statement is summarized in equation 28. Numerically solving this equation for $m$ gives a prediction for the total number of bacterial clones and total number of large phage clones.

$$m = \underbrace{\left[\frac{2s_0}{B(s_0+\delta_0)}\right]}_{\text{phage establishment fraction}} \underbrace{\alpha B(1-\mathrm{e}^{-\mu L})p_V n_V n_B (1-\frac{e\nu}{m})\frac{1}{gC_0}}_{\text{phage mutation rate}} \underbrace{\frac{2n_V^i(1-\ln\frac{n_V^i}{n_V})gC_0}{(B-1)^2\beta+\delta}}_{\text{large phage clone time to extinction}} \tag{28}$$

$s_0 = \beta_0(B-1) - \delta_0 = \alpha n_B(Bp_V - 1) - F$
$n_V^i$ = deterministic mean phage clone size at steady state
$\beta_0 = n_B \alpha p_V$
$\delta_0 = F + \alpha n_B(1-p_V)$
$\beta = n_B\alpha p_V - \alpha p_V e n_B^i$
$\delta = F + \alpha n_B(1-p_V) + \alpha p_V e n_B^i$

We now describe each of the three terms in equation 28. The phage establishment rate is the phage mutation rate multiplied by the fraction of new phage mutants which become established. Derivation of the phage establishment fraction and phage time to extinction can be found in Section 5.3.1 (equation 133). Derivation of the mean time to extinction for large phage clones can be found in Section 5.3.4 (equation 146).

The phage mutation rate is the mean-field phage reproduction rate $\alpha B n_V(p_V^a n_B^s + p_V n_B^0)$ multiplied by the probability of one or more mutations per burst $(1-\mathrm{e}^{-\mu L})$.

We assume that $p_V^a = p_V(1-e/m)$, which is true if all clones are equal in size (i.e. $n_V^i = n_V/m$ and $n_B^i = n_B^s/m$) or if the deviations from equal size are uncorrelated between matching bacteria and phage clones. This assumption means that the average immunity in the population is approximately $e/m$ which is accurate across a wide range of parameters: Figure 25 compares the full average immunity (equation 29) with $e/m$, where $m$ is the number of bacterial clones present at a given time point in a simulation. The assumption breaks down at low $\eta$ and high $\mu$ (points below the line in the upper right). Intuitively this happens when the sizes of matching clones become anti-correlated because bacteria acquire few spacers while phages acquire many mutations. The resulting matching pairs have more mismatched clone sizes than the mean number of bacterial clones would suggest. (In this particular case, phage clones are smaller than $n_V/m$.) Conversely, at large population sizes and large spacer acquisition rates, matching clone sizes become correlated, leading to average immunity $> e/m$. Figure 26 shows clone size distributions for four simulations with increasing $\eta$, showing that as $\eta$ increases the phage clone distribution becomes more narrow and the matching clone pairs become more correlated.

$$\text{Average immunity} = 1 - \frac{\sum_{i,j} n_B^i n_V^j p_V(i,j)}{p_V \sum_{i,j} n_B^i n_V^j} = \frac{e}{n_B^s n_V}\sum_i n_V^i n_B^i \tag{29}$$

For several parameter combinations, there is no $m$ that satisfies equation 28. This generally occurs for parameters resulting in small total population sizes. In these cases, no solution is found because the predicted mutation rate and/or the predicted establishment fraction and/or the predicted mean time to extinction for large clones are too low (Figure 29).

##### 4.1.1 Relationship of average immunity to Morisita-Horn index

In certain limits, average immunity can be related to the Morisita-Horn similarity index $\Psi$ [23, 17], where instead of comparing two populations from different timepoints or different regions, we are comparing

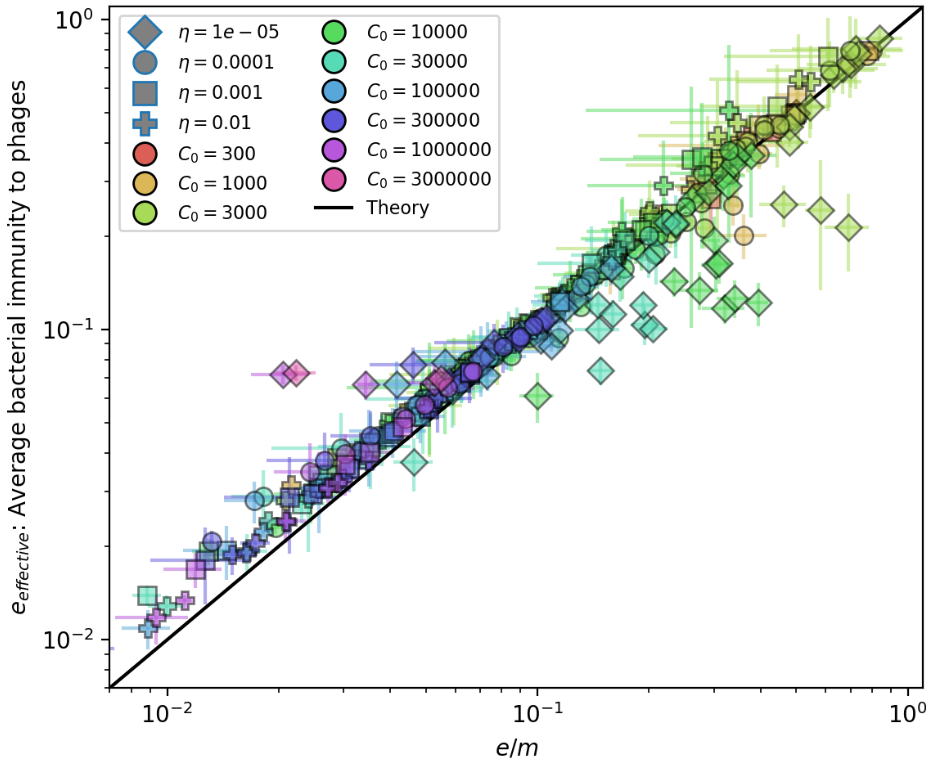

Figure 25: Effective $e = \frac{e}{n_B^s n_V} \sum_i n_V^i n_B^i$ vs. $e/m$ across all simulations where $m \geq 1$ on average. Error bars are the standard deviation across 3 or more independent simulations. The solid black line is $y = x$.

bacteria and phage populations scaled by the immune benefit of CRISPR.

If we assume that a matching spacer provides perfect immunity ($e = 1$) and there is no cross-reactivity, then $p_V(i,j) = p_V(1 - \delta_{ij})$ and average immunity is given by

$$1 - \frac{\sum_{i,j} n_B^i n_V^j p_V(i,j)}{p_V \sum_{i,j} n_B^i n_V^j} = 1 - \frac{p_V \sum_{i,j} n_B^i n_V^j (1 - \delta_{ij})}{p_V \sum_{i,j} n_B^i n_V^j} = \frac{\sum_i n_B^i n_V^i}{n_B^s n_V} \tag{30}$$

This is the Morisita overlap index between bacteria and phage without the factor of $\frac{2}{D_B + D_V}$, where $D_B$ and $D_V$ are the Simpson's diversity indices for bacteria with spacers and phages respectively. In the limit that all phage and bacteria clones are equal sizes ($n_B^i \approx n_B^s/m$, $n_V^i \approx n_V/m$), the factor $\frac{2}{D_B+D_V} \approx m$, so we have $\Psi \approx m e_{eff}$. We found in most simulations that effective $e \approx e/m$, which implies that the Morisita overlap between bacteria and phage is constant across all parameters we study. A constant Morisita overlap implies that the population diversity evolves to attain the highest possible overlap. The full average immunity includes pairwise comparisons between all species and may be quite different from the Morisita overlap index.

#### 4.2 Analytic approximations for diversity

We want to understand more intuitively how diversity depends on system parameters. Here we describe analytic approximations for diversity $m$ and its components. Equation 28 contains $m$ both explicitly and implicitly, so we approximate each of the three components to arrive at an analytic approximation for $m$.

We can make some initial simplifications by cancelling terms (note that $s_0 + \delta_0 = \beta_0(B - 1)$ and $\beta_0 = n_B \alpha p_V$:

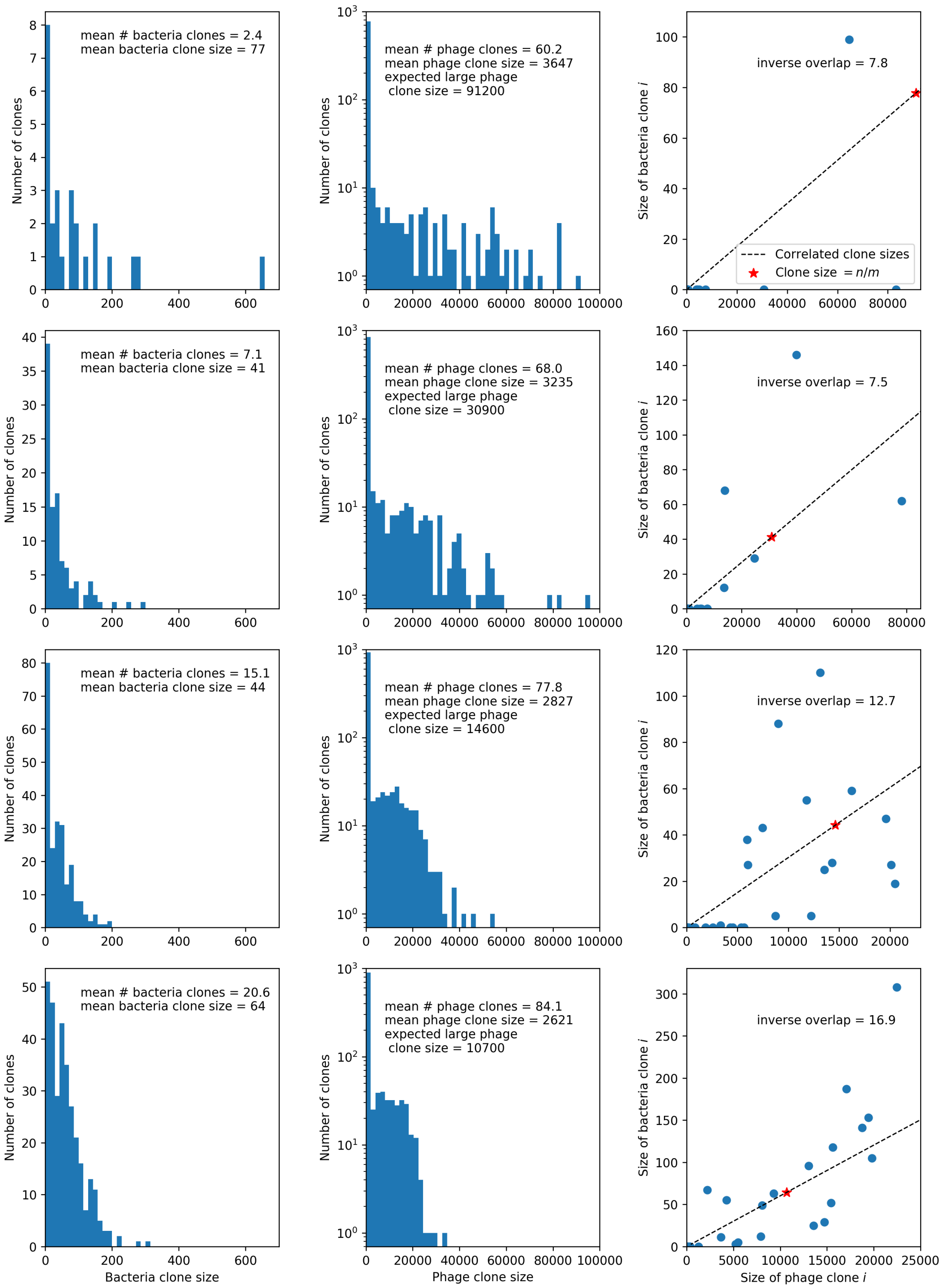

Figure 26: Four simulations with $C_0 = 10000$, $\mu = 10^{-5}$, and $e = 0.95$. From top to bottom, $\eta$ increases by a factor of 10 in each row, from $\eta = 10^{-5}$ in the top row to $\eta = 10^{-2}$ in the bottom row. The first two columns show clone size distributions combined from 15 time points between 2000 and 10000 bacterial generations. Bacteria are in the left column and phages in the middle column. The third column shows the pairwise clone sizes of matching clones at the last sampled time point (9467 generations). The expected large phage clone size is the total phage population divided by the mean number of bacterial clones. The inverse overlap is $\frac{e}{e_{\text{eff}}} = \frac{n_B^s n_V}{\sum_i n_V^i n_B^i}$, which we assume is $\approx m$ as shown in Figure 25. The dashed line in the third column indicates the line that clone size pairs would fall on if they were perfectly correlated, and the red star indicates the mean large clone size for phages and the mean clone size for bacteria.

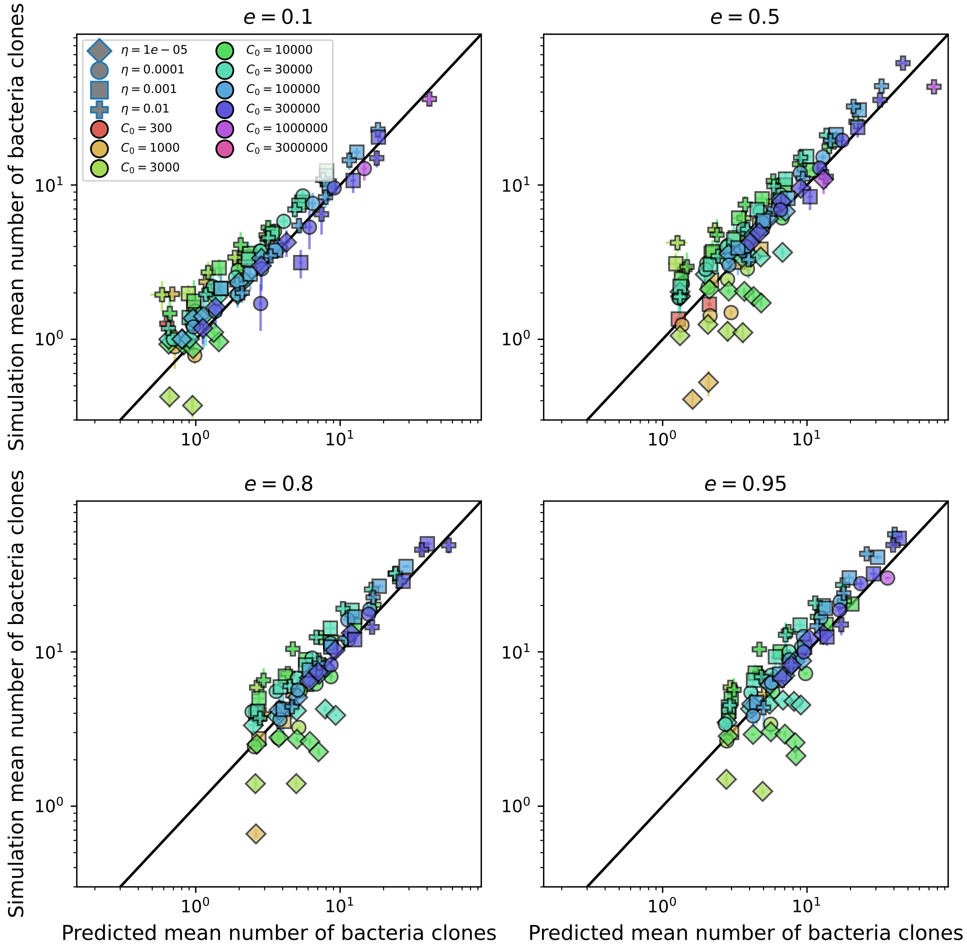

Figure 27: Simulation mean number of bacteria clones ($m$) vs. theoretical prediction for $m$ given by numerically solving equation 28, broken down by different values of $e$.

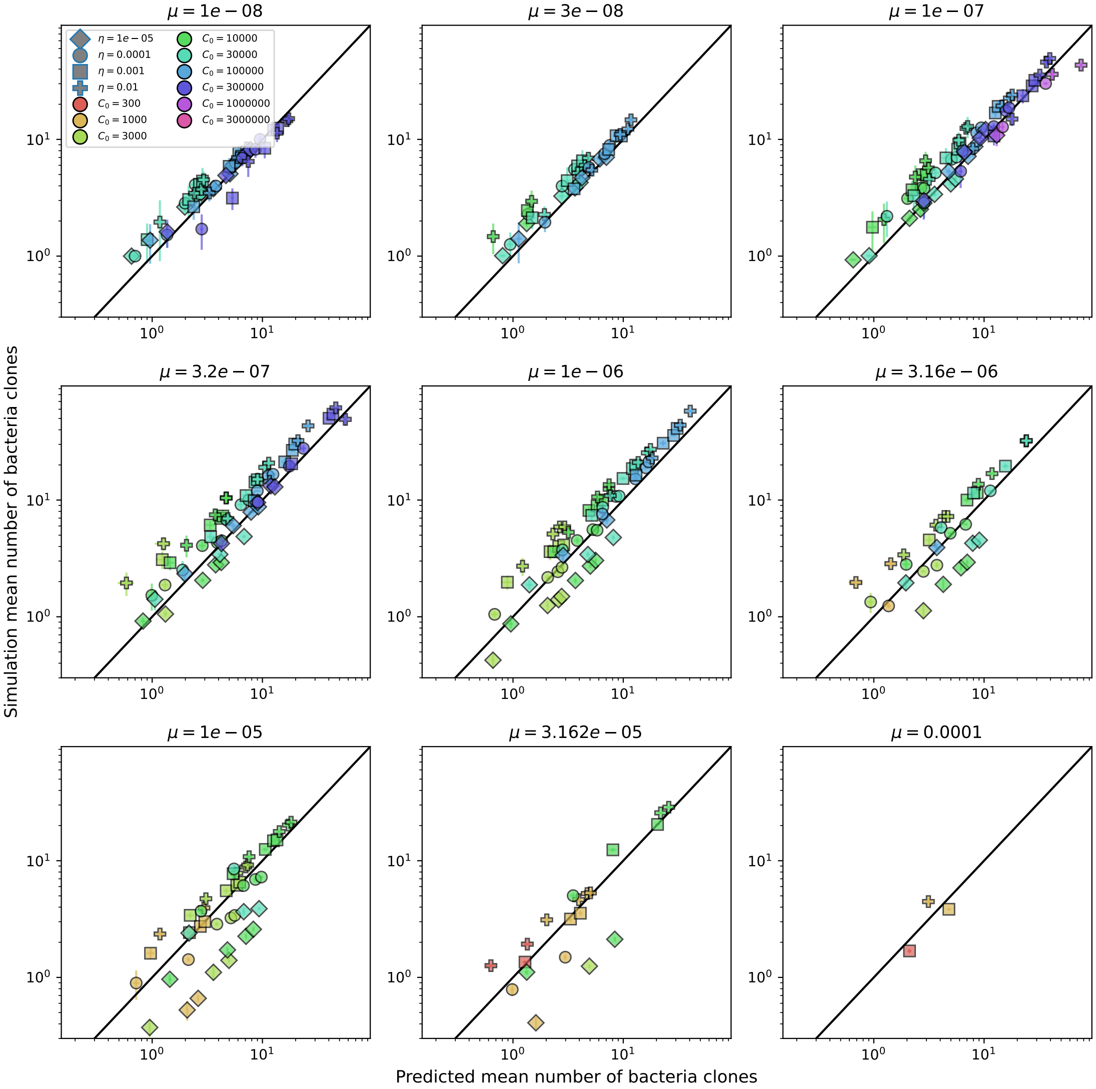

Figure 28: Simulation mean number of bacteria clones ($m$) vs. theoretical prediction for $m$ given by numerically solving equation 28, broken down by different values of $\mu$.

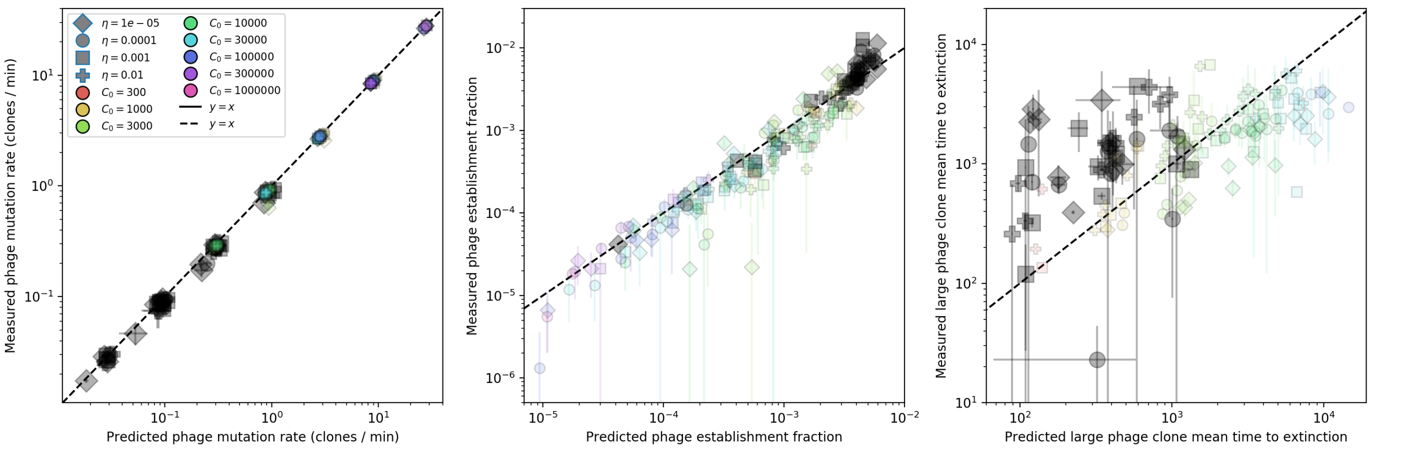

Figure 29: Measured simulation phage clone mutation rate, establishment fraction, and mean time to extinction as a function of the theoretical prediction for each. Highlighted in grey are parameter combinations for which no theoretically predicted $m$ could be determined; the predicted quantity is instead calculated with the simulation mean $m$.

$$m = \left(\frac{2s_0}{B-1}\right)(1-\mathrm{e}^{-\mu L})n_V(1-\frac{e\nu}{m})\frac{2n_V^i(1-\ln\frac{n_V^i}{n_V})}{(B-1)^2\beta+\delta} \tag{31}$$

Now expanding $s_0$ and the denominator and cancelling and collecting terms:

$$m = \frac{4(\alpha n_B(Bp_V-1)-F)(1-\mathrm{e}^{-\mu L})n_V n_V^i(1-\ln\frac{n_V^i}{n_V})}{(B-1)(B(B-2)\beta+\alpha n_B+F)}(1-\frac{e\nu}{m}) \tag{32}$$

In theory, $n_V^i \approx n_V/m$ and $n_B^i \approx n_B^s/m$. To see this, we start with the solutions for the deterministic mean bacteria and phage clone sizes (equations 17 and 18, reprinted here):

$$n_B^{i\,*} = \frac{1}{e}\left(n_B - \frac{F+\alpha n_B}{\alpha B P_0 p_V}\right) \tag{33}$$

$$n_V^{i\,*} = \frac{n_B^{i\,*}(\alpha p_V n_V - (gC-F-r))}{\alpha\eta n_B^0(1-p_V)+\alpha p_V e n_B^{i\,*}} \tag{34}$$

From the deterministic mean-field equation for $n_V$ (equation 6), we have that $F+\alpha n_B = \alpha B p_V n_B(1-\frac{e\nu}{m})$. Substituting this in equation 33, we get

$$n_B^{i\,*} = \frac{1}{e}\left(n_B - \frac{\alpha B p_V n_B(1-\frac{e\nu}{m})}{\alpha B P_0 p_V}\right) = \frac{n_B}{e}\left(1-\frac{(1-\frac{e\nu}{m})}{P_0}\right) \tag{35}$$

We can neglect $P_0$, both because it is always close to 1 and because, at steady state, the effect of mutants leaving type $i$ is partially balanced by mutants entering type $i$.

$$n_B^{i\,*} \approx \frac{n_B\nu}{m} = \frac{n_B^s}{m} \tag{36}$$

Similarly, from the deterministic mean-field equation for $n_B^s$ (equation 5), we have $gC-F-r = \alpha n_V p_V\left(1-\frac{e}{m}\right) - \alpha n_V(1-p_V)\eta\frac{n_B^0}{n_B^s}$. Substituting this in equation 34, we get

$$n_V^{i\,*} = \frac{n_B^{i\,*}(\alpha p_V n_V - (\alpha n_V p_V\left(1-\frac{e}{m}\right) - \alpha n_V(1-p_V)\eta\frac{n_B^0}{n_B^s}))}{\alpha\eta n_B^0(1-p_V)+\alpha p_V e n_B^{i\,*}} \tag{37}$$

Substituting $n_B^0 = n_B(1-\nu)$ and $n_B^s = n_B\nu$, we find ${n_V^i}^* = n_V/m$.
Replacing $n_V^i$ and $n_B^i$ with $n_V/m$ and $n_B^s/m$ respectively:

$$m = \frac{4(\alpha n_B(Bp_V - 1) - F)(1 - \mathrm{e}^{-\mu L})\frac{n_V^2}{m}(1 - \ln\frac{1}{m})}{(B-1)(B(B-2)\alpha p_V n_B(1-\frac{e\nu}{m}) + \alpha n_B + F)}(1 - \frac{e\nu}{m}) \tag{38}$$

Now we want to solve equation 38 for $m$. This is difficult because the total population sizes $n_B$, $n_V$, $C$, and $\nu$ also depend on $m$. To find the $m$ dependence of $n_V$, $n_B$, $C$, and $\nu$, we note that we can approximate the steady-state solutions to equations 5 to 9 as the corresponding solutions described in [3] with the simple replacement of the parameter $e$ with $e/m$. This is because $e/m$ is a good approximation for the full average immunity at most parameters (Figure 25), and average immunity enters these equations in place of $e$ in the model without phage mutations. This is analogous to average adaptive immunity replacing the rate of innate immunity in the Lotka-Volterra system described by Iranzo *et al.* [24].

We write the steady-state solutions for equations 6 and 9 below, where we have defined $p_V(i,j)$ as in equation 10 and approximated average immunity as $e/m$. For simplicity we rescale $n_B$ and $n_V$ by $C_0$: $n_V = C_0 y^*$ and $n_B = C_0 x^*$. Here $p = p_V\alpha/g$. These steady-state solutions are shown in Figure 30.

At low average immunity (high diversity), the population behaves as if there were no CRISPR system at all. Figure 30 shows total phage, total bacteria, and the fraction of bacteria with spacers as a function of average immunity (effective $e$). At low average immunity, total population sizes are the same as in a population without CRISPR entirely. Interestingly, the effective no-CRISPR case does not necessarily correspond to no CRISPR spacers: $\nu > 0$ even when effective $e \approx 0$ for these parameters. This is because spacer acquisition and growth of that clone can still happen even if the spacer confers no fitness benefit.

$$x^* = \frac{fp_V}{p}\frac{1}{Bp_V(1-\frac{e\nu^*}{m}) - 1} \tag{39}$$

$$y^* = \frac{(f-1)p(Bp_V(\frac{e\nu^*}{m} - 1) + 1) - fp_V}{p(\frac{e\nu^*}{m} - 1)(p(Bp_V(\frac{e\nu^*}{m} - 1) + 1) - p_V)} \tag{40}$$

We have defined $x^*$ and $y^*$ in terms of $\nu^*$, which is given in the following implicit cubic equation, where $R = r/(gC_0)$:

$$\begin{aligned} 0 = (1-\nu)\left[-p_V\frac{e\nu}{m} - \eta(1-p_V)\right]\left[(1-f)p(p_V B(1-\frac{e\nu}{m}) - 1) - fp_V\right] \\ + R\nu p_V(1-\frac{e\nu}{m})(Bpp_V(1-\frac{e\nu}{m}) - p + p_V) \end{aligned} \tag{41}$$

This cubic equation is analytically solvable (ignoring the dependence of $m$ on $\nu$), but the full solutions in terms of all parameters are cumbersome.

Only one of the three solutions of equation 41 is physical in the parameter range we use (real-valued and properly bounded):

$$\begin{aligned} \nu^* = &-\frac{\left(1+i\sqrt{3}\right)\sqrt[3]{\sqrt{\left(-27a^2d + 9abc - 2b^3\right)^2 + 4\left(3ac - b^2\right)^3} - 27a^2d + 9abc - 2b^3}}{6\sqrt[3]{2}a} \\ &+ \frac{\left(1-i\sqrt{3}\right)\left(3ac - b^2\right)}{3\ 2^{2/3}a\sqrt[3]{\sqrt{\left(-27a^2d + 9abc - 2b^3\right)^2 + 4\left(3ac - b^2\right)^3} - 27a^2d + 9abc - 2b^3}} - \frac{b}{3a} \end{aligned} \tag{42}$$

where the coefficients are

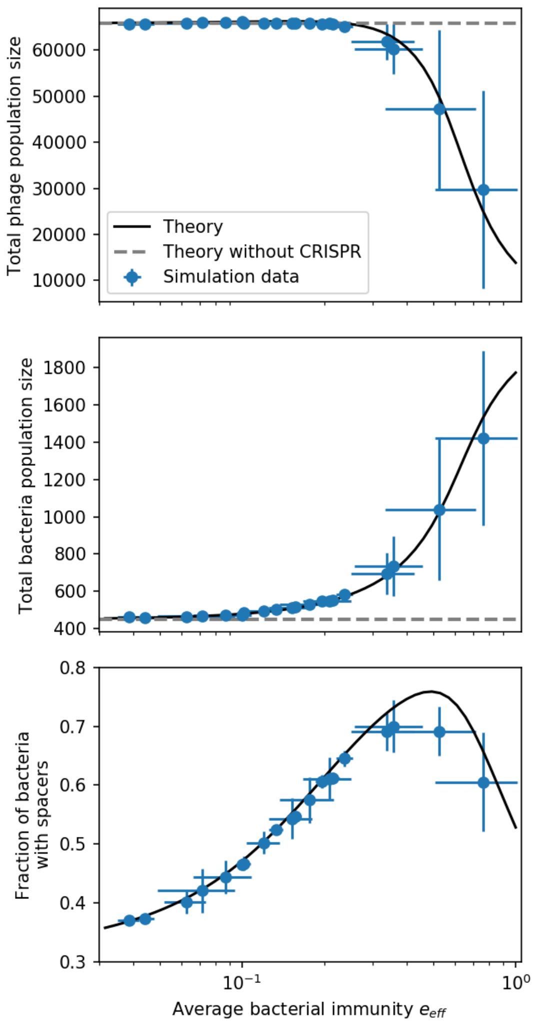

Figure 30: Total phage, total bacteria, and fraction of bacteria with spacers as a function of average immunity (effective $e$). Blue points are simulation results. Error bars are standard deviation across 3 or more independent simulations. The solid black line is the solution given by equations 39 to 41 (from [3]) with the parameter $e$ replaced by effective $e$. The horizontal grey dashed line corresponds to the no-CRISPR ($e = 0$) mean-field solution (derived in [3]).

$$a = B(\frac{e}{m})^2 f p p_V^2 (f + R - 1) \tag{43}$$

$$b = -\frac{e}{m} f p_V (p(f(B(p_V(\frac{e}{m} + \eta + 1) - \eta) - 1) + B(\eta - p_V(\frac{e}{m} + \eta - 2R + 1)) - R + 1) + p_V(f + R)) \tag{44}$$

$$c = fp\left[Bp_V^2(\frac{e}{m}(f-1)(\eta+1) + (f-1)\eta + R) - (\frac{e}{m} - 1)(f-1)p_V(B\eta + 1) - (2B+2)(f-1)\eta p_V + (f-1)\eta - p_V(f+R-1)\right] + fp_V(\frac{e}{m} f p_V - f\eta + p_V(f\eta + R)) \tag{45}$$

$$d = -f\eta(p_V - 1)((f-1)p(Bp_V - 1) + fp_V) \tag{46}$$

##### 4.2.1 Dominant balance approximations for $\nu$

The cubic equation for $\nu$ given by equation 41 can be simplified in certain parameter regimes using dominant balance. We write equation 41 as $a\nu^4 + b\nu^2 + c\nu + d = 0$ with the coefficients as written above. Comparing the numerical values of the coefficients in different parameter regimes, we arrive at Table 4 which outlines three different approximations for $\nu$ obtained by dropping coefficients of the original cubic equation. Broadly speaking, at low $\eta$ and low $e$, both the cubic and quadratic coefficients ($a$ and $b$) are small, so $\nu^* \approx -d/c$, while at high $\eta$ and low $e$ the cubic coefficient ($a$) can be dropped, and at high $e$ and low $\eta$ the constant $d$ can be dropped. Figures 31 to 33 show total population sizes as a function of the three approximations in Table 4.

Table 4: $\nu$ approximations

| | $e_{\text{effective}}$ | | | | |
|---|---|---|---|---|---|
| $\eta$ | $\leq 0.01$ | 0.01 to 0.05 | 0.05 to 0.1 | 0.1 to 0.5 | 0.5 to 1 |
| $\leq 10^{-5}$ | $-\frac{d}{c}$ | $-\frac{d}{c}$ | $-\frac{d}{c}$ | $\frac{-b+\sqrt{b^2-4ac}}{2a}$ | $\frac{-b+\sqrt{b^2-4ac}}{2a}$ |
| $10^{-5}$ to $10^{-4}$ | $-\frac{d}{c}$ | $-\frac{d}{c}$ | $\frac{-c+\sqrt{c^2-4bd}}{2b}$ | $\frac{-b+\sqrt{b^2-4ac}}{2a}$ | |
| $10^{-4}$ to $10^{-2}$ | $-\frac{d}{c}$ | $-\frac{d}{c}$ | $\frac{-c+\sqrt{c^2-4bd}}{2b}$ | | |
| $\geq 10^{-2}$ | $-\frac{d}{c}$ | $\frac{-c+\sqrt{c^2-4bd}}{2b}$ | $\frac{-c+\sqrt{c^2-4bd}}{2b}$ | | |

Note: the drop-$a$ solution also works for below $e = 0.1$ for all values of $\eta$ (since $\frac{2bd}{c}^2 \approx 0$), but the $-d/c$ solution is simpler and so preferred for the very low $e$ range.

For small effective $e$ and small $\eta$, $\nu \approx -d/c$:

$$\nu \approx \frac{\eta(1-p_V)(\alpha(1-f)(Bp_V - 1) - fg)}{\eta(1-p_V)(\alpha(1-f)(B(\frac{e}{m}+1)p_V - 1) - fg) + \alpha p_V(Bp_V - 1)(R - \frac{e}{m}(1-f)) + gp_V(\frac{e}{m}f + R)} \tag{47}$$

We define two combined parameters: $A = \frac{(Bp_V-1)(1-f)\alpha}{fg}$, and $\hat{\eta} = \eta(1-p_V)$. Each of these has an intuitive meaning: $A > 1$ is the stability condition for phage existence, and $\hat{\eta}$ is the probability of spacer acquisition following escape from naive phage predation, so it can be thought of as the overall spacer acquisition probability at the start of an infection. With these variable substitutions, we get

$$\nu \approx \frac{\hat{\eta}(A-1)}{p_V R\left(\frac{1-f+Af}{f(1-f)}\right) + \hat{\eta}(A - 1 + \frac{ABp_V\frac{e}{m}}{(A-1)(Bp_V-1)}) - \frac{e}{m}p_V(A-1)} \tag{48}$$

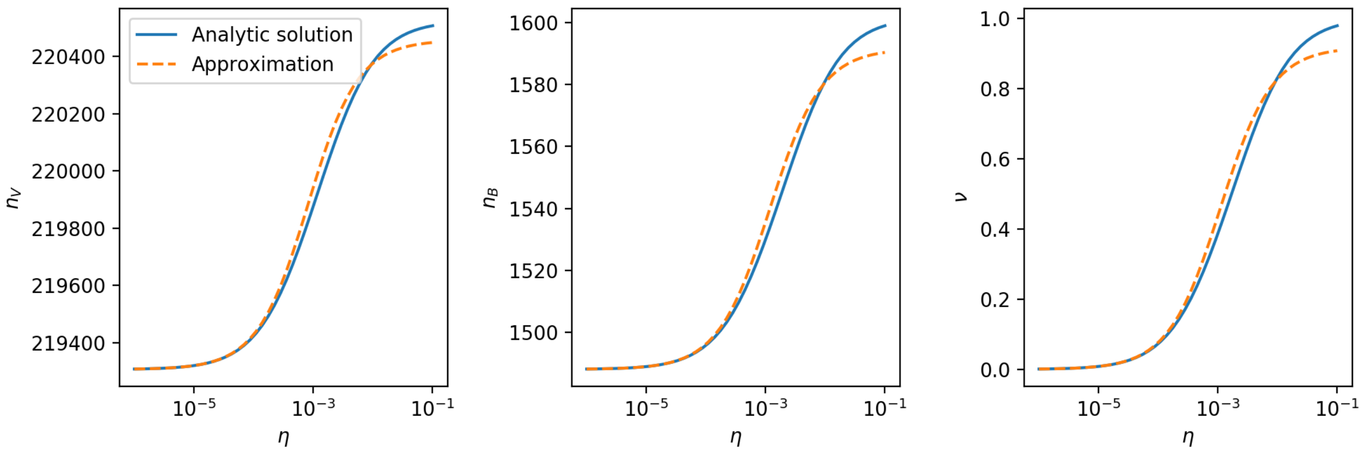

Figure 31: $n_V$, $n_B$, and $\nu$ vs. $\eta$ for $e = 0.05$, approximating $\nu \approx -d/c$.

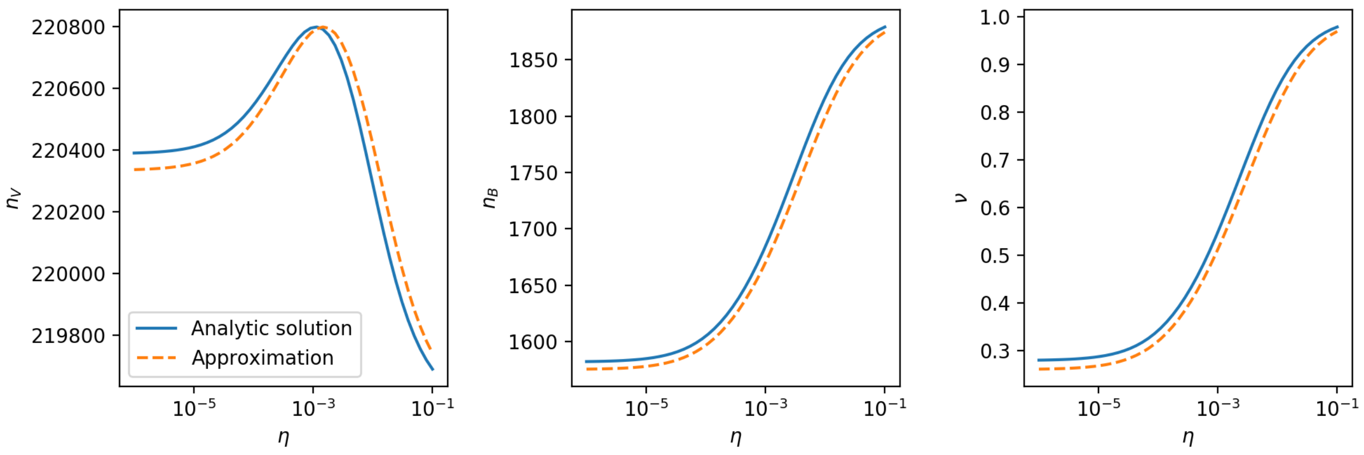

Figure 32: $n_V$, $n_B$, and $\nu$ vs. $\eta$ for $e = 0.15$, approximating $\nu \approx \frac{-c+\sqrt{c^2-4bd}}{2b}$

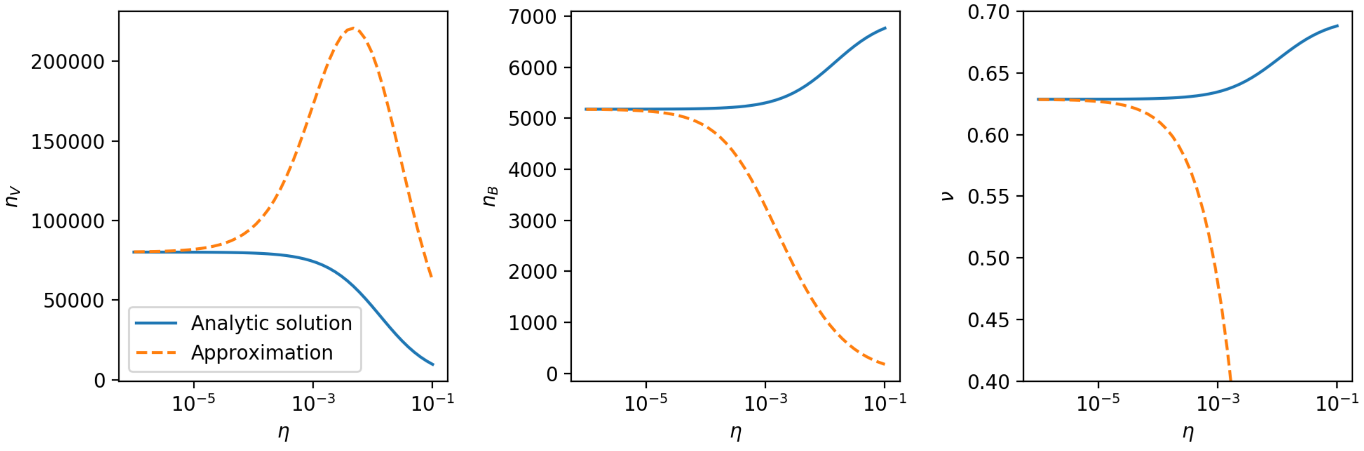

Figure 33: $n_V$, $n_B$, and $\nu$ vs. $\eta$ for $e = 0.8$, approximating $\nu \approx \frac{-b+\sqrt{b^2-4ac}}{2a}$

Now we notice that the deterministic total phage population size when $e = 0$ is given by

$$n_{V\text{no CRISPR}} = \tilde{n_V} = \frac{gC_0 f(1-f)(A-1)}{\alpha p_V(1-f+Af)} \tag{49}$$

We also include $n_B$ for $e = 0$ for completeness:

$$n_{B\text{no CRISPR}} = \tilde{n_B} = \frac{C_0(1-f)}{A} \tag{50}$$

We can replace some terms in $\nu$ with $\tilde{n_V}$ by comparison with equation 49:

$$\nu \approx \frac{1}{1 + \frac{r}{\hat{\eta}\alpha\tilde{n_V}} - \frac{e}{m}\left(\frac{p_V}{\hat{\eta}} - \frac{ABp_V}{(A-1)(Bp_V-1)}\right)} \tag{51}$$

where $r = RgC_0$ is the spacer loss rate per minute. The second term in the denominator is the balance of spacer acquisition and spacer loss per naive bacterium: $\hat{\eta}\alpha\tilde{n_V}$ is the rate of spacer acquisition per naive bacterium in the low average immunity limit.

The third term in the denominator is negative for the parameters we use, but this is not necessarily true in general. It's negative if $\hat{\eta}AB < (A-1)(Bp_V - 1)$, which for the values of $A$ and $Bp_V$ we use (fixed across all simulations) is true for $\eta < 0.0113$.

From this expression for $\nu$ we learn the following:

- $\nu$ decreases if $r$ goes up. More spacer loss means a smaller fraction of bacteria with spacers.
- $\nu$ increases if spacer acquisition goes up (provided $e/m$ is small).
- $\nu$ increases if $\frac{e}{m}$ increases: higher CRISPR effectiveness means higher $\nu$.

For $e/m \to 0$, we define $\tilde{\nu}$:

$$\tilde{\nu} = \frac{1}{1 + \frac{r}{\hat{\eta}\alpha\tilde{n_V}}} \tag{52}$$

Equation 51 expression breaks down at high $\eta$ where the true value of $\nu$ is largely independent of $e$. In this case, when both $\frac{e}{m}$ is small and $\hat{\eta}$ is large, we can drop the third term in the denominator in equation 51).

There is a discontinuity in equation 51 at the value of $e/m$ where the denominator equals zero. This critical value of $e/m$, $\frac{e}{m}^*$, is given by equation 53.

$$\frac{e}{m}^* = \frac{(A-1)(Bp_V-1)(\alpha\hat{\eta}\tilde{n_V} + r)}{\alpha\tilde{n_V}p_V((A-1)(Bp_V-1) - AB\hat{\eta})} \tag{53}$$

Taking a series expansion in $\hat{\eta}$ and keeping the first two terms:

$$\frac{e}{m}^* \approx \frac{r}{\alpha\tilde{n_V}p_V} + \frac{\hat{\eta}}{p_V} + \frac{ABr\hat{\eta}}{\alpha\tilde{n_V}p_V(A-1)(Bp_V-1)} \tag{54}$$

At high $\eta$, the middle term dominates, and at small $\eta$, the first two terms dominate (since the third term is $\approx 9.8\hat{\eta}$ and $1/p_V = 50$). This transition governs the change between the low $e$ and high $e$ regimes; equation 55 is plotted in Figures 37 and 38.

$$\frac{e}{m}^* \approx \frac{r}{\alpha\tilde{n_V}p_V} + \frac{\hat{\eta}}{p_V} \tag{55}$$

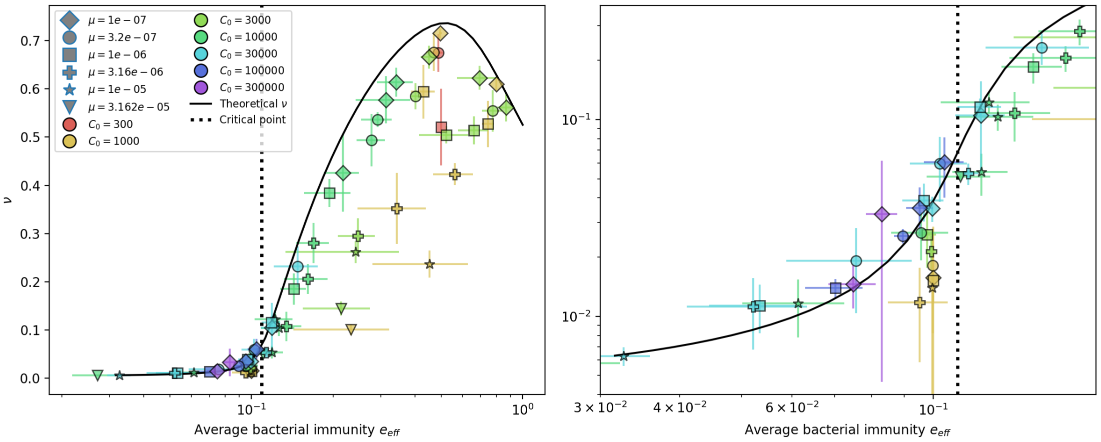

Figure 34: Fraction of bacteria with spacers $\nu$ vs. effective $e$. The solid line is the full numerical theoretical solution. The dashed black line is given by equation 55.

If we substitute this critical value of $e/m$ into the mean-field equations and take $\eta$ to 0, we recover the no-CRISPR mean-field solutions for $n_V$, $n_B$, and $C$. Since this also corresponds to $\nu = 0$, this seems to explain why several simulation values of the phage establishment probability go to 0 around effective $e = 0.1$ for small $\eta$ and small $C_0$ (Figure 34). Despite the presence of spacers with non-zero effectiveness, the population appears to behave as if there is no CRISPR near this point, which removes the fitness advantage of new phage mutants and lowers their establishment probability. We also see large fluctuations in total bacteria population size near this critical point (Figure 35).

Another way to understand the critical point is to look at the difference in relative fitness between bacteria with and without spacers, $n_B^s$ and $n_B^0$:

$$\frac{\dot{n_B^s}}{n_B^s} - \frac{\dot{n_B^0}}{n_B^0} = -r\left(1 + \frac{n_B^s}{n_B^0}\right) + \alpha n_V p_V e_{eff} + \alpha(1-p_V)\eta n_V\left(\frac{n_B^0}{n_B^s} - 1\right) \tag{56}$$

Now if effective $e$ is given by equation 55:

$$\frac{\dot{n_B^s}}{n_B^s} - \frac{\dot{n_B^0}}{n_B^0} = -r\frac{n_B^s}{n_B^0} + \alpha(1-p_V)\eta n_V\frac{n_B^0}{n_B^s} \tag{57}$$

Equation 57 indicates that at the critical value of effective $e$, the fitness difference is exactly given by the balance of spacer acquisition and spacer loss. Below this critical point, there are additional constant negative terms, and for the same values of $n_B^s$ and $n_B^0$, the relative fitness of spacer-containing bacteria is lower than naive bacteria; above this point, it is higher. At the critical point, the immunity boost from spacers exactly cancels out spacer loss. It is interesting that for the same reasons that many parameters collapse onto the same curves as a function of average immunity, the value of this critical point is largely independent of the parameters of CRISPR immunity and occurs at a value of effective $e \approx 10^{-1}$ regardless of $\mu$, $e$, or $C_0$.

##### Large $e$ approximation for $\nu$

In the limit of $e \to 1$ and small $\eta$, we find the following approximate solution for $\nu$ by setting $e = 1$ and taking a series expansion in $\eta$.

$$\nu \approx \frac{\alpha(Bp_V - 1)(f + R - 1) + g(f + R)}{\alpha B p_V(f + R - 1)} + \frac{gR\hat{\eta}}{p_V(f + R - 1)(\alpha(Bp_V - 1)(f + R - 1) + g(f + R))} \tag{58}$$

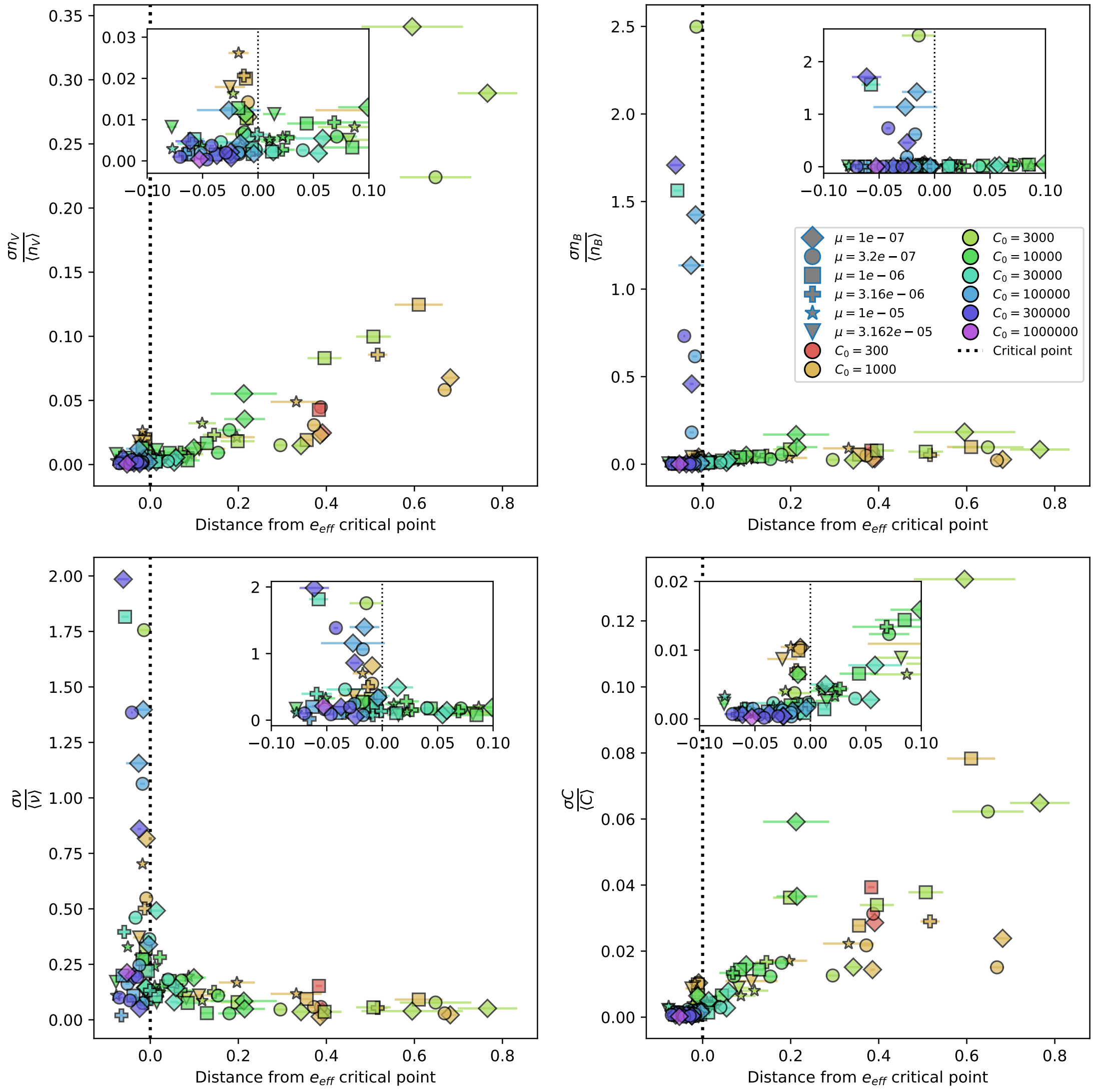

Figure 35: Population standard deviation divided by population mean as a function of the distance from the average immunity critical point given by equation 55. Insets show a smaller x-axis range for the same quantities. Total phage (top left), total bacteria (top right), fraction of bacteria with spacers (bottom left) and total nutrients (bottom right) are plotted.

Equation 58 is plotted in Figure 37 in red. The second term proportional to $\hat{\eta}$ is tiny compared to the first for all values of $\eta$ we use.

##### 4.2.2 Approximation for $T_{\text{ext}}$

If we evaluate the mean phage time to extinction (Section 5.3.4) at the deterministic mean phage clone size, we can approximate $n_V^i \approx \frac{n_V}{m}$ and $n_B^i \approx \frac{n_B^s}{m}$ as before:

$$T_{\text{ext}} \approx \frac{2\frac{n_V}{m}(1+\ln m)gC_0}{B(B-2)\alpha p_V n_B(1-\nu\frac{e}{m}) + F + \alpha n_B} \tag{59}$$

We expand $n_V$ and $n_B$ in powers of $e/m$:

$$n_V = \frac{gC_0 f(1-f)(A-1)}{\alpha p_V(1-f+Af)} + \frac{e\nu}{m}C_0\left[-\frac{fg}{\alpha p_V} - \frac{1-f}{p_V(1-f+Af)} + \frac{ABf(1-f)}{(1-f+Af)^2}\right] + \mathcal{O}(e/m)^2 \tag{60}$$

$$n_B = \frac{C_0(1-f)}{A} + \frac{e\nu}{m}\left[\frac{BFp_V}{\alpha(Bp_V-1)^2}\right] + \mathcal{O}(e/m)^2 \tag{61}$$

Note that $\nu$ never appears without $e/m$ beside it in these expressions, so we will use the 0th order expression for $\nu$ (equation 52) which still allows us to keep $e/m$ to first order elsewhere.

$$\tilde{\nu} = \frac{1}{1+\frac{r}{\hat{\eta}\alpha\tilde{n_V}}} + \mathcal{O}(e/m) \tag{62}$$

Substituting equations 60 to 62 into $T_{\text{ext}}$:

$$\begin{aligned} T_{\text{ext}} &\approx \frac{2(1+\ln m)}{f(B-1)}\frac{\tilde{n_V}}{m}\left(1-\frac{1}{Bp_V}\right) \\ &+ \frac{e}{m^2}(1+\ln m)\left[\frac{2C_0^2 fg\hat{\eta}(1-A)((Bp_V-2)(g-\frac{A^2fg}{1-f}) + \frac{2gA}{1-f}(1+f(Bp_V-2))}{\alpha p_V^2 B(B-1)(1+\frac{Af}{1-f})^2(\hat{\eta}gfC_0(1-A) - p_V r(1+\frac{Af}{1-f})}\right] \end{aligned} \tag{63}$$

The second term is order $\frac{e}{m^2}$, which we drop, giving the following approximation for the mean phage time to extinction (plotted in Figure 36):

$$T_{\text{ext}} \approx \frac{2(1+\ln m)}{f(B-1)}\frac{\tilde{n_V}}{m}\left(1-\frac{1}{Bp_V}\right) \tag{64}$$

Here again we are using the phage population size independent of CRISPR ($\tilde{n_V}$). Notably this is independent of $\eta$ and $e$ (except implicitly in $m$) but is still extremely close to the full theoretical quantity.

From equation 64 we learn:

- $T_{\text{ext}}$ is insensitive to the changes in total phage population size caused by changing CRISPR immunity.
- $T_{\text{ext}}$ increases if the phage clone size $\frac{\tilde{n_V}}{m}$ increases.
- $Bp_V$ is the effective burst size: for every phage adsorption, $Bp_V$ new phages emerge on average. This can be thought of as the phage growth rate. $1-\frac{1}{Bp_V}$ lies between 0 and 1; if $Bp_V$ is larger, $T_{\text{ext}}$ is also larger.

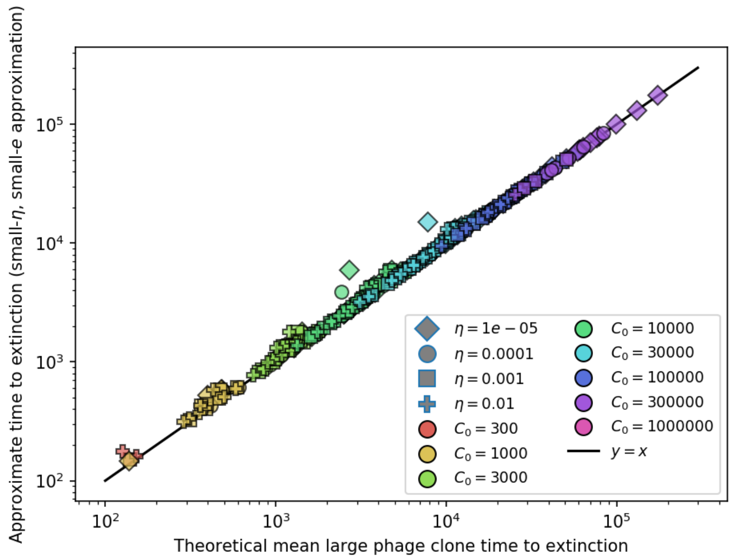

Figure 36: Approximate phage time to extinction vs. numerically calculated theoretical time to extinction, where the approximate time to extinction is given by 64. The full theoretical predicted value of $m$ is used in the approximate expression.

- The burst size $B$ also appears alone in the denominator $(B-1)$. A larger burst size means larger fluctuations: for larger $B$, more deaths tend to happen between birth events at steady-state, so extinction happens sooner on average (shorter $T_{\text{ext}}$ from $B-1$ in the denominator).
- The normalized chemostat flow rate $f$ lies between 0 and 1 and is a death rate for phages; if $f$ is larger $T_{\text{ext}}$ is smaller.

The $1+\ln m$ term is more difficult to parse. A hand-wavy intuition is that while larger $m$ sometimes means smaller clone size and shorter extinction times $(\frac{\tilde{n_V}}{m})$, it is also related to increased phage success in the population, and so the $1+\ln m$ in the numerator softens the straight inverse dependence on $m$ through clone size.

##### 4.2.3 Approximation for $P_{\text{est}}$

Now we approximate the probability of phage clone establishment.

$$P_{\text{est}} = \frac{2s_0}{B(s_0+\delta_0)} \tag{65}$$

Recall that $s_0 = \beta_0(B-1) - \delta_0 = \alpha n_B(Bp_V - 1) - F$, $\beta_0 = n_B \alpha p_V$, and $\delta_0 = F + \alpha n_B(1-p_V)$. These all depend on $n_B$, so plugging in equation 39 for $n_B$, we get

$$P_{\text{est}} = \frac{2e\nu}{m(B-1)} \tag{66}$$

Equation 66 with $\nu$ given by equation 51 is plotted in blue in Figure 37. The same expression with $e=0$ is plotted in green. There is a discontinuity in $\tilde{\nu}$ when the third term in the denominator becomes large, which is why the blue lines don't extend to large effective $e$. These two approximations bound the full solution at low effective $e$.

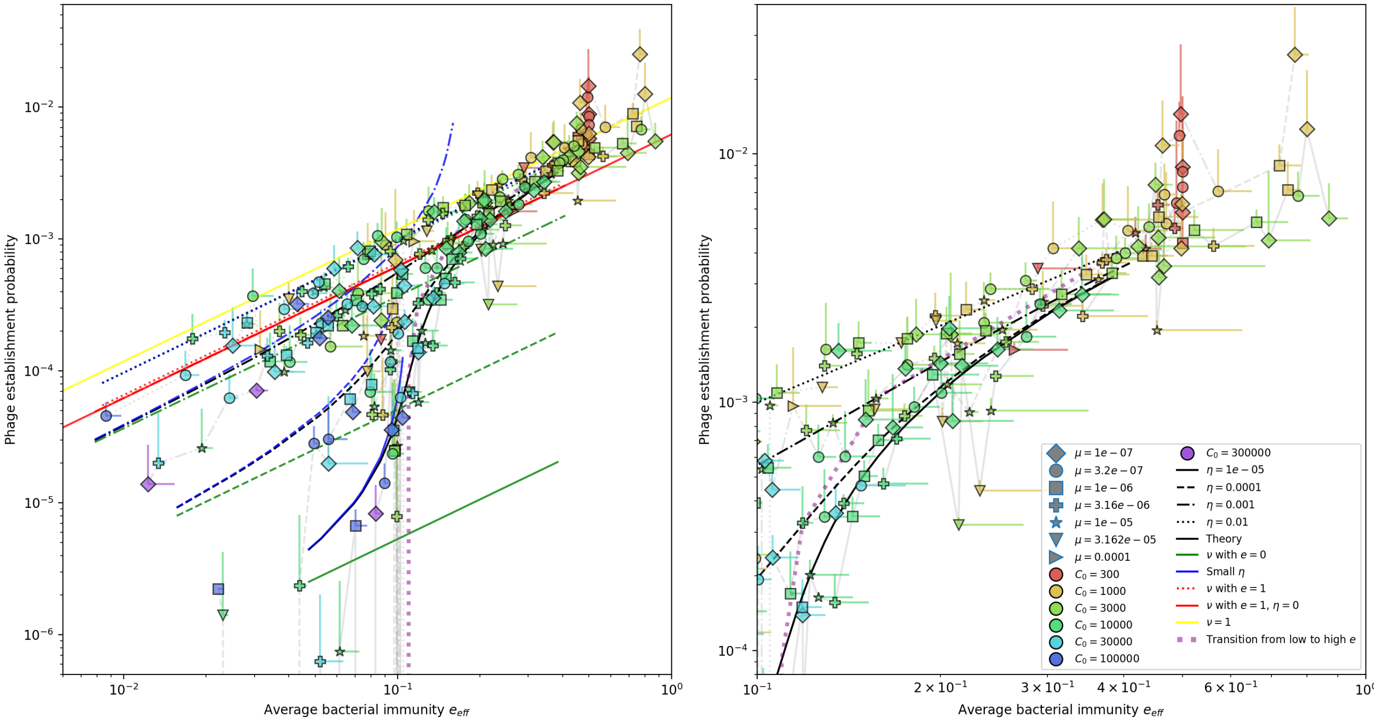

Figure 37: Phage establishment probability vs. effective $e$. Markers are simulation results; error bars are standard deviation across three or more independent simulations. Black lines are the numerical theoretical solution. Blue lines are the approximate solution given by equation 51 (with the full theoretical predicted value of $m$), and green lines are the same but with $e = 0$. All values of $e$ collapse on the same $\eta$ lines; the establishment probability depends only on $e/m$ and not on $e$ by itself. Interestingly, the $e = 0$ approximation is close to the small $e$ approximation at high $\eta$. This implies that high $e$ matters more at small $\eta$. For some simulations near the transition from low to high $\eta$, the probability of establishment is zero, which is why the grey connecting lines drop below the x-axis near effective $e \approx 10^{-1}$.

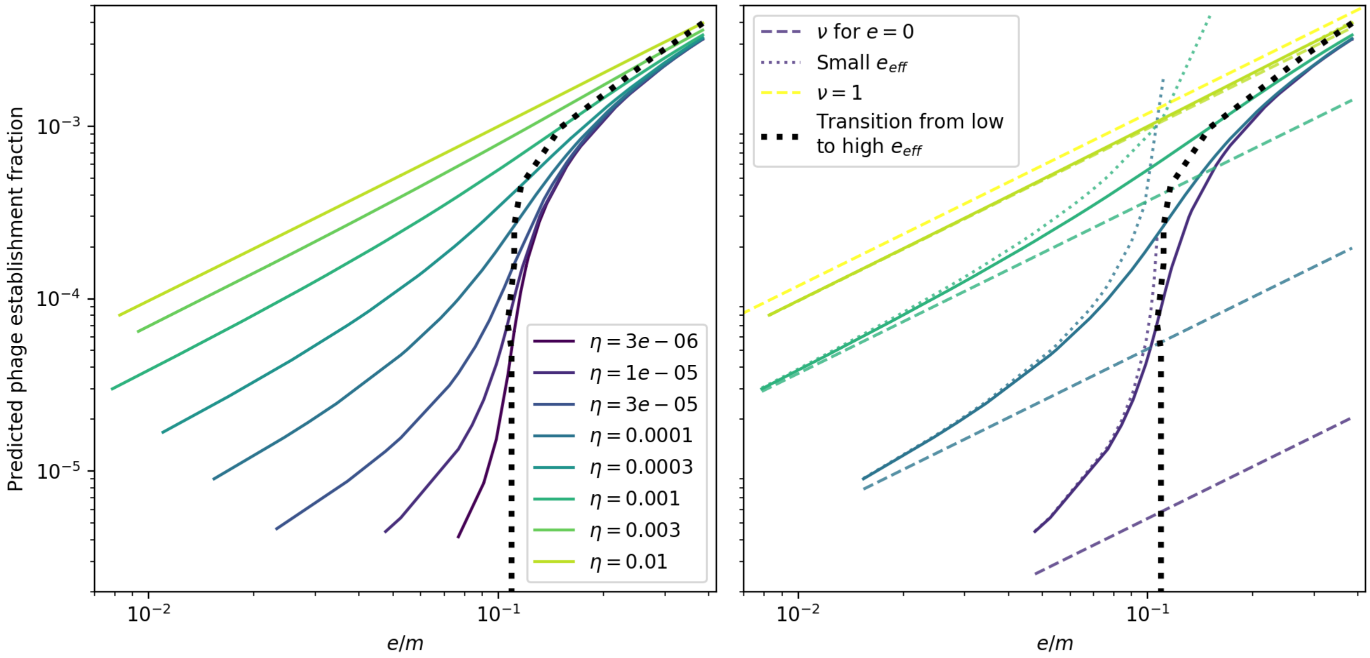

Figure 38: Theoretical phage establishment probability vs. $e/m$. Solid lines are the numerical theoretical solution. Light dashed lines ("small $e_{eff}$") are the approximate solution given by equation 51 (with the full theoretical predicted value of $m$), heavy dashed lines are the same but with $e = 0$. All values of $e$ collapse on the same $\eta$ lines; the establishment probability depends only on $e/m$ and not on $e$ by itself. The inflection point in $P_{est}$ (dashed black line) is given by equation 55.

###### 4.2.4 Approximation for phage mutation rate

The effective phage mutation rate (new phages per minute, $\bar{\mu}$) is given by equation 67. In general, $\bar{\mu}$ depends both explicitly on $e$ and $m$ and implicitly through $n_V$, $n_B$, and $\nu$. We do the same small $e/m$ approximation for $\bar{\mu}$ as above, where we expand $n_V$, $n_B$, and $\nu$ in $e/m$ ($n_V \approx \tilde{n_V} + \frac{e}{m} n_V^1$, etc). The 0th order approximation ($e/m = 0$) does not change $\bar{\mu}$ much at all across the whole range of parameters we investigated (equation 68 and Figure 39).

$$\bar{\mu} = \alpha B(1 - \mathrm{e}^{-\mu L}) p_V n_V n_B (1 - \frac{e\nu}{m}) \frac{1}{g C_0} \tag{67}$$

$$\bar{\mu} = \frac{\alpha B(1 - \mathrm{e}^{-\mu L}) p_V}{g C_0} \left[ \tilde{n_V} \tilde{n_B} + \frac{e}{m} (n_V^1 \tilde{n_B} + \tilde{n_V} n_B^1 - \tilde{\nu} \tilde{n_V} \tilde{n_B}) \right] + \mathcal{O}(e/m)^2 \tag{68}$$

###### 4.2.5 Approximation for $m$

Finally, we do the same approximation for the complete expression for $m$, expanding all variables in powers of $e/m$. The result turns out to be the same as if we combined the individual approximations shown above.

$$m = \underbrace{\frac{2e\nu}{m(B-1)}}_{\text{phage establishment fraction}} \quad \underbrace{\alpha B(1 - \mathrm{e}^{-\mu L}) p_V n_V n_B (1 - \nu \frac{e}{m})}_{\text{phage mutation rate}} \underbrace{\frac{2^{\frac{n_V}{m}} (1 + \ln m)}{B(B-2)\alpha p_V n_B (1 - \nu \frac{e}{m}) + F + \alpha n_B}}_{\text{large phage clone time to extinction}} \tag{69}$$

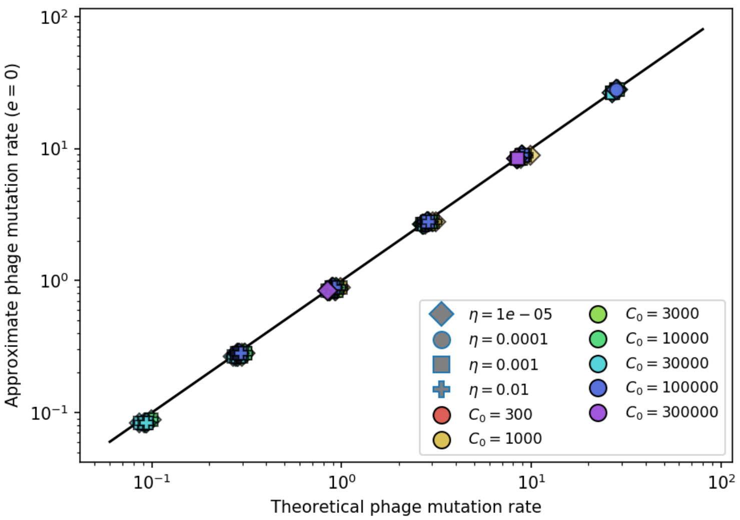

Figure 39: Approximate phage mutation rate for $e = 0$ vs. theoretical phage mutation rate.

$$
\begin{aligned}
m =& \frac{4(1-\mathrm{e}^{-\mu L})\tilde{\nu}\tilde{n_V}^2}{(B-1)^2}\frac{e(1+\ln m)}{m^2} \\
&+\frac{4\tilde{n_V}(1-\mathrm{e}^{-\mu L})\left(BC_0 fgp_V\left(2(B-1)\tilde{\nu}n_V^1+(B-1)\nu^1\tilde{n_V}+\tilde{\nu}^2(-\tilde{n_V})\right)+\alpha n_B^1\tilde{\nu}\tilde{n_V}(Bp_V-1)^2\right)}{(B-1)^3 BC_0 fgp_V}\frac{e^2(1+\ln(m)}{m^3} \\
&+\mathcal{O}(e/m)^3
\end{aligned}
\tag{70}
$$

Let's look at the first term by itself, neglecting $(e/m)^2$ and higher. Rearranging to collect $m$ terms:

$$\frac{m^3}{(1+\ln m)} = \frac{4e(1-\mathrm{e}^{-\mu L})\tilde{\nu}\tilde{n_V}^2}{(B-1)^2} \tag{71}$$

Let's let the right-hand side of equation 71 equal $a$, a new parameter that is independent of $m$. Now we are approximating:

$$m^3 = a(1+\ln m) \tag{72}$$

Changing variables to $\frac{m^3}{a} = z$:

$$z = 1 + \frac{\ln a}{3} + \frac{\ln z}{3} \tag{73}$$

We solve this perturbatively. Let $z_0 = 1 + \frac{\ln a}{3}$, then let $z = z_0(1+\delta)$ where we assume $\delta$ is small. Now we are solving for the perturbation $\delta$:

$$z_0(1+\delta) = z_0 + \frac{1}{3}\ln(z_0(1+\delta)) \tag{74}$$

We approximate $\ln(1+\delta) \approx \delta$ for small $\delta$. This gives

$$\delta \approx \frac{\ln z_0}{3z_0 - 1} \tag{75}$$

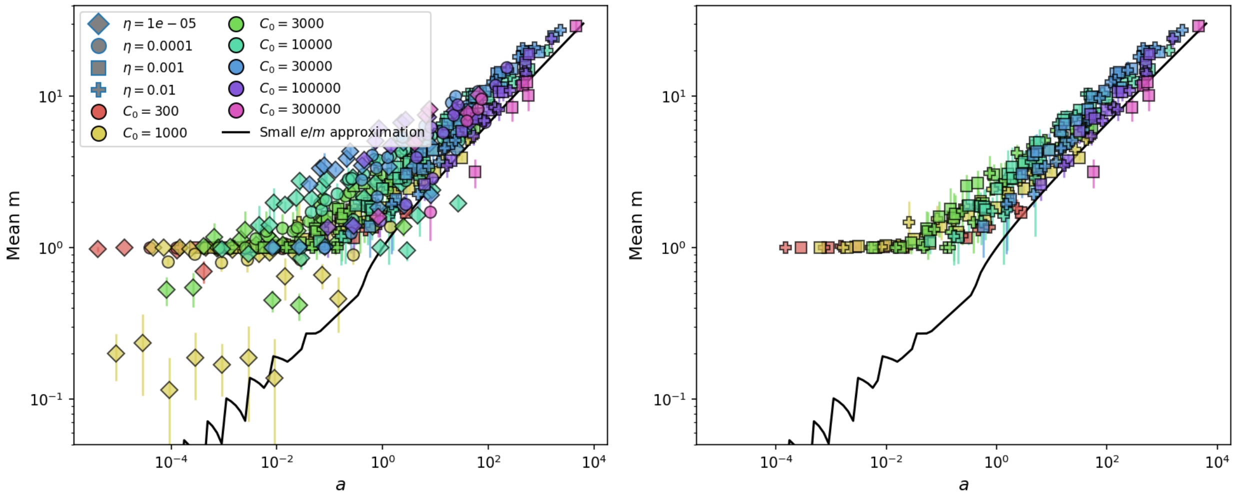

Figure 40: Measured mean $m$ in simulations vs $a$ as given by equation 78. The solid line is $m$ vs. $a$ solved numerically using equation 167. The right panel shows the same but with the two lowest values of $\eta$ removed.

$$m \approx \left( a z_0 (1 + \frac{\ln z_0}{3z_0 - 1}) \right)^{\frac{1}{3}} \tag{76}$$

$$m \approx \left[ \frac{a(\ln(a) + 3) \left( \ln(a) + \ln\left(\frac{1}{3}(\ln(a) + 3)\right) + 2 \right)}{3(\ln(a) + 2)} \right]^{\frac{1}{3}} \tag{77}$$

If $a$ is large, the leading order contribution to $m$ is $a^{1/3}$.

$$a = \frac{4e\tilde{\nu}\tilde{n_V}^2(1 - \mathrm{e}^{-\mu L})}{(B-1)^2} \tag{78}$$

To get more insight into the dependence of $a$ and $m$ on parameters, we make some approximations. First, since $\mu L << 1$, we approximate $(1 - \mathrm{e}^{-\mu L}) \approx \mu L$. Then substituting $\tilde{\nu}$ in $a$:

$$a \approx \frac{4e\mu L \frac{\hat{\eta}\alpha\tilde{n_V}}{\hat{\eta}\alpha\tilde{n_V} + r} \tilde{n_V}^2}{(B-1)^2} \tag{79}$$

Substituting in $\tilde{n_V}$:

$$\tilde{n_V} = \frac{C_0}{\alpha p_V} \frac{(Bp_V - 1)(1-f)\alpha - fg}{1 + (Bp_V - 1)\alpha/g} \tag{80}$$

$$a \approx \frac{4e\mu L \frac{\hat{\eta}\alpha(\frac{C_0}{\alpha p_V} \frac{(Bp_V-1)(1-f)\alpha - fg}{1+(Bp_V-1)\alpha/g})}{\hat{\eta}\alpha(\frac{C_0}{\alpha p_V} \frac{(Bp_V-1)(1-f)\alpha - fg}{1+(Bp_V-1)\alpha/g}) + r} (\frac{C_0}{\alpha p_V} \frac{(Bp_V-1)(1-f)\alpha - fg}{1+(Bp_V-1)\alpha/g})^2}{(B-1)^2} \tag{81}$$

Now we do an expansion assuming large burst size, $B >> 1$. Expanding in $1/B$:

$$a \approx \frac{4g^3 C_0^3 (1-f)^3 e\eta\mu L(1-p_V)}{\alpha^2 B^2 p_V^2 (gC_0(1-f)\eta(1-p_V) + p_V r)} + \mathcal{O}\left(\frac{1}{B}^3\right) \tag{82}$$

Now if $\eta$ is small, specifically if $p_V r > \eta(1-p_V)gC_0(1-f)$, we can expand in $\eta$. This assumption roughly amounts to assuming bacterial survival followed by spacer acquisition is rare compared to phage predation and spacer loss.

$$a \approx \frac{4g^3C_0^3(1-f)^3 e\mu L}{\alpha^2 B^2 p_V^3 r}\left[\eta(1-p_V) - \frac{gC_0(1-f)}{p_V r}\eta^2(1-p_V)^2 + ...\right] \tag{83}$$

The 0th order term captures the trend across a wide range of parameters and gives some insight into parameter dependence (Figure 2C). To summarize: For $\mu L << 1$, $B >> 1$ and $p_V r > \hat{\eta} gC_0(1-f)$:

$$a \approx \frac{4e\mu L\hat{\eta}(gC_0(1-f))^3}{B^2\alpha^2 p_V^3 r} \tag{84}$$

Equation 84 implies that $m$ goes like $(e\mu\eta)^{1/3}$, a non-intuitive dependence.

- $m$ increases as the overall phage mutation rate increases.
- $m$ increases as $\nu$ increases.
- $m$ increases for the same reasons that phage time to extinction increases, except that it depends on the total phage population size (instead of the phage clone size).

#### 4.3 Speed of evolution

The speed of evolution of a population is often taken as the rate at which its genetic structure changes over time. For example, Betts *et al.* measure the rate of evolution in a bacteria-phage coevolution experiment by calculating the Euclidean genetic distance between populations at different time points [25], and Rouzine and Rozhnova calculate the rate of substitution [26]. Speed of evolution is sometimes defined more functionally as the rate of change of a population's fitness over time (see [27, 28, 26]).

In our context, the average population fitness does not change over time at steady state (more like a Red Queen scenario than an arms race scenario) and so we use a measure of genetic distance to calculate the speed of evolution. Instead of Euclidean distance (L2 norm), we use Hamming distance or edit distance (L1 norm). This distance metric is more representative of the process involved in changing sequences, since the sequence [1,1] is mutationally twice as far from [0,0] as [0,1]. Our sequence space has as many dimensions as the number of genetic sites, and a population's "centre of mass" can be calculated by weighting each sequence ("position") in this space by the frequency of each subpopulation with that sequence. Our effective space is dimension $L = 30$, since each protospacer has 30 sites that can mutate. Each site can have the value 0 or 1 meaning there are $2^{30}$ possible sequences, but a sequence can differ by at most 30 changes from another sequence. The maximum distance possible in our model using the L1 norm is 30.

For example: if $L = 2$ and there are 20 phages with the sequence [0,1] and 10 phages with the sequence [1,1], then the centre of mass in the 2-dimensional space is $[\frac{1}{3}, 1]$:

$$R_x = \frac{1}{30}(20 \times 0 + 10 \times 1) = 1/3x$$

$$R_y = \frac{1}{30}(20 \times 1 + 10 \times 1) = 1y$$

If the ancestor phage is [0,0], then the centre of mass distance is $1 + 1/3 = \frac{4}{3}$. If the ancestor sequence is at one of the corners (i.e. only values of 0 or 1), calculating the centre of mass in this manner is the same as calculating the population-weighted average distance from the ancestor: 20 phages are 1 mutation away, 10 are 2 mutations away, therefore the average distance is $(20 + 20)/30 = \frac{4}{3}$. However,

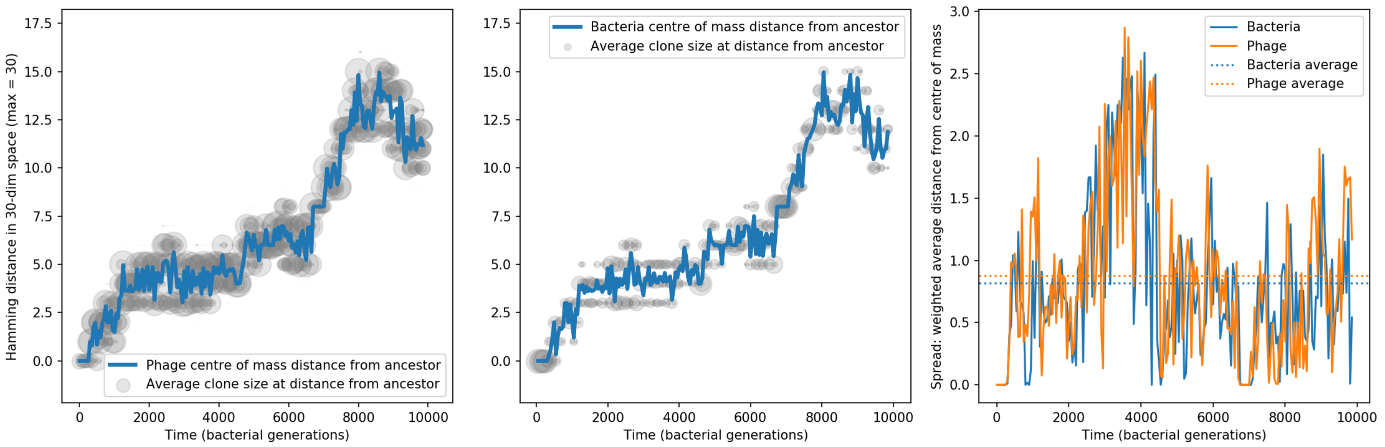

Figure 41: Phage and bacteria centre of mass distance from the original phage and bacteria sequences. The centre of mass distance is plotted in blue for phage (left) and bacteria (centre). Grey circles represent the size of clonal subpopulations at each distance from the ancestor sequence (arbitrary scale). The third panel shows the weighted average distance of the population from the centre of mass at that timepoint, a measure of the spread in sequences present at any time. In this simulation $C_0 = 10^4$, $\eta = 10^{-5}$, $\mu = 10^{-6}$, $e = 0.95$, and mean $m = 2.8$.

if the ancestor sequence is not at a corner (i.e. if comparing two centres of mass), the sum of distances gives larger values than the difference of centres of mass. The sum of distances is a measure of the spread, while the change in centre of mass is a measure of the speed of evolution.

We can look at the distance of the phage and bacteria population in simulations from the ancestor sequence over time to get an idea of how the population moves through sequence space. Figures 41 and 42 show the mutational distance for phage and bacteria over time from either the ancestor phage sequence or a centre-of-mass starting point later in the simulation, for low (Figure 41) and high (Figure 42) values of $\eta$.

Since the maximum distance in our model is 30, there comes a point in distance where mutations back in the direction of the ancestor become likely. The fraction of available mutations $f_m$ that move away from the ancestor decreases as the distance increases: $f_m = \frac{L - \text{distance}}{L}$. For $L = 30$, this means that at a distance of 15 the population is equally likely to move towards or away from the ancestor. This is apparent in Figure 41 - the distance from the ancestor phage reaches 15 and then begins to decrease. In other simulations, the distance continues to increase and doesn't come close to 15 (Figure 42).

##### 4.3.1 Measuring speed of evolution

The speed of evolution in this framework is the distance the population travels in spacer space in a certain amount of time. In principle this is straightforward to calculate, but we find that the distance between populations does not increase linearly with the time interval used to calculate the distance (Figure 43 left panel). This means that measured speed will depend on the time interval used to calculate it (Figure 43 right panel), and there is no clear way to choose one particular time interval over another. We go to the long $\Delta t$ limit and measure the maximum distance from the ancestor reached by a population in a set number of bacterial generations. The maximum distance reached by bacteria and phage populations is highly correlated with each other within a simulation and is also repeatable across independent simulations (Figure 44).

The total number of phage establishments in a simulation determines the maximum distance that the population can move away from the ancestor, but if diversity is high, many of those establishments won't actually move the population further away. A simple approximate scaling for this process is that

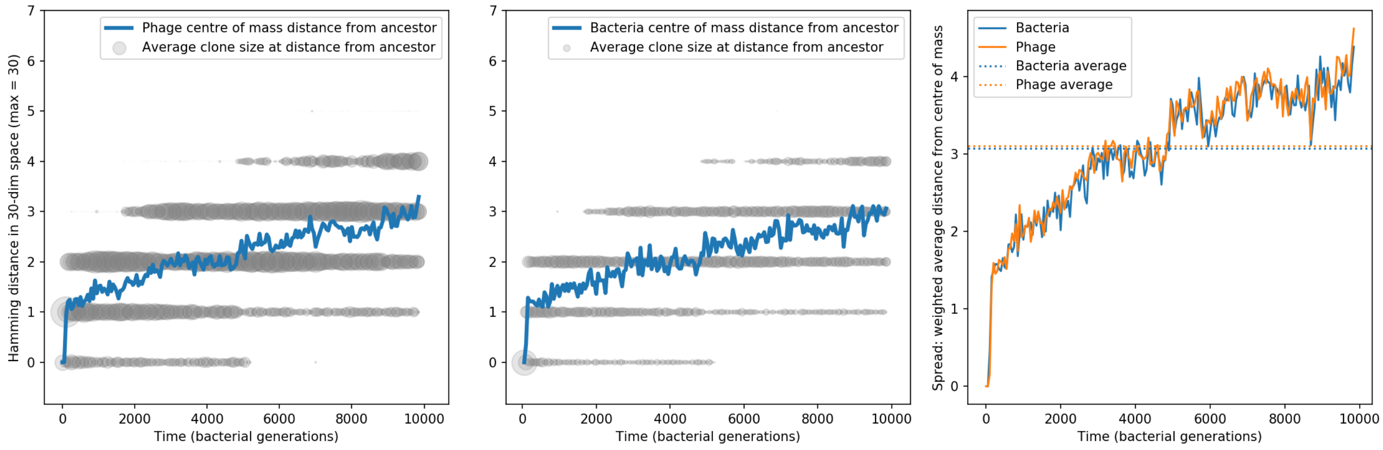

Figure 42: Phage and bacteria centre of mass distance from the original phage and bacteria sequences. The centre of mass distance is plotted in blue for phage (left) and bacteria (centre). Grey circles represent the size of clonal subpopulations at each distance from the ancestor sequence (arbitrary scale). The third panel shows the weighted average distance of the population from the centre of mass at that timepoint, a measure of the spread in sequences present at any time. In this simulation $C_0 = 10^4$, $\eta = 10^{-2}$, $\mu = 10^{-6}$, $e = 0.95$, and mean $m = 14.5$.

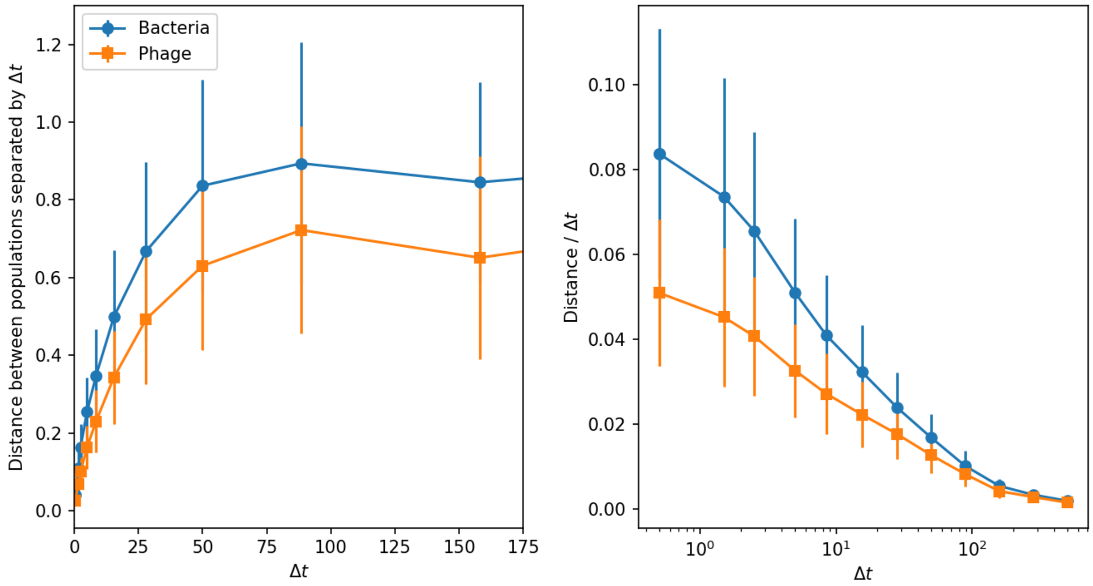

Figure 43: Phage and bacteria centre-of-mass distance from the centre-of-mass at time $t - \Delta t$ (left) and the distance divided by the time interval $\Delta t$ (right). Distances are averaged over the entire simulation at steady-state. Simulation parameters are $C_0 = 10^4$, $\eta = 10^{-2}$, $\mu = 10^{-6}$, $e = 0.95$, and mean $m = 14.5$.

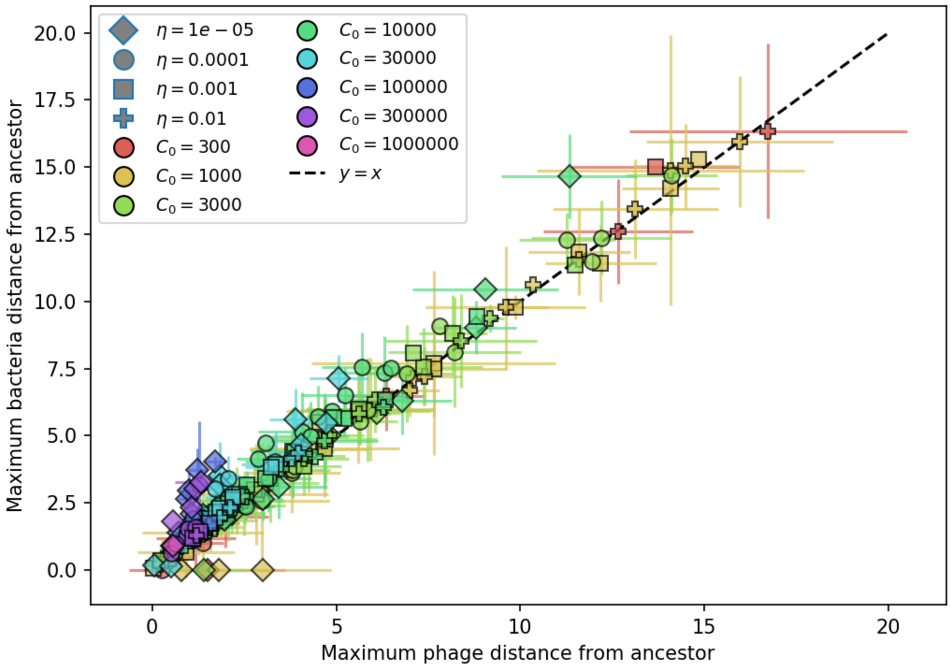

Figure 44: Maximum distance from ancestor population for bacteria vs. phage. The maximum distance is highly correlated, indicating that the bacteria population tracks the phage population closely. Error bars are the standard deviation across multiple independent simulations.

total distance is proportional to $\frac{1}{m}$: if there are $m$ established clones and one of them is the furthest away from the ancestor, there is a $\frac{1}{m}$ probability that the next establishment will come from the furthest-away clone. This is similar to Childs *et al.* [17] where if they have $v$ protospacers per phage (and one dominant bacterial species with a spacer), there is only a $1/v$ chance that a random mutation will actually be an escape mutation.

Since $m$ is a product of mutation rate, establishment probability, and extinction time, plotting distance per establishment vs. $m$ is equivalent to plotting speed vs. time to extinction (equation 85). We have $m = P_{\text{est}}\bar{\mu}T$, where $P_{\text{est}}$ is the probability of establishment for a new phage clone, $\bar{\mu}$ is the effective phage mutation rate, and $T$ is the mean time to extinction for large phage clones. Let the mutational distance from the ancestor be $\Delta = v\tau$, where $v$ is the "speed" and $\tau$ is a fixed time interval used to measure distance.

$$\begin{aligned} \frac{\Delta}{P_{\text{est}}\bar{\mu}\tau} &\propto \frac{1}{m} \\ \frac{v\tau}{P_{\text{est}}\bar{\mu}\tau} &\propto \frac{1}{P_{\text{est}}\bar{\mu}T} \\ v &\propto \frac{1}{T} \end{aligned} \tag{85}$$

Figure 45 shows distance per establishment vs. $m$ and speed vs $T$. Relating distance and speed to the time to extinction gives further insight into the underlying parameter dependencies: we know from other approximations (section 4.2) that $T \propto \tilde{n_V}\frac{1+\ln m}{m}$, and for small $m$ we can substitute our approximation for $m$, giving $T \propto \frac{1}{e\mu\eta}^{\frac{1}{3}}$ and $v \propto e\mu\eta^{\frac{1}{3}}$. Applying the same approximation here as for $m$ above (equation 84), with $T_{ext} \approx \frac{2\tilde{n_V}}{f(B-1)m}\left(1 - \frac{1}{Bp_V}\right)$ (Figure 6):

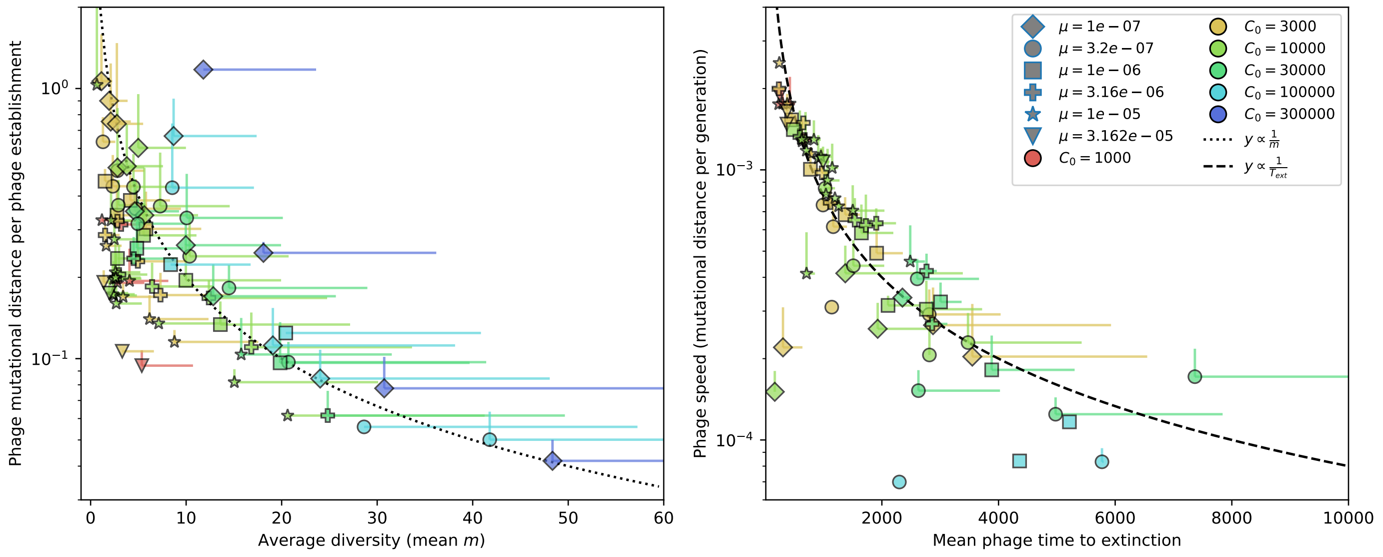

Figure 45: Phage mutational distance reached in simulations divided by the number of phage establishments vs. diversity (left) and phage mutational distance per generation vs. mean phage time to extinction (right). Error bars are the standard deviation across multiple independent simulations and are shown in the positive direction only.

$$v \approx \alpha B f \left( \frac{e\mu\eta L(1-p_V)}{2\alpha^2 B^2 r} \right)^{\frac{1}{3}} \tag{86}$$

##### 4.3.2 Spread in sequence space

We can ask how spread out the population is in sequence space - how far is the average clone from the centre-of-mass? When there are more clones (larger $m$), the population spread is larger (Figures 48 and 49). The spread also depends on initial conditions, at least on the timescale of our simulations: simulations that start with one phage clone (Figure 48) have lower spread than simulations that start with 50 phage clones (Figure 49). In principle, waiting for a very long time in simulations that begin with 1 clone should produce results in line with simulations that start with 50 clones as different clans diverge from each other over time. This suggests that there are multiple features with which we could define steady-state, and they don't all reach true steady-state at the same time. Total population size equilibrates fastest, followed by diversity (SI Figure 1), and then spread in sequence space.

##### 4.3.3 Number and size of clone clans

Watching a simulation visualization of spacer and protospacer types, it is apparent that over time several different lineages may appear and become separated from each other in genome space (Figure 50). What determines how many separate "clans" there are at steady state? What is the characteristic size of a clan? How does this depend on parameters? To address this question, we perform agglomerative clustering on protospacer and spacer sequences to identify separated groups of sequences. We use the L1 norm to define distance, which as described above is the most natural distance metric for sequence changes by mutation. Clusters are grouped using the 'single' linkage criterion in scikit-learn AgglomerativeClustering: the cluster distance is the minimum of the distances between all observations of the two sets. In other words, a collection of sequences that differ from any other sequence in the cluster by 1 mutation are all grouped together, even if some of the sequences differ by more than 1 from

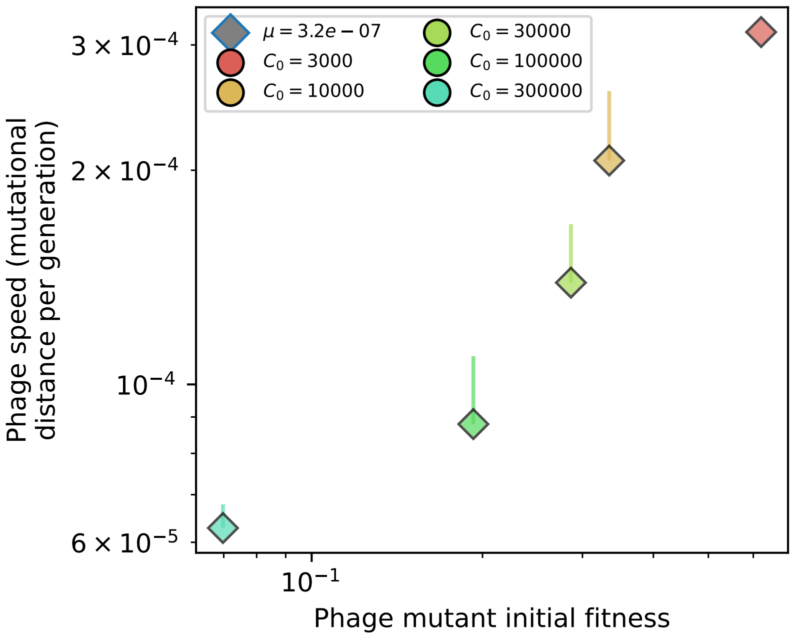

Figure 46: Phage mutational distance per generation vs. initial phage mutant fitness for simulations with $e = 0.95$, $\eta = 10^{-3}$, and $\mu = 3 \times 10^{-7}$. Error bars are the standard deviation across multiple independent simulations and are shown in the positive direction only.

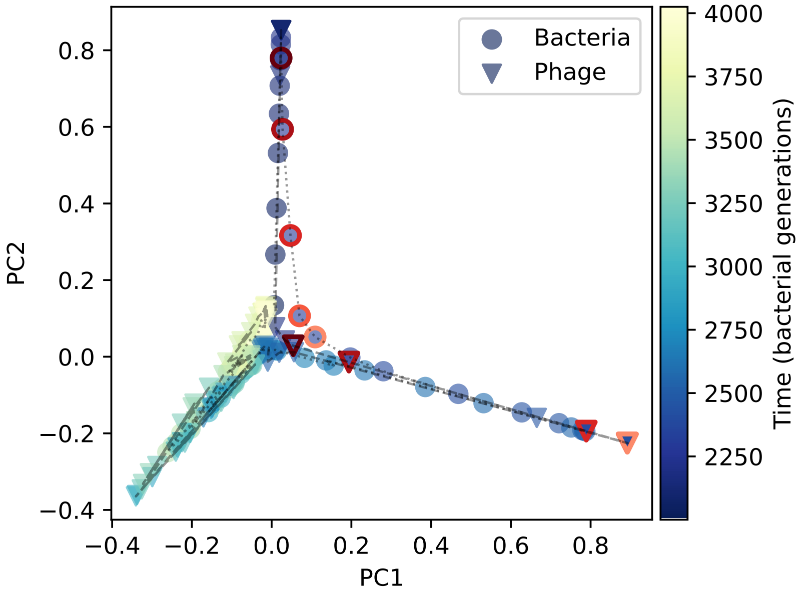

Figure 47: PCA decomposition of phage and bacteria clone abundances for a simulation with exponential cross-reactivity and $C_0 = 10^4$, $e = 0.95$, $\eta = 10^{-4}$, and $\mu = 10^{-6}$. Clone abundances are normalized at each time point, then PCA is performed for the entire phage time series over $\approx 500$ generations (4 times the mean extinction time for phage clones). Bacteria and phage clone abundances are transformed into the PCA coordinates; colours indicate simulation time. Five time points are highlighted in progressively lighter shades of red for emphasis.

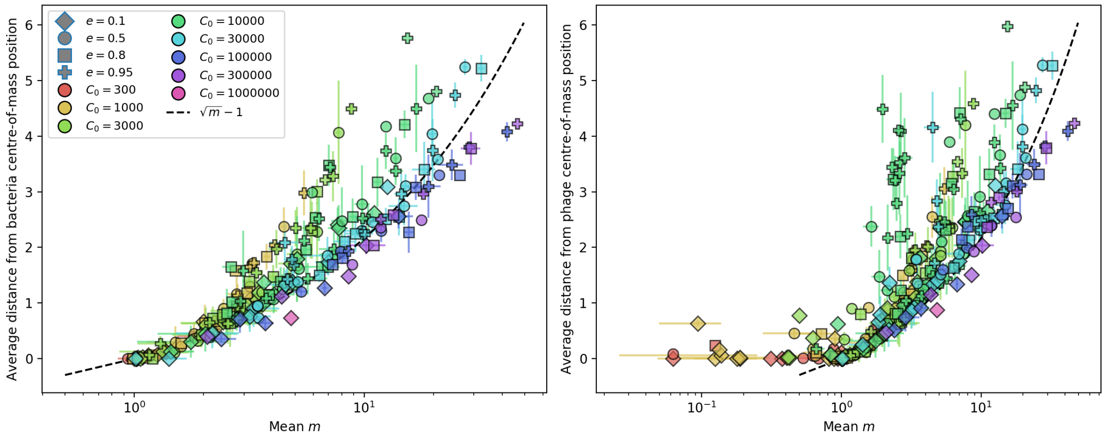

Figure 48: Average population distance from the centre of mass at steady-state for bacteria (left) and phages (right) vs. mean $m$ for simulations with 1 original phage clone ancestor. The dashed line is $\sqrt{m} - 1$, a purely phenomenological choice.

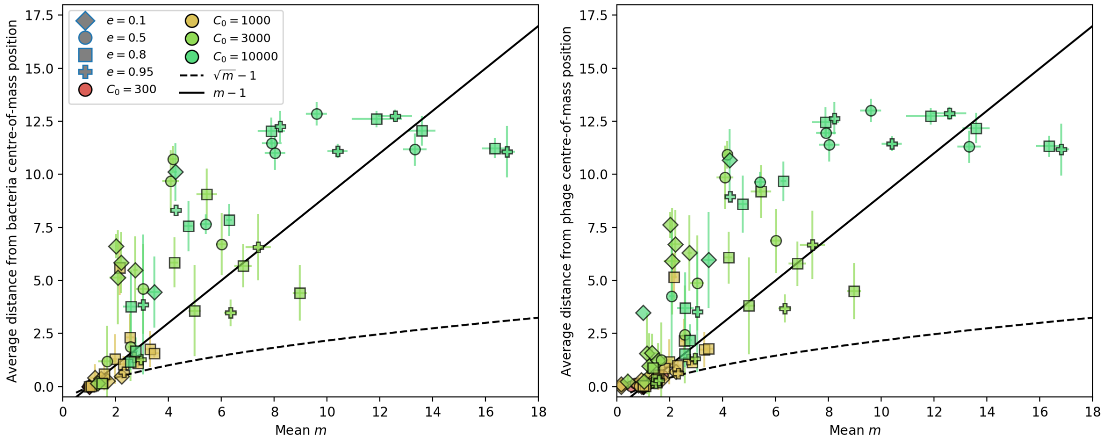

Figure 49: Average population distance from the centre of mass for bacteria (left) and phages (right) vs. mean $m$ for simulations with 50 original phage clones. The dashed line is $\sqrt{m} - 1$ and the solid line is $m - 1$.

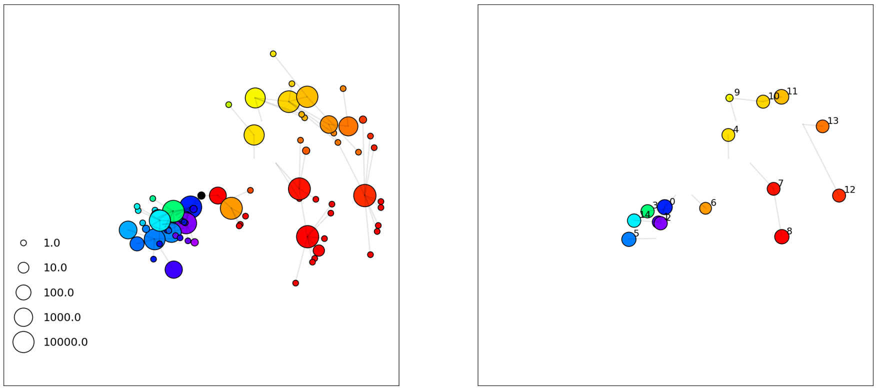

Figure 50: A frame from a simulation movie at 5000 generations with $C_0 = 10^4$, $\eta = 10^{-3}$, $\mu = 10^{-5}$, $e = 0.95$, initial $m = 1$. Phages are on the left, bacteria with spacers on the right.

each other. This is a reasonable criterion for identifying groups of sequences that are related by descent and are actively feeding each other with mutations.

Figures 51 and 52 show the resulting dendrogram from this clustering process for phages and bacteria at one time point for one initial phage clone and 10 initial phage clones. To determine the number of clusters, we use a distance threshold of 2. This separates groups that have no members closer than 2 mutations away. In reality these groups may still be linked by descent more recently than other groups (apparent in the dendrograms), but a distance of 2 or more means these groups are likely to remain separate going forward in time since they lack a connecting clone that can be accessed by a single mutation.

Simulations with an initial number of clones greater than 1 converge to a steady-state number of clusters at a different rate than beginning with 1 clone, since clones from a single ancestor remain more related on average than clones from multiple different ancestors (Figures 53 and 54).

We calculate the average number of clans at steady state and the average clan size for bacteria and phages across all simulations. The number of clans is proportional to $m$; at high mutation rates the number of clans is approximately $m/2$. Low mutation rates mean that it takes a long time for the number of clans to equilibrate, which can be seen by contrasting the mutation rate dependence of 10 initial clones (Figure 55) with 1 initial clone (Figure 56).

The regimes of low and high clan number in our model are similar to regimes of virus-host coevolution identified in ref. [29]: at low virus mutation rates and low mutational jump distances, trajectories move in a straight line in antigenic space, while at higher jump distances, trajectories become more diffusive, the way ours look. At even higher mutation rates and jump sizes, trajectories split into multiple stable coexisting lineages, which is what we see at high diversity.

Based on their parameters and regimes, we expect that increasing cross-reactivity relative to mutation rate in our model should bring our results closer to the ballistic regime.

#### 4.4 Cross-reactivity

In our standard simulations and theory, we assume that $p_V(i, j)$ is binary (equation 87): any mismatch between protospacers and spacers means bacteria no longer have any immune advantage against phage. Here, we discuss results of simulations where $p_V$ includes some cross-reactivity so that the more mutations a protospacer contains, the less immunity bacteria have against it.

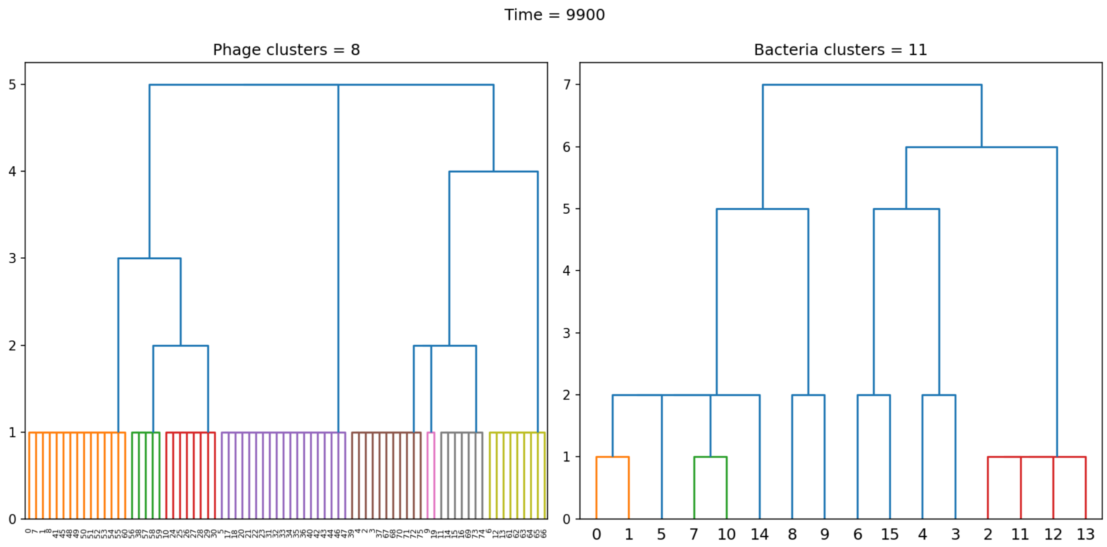

Figure 51: Dendrogram resulting from agglomerative clustering with the L1 norm and linking clusters using the minimum distance between members. The number of clusters is determined with a cutoff at a distance of 2. $C_0 = 10^4$, $\eta = 10^{-3}$, $\mu = 10^{-5}$, $e = 0.95$, initial $m = 1$.

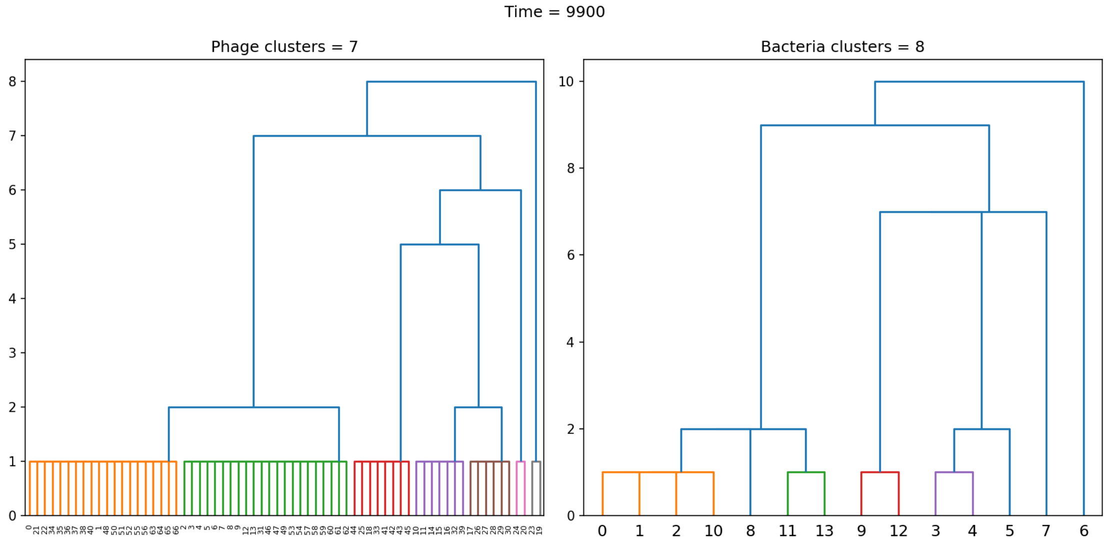

Figure 52: Dendrogram resulting from agglomerative clustering with the L1 norm and linking clusters using the minimum distance between members. The number of clusters is determined with a cutoff at a distance of 2. $C_0 = 10^4$, $\eta = 10^{-3}$, $\mu = 10^{-5}$, $e = 0.95$, initial $m = 10$.

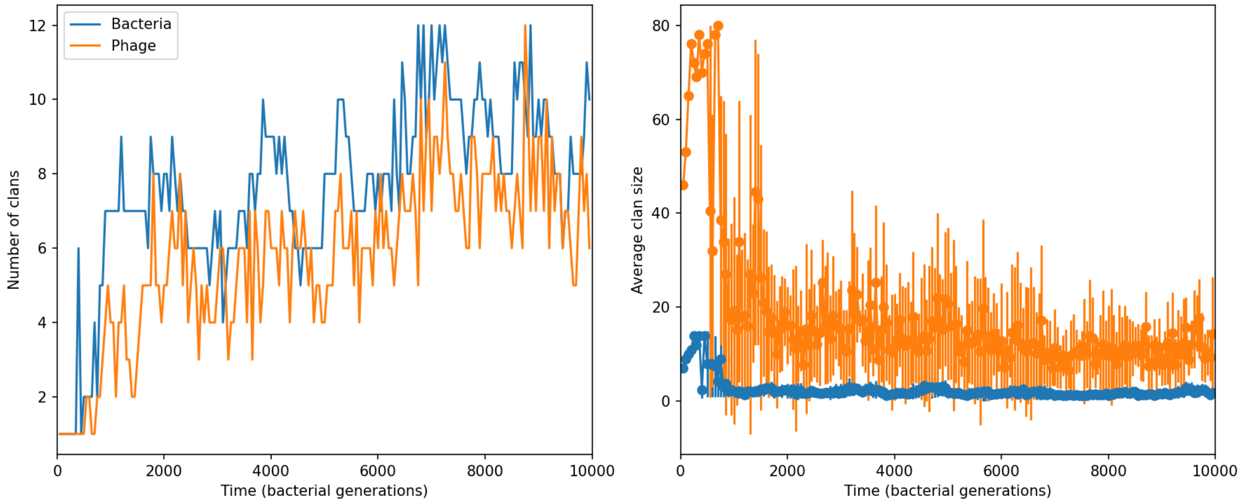

Figure 53: Clan number and size over time in a simulation with $C_0 = 10^4$, $\eta = 10^{-3}$, $\mu = 10^{-5}$, $e = 0.95$, initial $m = 1$.

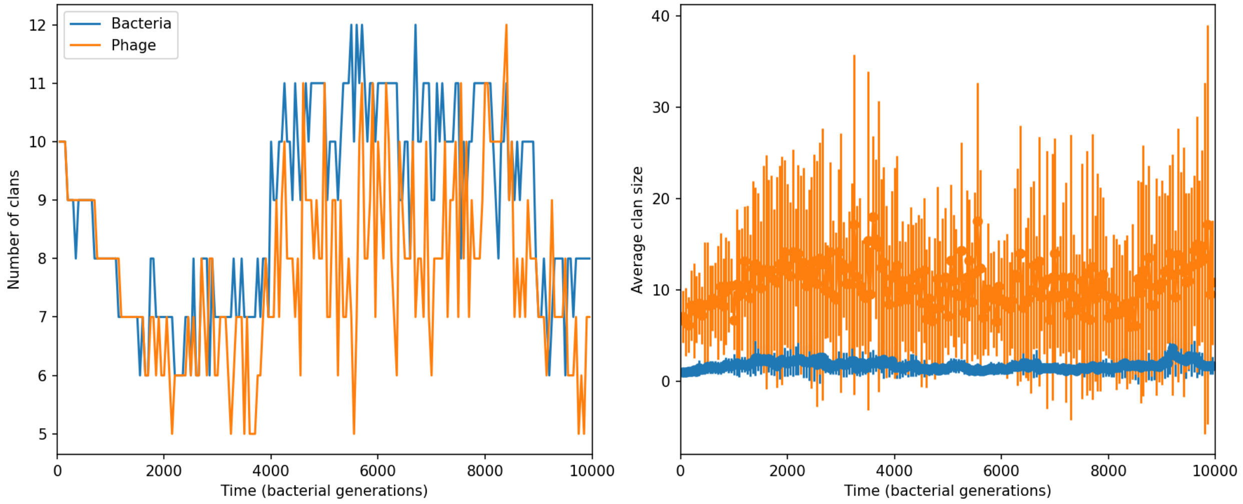

Figure 54: Clan number and size over time in a simulation with $C_0 = 10^4$, $\eta = 10^{-3}$, $\mu = 10^{-5}$, $e = 0.95$, initial $m = 10$.

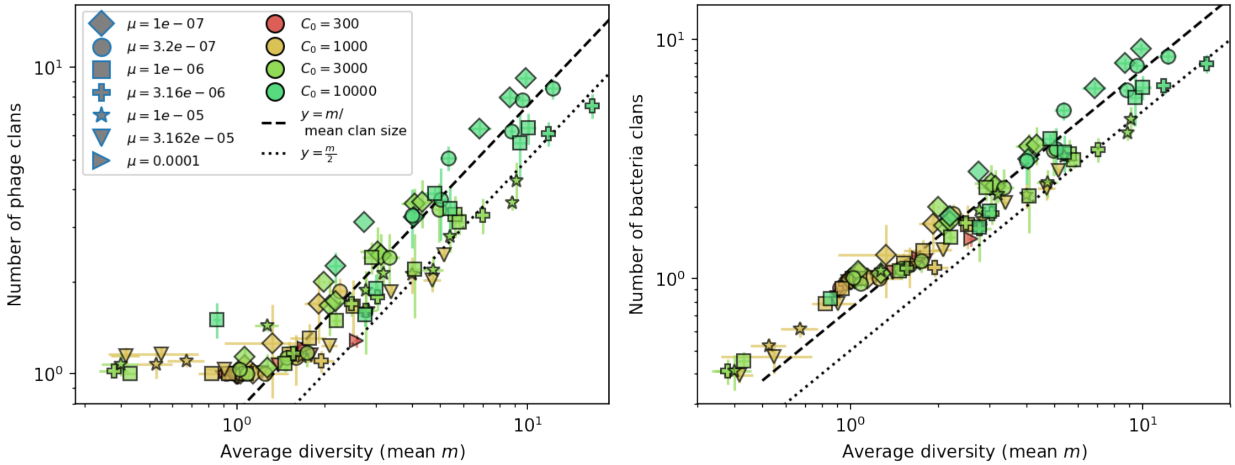

Figure 55: Average clan number vs. $m$ for all simulations that begin with 10 clones. The dashed lines are $m$ divided by the mean bacterial clan size ($\approx 1.3$) and $m/2$.

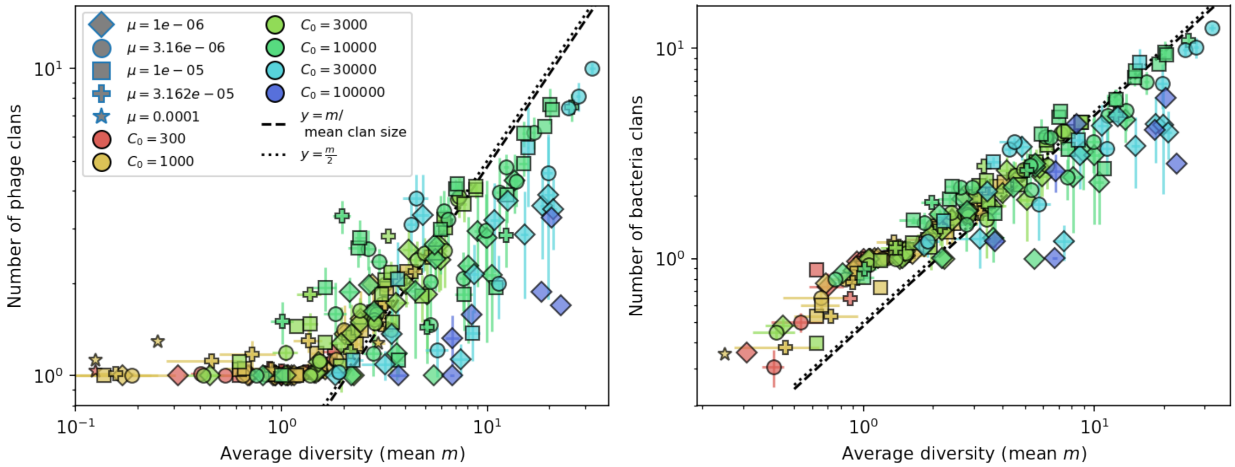

Figure 56: Average clan number vs. $m$ for all simulations that begin with 1 clones with $\mu \geq 10^{-6}$. The dashed lines are $m$ divided by the mean bacterial clan size and $m/2$.

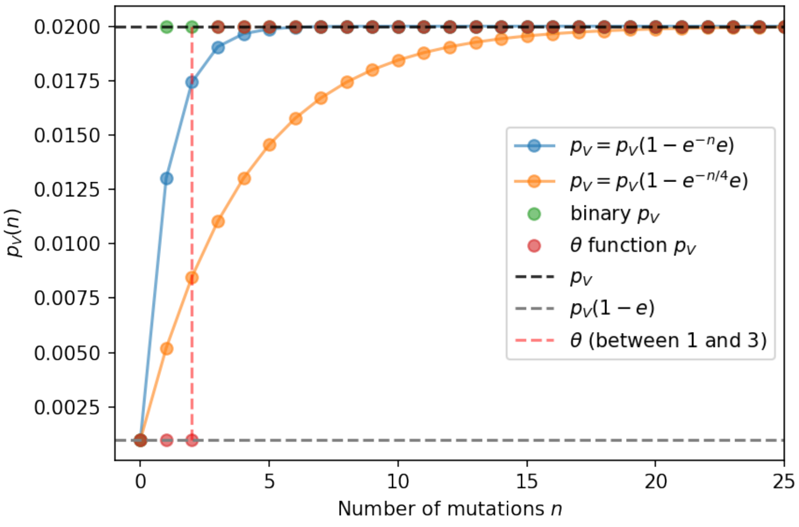

Figure 57: Phage infection success probability $p_V$ as a function of mutational distance between spacers and protospacers for different definitions and degrees of cross-reactivity. Definitions are plotted for $p_V = 0.02$, $e = 0.95$.

We explore two types of cross-reactivity: one in which we define $p_V(i,j)$ as an exponential function of the mutational distance between a spacer and protospacer (equation 88), and one in which $p_V(i,j)$ is a $\theta$-function of mutational distance (equation 89). The exponential definition of cross-reactivity is the same as that used in refs. [29, 30] to model viruses evolving in abstract antigen space [29] and in genomic distance space [30].

$$p_V(i,j) = \begin{cases} p_V(1-e) & \text{if } i = j \\ p_V & \text{if } i \neq j \end{cases} \tag{87}$$

$$p_V(i,j) = p_V\left(1 - e\,\exp\left[-\frac{n_{ij}}{d}\right]\right) \tag{88}$$

$$p_V(i,j) = \begin{cases} p_V(1-e) & \text{if } n_{ij} \leq \theta \\ p_V & \text{if } n_{ij} > \theta \end{cases} \tag{89}$$

The mutational distance $n_{ij} = \sum |(V_i - V_j)|$ is the number of mutations between a protospacer and spacer, and $d$ scales the radius of cross-reactivity, with larger $d$ meaning more mutations are required to achieve the same immune escape [30]. Note that our usual binary definition of $p_V$ is a special case of 89 with $\theta = 0$. Figure 57 shows these different definitions of $p_V(i,j)$ as a function of $n_{ij}$.

Cross-reactivity leads to a higher average bacterial immunity against phages for the same parameters (Figures 58 and 60). This means that our assumption that effective $e \approx e/m$, valid for nearly all parameters without cross-reactivity, breaks down when we introduce cross-reactivity. Importantly, this is a quasi-steady-state effect: if we start simulations with more initial clones, clone clans are genetically separated and both mean $m$ and average immunity converge to the binary $p_V$ case (Figure 58 right panel).

Phages are less likely to establish when there is cross-reactivity because phage escape mutations are incomplete (main text Figure 3E). However, the clones that do survive grow quicker: simulations with high cross-reactivity appear to have a faster mean growth rate for clones, regardless of conditioning on survival (Figure 59).

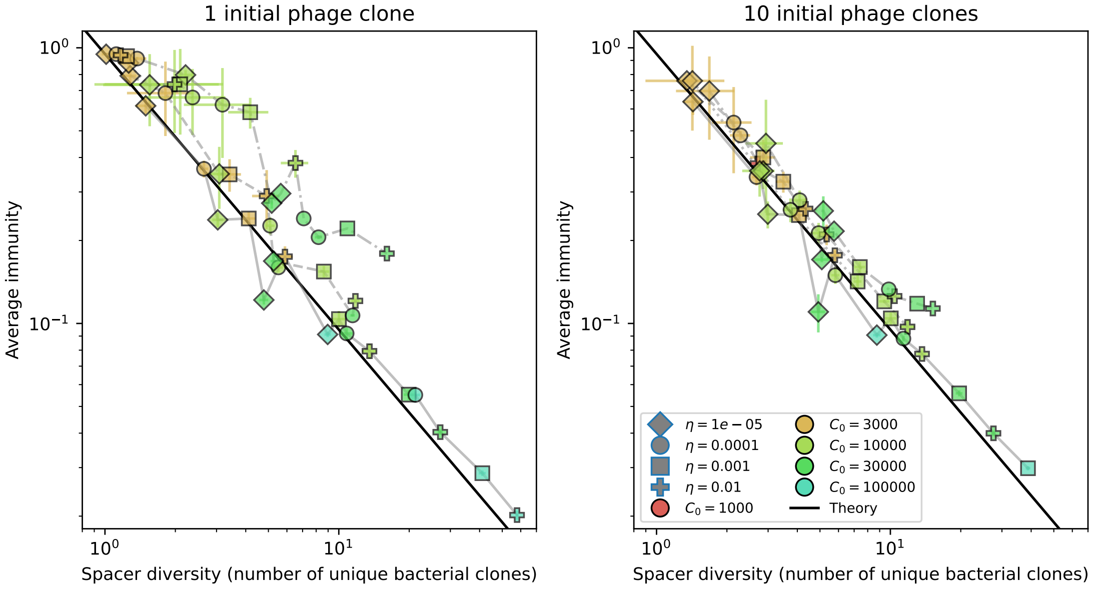

Figure 58: Average immunity vs. diversity with differnet degrees of cross-reactivity for simulations with $e = 0.95$, $\mu = 10^{-6}$. Dashed lines are simulations with cross-reactivity, solid line is simulations without cross-reactivity.

Adding cross-reactivity causes some unexpected dynamical behaviours. We observed a traveling wave regime in some simulations with large spikes in phage clone size (Figures 63 and 64). This happens because cross-reactivity creates a fitness gradient for new phage mutants such that some mutants are much more fit and establish much more quickly.

To see this happening, let's look closely at the simulation in Figure 63 at time 3565 (Figure 65). There are 3 large phage clones and 2 large bacteria clones (soon to be 3). Phage clone 0 is the largest and phage clone 2 is the smallest. Phage clone 0 and phage clone 1 are both either 0 or 1 mutation away from bacteria clones 0 and 1 (Figure 66). Because $\theta = 1$, they are all equally fit; effectively they could be considered one large clone. Phage 2, however, is 1 mutation from bacteria clone 1 but 2 mutations from bacteria clone 0, so it now has a much higher fitness than the other two phage clones and grows quickly. But now bacteria clone 1 is also fit against this new mutant, so it too continues growing. This push-pull keeps going with new phage mutants experiencing runaway fitness for a while. This also leads to an asymmetry in clone identity between the populations: because a bacterial clone may be immune to a phage clone with a different but related sequence, we see bacteria clones becoming large *after* their matching phage clone has gone extinct because they are still resistant to an existing phage clone that is genetically related (Figures 65 and 68).

This fitness and growth pattern can be seen as a function of the marginal immunity of bacterial clones to the phage population and of the bacterial population to individual phage clones (Figure 68). Average immunity for individual clones, which we call marginal immunity in analogy with ref. [31], captures the fitness differences between individual clones that determine their dynamical behaviour. Without cross-reactivity, bacteria clones are only immune to at most one phage clone at a time, though their overall marginal immunity still changes smoothly as the matching phage clone changes size (Figure 69B). When cross-reactivity is added, bacteria may be immune or partially immune to multiple phage clones, and calculating marginal immunity summarizes all of these combined effects on the fitness of a particular clone. Bacteria clones grow larger once their marginal immunity is high, and phage clones grow large once bacterial marginal immunity against them is low. These opposing forces generate Lotka-Volterra oscillations even in the case with no cross-reactivity, though these oscillations are more rapid

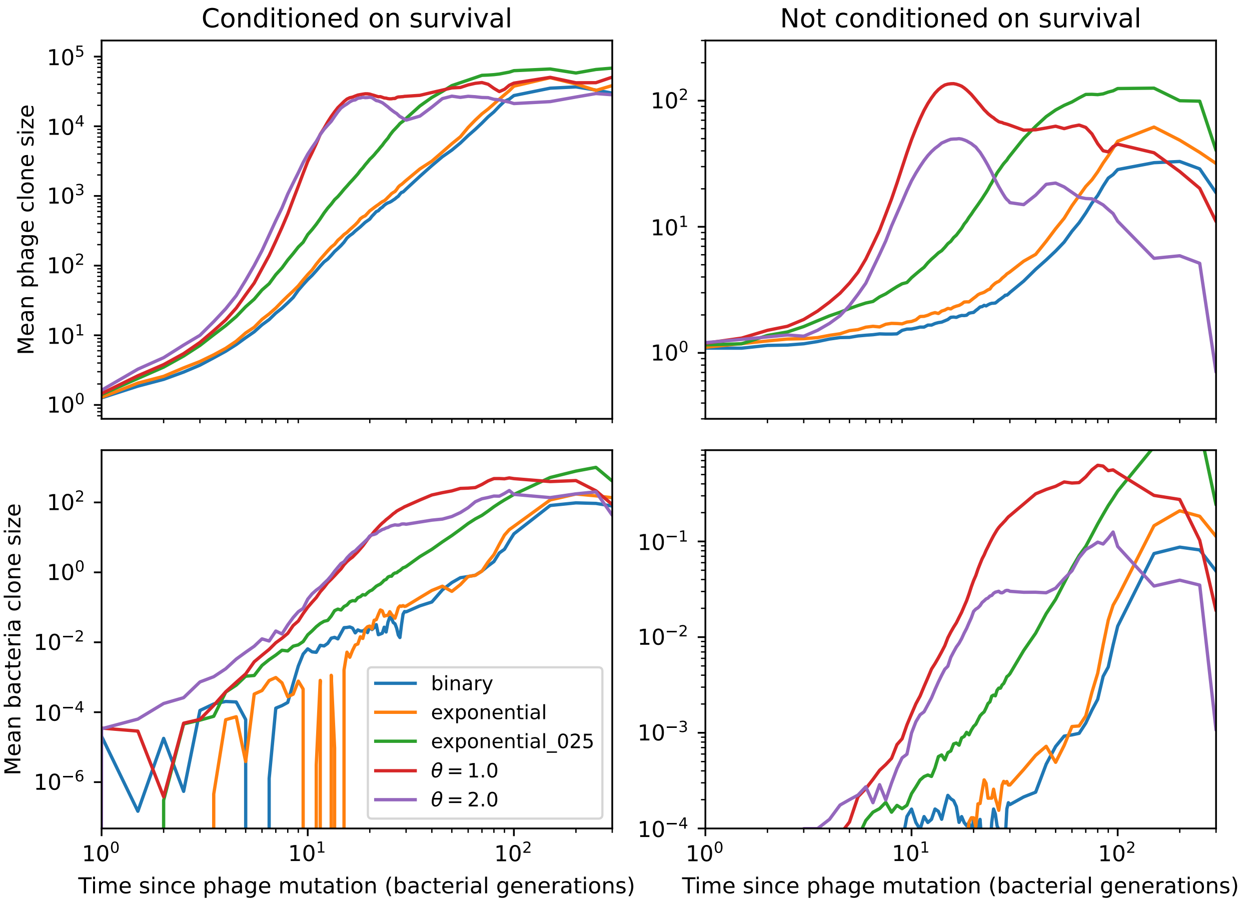

Figure 59: Mean phage clone size (top) and mean bacteria clone size (bottom) relative to time of phage mutation for different definitions of $p_V$. These simulations begin with 1 initial phage clone and parameters $C_0 = 10^4$, $\mu = 10^{-6}$, $\eta = 0.0001$, $e = 0.95$.

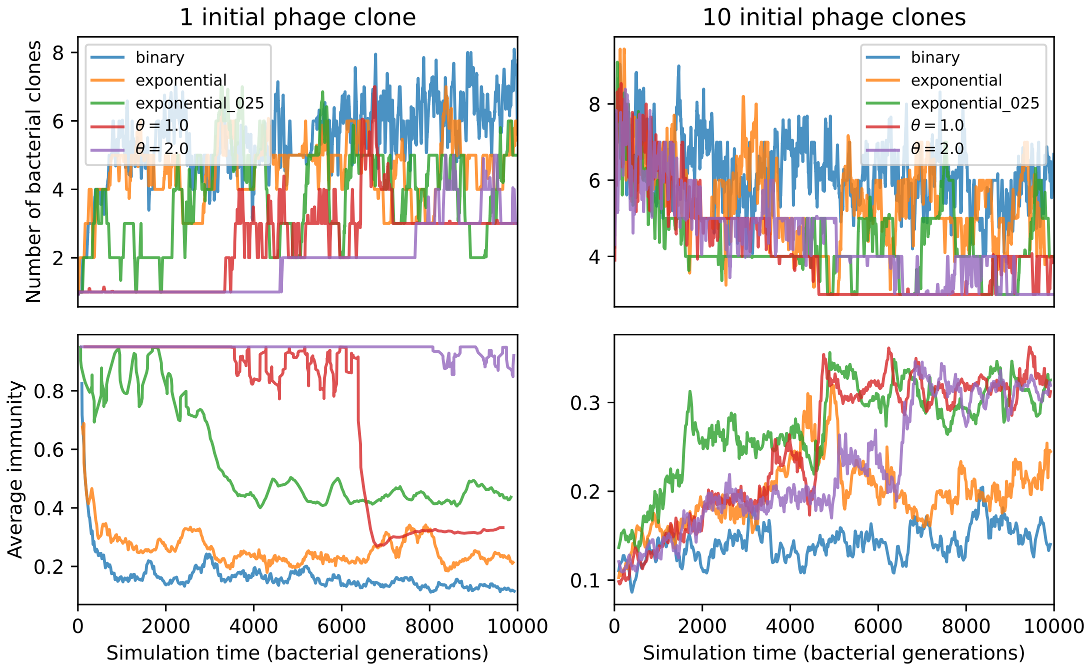

Figure 60: Number of bacterial clones (top) and average immunity (bottom) for simulations beginning with either 1 phage clone (left) or 10 phage clones (right). These simulations have parameters $C_0 = 10^4$, $\mu = 10^{-6}$, $\eta = 0.0001$, $e = 0.95$.

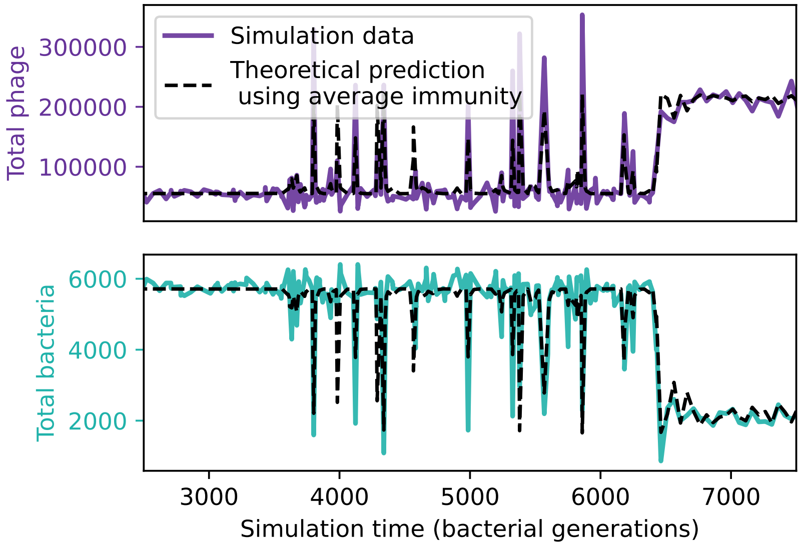

Figure 61: Total phage (top) and total bacteria (bottom) in a simulation with cross-reactivity (step function CRISPR effectiveness with $\theta = 1$). The dashed black line uses the measured value of average immunity from the simulation at each time point to predict population sizes using the solutions to the system of equations 5 through 9. This simulation has 1 initial phage clone and parameters $C_0 = 10^4$, $\mu = 10^{-6}$, $\eta = 10^{-4}$, and $e = 0.95$.

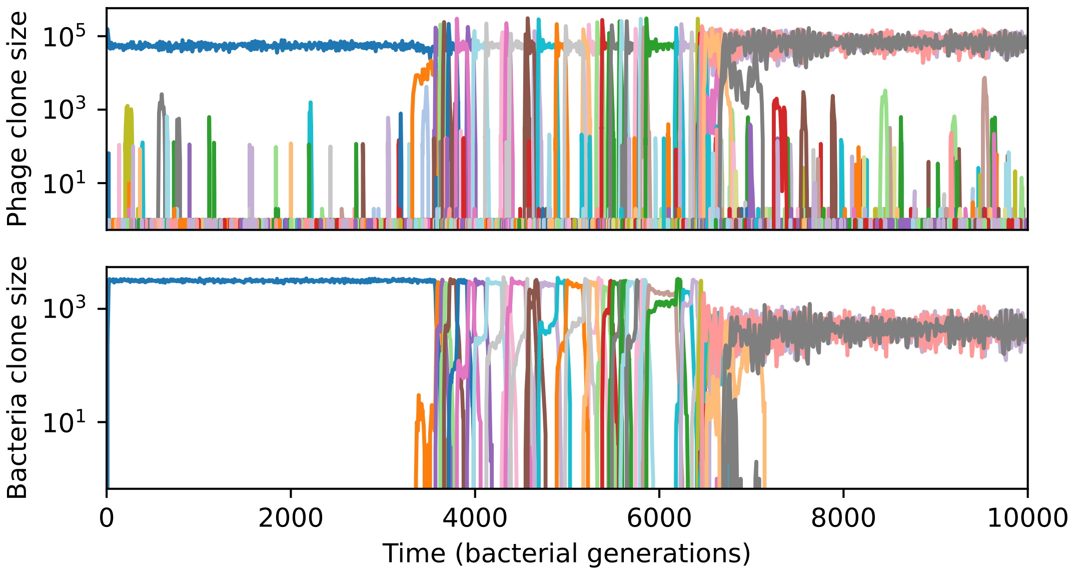

Figure 62: Phage clone size (top) and bacteria clone size (bottom) in a simulation with cross-reactivity (step function CRISPR effectiveness with $\theta = 1$). This simulation has 1 initial phage clone and parameters $C_0 = 10^4$, $\mu = 10^{-6}$, $\eta = 10^{-4}$, and $e = 0.95$. Later times in this simulation are shown in Figure 64), earlier times in this simulation are shown in Figure 63).

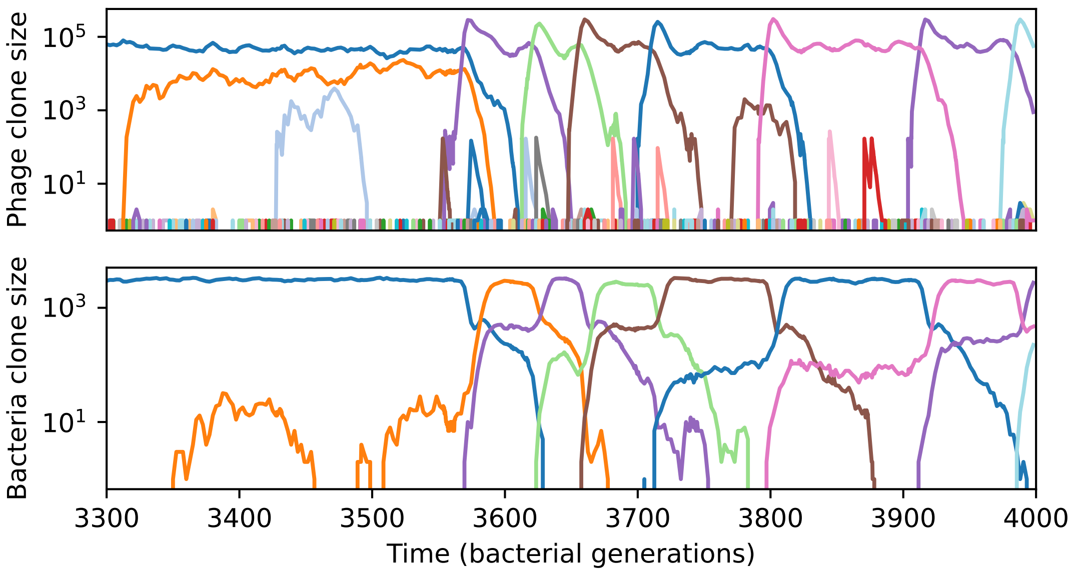

Figure 63: Phage clone size (top) and bacteria clone size (bottom) in a simulation with cross-reactivity (step function CRISPR effectiveness with $\theta = 1$). This simulation has 1 initial phage clone and parameters $C_0 = 10^4$, $\mu = 10^{-6}$, $\eta = 10^{-4}$, and $e = 0.95$. Later times in this simulation are shown in Figure 64)

.

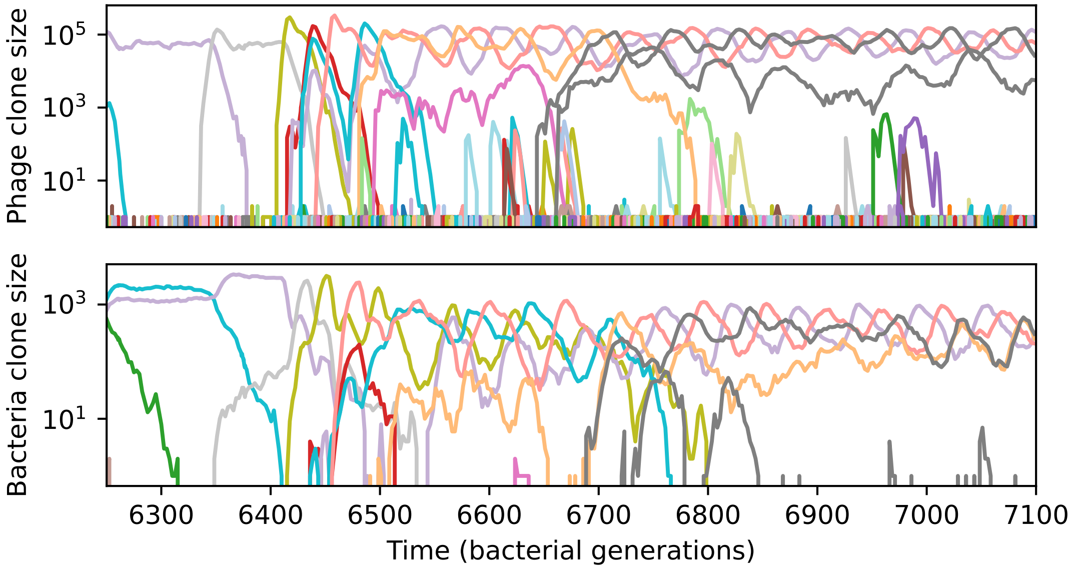

Figure 64: Phage clone size (top) and bacteria clone size (bottom) in a simulation with cross-reactivity (step function CRISPR effectiveness with $\theta = 1$) showing the switch between a traveling wave and low turnover regime. This simulation has 1 initial phage clone and parameters $C_0 = 10^4$, $\mu = 10^{-6}$, $\eta = 10^{-4}$, and $e = 0.95$. Earlier times in this simulation are shown in Figure 63).

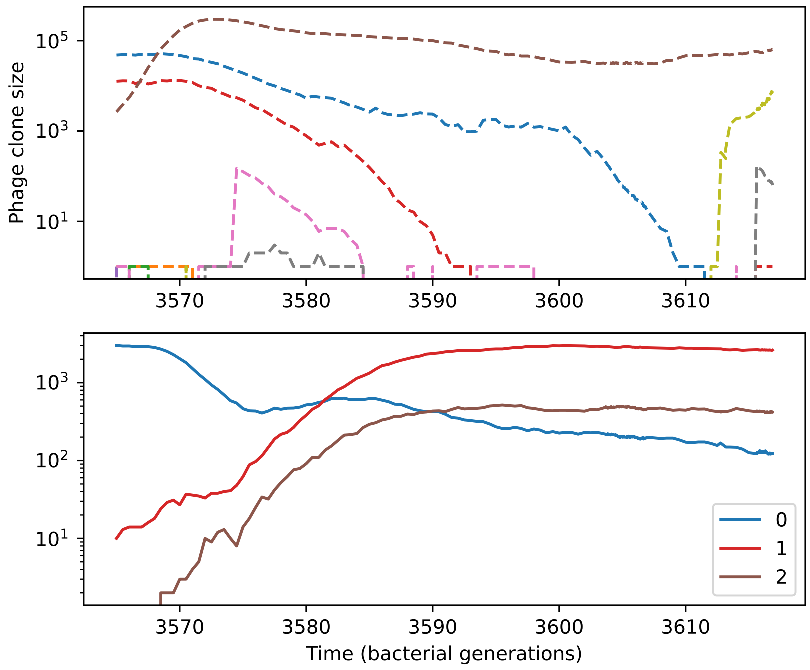

Figure 65: Phage clone size (top) and bacteria clone size (bottom) for a short time window of the simulation shown in Figure 63. Large phage and bacteria clones are numbered in the legend; these numbers correspond to the numbers in Figure 66.

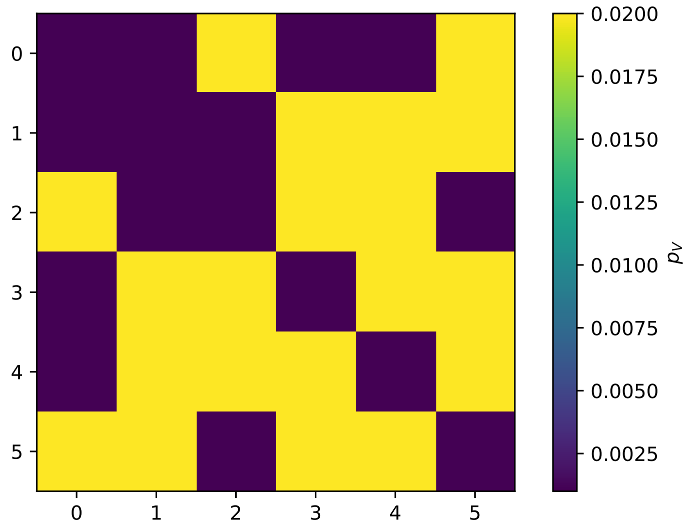

Figure 66: Phage infection success probability matrix for each clone shown in Figure 65. Dark blue is low infection success, light blue is high infection success.

and persistent when cross-reactivity is added (Figure 69C-D). Phage clones grow quickly when bacteria have low marginal immunity to their sequence, while bacteria clones grow quickly when they have high marginal immunity.

Clones grow more quickly on average in the travelling wave regime than in other regimes in the same simulation (SI Figure 67).

In main text Figure 4 we show phylogenies of four simulations with different amounts and types of cross-reactivity. We used DendroPy [32] and Toytree [33] to construct and plot phylogenies for different simulations. Figure 70 shows simulations for the same parameters as in the main text but a ten-fold higher phage mutation rate: most simulations have multiple coexisting lineages, and the traveling wave regime is less long-lived before periods of low turnover. In the low-turnover regime, a relatively small number of large clones that are all outside of each other's cross-reactivity radius can coexist for a very long time, oscillating out of phase from each other (Figures 71 and 72).

The durability and turnover of immune memory is qualitatively different in the different regimes of the simulation with cross-reactivity shown in Figure 63. We calculated time-shifted average immunity for each of the three regimes (initial regime with no turnover, traveling wave regime, and persistent oscillation regime) and found that turnover was extremely low in all but the traveling wave regime, which had a rapid decay of immune memory to zero (Figure 73). In contrast, the immune memory for the same parameters without cross-reactivity had a much more gradual long-term decay towards zero immunity (Figure 74). In the traveling wave regime, different amounts of cross-reactivity result in different time-shifted immunity patterns. Turnover is slow but smooth with exponential cross-reactivity, while turnover is much more rapid with step-function cross-reactivity (Figure 75). The differences in rate of turnover can also be seen by comparing timescale of sequence change in Figure 4.

Adding cross-reactivity appears to slightly decrease average clan sizes (Figure 76). This is what we intuitively expect, since cross-reactivity means phages are under pressure to get as far away from other clones as possible.

### 5 Stochastic clone dynamics

In this section we define $p_V(i,j)$ as given by equation 10.

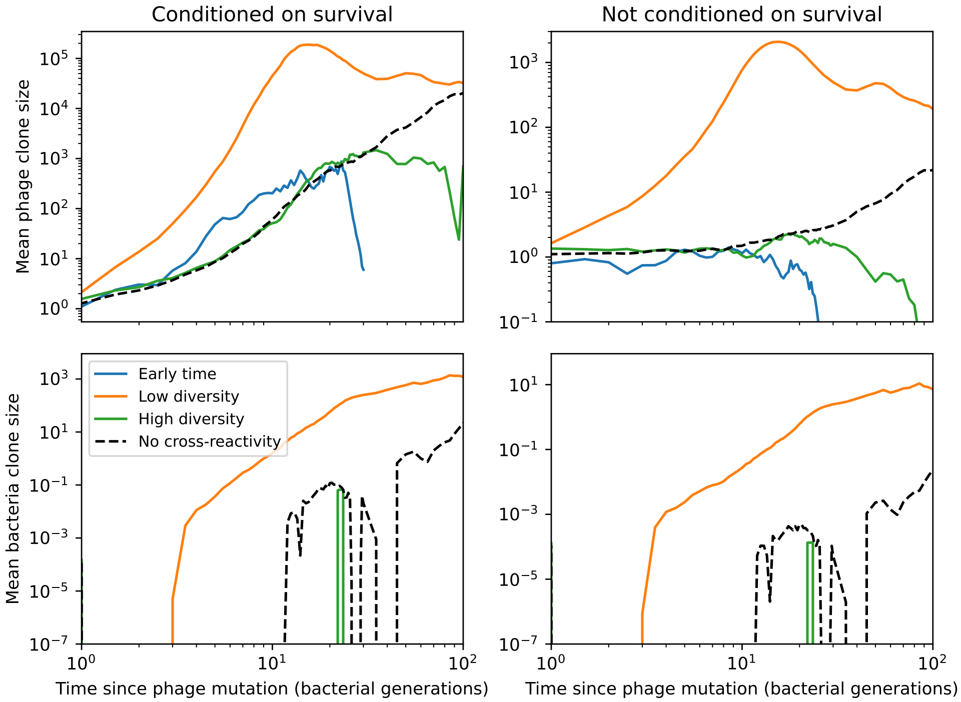

Figure 67: Mean phage clone size (top) and mean bacteria clone size (bottom) relative to the time of phage clone mutation, either normalized to surviving clones (left) or averaged over all trajectories (right) for the simulation shown in Figure 63 ($C_0 = 10^4$, $\mu = 10^{-6}$, $\eta = 10^{-4}$). Clones in the traveling wave regime (4000 to 6200 generations, orange) grow much more quickly than clones in the initial low-turnover regime (1000 to 3200 generations, blue) or the final low turnover regime (7000 to 10000 generations, green). The black dashed line is the mean clone size for a simulation with the same parameters but without cross-reactivity.

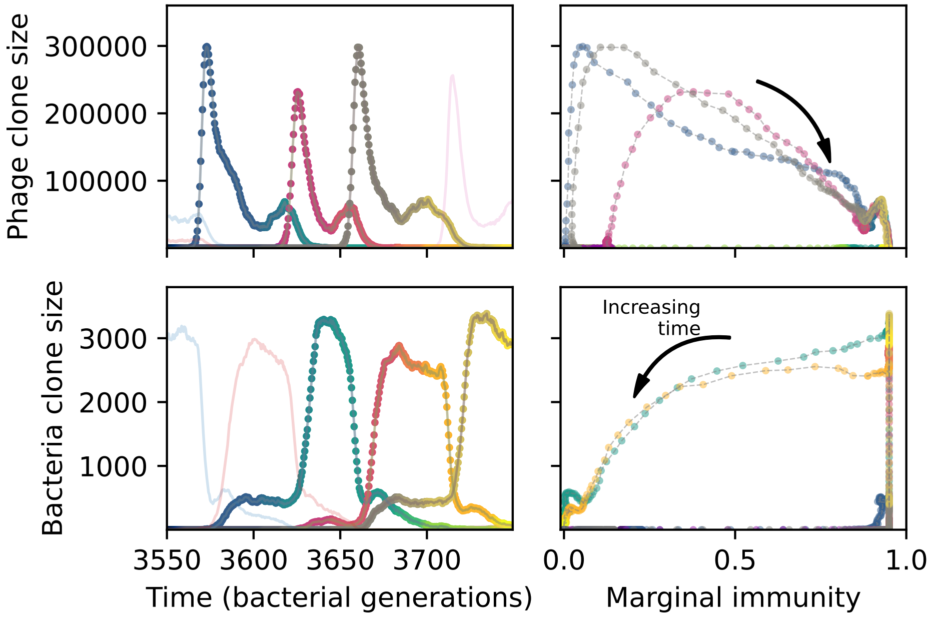

Figure 68: A slow-switching cross-reactivity regime for the simulation shown in Figure 63. (Left) A subset of clone trajectories for phages (top) and bacteria (bottom) in a simulation with cross-reactivity (step function CRISPR effectiveness with $\theta = 1$). Three trajectories are highlighted and coloured to show increasing time. The population is in a regime where matching clones are offset: because clones that are one mutation apart have the same complete immune overlap, the highlighted clones have large bacterial clone size well after the matching phage clone goes extinct. (Right) The clone size of the highlighted trajectories shown as a function of the bacterial marginal immunity against a particular phage clone (top) or the marginal immunity of a particular bacteria clone (bottom).

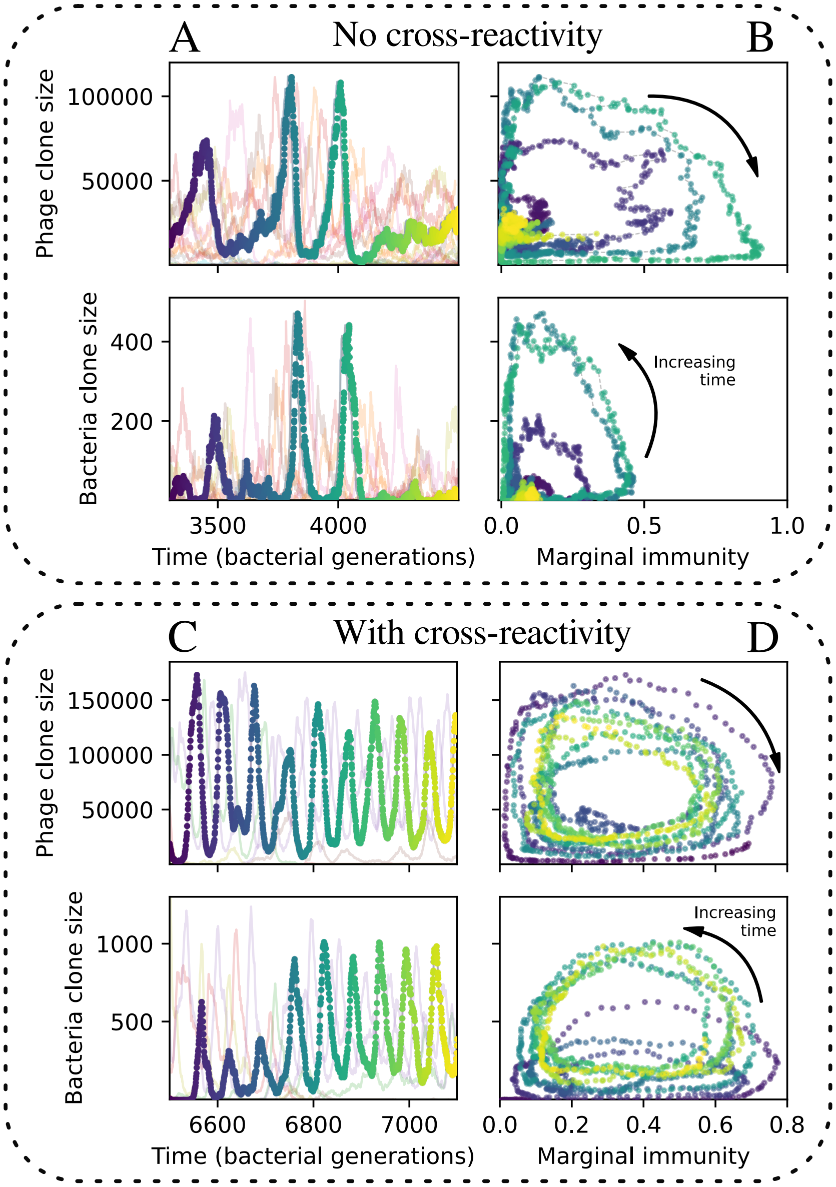

Figure 69: **Cross-reactivity leads to persistent oscillations.** (A) A subset of clone trajectories for phages (top) and bacteria (bottom) in a simulation with no cross-reactivity. Transient oscillations occur. One trajectory is highlighted and coloured to show increasing time. (B) The highlighted trajectory in (A) is shown as a function of the marginal immunity for phages (top) and bacteria (bottom). Clones experience an oscillating fitness that depends on their overlap from the other population. Arrows indicate the direction of increasing time in the oscillation. (C) A subset of clone trajectories for phages (top) and bacteria (bottom) in a simulation with cross-reactivity (step function CRISPR effectiveness with $\theta = 1$). The population is in a regime where several clones experience persistent and rapid oscillations. One trajectory is highlighted and coloured to show increasing time. (D) The highlighted trajectory in (C) is shown as a function of the marginal immunity for phages (top) and bacteria (bottom). Clones experience an oscillating fitness that depends on their overlap from the other population. Arrows indicate the direction of increasing time in the oscillation. For all simulations $C_0 = 10^4, e = 0.95, \eta = 10^{-4}$, and $\mu = 10^{-5}$.

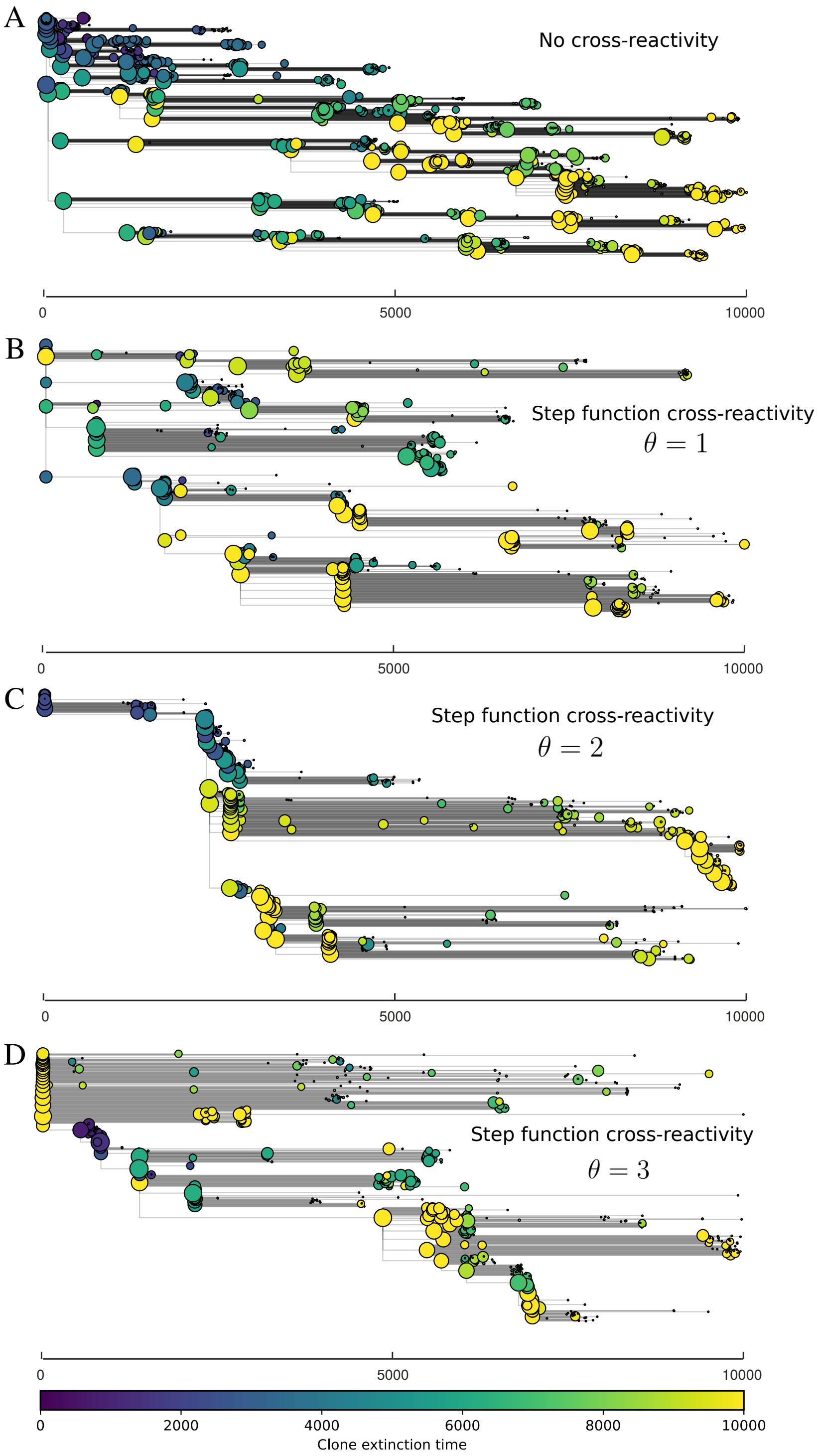

Figure 70: Phage clone phylogenies for four simulations with different cross-reactivities and a higher mutation rate than shown in main text Figure 4: no cross-reactivity (A) and step-function cross-reactivity with $\theta = 1$ (B, $\theta = 2$ (C), and $\theta = 3$ (D). All simulations share all other parameters: $C_0 = 10^4, \eta = 10^{-4}, \mu = 10^{-5}, e = 0.95$. Phage clones are plotted at the first time they pass a population size of 2 (to remove clutter from many new mutations destined for extinction), and the size of each circle is logarithmically proportional to the maximum size reached by that clone. Colours indicate the time of extinction of each clone. For each simulation with cross-reactivity, the left inset shows phage (top) and bacteria (bottom) clone sizes over time; colours indicate unique clone identities.

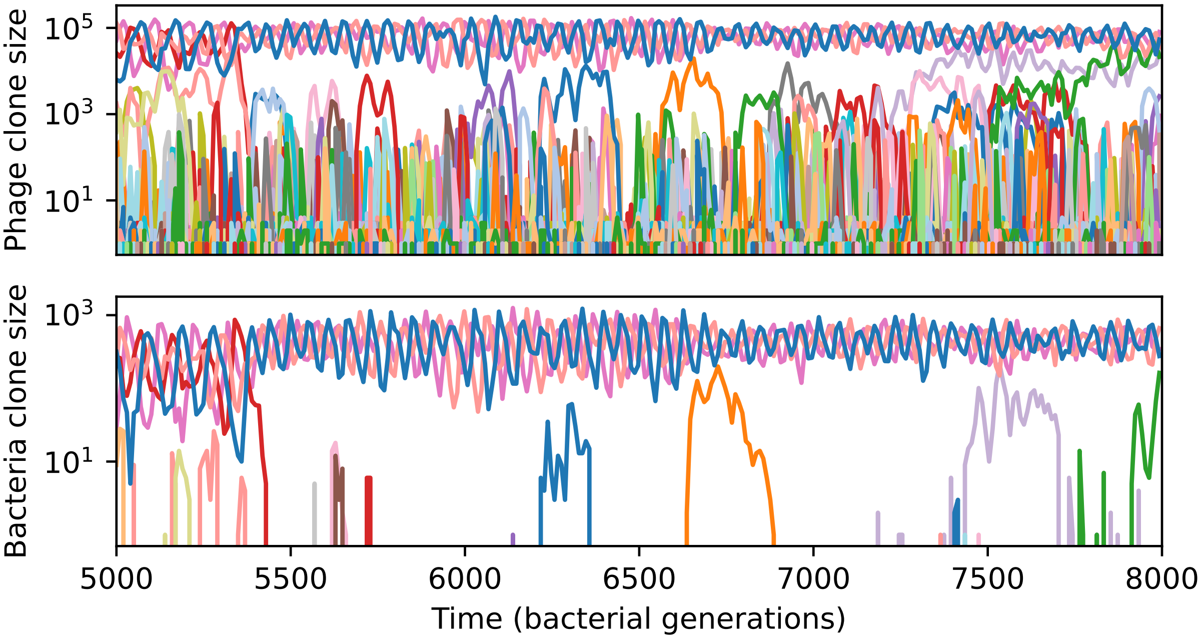

Figure 71: Phage clone size (top) and bacteria clone size (bottom) for a short time window of the $\theta = 2$ simulation shown in Figure 70C. Each of the large trajectories oscillating out of phase between 5000 and 7000 generations is at least three mutations away from all of the others; they are all outside each other's cross-reactivity radius.

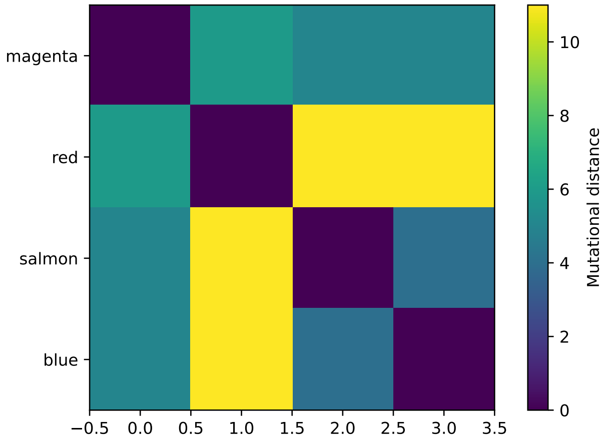

Figure 72: Matrix of mutational distance between each of the four largest phage clones shown in Figure 71; colours of those trajectories are labeled on the y axis. Each clone is at least three mutations away from all other large clones.

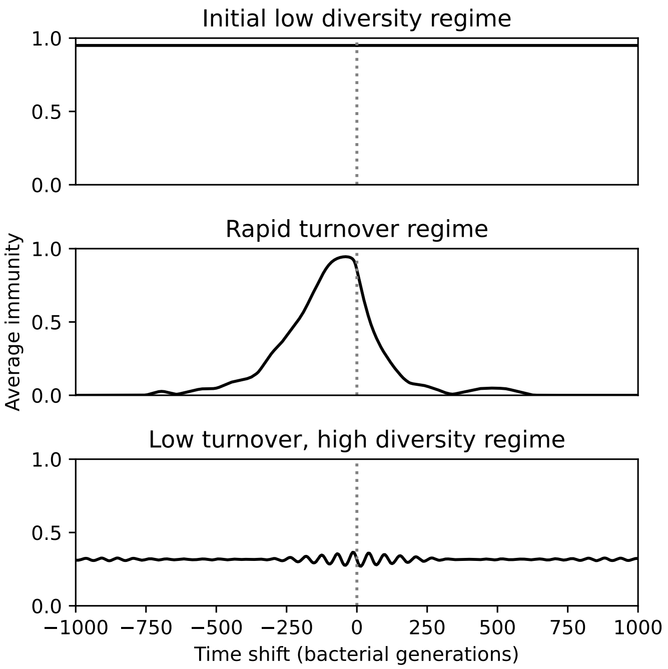

Figure 73: Time shifted average immunity for three regimes of the simulation shown in Figure 63 ($C_0 = 10^4$, $\mu = 10^{-6}$, $\eta = 10^{-4}$, and $e = 0.95$, step-function cross-reactivity with $\theta = 1$). The initial low-diversity regime (1000 to 3200 generations) and the low turnover, high diversity regime (7000 to 10000 generations) had extremely low turnover, while the traveling wave regime (4000 to 6200 generations) had high average immunity near 0 delay that rapidly decayed to zero both in the past and future.

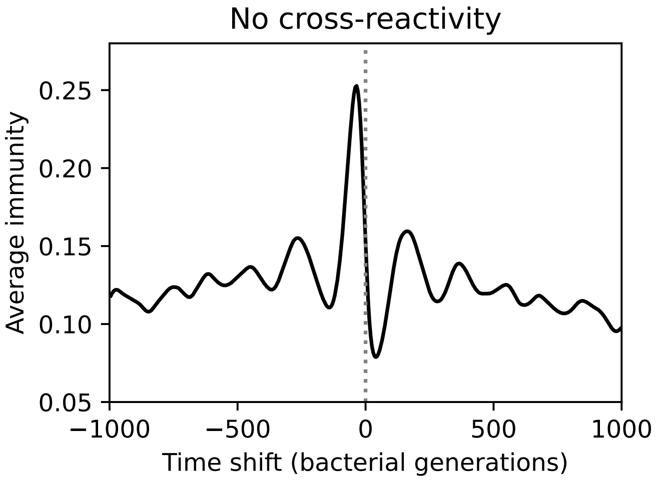

Figure 74: Time shifted average immunity for the corresponding simulation to Figure 73 without cross-reactivity ($C_0 = 10^4$, $\mu = 10^{-6}$, $\eta = 10^{-4}$, and $e = 0.95$). Peak average immunity is low because of high diversity, and immunity decays very gradually to zero in both the past and future.

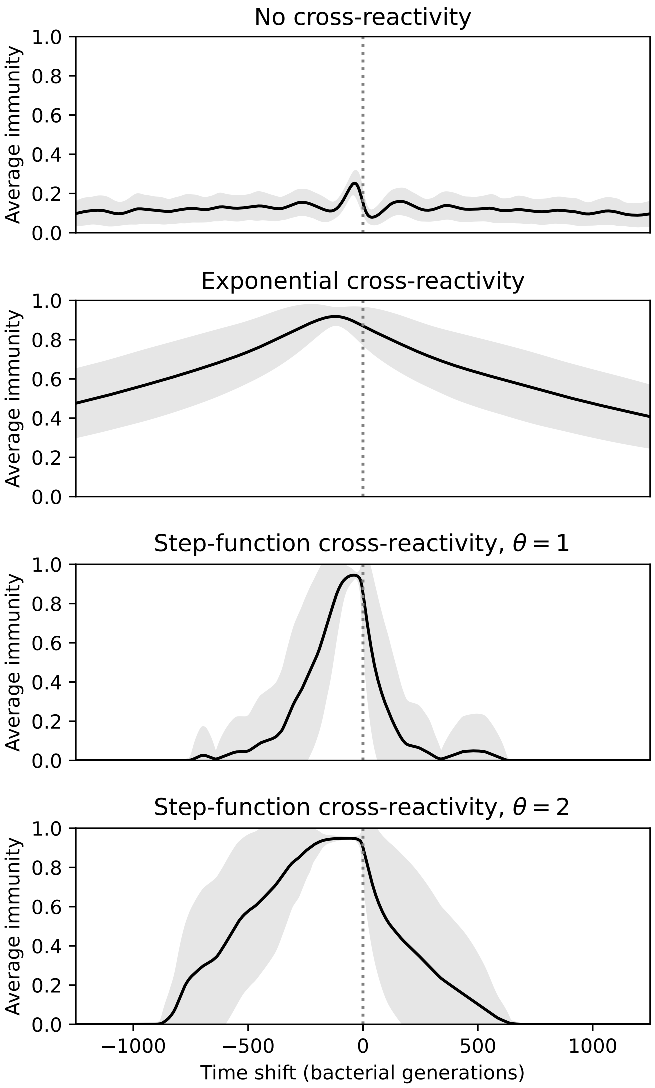

Figure 75: Time shifted average immunity for four simulations with the same parameters but different types of cross-reactivity: no cross-reactivity (top), exponential cross-reactivity (middle top), step-function cross-reactivity with $\theta = 1$ (middle bottom) and $\theta = 2$ (bottom). Shared parameters are $C_0 = 10^4$, $\mu = 10^{-6}$, $\eta = 10^{-4}$, and $e = 0.95$. Only the traveling-wave regime of each simulation with cross-reactivity was used to compare turnover in this regime.

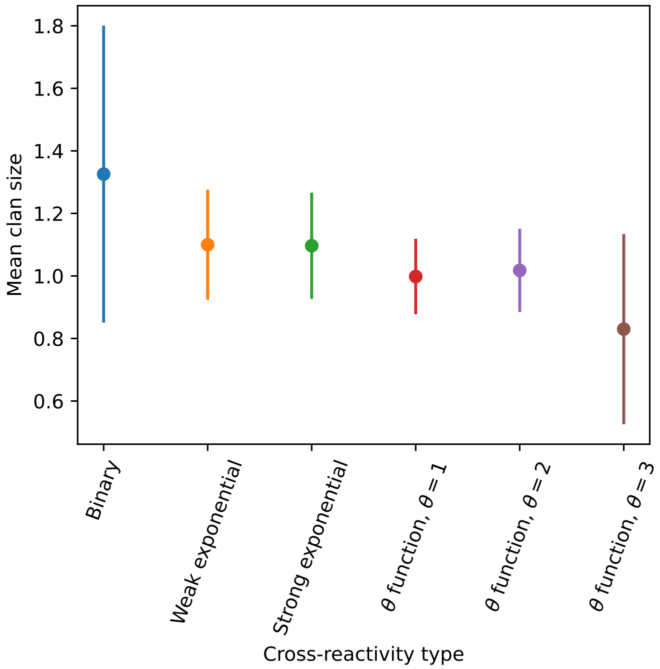

Figure 76: Average clan size across simulations with different parameters for different degrees of cross-reactivity with $m_{\text{init}} = 10$.

##### 5.0.1 Clone fitness

We investigated the fitness of new phage mutants to understand the effect of bacterial spacer acquisition on phage mutant growth. We define the fitness of phage clones to be their per-capita average growth rate: their average growth rate in bacterial generations divided by their average size. We calculate the fitness from simulation data by calculating the mean phage clone size conditioned on survival (as in Figure 3), then taking the time derivative, then dividing by the mean phage clone size. (In principle, this is equivalent to first taking the time derivative of each individual clone trajectory and then averaging across all trajectories, but we found that edge effects from trajectories that go extinct skewed the result, so we first average across all trajectories before taking a derivative.) Phage clone fitness over time in a single simulation is plotted in Figure 77 (orange markers).

We calculate the theoretical phage clone fitness by taking the time derivative of the predicted mean phage clone size and dividing by the predicted mean clone size. The predicted mean phage clone size is piecewise-defined at short times as the numerical solution of the system of equations 13 and 14, and at long times as the numerical solution steady-state clone size (equation 18). This prediction is plotted as a solid black line in Figure 77.

New phage mutants have a positive growth rate on average (initial fitness $> 0$), and their growth rate drops to zero on average as bacteria acquire matching spacers and gain immunity to new mutants. We can define the time or size at which phage clones become "established", after which they are safe from rapid stochastic extinction and behave like neutral clones (having a fitness that is close to 0). We define one minus the long-time limit of the probability of phage clone extinction (equation 134) as the probability of establishment for new phage clones:

$$P_{\text{est}} = 1 - P_0^{N_{\text{est}}} = 1 - \left(1 - \frac{2s_0}{B(s_0 + \delta_0)}\right)^{N_{\text{est}}} \tag{90}$$

where $s_0 = \alpha B p_V - \alpha n_B - F$ is the average initial growth rate of phage clones and $\delta_0 = F + \alpha n_B(1 - p_V)$

is the average initial death rate of phage clones (more in Section 5.3). The establishment clone size can then be defined as the value of $N_{\text{est}}$ for which $P_{\text{est}} \approx 1$. Since $N_0 \frac{2s_0}{B(s_0+\delta_0)}$ is not necessarily small, we approximate $P_{\text{est}}$ as an exponential function and set $N_{\text{est}}$ as the scale of the exponent:

$$\left(1 - \frac{2s_0}{B(s_0+\delta_0)}\right)^{N_{\text{est}}} \approx e^{-N_0 \frac{2s_0}{B(s_0+\delta_0)}} = e^{-\frac{N_0}{N_{\text{est}}}}$$

$$N_{\text{est}} \approx \frac{B(s_0+\delta_0)}{2s_0} \tag{91}$$

Equation 91 is the size at which phage clones become established on average (where $P_0 \approx 1/e$). This is plotted as a horizontal dashed line for one simulation in Figure 77. If $B = 2$ (birth-death with no bursts and positive selection), then $N_{\text{est}} = \frac{s_0+\delta_0}{s_0} \approx \delta_0/s_0$. This is the expected result for a simple birth-death process with selection as given in [27].

We can also find the time at which phage clones reach the establishment size. In the absence of matching bacterial spacers, new phage mutants grow deterministically as $n_V^i(t) = e^{s_0 t}$. We condition on survival by dividing by the probability of establishment; for this calculation we use the long-time probability of establishment given by equation 131 with $s = s_0$ and $\delta = \delta_0$. This curve does not match the measured growth at short times, but in the region of phages reaching their establishment size it agrees well (solid pink line in Figure 3).

With these assumptions, the growth curve for phage clones is

$$n_V^i(t) \approx \frac{e^{s_0 t}}{1 - \frac{(B(\delta_0+s_0)-2s_0)(e^{s_0 t}-1)}{2s_0+B(e^{s_0 t}-1)(\delta_0+s_0)}} = 1 + \frac{B(e^{s_0 t}-1)(\delta_0+s_0)}{2s_0} \tag{92}$$

Replacing $n_V^i$ with the establishment clone size (91) and solving for $t$, we find

$$t_{\text{est}} = \frac{1}{s_0} \ln\left(\frac{2B(s_0+\delta_0)-2s_0}{B(s_0+\delta_0)}\right) = \frac{1}{s_0}\left[\ln 2 + \ln\left(1 - \frac{s_0}{B(\delta_0+s_0)}\right)\right] \tag{93}$$

Now $s_0/(B(\delta_0+s_0)) << 1$, so the second logarithm can be approximated as $-s_0/(B(\delta_0+s_0))$.

$$t_{\text{est}} \approx \frac{1}{s_0}\left[\ln 2 - \frac{s_0}{B(\delta_0+s_0)}\right] \tag{94}$$

We can further approximate by dropping the second term entirely since $\ln 2 >> s_0/(B(\delta_0+s_0))$.

$$t_{\text{est}} \approx \frac{\ln 2}{s_0} \tag{95}$$

We get the same approximate value of $t_{\text{est}}$ if we take $B = 2$ directly and assume $s_0 << 1$. Equation 95 is also numerically close to the mean establishment time calculated by Desai and Fisher for positive selection, $\langle \tau_{\text{est}} \rangle = \frac{\gamma}{s}$ (where $\gamma \approx 0.577216$) [27]. This implies that the presence of a burst size $B > 2$ does not dramatically change the dependence of the time to establishment on phage fitness. The phage growth rate $s_0$ does still depend on $B$, however. (Note that Desai and Fisher define establishment time differently than we do, so our cases are not directly comparable.) Equation 95 is plotted as a vertical dashed line in Figure 77.

The initial fitness $f_0$ of a new phage mutant can be computed analytically by using a different early-time approximation for $n_V^i(t)$, this time conditioning on survival using the short-time approximation for $P_0(t)$ given by equation 138.

$$n_V^i(t) \approx \frac{e^{s_0 t}}{1 - \frac{\delta_0}{\beta_0+\delta_0}\left(1 - e^{-(\beta_0+\delta_0)t}\right)} = \frac{e^{s_0 t}(\beta_0+\delta_0)}{\beta_0 + \delta_0 e^{-(\beta_0+\delta_0)t}} \tag{96}$$

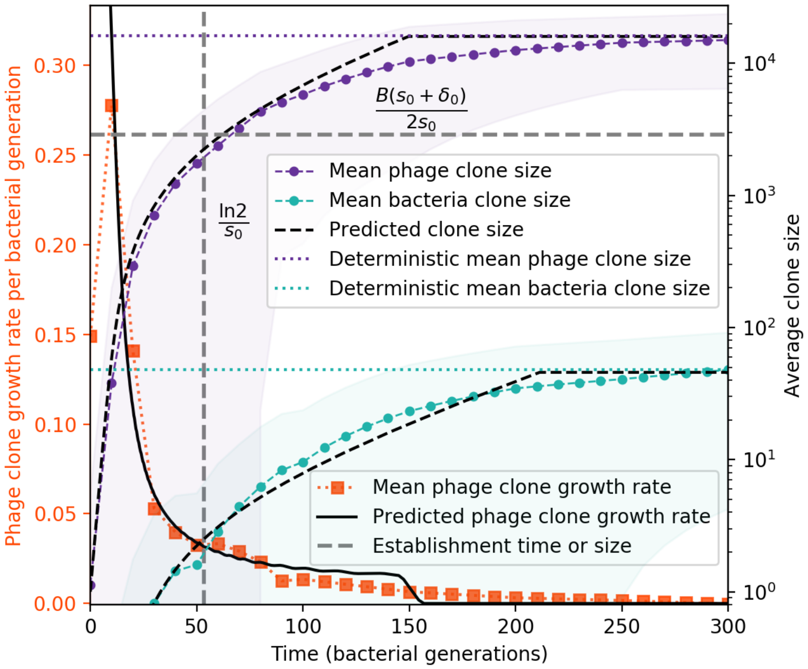

Figure 77: Average phage and bacteria clone size vs. time (right vertical axis, purple and green markers) and average phage clone growth rate vs. time (left vertical axis, orange markers). Markers are the average over all clone trajectories after steady-state in a single simulation. Phage clones appear at size 1 and grow until they reach the deterministic mean phage clone size on average. Once a phage clone becomes large, bacteria encounter it often enough to acquire a matching spacer, and bacteria clones grow until they reach their deterministic mean clone size. New phage mutants have a selective advantage (fitness $> 0$, positive growth rate) until they reach the deterministic mean clone size, at which point they evolve neutrally (fitness $\approx 0$). Individual clone trajectories are highly variable, leading to a large standard deviation on the mean phage clone fitness and a fitness trend which is only evident on average. The predicted clone size is piecewise-defined as the theoretical clone trajectory until the theoretical trajectory reaches the predicted mean clone size.

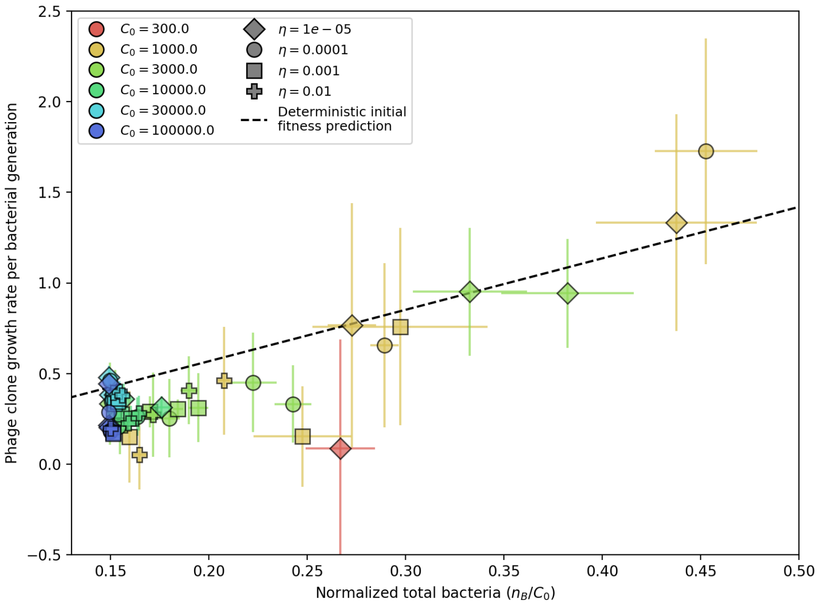

Figure 78: Phage clone initial growth rate vs. total bacteria normalized by the initial nutrient concentration $C_0$. Phage clone growth rate is as defined in Figure 77 - for each simulation, the average phage clone growth rate is the derivative of the average phage clone size, averaged across all trajectories after steady-state; plotted points and error bars are the average across 3 or more simulations. The phage initial fitness depends slightly on the phage mutation rate (mutants decrease the growth rate of a particular phage clone), but this dependence is slight enough that all mutation rates collapse onto the theoretical line. Here we plot data with $\mu = 10^{-6}$. The phage clone initial growth rate also does not depend on $e$ or $\eta$ because new phage mutants see the bacteria population as if it did not have any CRISPR immunity. The theoretical initial phage clone growth rate is given by equation 98. The effective lower bound of $n_B/C_0$ is set by the steady-state population size without CRISPR immunity: $n_B/C_0 = \frac{fg}{\alpha(Bp_V - 1)} \approx 0.15$.

To estimate the initial fitness, we differentiate 96 with respect to $t$, divide by $n_V^i(t)$ to get the per-capita growth rate, and evaluate at $t = 0$:

$$f_0 = \frac{1}{n_V^i(t)} \frac{dn_V^i(t)}{dt}|_{t=0} = s_0 + \frac{\delta_0(\beta_0 + \delta_0)\mathrm{e}^{-(\beta_0+\delta_0)t}}{\beta_0 + \delta_0 \mathrm{e}^{-(\beta_0+\delta_0)t}}|_{t=0} = s_0 + \delta_0 \tag{97}$$

Interestingly, if we had not conditioned on survival, the initial per-capita growth rate would simply be $s_0$: conditioning on survival increases the apparent growth rate of phage clones by effectively ignoring their death rate.

We can evaluate $s_0 + \delta_0$ in terms of our original parameters for insight. We replace $B$ with $B\mathrm{e}^{-\mu L}$ to capture the decrease in phage growth clone growth due to mutations away from a clone.

$$f_0 = s_0 + \delta_0 = \alpha p_V n_B (B\mathrm{e}^{-\mu L} - 1) \tag{98}$$

The initial fitness does not directly depend on characteristics of CRISPR immunity such as $\eta$ and $e$ because new phage mutants see the bacterial population as if it did not have any CRISPR immunity; $f_0$ depends only on the total bacterial population size (SI Figure 78).

#### 5.1 Bacteria clone dynamics

To solve for the dynamics of individual bacteria clones, we write a one-dimensional master equation just for $n_B^j$ (equation 99).

$$\begin{aligned}\frac{dP_n}{dt} =&(n+1)P_{n+1}[F+r+\alpha p_V(n_V - e n_V^i)] \\ &+ (n-1)P_{n-1}[gC] \\ &+ P_{n-1}[\alpha\eta n_B^0 n_V^i(1-p_V)] \\ &- nP_n[F+r+\alpha p_V(n_V - e n_V^i) + gC] \\ &- P_n[\alpha\eta n_B^0 n_V^i(1-p_V)]\end{aligned} \tag{99}$$

For brevity we write $P_n = P_{n_B^i}(t|N_0)$, the probability of having $n$ bacteria of type $i$ at time $t$ given $N_0$ bacteria of type $i$ at $t=0$.

Bacteria clone growth ($gC$), phage predation ($\alpha p_V(n_V - e n_V^i)$), outflow ($F$), and spacer loss ($r$) all depend on the number of bacteria $n$, but spacer acquisition ($\alpha\eta n_B^0 n_V^i(1-p_V)$) adds new bacteria independent of the current size of the clone.

We assume that the total population is in steady state so that the total population sizes $n_V$, $C$, and $n_B^0$ are constant. In general, $n_V^i$ is time-dependent and varies for each clone $i$, but we will assume that it is also constant at steady-state.

This equation is very nearly identical in form to the bacteria clone size equation solved in our previous work [3] as well as the clone size equation described in [34], and we repeat our derivation of the solution here in brief.

We solve equation 99 using a generating function approach: $G(z,t) = \sum_n z^n P_n(t)$. Multiplying equation 99 by $\sum_n z^n$, we get the corresponding generating function partial differential equation:

$$\partial_t G(z,t) = \partial_z G(z,t)\left(d + bz^2 - (b+d)z\right) + DG(z-1) \tag{100}$$

Here $b$ and $d$ are the birth and death rates for bacterial clones: $b = gC$ and $d = F+r+\alpha p_V(n_V - e n_V^i)$. $D$ is the rate of spacer acquisition from naive bacteria: $D = \alpha\eta n_B^0 n_V^i(1-p_V)$.

We solve equation 100 using the method of characteristics [35]. We parameterize the function $G(z,t)$ with a new variable $x$. Applying the chain rule:

$$\partial_x G(z(x), t(x)) = \frac{\partial G}{\partial z}\frac{\partial z}{\partial x} + \frac{\partial G}{\partial t}\frac{\partial t}{\partial x} \tag{101}$$

Comparing equation 100 with equation 101 gives the following characteristic equations:

$$\frac{\partial t}{\partial x} = 1 \tag{102}$$

$$\frac{\partial z}{\partial x} = (1-z)(bz-d) \tag{103}$$

$$\frac{\partial G}{\partial x} = DG(z-1) \tag{104}$$

From equation 102 we see $t = x + c_1$, so we can choose $t_0 = c_1 = 0$ and replace $x$ with $t$ going forward.

Solving the characteristic equation for $z$ by integrating both sides gives equation 105.

$$\frac{1-z}{d-bz}\mathrm{e}^{(b-d)t} = c_2 \tag{105}$$

At $t = 0$, $z$ will pass through some point $z_0$, so we have the initial condition $z(0) = z_0$. With $z_0$ in equation 105 at $t = 0$, we get equation 106, where $c_2$ is given by equation 105.

$$z_0 = \frac{c_2 d - 1}{c_2 b - 1} \tag{106}$$

The variation of $G$ along the $z - t$ curve is

$$\frac{\partial G}{\partial z} = -\frac{DG(z-1)}{(1-z)(bz-d)} = -\frac{DG}{(bz-d)} \tag{107}$$

Integrating both sides, we get

$$G(z) = \Omega(c_2)(bz-d)^{-\frac{D}{b}}$$

The constant $\Omega$ is a function of the characteristic $z$-$t$ curve (equation 105). To find the particular form of $\Omega(c_2)$, we apply the initial condition $P_{N0}(0) = 1$ which gives $G(z, 0) = z^{N_0}$, meaning that the clone starts at size $N_0$ at time $t = 0$.

$$G(z,0) = z^{N_0} = \Omega\left(\frac{1-z}{d-bz}\right)(bz-d)^{-\frac{D}{b}}$$

Let $\xi = \frac{1-z}{d-bz}$, therefore $z = \frac{\xi d-1}{\xi b-1}$.

$$\Omega(\xi)\left(b\left(\frac{\xi d-1}{\xi b-1}\right)-d\right)^{-\frac{D}{b}} = \left(\frac{\xi d-1}{\xi b-1}\right)^{N_0}$$

Solving for $\Omega(\xi)$:

$$\Omega(\xi) = \left(\frac{\xi d-1}{\xi b-1}\right)^{N_0}\left(b\left(\frac{\xi d-1}{\xi b-1}\right)-d\right)^{\frac{D}{b}}$$

The full solution for $G(z,t)$ can be written by replacing the constant $\Omega(c_2)$ with the expression for $\Omega(\xi)$ and replacing $\xi$ with $\xi\epsilon$, where $\epsilon = \mathrm{e}^{(b-d)t}$ is the time-dependent part of the $z - t$ curve.

$$G(z,t) = (bz-d)^{-\frac{D}{b}}\left(\frac{\xi\epsilon d-1}{\xi\epsilon b-1}\right)^{N_0}\left(b\left(\frac{\xi\epsilon d-1}{\xi\epsilon b-1}\right)-d\right)^{\frac{D}{b}}$$

Finally, replacing $\xi$ with $\frac{1-z}{d-bz}$, we get

$$G(z,t) = (bz-d)^{-\frac{D}{b}}\left(\frac{(1-z)\epsilon d+bz-d}{(1-z)\epsilon b+bz-d}\right)^{N_0}\left(b\left(\frac{(1-z)\epsilon d+bz-d}{(1-z)\epsilon b+bz-d}\right)-d\right)^{\frac{D}{b}}$$

$G(1,t) = \sum_n P_n(t) = 1$, meaning that the total probability is conserved.
Assuming $d > b$, the limit as $t \to \infty$ of $G(z,t)$ is

$$G(z) = \left(\frac{bz-d}{b-d}\right)^{-\frac{D}{b}}$$

The limit is independent of the initial clone size $N_0$ as we expect.
We can construct $P_n$ at steady-state by taking successive derivatives of $G(z)$: $P_n = \frac{1}{n!}\frac{\partial^n G}{\partial z^n}|_{z=0}$

$$P_n = \frac{1}{n!d^n}\left(\frac{d-b}{d}\right)^{D/b}\prod_{i=1}^{n}[D+(i-1)b] \tag{108}$$

$$P_0 = \left(\frac{d-b}{d}\right)^{D/b}$$

This is a negative binomial distribution with parameters $D/b$ and $b/d$. We can re-write this expression using Stirling's approximation for $n!$ to facilitate evaluation at large $n$.

$$P_n = \frac{1}{\sqrt{2\pi n}} \exp\left[\frac{D}{b}\ln\left(\frac{d-b}{d}\right) + \sum_{i=1}^{n} \ln\left(\frac{e}{nd}(D+(i-1)b)\right)\right] \tag{109}$$

Equation 109 is an analytic expression describing the steady-state spacer abundance distribution that results from our simulations. To compare this prediction with our simulations, we assume that $n_V^i$ on average is equal to $n_V/m$, where $n_V$ is the predicted total phage population size and $m$ is the number of large phage clones approximated by the predicted bacterial diversity.

Figure 79 compares the analytic distribution to the steady-state spacer clone size distribution from our simulations at several values of the spacer acquisition probability $\eta$. The theoretical prediction captures the qualitative impact of increasing $\eta$ fairly well: as $\eta$ increases, the clone size distribution gains a more pronounced peak.

The discrepancy between the theoretical prediction and simulation data at high $\eta$ results in part from the predicted large phage clone size being larger than the measured large phage clone size in simulations. This can happen when the predicted number of clones $m$ is smaller than the simulation result. To assess whether this influenced our prediction, we also used Maximum Likelihood Estimation to calculate the value of $n_V^i$ that gave the best fit between equation 109 and the data; for the two largest values of $\eta$, this does return a smaller value of $n_V^i$ and hence a distribution peak further to the left.

The previous two calculations assumed that the phage clone size is single-valued and constant in time. In reality, however, the phage clone size is both broadly distributed (Figure 26) and changing in time (Figure 77). We relax the first assumption by calculating an average bacterial distribution using the observed distribution of phage clones: we solve equation 109 at each observed large phage clone size, then average across the distribution of clone sizes to calculate $P(n_B^i) = P(n_B^i|n_V^i)P(n_V^i)$. (The large phage clone distribution is given by equation 24.) This average distribution more accurately predicts the presence of small bacterial clones at high $\eta$, but it actually behaves worse than the single-$n_V^i$ solution at small $\eta$. This is related to the deviation of the number of large phage clones from the number of bacterial clones at small $\eta$ (Figure 24): at small $\eta$, bacteria don't acquire spacers as readily and so phage clones experience clonal interference largely without bacterial influence. The average distribution then predicts more small bacterial clones than there are because the large phage clone distribution includes intermediate phage clone sizes that the bacteria don't end up acquiring spacers from. Essentially, the bacteria "see" fewer phage clones than the theory predicts, so the observed bacteria clone size distribution has fewer smaller clones than predicted.

Equation 109 can be approximated for large $n$ as a gamma distribution:

$$P_n \approx \frac{\left(1-\frac{b}{d}\right)^{\frac{D}{b}}}{\Gamma\left(\frac{D}{b}\right)} \mathrm{e}^{-\ln(d/b)n} \left(\frac{1}{n}\right)^{1-\frac{D}{b}} \tag{110}$$

This is a gamma distribution with shape parameter $\frac{D}{b}$ and rate parameter $\ln(d/b)$. Note that $\left(1-\frac{b}{d}\right)^{\frac{D}{b}} \approx \ln(d/b)^{\frac{D}{b}}$, consistent with the canonical form of the gamma distribution. The shape parameter $D/b$ describes the relative balance between spacer acquisition and growth: if $D/b > 1$, then spacer acquisition is the dominant means by which bacterial clones grow. This often also means that the clone size distribution has a peak at clone size $> 1$ (provided $d \gtrsim b$). Specifically, the mode of the distribution is greater than 1 if $\frac{D-b}{b} > \frac{d-b}{b}$. The rate parameter describes the decay constant of the exponential distribution resulting if the shape parameter equals 1.

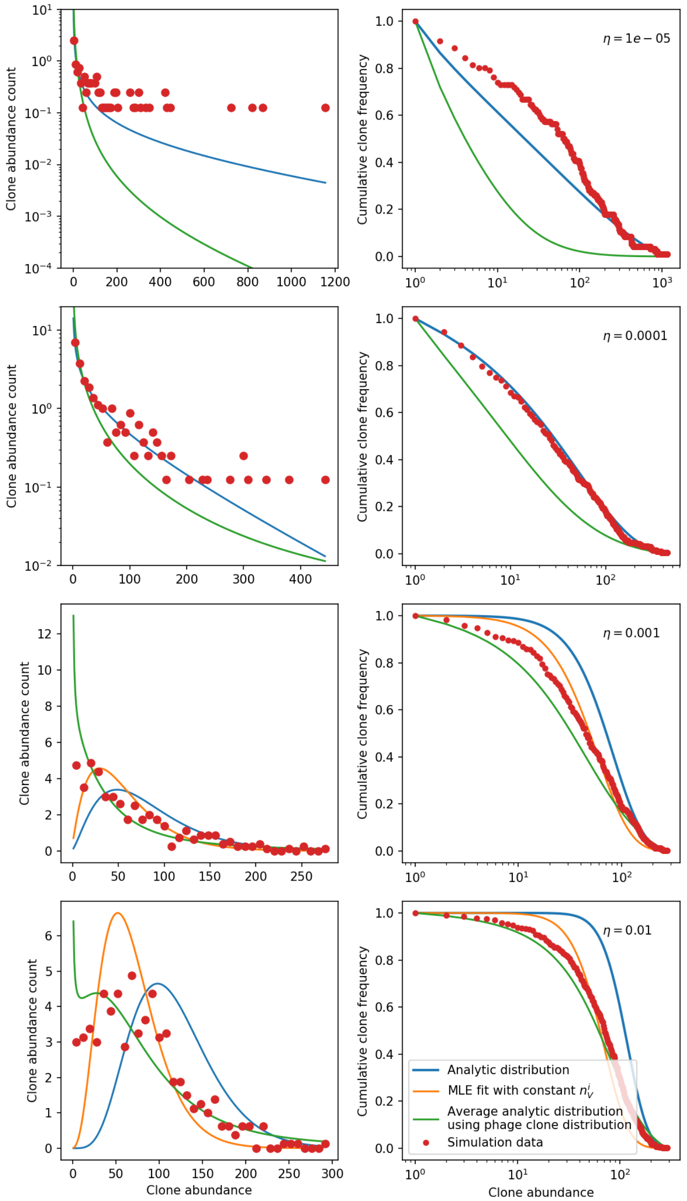

Figure 79: Clone size histograms (left) and cumulative distributions (right) for four different values of the spacer acquisition probability $\eta$. In all simulations $C_0 = 10^4$, $e = 0.95$, and $\mu = 3 \times 10^{-6}$. We sample 30 evenly-spaced time points between 2000 and 10000 bacterial generations and combine the clone sizes at each of the sampled points to create the clone size distributions plotted. Solid lines show three different theoretical solutions. The solid blue line is given by equation 109 with all population quantities predicted from solving the system of equations 5 - 9 with $m$ given by equation 28 and $n_V^i = n_V/m$. The solid orange line is given by equation 109, with the value $n_V^i$ determined by Maximum Likelihood Estimation to give the best fit to the data. For the two largest values of $\eta$, the value of $n_V^i$ returned by the MLE fit is smaller than the theoretical value of $n_V^{i\,*}$, while for the two smallest values of $\eta$ the MLE value of $n_V^i$ is larger. For large enough values of $n_V^i$, the bacteria clone death rate $d$ is smaller than the birth rate $b$ which violates the assumptions used to derive equation 109. This happens for the MLE fit at the two smallest $\eta$ values and hence no MLE solution is plotted. The solid green line is an average distribution calculated by solving equation 109 at each observed large phage clone size and averaging across the distribution of clone sizes; i.e. $P(n_B^i) = P(n_B^i | n_V^i) P(n_V^i)$. The large phage clone distribution is given by equation 24.

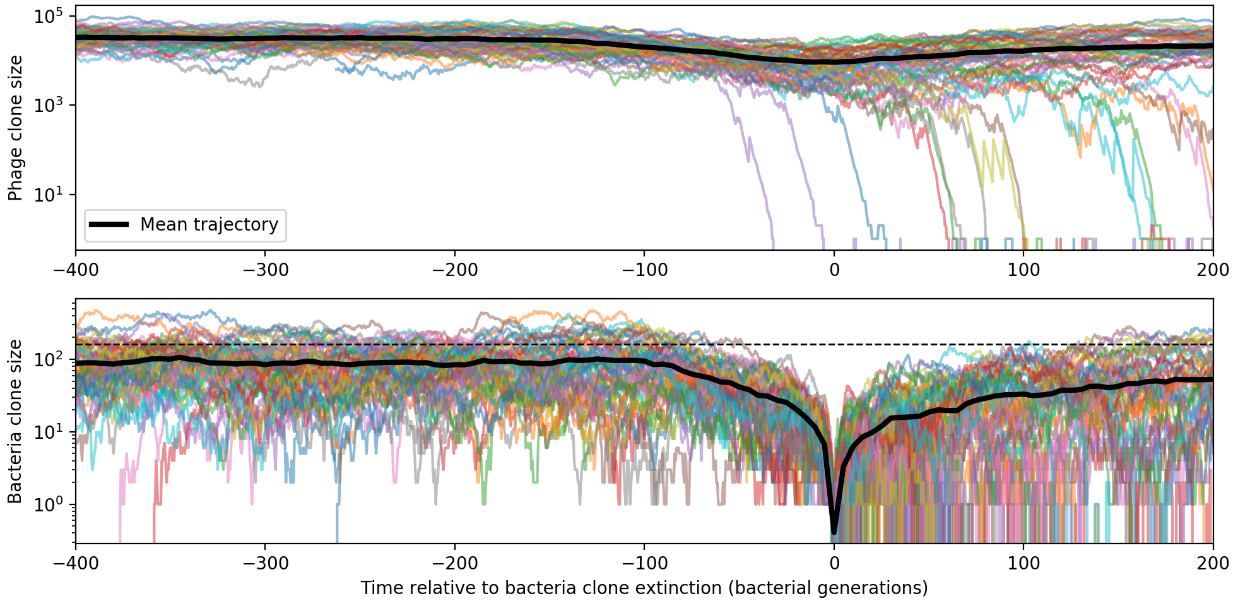

Figure 80: Bacteria and phage clone trajectories aligned to the time at which bacteria trajectories go extinct. Bacteria trajectories are included if they reach size ${n_B^i}^*$ given by equation 17 and all corresponding phage trajectories are plotted. In this simulation $C_0 = 3 \times 10^4$, $e = 0.8$, $\eta = 10^{-3}$, and $\mu = 10^{-6}$.

##### 5.1.1 Bacteria clone extinction

When a matching phage clone exists in the population, bacteria with a particular spacer have a fitness advantage if they encounter that phage. Once the matching phage goes extinct, bacteria tend to quickly go extinct as well; in fact, bacteria often go extinct *before* their matching phage clone (Figure 80). To understand why this might be, we derive a theoretical prediction for the mean time to extinction under the assumption that bacterial clones evolve neutrally once they become large. This prediction does describe the extinction time distribution well in some regimes, but it underestimates the time to extinction at large total population sizes. If the neutral assumption is valid, this means that bacterial clones go extinct stochastically and are not necessarily driven to extinction by the extinction of their matching phage clone. This is true at small-to-medium total population sizes ($C_0 \leq 10^4$). On the other hand, when population sizes are large, bacteria clones go extinct more slowly than neutral theory would predict, likely because they are still being challenged by a matching phage and are also able to acquire spacers from that phage clone.

We calculate the time to extinction using the backward master equation corresponding to equation 99. The backwards equation is an equation for the time to extinction $T_n$ from a given state. Instead of working with frequencies, we write this in terms of the number of bacteria belonging to a clone, $n$. Here $b = gC$, $d = F + r + \alpha p_V(n_V - e n_V^i)$, and $D = \alpha \eta n_B^0 n_V^i (1 - p_V)$.

$$T_n - \Delta t = bn\Delta t T_{n+1} + D\Delta t T_{n+1} + dn\Delta t T_{n-1} + (1 - bn\Delta t - D\Delta t - dn\Delta t)T_n \tag{111}$$

Notice that the rate terms in the backward equation depend only on $n$, not on $n-1$ or $n+1$ like in the forward master equation. The forward equation is a sum of all the ways in which the system could end up at state $n$ at time $t$ from where it could have been at time $t - \Delta t$, so those rates depend on the other states. The backward equation goes in the other direction, looking backwards: it is a sum of all the ways in which the system *could have been* in state $n$ at time $T$ now that time $\Delta t$ has elapsed.

Rearranging equation 111, we arrive at equation 112.

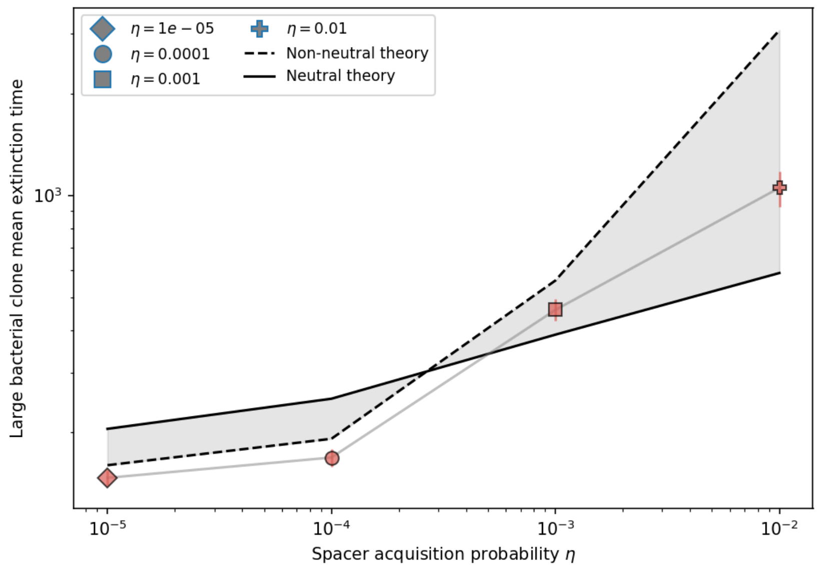

Figure 81: Mean time to extinction for bacterial clones after reaching size ${n_B^i}^*$ as a function of $\eta$ for $C_0 = 10^4$, $e = 0.95$, and $\mu = 3 \times 10^{-6}$. The solid line is given by equation 115 with $n = {n_B^i}^*$, and the dashed line is given by numerically solving equation 113.

$$-1 = (bn + D)T_{n+1} + dnT_{n-1} - (bn + dn + D)T_n \tag{112}$$

For boundary conditions, we have $T(n = 0) = 0$ (time to extinction is 0 when already extinct) and $\frac{dT}{dn}|_{n=n_B^s} = 0$ (reflecting boundary at $n = n_B^s$). The reflecting boundary is harder to justify because in reality $n_B^s$ is a flexible upper limit on clone size, but in steady state when $n_B^s$ is approximately constant, it is true that no single clone will grow larger than $n_B^s$.

To solve equation 112, we expand about $n$ and keep terms up to 2nd order to get the Fokker-Planck equation:

$$-1 = \frac{dT}{dn}(bn + D - dn) + \frac{1}{2}\frac{d^2T}{dn^2}(bn + D + dn) \tag{113}$$

To get an approximate solution, we drop the drift term, assuming that when clones are large their net growth rate is approximately zero so $bn + D - dn \approx 0$ (this is the same as setting $\dot{n}_B^i = 0$ in equation 14). This gives the following differential equation:

$$-1 \approx \frac{1}{2}\frac{d^2T}{dn^2}(bn + D + dn) \tag{114}$$

The solution to equation 114 with the boundary conditions described above is:

$$T(n) = \frac{2}{(b+d)^2}\left[D\ln D - (D + (b+d)n)\ln(D + (b+d)n) + (b+d)n(1 + \ln(D + (b+d)n_B^s))\right] \tag{115}$$

Equation 115 with $n = {n_B^i}^*$ is plotted in Figure 81. We also solved the full Fokker-Planck equation numerically without assuming the drift term is 0 (equation 113). In this numerical solution, we change the value of $n_V^i$ for different values of $n = n_B^i$ to reflect the fact that at small bacteria clone sizes phage clones also tend to be smaller; we use the numerical solutions for average $n_v^i(t)$ and $n_B^i(t)$ shown as dashed lines in Figure 77.

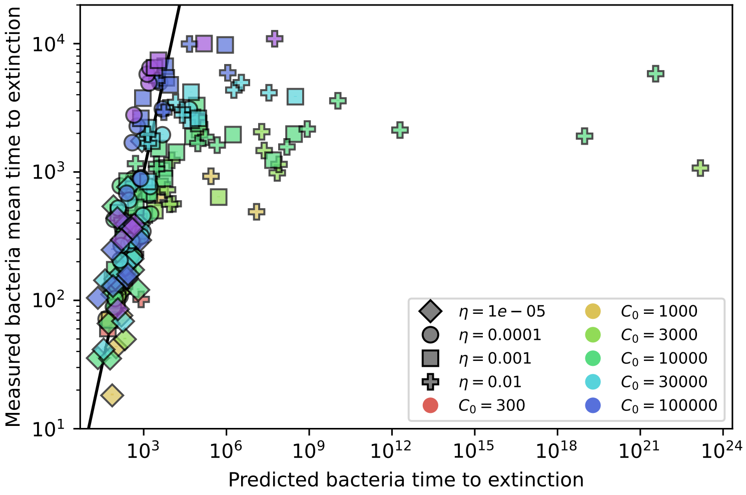

Figure 82: Measured vs. predicted mean time to extinction for bacterial clones after reaching size ${n_B^i}^*$. The predicted time to extinction is the solution with drift, given by numerically solving equation 113.

##### 5.1.2 Approximate time to extinction

Since we are interested in the time to extinction once bacterial clones reach a large size, we can substitute $n = {n_B^i}^*$ in equation 115 to gain insight into the time to extinction for bacteria. The average clone size ${n_B^i}^* = \frac{n_B^s}{m}$.

$$T({n_B^i}^*) = \frac{2}{(b+d)^2}\left[\frac{n_B^s(b+d)}{m}(1+\ln(n_B^s(b+d)+D)) - \left(\frac{n_B^s(b+d)}{m}+D\right)\ln\left(\frac{n_B^s(b+d)}{m}+D\right)+D\ln(D)\right] \tag{116}$$

We can substitute values for $b+d$ using the steady-state deterministic solution for $n_B^s$ and assuming $n_V^i = n_V/m$:

$$b+d = gC + F + r + \alpha p_V n_V(1-\frac{e}{m}) = 2F + 2r + 2\alpha p_V n_V(1-\frac{e}{m}) - \alpha(1-p_V)\eta n_V \frac{1-\nu}{\nu} \tag{117}$$

To approximate the extinction time expression, we decompose $n_B$, $n_V$, and $\nu$ into series expansions in $e/m$, i.e. $n_V \approx n_{V0} + \frac{e}{m}n_{V1} + \frac{e}{m}^2 n_{V2}$. We substitute these expressions into equation 116 and perform an overall series expansion in $e/m$. The 0th order term is shown here and plotted in Figure 83B, and the solution to 1st order is plotted in Figure 83A.

$$
\begin{gathered}
T\approx-\bigg[\bigg(2C_0^2f^2g^2\eta(\alpha(Bp_V-1)+g)^2\Big(-\frac{\alpha(f-1)(Bp_V-1)}{fg}-1\Big) \\
\bigg(-r\ln\bigg(-\frac{C_0^2fg^2\eta r(\alpha+f(\alpha(Bp_V-1)+g)+\alpha(-B)p_V)}{\alpha m(Bp_V-1)\Big(g\big(\alpha BC_0(f-1)\eta p_V^2+p_V(r-\alpha(B+1)C_0(f-1)\eta)+\alpha C_0(f-1)\eta\big)+\alpha p_V r(Bp_V-1)+C_0f\eta g^2(p_V-1)\Big)}\bigg) \\
-((g(2\alpha C_0(p_V-1)(Bp_V-1)+(3p_V-4)r)+\alpha(3p_V-4)r(Bp_V-1)) \\
\ln\bigg(\frac{C_0^2fg^2\eta(\alpha+f(\alpha(Bp_V-1)+g)+\alpha(-B)p_V)(g(2\alpha C_0(p_V-1)(Bp_V-1)+(3p_V-4)r)+\alpha(3p_V-4)r(Bp_V-1))}{\alpha m(Bp_V-1)(\alpha(Bp_V-1)+g)\Big(g\big(\alpha BC_0(f-1)\eta p_V^2+p_V(r-\alpha(B+1)C_0(f-1)\eta)+\alpha C_0(f-1)\eta\big)+\alpha p_V r(Bp_V-1)+C_0f\eta g^2(p_V-1)\Big)}\bigg) \\
-(p_V-1)(g(2\alpha C_0(Bp_V-1)+3r)+3\alpha r(Bp_V-1)) \\
\bigg(\ln\bigg(\frac{C_0^2fg^2\eta(\alpha+f(-\alpha+\alpha Bp_V+g)+\alpha(-B)p_V)(g(2\alpha C_0m(p_V-1)(Bp_V-1)+r(3m(p_V-1)-1))+\alpha r(Bp_V-1)(3m(p_V-1)-1))}{\alpha m(Bp_V-1)(\alpha(Bp_V-1)+g)\Big(g\big(\alpha BC_0(f-1)\eta p_V^2+p_V(r-\alpha(B+1)C_0(f-1)\eta)+\alpha C_0(f-1)\eta\big)+\alpha p_V r(Bp_V-1)+C_0f\eta g^2(p_V-1)\Big)}\bigg)+1\bigg)) \\
/(\alpha(Bp_V-1)+g)))\div\Big(\alpha m(Bp_V-1)(g(2\alpha C_0(Bp_V-1)+3r)+3\alpha r(Bp_V-1))^2\Big(C_0\eta(p_V-1)(\alpha+f(\alpha(Bp_V-1)+g)+\alpha(-B)p_V)+\frac{p_V r(\alpha(Bp_V-1)+g)}{g}\Big)\Big)\bigg]
\end{gathered}
\tag{118}
$$

This expression remains unwieldy, so we also do a series expansion in $r$ and keep the 0th order term (equivalent to setting $r = 0$). Spacer loss is not the dominant death rate for spacer-containing clones; the rate of spacer loss we use is an order of magnitude lower than the rate of cell death due to outflow ($R = 0.04$, $f = 0.3$). This expression becomes very simple:

$$
T({n_B^i}^*) \approx \frac{f\,(1+\ln m)\,(\alpha(Bp_V-1)+g)}{\alpha^2 m(Bp_V-1)^2} \tag{119}
$$

We can write equation 119 in terms of total population sizes without CRISPR ($\tilde{n_B}$ and $\tilde{n_V}$) to gain insight.

$$
T({n_B^i}^*) \approx \frac{\tilde{n_B}}{F+p_V\alpha\tilde{n_V}}\frac{(1+\ln m)}{m} \tag{120}
$$

If average immunity is low, then $F + \alpha p_V \tilde{n_V} \approx gC$ (from the mean-field equation for $n_B$).

$$
T({n_B^i}^*) \approx \frac{n_B}{g\tilde{C}}\frac{(1+\ln m)}{m} = \frac{n_B}{b}\frac{(1+\ln m)}{m} \tag{121}
$$

Equation 119 is compared to the measured time to extinction in Figure 83C. It is not a good approximation for low $\eta$, but at the two highest values of $\eta$ it captures the trend reasonably well.

Interestingly the dependence on $m$ in this approximate expression is identical to the dependence on $m$ in the phage extinction approximation (equation 64). Furthermore, there is also a proportionality to the mean clone size in the form $n_B/m$ in the bacteria extinction equation (since $n_B^s = n_B$ if $r = 0$, which is why this approximation works best at high $\eta$ when $\nu$ is closer to 1). This is reasonable considering that the dynamics of bacteria and phage are ultimately matched to each other at steady state.

#### 5.2 Phage clone dynamics

To solve for the dynamics of individual phage clones, we can write a one-dimensional master equation just for $n_V^j$ (equation 122). Here we neglect mutations; we assume that mutations are rare and a burst always contributes to the clonal population being tracked. This master equation is a birth-death master equation with bursts – it is in the form of a classic birth-death master equation except that the population grows with a jump of size $B - 1$. $P_n = P_{n_V^i}(t|N_0)$ is the probability of having $n$ phages of type $i$ at time $t$ given $N_0$ phages of type $i$ at $t = 0$.

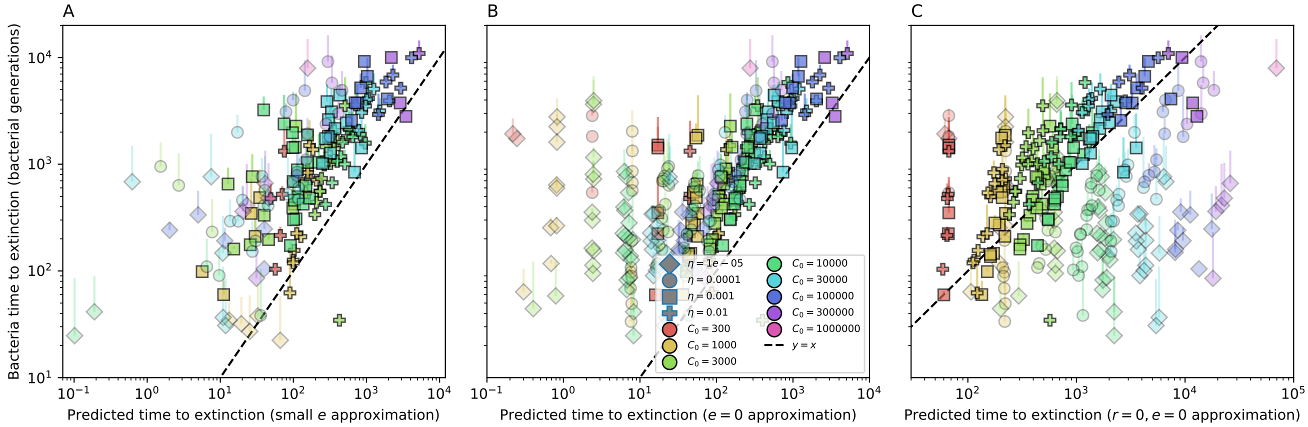

Figure 83: Measured mean time to extinction for large bacterial clones vs. three successively more aggressive analytic approximations for the mean time to extinction. (A) The predicted time to extinction is given by taking a series expansion in $e/m$ and keeping terms to 1st order. (B) A series expansion in $e/m$ and keeping the 0th order term ($e = 0$) only. (C) Series expansion in $e/m$ and $r$ and keeping the 0th order term only (equation 119).

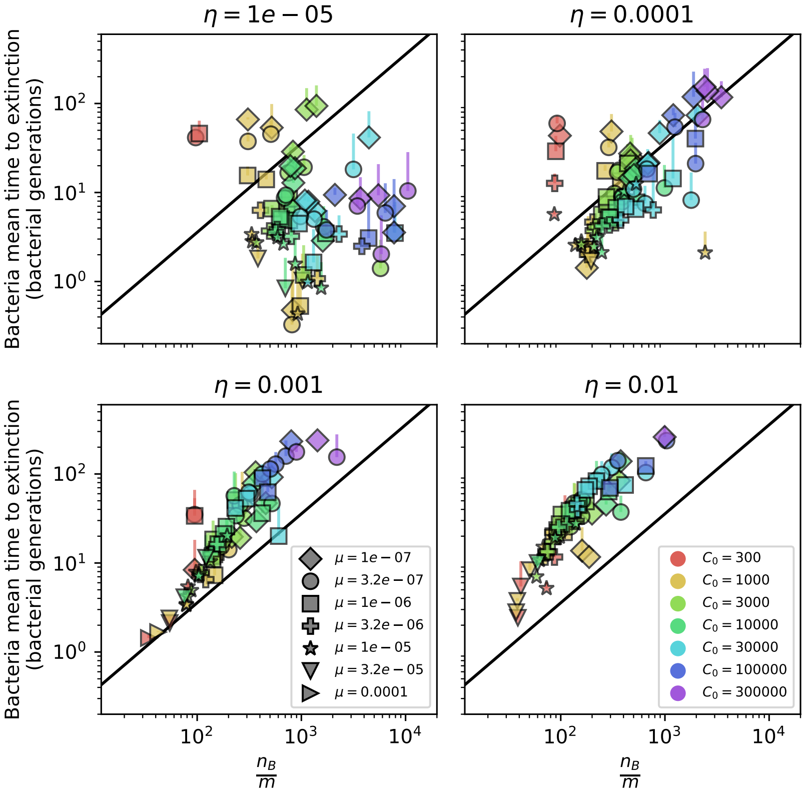

Figure 84: Bacteria extinction time as a function of mean clone size. Large bacteria clone mean time to extinction as a function of $\frac{n_B}{m}$, where $m$ is measured from simulations and $n_B$ is the total bacteria population size. This parameter combination describes the trend in both phage and bacteria extinction reasonably well at large values of $\eta$ (bottom panels). The solid line is given by equation 119 without the $1 + \ln m$ term..

$$
\begin{aligned}
\frac{dP_n}{dt} =&(n+1)P_{n+1}[F+\alpha(1-p_V)n_B+\alpha p_V e n_B^i] \\
&+(n-B+1)P_{n-B+1}[\alpha p_V n_B-\alpha p_V e n_B^i] \\
&-nP_n[\alpha n_B+F]
\end{aligned} \tag{122}
$$

#### 5.3 Phage clone probability of extinction

We can find the probability of extinction $P_0$ for new phage mutants under the assumption that $n_B$ and $n_B^i$ are constant using equation 122. We solve this equation using a generating function approach: $G(z,t)=\sum_n z^n P_n(t)$. Multiplying equation 122 by $\sum_n z^n$, we get the corresponding generating function partial differential equation:

$$
\frac{1}{\beta+\delta}\partial_t G(z,t)=\partial_z G(z,t)\left(1-p+pz^B-z\right) \tag{123}
$$

Here $p=\frac{\beta}{\beta+\delta}$ where $\beta$ and $\delta$ are the birth and death rates for phage mutants: $\beta=\alpha p_V n_B-\alpha p_V e n_B^i$ and $\delta=F+\alpha n_B(1-p_V)+\alpha p_V e n_B^i$. We distinguish two cases of constant $n_B^i$: $n_B^i=0$, valid at early times after phage mutation, and $n_B^i={n_B^i}^*$, the deterministic steady-state value of $n_B^i$ given by equation 17, valid at long times after phage mutation. When we are assuming $n_B^i=0$, we use $\beta=\beta_0$ and $\delta=\delta_0$.

We solve equation 123 using the method of characteristics:

$$
\partial_x G(z(x),t(x))=\frac{\partial G}{\partial z}\frac{\partial z}{\partial x}+\frac{\partial G}{\partial t}\frac{\partial t}{\partial x} \tag{124}
$$

This gives the following characteristic equations:

$$
\frac{\partial t}{\partial x}=\frac{1}{\beta+\delta} \tag{125}
$$

$$
\frac{\partial z}{\partial x}=-(1-p+pz^B-z) \tag{126}
$$

Integrating these two equations, we get $t=\frac{x}{\beta+\delta}$, and

$$
x=-\int_{z(0)}^{z(x)}\frac{dw}{1-p+pw^B-w} \tag{127}
$$

The initial phage clone size is $N_0$. The initial condition is

$$
G(z(0),x=0)=z(0)^{N_0}=G(z(x),t(x))
$$

The last equality is from the method of characteristics: the solution for $G$ at the initial condition gives the full solution which is constant as parameterized by $x$.

The probability of extinction is given by setting $z=0$ in $G(z,t)$.

$$
P_0(t)=G(z(t)=0,t)=z(0)^{N_0}|_{z(t)=0}=\zeta^{N_0}
$$

where $\zeta$ solves equation 128, which we obtain by setting $z(x)=0$ in equation 127 and replacing $z(0)$ with $\zeta$, since $G(z,t)|_{z=0}=z(0)^{N_0}|_{z=0}=P_0(t)$.

$$
x=\int_0^{\zeta}\frac{dw}{1-p+pw^B-w} \tag{128}
$$

##### 5.3.1 Long-time approximation for $P_0(t)$

To approximate $P_0(t)$ at long times, we notice that since extinction probability becomes large at $t = \infty$, then the large-$t$ limit corresponds to $\zeta \to 1$.

The denominator of equation 128 will be smallest when $w$ is near 1, so the largest contribution to the integral will come from $w$ near 1. Expanding the denominator to 2nd order with $w = 1 - \epsilon$, where $\epsilon$ is small:

$$1 - p + pw^B - w \approx 1 - p + p(1 - \epsilon B + \frac{1}{2}\epsilon^2 B(B-1)) - 1 + \epsilon = \epsilon - p\epsilon B + \frac{p}{2}\epsilon^2 B(B-1)$$

Changing back to the variable $w$ and substituting back into equation 128, the integral becomes:

$$x \approx \int_0^{\zeta} \frac{dw}{1 - w - pB(1-w) + \frac{p}{2}(1-w)^2 B(B-1)} \tag{129}$$

which when evaluated gives

$$\zeta = \frac{(2 - 3Bp + B^2 p)(1 - \mathrm{e}^{st})}{2 - Bp(3 - \mathrm{e}^{st}) + B^2 p(1 - \mathrm{e}^{st})} \tag{130}$$

$$\zeta = \frac{(B(\delta + s) - 2s)(\mathrm{e}^{st} - 1)}{2s + B(\mathrm{e}^{st} - 1)(\delta + s)} \tag{131}$$

where $t$ is in units of minutes and we introduce the variable $s = \beta(B-1) - \delta = \delta\left(\frac{p}{1-p}(B-1) - 1\right)$ (and $p = \frac{s+\delta}{s+B\delta}$). The parameter $s$ is the average growth rate of phage clones. When phage mutants first appear and $n_B^i = 0$, this corresponds to $s > 0$ and we write $s = s_0$. As $n_B^i \to {n_B^i}^*$, $s \to 0$.

The $s = 0$ limit of 130 is

$$\zeta = \frac{B\delta t}{2 + B\delta t} \tag{132}$$

The long-time limit of 131 is

$$\zeta = \frac{2 - 3Bp + B^2 p}{Bp(B-1)} = 1 - \frac{2s}{B(s+\delta)} \tag{133}$$

For a phage clone that begins at size $N_0 = 1$ and for which there are no matching bacterial clones ($n_B^i = 0$), equation 134 gives the probability of that clone going extinct.

$$P_0 = 1 - \frac{2s_0}{B(s_0 + \delta_0)} \tag{134}$$

If $B = 2$ (a return to a simple birth-death equation where the population grows and dies by increments of 1), equation 131 becomes

$$\zeta = \frac{\delta(\mathrm{e}^{st} - 1)}{\mathrm{e}^{st}(\delta + s) - \delta} \tag{135}$$

This is the expected probability of extinction for a simple birth-death process with selection given in [27]. (Note that time can be rescaled so that $\delta = 1$.)

The long-time limit of 135 is

$$\frac{1-p}{p} = \frac{\delta}{s+\delta} = \frac{\delta}{\beta} \tag{136}$$

This is the expected limit for the extinction probability for a simple birth-death process with selection. As $s \to 0$, the probability of extinction goes to 1.

Note that the probability of extinction for $B > 2$ (equation 133) is larger than for $B = 2$ (equation 136). Somewhat paradoxically, this implies that one effect of the burst size $B > 2$ is to increase the probability of extinction for new phage mutants. This happens because bursts are infrequent events; on average $B$ death events must occur for every one birth event, making death a more common stochastic outcome. The extinction probability for $B > 2$ is $\frac{1}{B}(\frac{2}{B-1} + \frac{\beta(B-2)}{\delta})$ times greater than for $B = 2$. Expanding in $1/B$ for large $B$, this is $\frac{\beta}{\delta}(B-3) + \mathcal{O}(1/B)$.

##### 5.3.2 Small-time approximation for $P_0(t)$

When $t$ is small, $P_0$ will be close to 0, which means $\zeta$ will be close to 0 and therefore we can approximate $w$ as being near 0.

The denominator simplifies to $1 - p - w$ for small $w$, and we get

$$x \approx \int_0^{\zeta} \frac{dw}{1-p-w} \tag{137}$$

$$\zeta = (1-p)(1-\mathrm{e}^{-x}) = (1-p)\left(1-\mathrm{e}^{-\beta_0 t/p}\right) = \frac{\delta_0}{\beta_0+\delta_0}\left(1-\mathrm{e}^{-(\beta_0+\delta_0)t}\right) \tag{138}$$

##### 5.3.3 Long-time approximation for $P_0(t)$ with no selection

In the above derivations of the probability of phage clone extinction, we generally assumed that phages had a selective advantage because there were no bacteria with matching spacers at the time a phage clone arose by mutation. If instead a phage clone does not have a selective advantage, this corresponds to $s = \beta(B-1) - \delta \approx 0$, or $p \approx 1/B$. $p$ is exactly $1/B$ where $n_B$ is given by the mean-field steady state with no CRISPR; in other words, phage clones do not have a selective advantage if the population is in steady state and there is no CRISPR immunity to distinguish clones from each other.

Expanding the denominator with $w = 1 - \epsilon$ as before:

$$1 - p + pw^B - w \approx \epsilon - p\epsilon B + \frac{p}{2}\epsilon^2 B(B-1)$$

When $p = 1/B$, the first two terms cancel out. We get the integral

$$x \approx \frac{2}{pB(B-1)} \int_0^{\zeta} \frac{dw}{(w-1)^2} \tag{139}$$

which gives

$$\zeta = \frac{(-B+B^2)px}{2-Bpx+B^2px} = \frac{(B-1)Bpx}{2+(B-1)Bpx} = \frac{(B-1)B\beta t}{2+(B-1)B\beta t} \tag{140}$$

Equation 140 is equivalent to equation 132, assuming $\beta(B-1) = \delta$. This is true when $s = 0$, since $s = 0$ corresponds to $(B-1)\beta = \delta$.

##### 5.3.4 Neutral time to extinction from backward master equation

We can calculate the time to extinction using the backward master equation corresponding to equation 122. The backwards equation is an equation for the time to extinction $T_n$ from a given state. Instead of working with frequencies, we write this in terms of the number of phages belonging to a clone, $n$. Here $\beta = \alpha p_V n_B - \alpha p_V e n_B^i$ and $\delta = F + \alpha n_B(1-p_V) + \alpha p_V e n_B^i$.

$$T_n - \Delta t = \beta n \Delta t T_{n+B-1} + \delta n \Delta t T_{n-1} + (1 - \beta n \Delta t - \delta n \Delta t) T_n \tag{141}$$

Rearranging equation 141, we arrive at equation 142.

$$-1 = \beta n T_{n+B-1} + \delta n T_{n-1} - (\beta n + \delta n) T_n \tag{142}$$

For boundary conditions, we have $T(n = 0) = 0$ (time to extinction is 0 when already extinct) and $\frac{dT}{dn}|_{n=n_V} = 0$ (reflecting boundary at $n = n_V$). As with bacteria clones, we assume here that the upper limit on phage clone size is constant at $n_V$ at steady state.

To solve equation 142, we expand about $n$ and keep terms up to 2nd order to get the Fokker-Planck equation:

$$-1 = \frac{dT}{dn} \left(\beta n(B-1) - \delta n\right) + \frac{1}{2} \frac{d^2T}{dn^2} \left(\beta n(B-1)^2 + \delta n\right) \tag{143}$$

At steady state and for large clones, $\beta(B - 1) \approx \delta$, which means clones are approximately neutral and we can drop the drift term:

$$-1 \approx \frac{1}{2} \frac{d^2T}{dn^2} \left(\beta n(B-1)^2 + \delta n\right) \tag{144}$$

This is a straightforward second-order ordinary differential equation, which has the solution

$$T(n) = c_1 + n c_2 - \frac{2n\left(-1 + \ln[n(\beta(B-1)^2 + \delta)]\right)}{\beta(B-1)^2 + \delta} \tag{145}$$

Using the boundary condition $T(0) = 0$, the constant $c_1 = 0$. Using the second boundary condition, $c_2 = \frac{2\ln[n_V(\beta(B-1)^2+\delta)]}{\beta(B-1)^2+\delta}$, and the full solution is

$$T(n) = \frac{2n}{\beta(B-1)^2 + \delta} \left(1 - \ln \frac{n}{n_V}\right) \tag{146}$$

Once phage clones become large (by reaching size ${n_V^i}^*$ as given by equation 18), they behave neutrally and equation 146 agrees well with observed extinction times in simulations. We set $n = {n_V^i}^*$ and use the deterministic steady-state clone size for $n_B^i$ given by equation 17 in $\beta$ and $\delta$. Figure 86 compares observed large clone extinction times in a simulation (blue) to the prediction given by equation 146 (orange dashed).

If ${n_V^i}^* \approx \frac{n_V}{m}$, which it is at steady-state, we can rewrite equation 146 as:

$$T(n) = \frac{2n}{\beta(B-1)^2 + \delta} (1 + \ln m) \tag{147}$$

The factor $1 + \ln m$ comes from the integral of the second-order differential equation and from the reflecting boundary assumption, and interestingly it would happen even if $B = 2$ (no burst): integrating the Fokker-Planck equation gives a factor of $1-\ln n$; if we start from the steady-state value of $n \approx n_V/m$ this becomes $1 + \ln m - \ln n_V$. Applying the reflecting boundary at $n = n_V$ cancels out the $\ln n_V$ term. Very simply: the integral we're solving is $\frac{-1}{n} \propto \frac{d^2T}{dn^2}$, and this gives $n(1 - \ln n)$ up to some constants.

#### 5.4 A neutral model of non-coevolving bacteria

In our results for the diversity in a coevolving population of bacteria and phage, we find that diversity approximately scales like $m \approx e\mu\eta^{\frac{1}{3}}$ (Section 4.2.5). Intuitively this dependence seems quite weak, but we'd like to know what to compare this to: what is the expectation for diversity in a population with mutations but without coevolution?

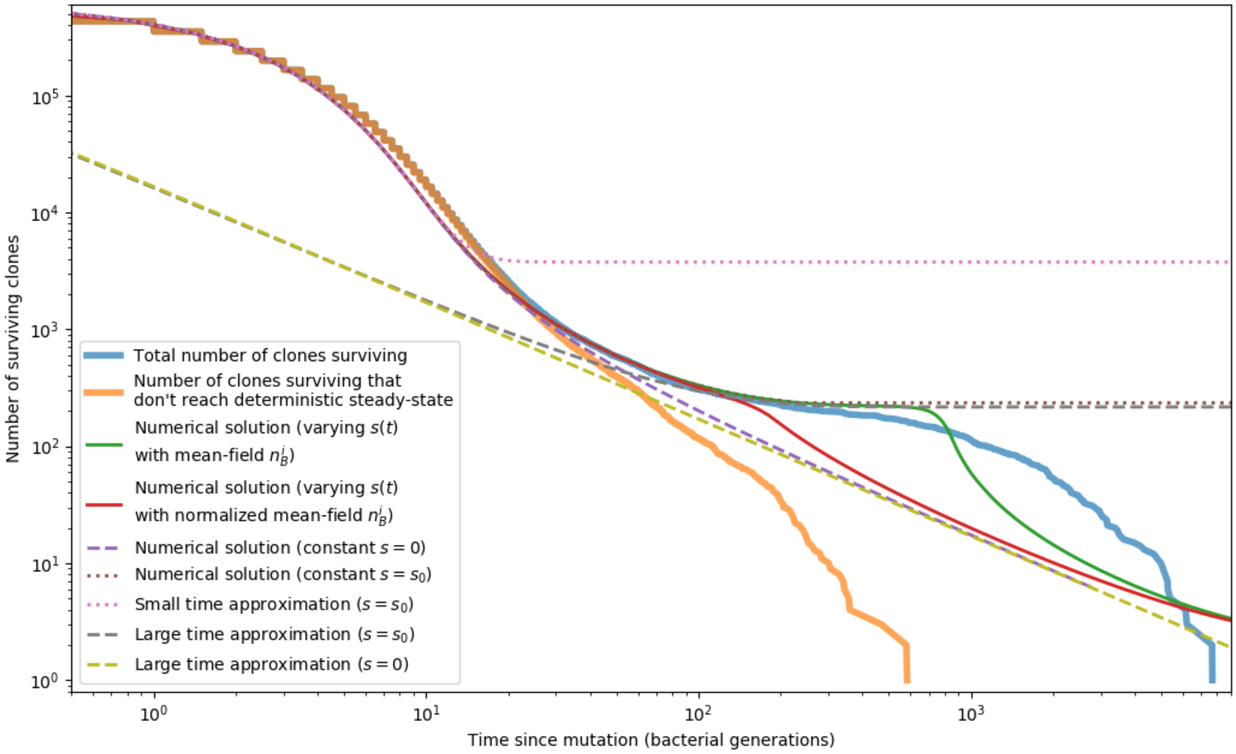

Figure 85: Phage clone extinction times and theoretical predictions for a simulation with parameters $C_0 = 10^4, \eta = 0.001, e = 0.95, \mu = 10^{-5}$. Time zero for each trajectory is the time at which that clone arose by mutation. All simulation trajectories are plotted in blue, and a subset of trajectories that don't reach a size of ${n_V^i}^* = 16170$ as given by equation 18 are plotted in orange. Trajectories that don't become established by this definition go extinct more quickly than ones that do. All other curves are theoretical predictions for the extinction time. The green and red solid lines and purple and brown dashed lines show a numerical solution to equation 128 with different values for $s$. The remaining dashed lines are a small time approximation given by equation 138, and a large time approximation given by equation 131 with either $s = s_0$ or $s = 0$. All of these predictions agree with the simulation data in different regimes; none accurately captures the entire timecourse of extinctions.

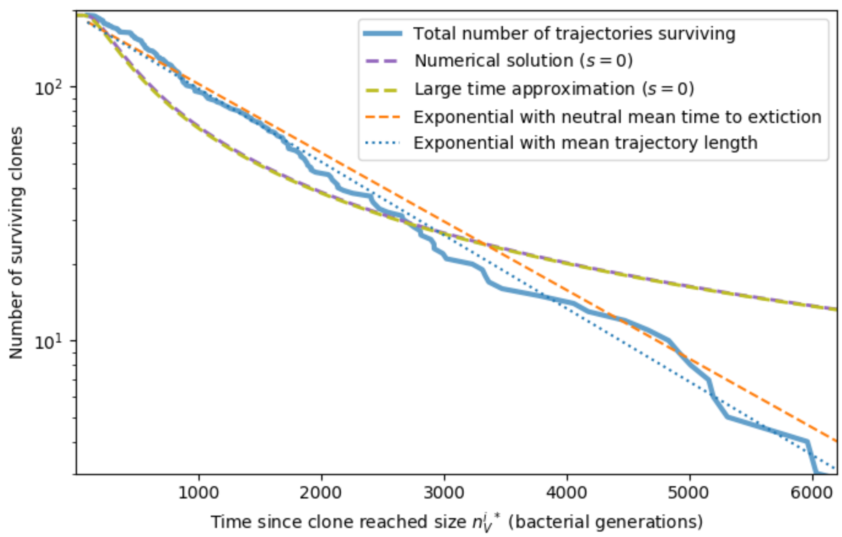

Figure 86: Large clone extinction times from a simulation with parameters $C_0 = 10^4, \eta = 0.001, e = 0.95, \mu = 10^{-5}$. Trajectories are counted as large if the phage clone size passes $n_V^{i\,*} = 16170$, the theoretical deterministic mean phage clone size for these parameters given by equation 18. Once a trajectory reaches $n_V^{i\,*}$, we count that point as time zero to measure the extinction time of large clones. The numerical solution is given by solving equation 128 with $s = 0$ ($p = 1/B$). The large time approximation is given by equation 132. The orange dashed line gives an exponential decay prediction with the mean time to extinction given by equation 146 with $n = n_V^{i\,*}$.

An appropriate comparison point is a neutral model of a bacterial population that grows and dies; the birth and death rates are exactly matched to give a constant size at steady state. We also include the possibility of mutations: a bacterium may mutate to a new type with probability $\mu$ per division, and we assume that each mutation is to a new, never-before-seen type (infinite alleles model).

Let's consider a population of bacteria that divides with rate $g$, dies with rate $F$, and mutates with rate $\mu$. We can write a master equation for the number of bacterial clones of size $k$. New mutants enter at clone size 1, and we assume that the total population size is constant at $N_b = \sum_k kb_k$. Mutations effectively lower the growth rate of a particular clone.

$$\partial_t b_k = (g - \mu)[(k-1)b_{k-1} - kb_k] + F[(k+1)b_{k+1} - kb_k] + \delta_{k,1}\mu N_b \tag{148}$$

#### Generating function solution

The generating function for the probability distribution $b_k(t)$ is $G(z,t) = \sum_k z^k b_k(t)$. Let's replace the birth and death rates by $\beta$ and $\delta$, respectively. $\beta = g - \mu$, $\delta = F$. Multiplying equation 148 with $\sum_k z^k$ and noting that $\partial_z G(z,t) = \sum_k kz^{k-1} b_k(t)$, we get the following differential equation:

$$\partial_t G(z,t) = \partial_z G(z,t)\left(z^2\beta - z(\beta+\delta) + \delta\right) + Dz \tag{149}$$

The term $Dz$ comes from the source term with $D = \mu N_b$.

Equation 149 can be solved with the method of characteristics [35]. We parameterize the function $G(z,t)$ with a new variable $s$. Applying the chain rule:

$$\partial_s G(z(s), t(s)) = \frac{\partial G}{\partial z}\frac{\partial z}{\partial s} + \frac{\partial G}{\partial t}\frac{\partial t}{\partial s} \tag{150}$$

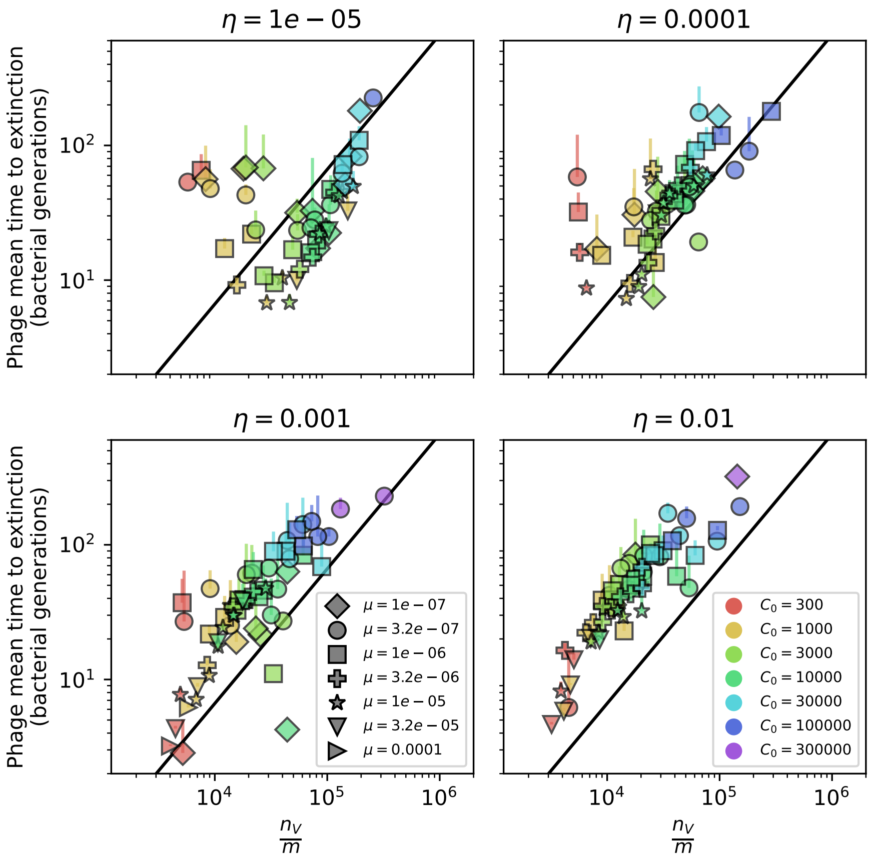

Figure 87: Phage extinction time as a function of mean clone size. Large phage clone mean time to extinction as a function of $\frac{n_V}{m}$, where $m$ is measured from simulations and $n_V$ is the total phage population size. The solid line is given by equation 147 without the $1 + \ln m$ term.

And by comparison with equation 149, the characteristic equations are

$$\frac{\partial t}{\partial s} = 1 \tag{151}$$

$$\frac{\partial z}{\partial s} = (1-z)(\beta z - \delta) \tag{152}$$

$$\frac{\partial G}{\partial s} = Dz \tag{153}$$

From equation 151 we see $t = s + C$, so we can choose $t_0 = C = 0$ and replace $s$ with $t$ going forward. Solving the characteristic equation for $z$ by integrating both sides gives equation 154.

$$\frac{1-z}{\delta - \beta z} \mathrm{e}^{(\beta-\delta)t} = C \tag{154}$$

At $t = 0$, $z$ will pass through some point $z_0$, so we have the initial condition $z(0) = z_0$. With $z_0$ in equation 154 at $t = 0$, we get equation 155, where $C$ is given by equation 154.

$$z_0 = \frac{C\delta - 1}{C\beta - 1} \tag{155}$$

The variation of $G$ along this $z$-$t$ curve is given by

$$\partial_z G = \frac{-Dz}{z^2\beta - z(\beta+\delta) + \delta} \tag{156}$$

which has the solution

$$G(z) = \frac{D}{\beta - \delta}\left[-\ln(1-z) + \frac{\delta}{\beta}\ln(\delta - \beta z)\right] + \Omega(C) \tag{157}$$

The constant $\Omega$ is a function of the characteristic $z$-$t$ curve (equation 154). To find the particular form of $\Omega(C)$, we apply the initial condition $G(z, 0) = z^{N_b}$, meaning we start with 1 clone of size $N_b$ at $t = 0$.

$$z^{N_b} = \frac{D}{\beta - \delta}\left[-\ln(1-z) + \frac{\delta}{\beta}\ln(\delta - \beta z)\right] + \Omega\left[\frac{1-z}{\delta - \beta z}\right] \tag{158}$$

Let's use the temporary variable $\xi = \frac{1-z}{\delta - \beta z}$. Then $z = \frac{\xi\delta - 1}{\xi\beta - 1}$, and the solution for $\Omega(\xi)$ is

$$\Omega(\xi) = \frac{\xi\delta - 1}{\xi\beta - 1}^{N_b} - \frac{D}{\beta - \delta}\left[-\ln(1 - \frac{\xi\delta - 1}{\xi\beta - 1}) + \frac{\delta}{\beta}\ln(\delta - \beta\frac{\xi\delta - 1}{\xi\beta - 1})\right] \tag{159}$$

Now we can write the full solution for $G(z, t)$, replacing $\xi$ with $\xi\epsilon$, where $\epsilon = \mathrm{e}^{(\beta-\delta)t}$.

$$\begin{aligned} G(z,t) \;&=\; \left(\frac{\epsilon\delta(1-z) - \delta + \beta z}{\epsilon\beta(1-z) - \delta + \beta z}\right)^{N_b} + \frac{D}{\beta-\delta}\left[-\ln(1-z) + \frac{\delta}{\beta}\ln(\delta - \beta z)\right] \\ &\quad - \frac{D}{\beta - \delta}\left[-\ln(1 - \frac{\epsilon\delta(1-z) - \delta + \beta z}{\epsilon\beta(1-z) - \delta + \beta z}) + \frac{\delta}{\beta}\ln(\delta - \beta\frac{\epsilon\delta(1-z) - \delta + \beta z}{\epsilon\beta(1-z) - \delta + \beta z})\right] \end{aligned} \tag{160}$$

$b_k$ can be found by Taylor-expanding $G(z)$ about $z = 0$: $b_k(t) = \frac{1}{k!}\frac{\partial^k G}{\partial z^k}|_{z=0}$. The full distribution is cumbersome for this initial condition, but the limit as $t \to \infty$, if $\beta < \delta$, is independent of the initial condition:

$$b(k) = \frac{D}{k}\frac{\beta^{k-1}}{\delta^k} = \frac{D}{\beta k}e^{-k\ln(\delta/\beta)} \tag{161}$$

Now we have required constant $N_b = \sum_k kb_k$, which means

$$N_b = \sum_k D\frac{\beta^{k-1}}{\delta^k} = \frac{D}{\delta-\beta} = \frac{\mu N_b}{F-g+\mu} \tag{162}$$

So to maintain constant population size, $F = g$, as expected.

The diversity is the total number of clones, defined as $\sum_k b_k$. We represent diversity as $m$.

This gives the following result for steady-state diversity:

$$\sum_k b_k = m = -\frac{D}{\beta}\ln\left(\frac{\delta-\beta}{\delta}\right) = -\frac{\mu N_b}{g-\mu}\ln\left(\frac{\mu}{g}\right) \tag{163}$$

In our definition of the birth and death rates, it is implied that $\mu < g$, and under that condition, the diversity is positive. If we let $\frac{\mu}{g} = u$, we can write the diversity as

$$m = -N_b\frac{u}{1-u}\ln u \tag{164}$$

For small $u$ (i.e. mutation rate much smaller than division rate, usually true), $m \approx -N_b u \ln u$.

Equation 161 is equivalent to Fisher's logseries with $\alpha = \frac{D}{\beta}$ and $x = \frac{\beta}{\delta}$. This is also equivalent to Hubbell's unified neutral theory in the limit of large sample size. This result for diversity (equation 164) is also approximately equivalent to the result obtained in [36] and also in [37] when the mutation rate is within a few orders of magnitude of $g$.

##### 5.4.1 Comparison to coevolution model

In our bacteria-phage coevolution model, we find that diversity scales approximately like $m \sim a^{\frac{1}{3}}$, where

$$a \approx \frac{4e\mu L\frac{\hat{\eta}\alpha\tilde{n_V}}{\hat{\eta}\alpha\tilde{n_V}+r}\tilde{n_V}^2}{(B-1)^2} \tag{165}$$

For $\mu L << 1$, $Bp_V - 1 >> 1$ and $p_V r > \hat{\eta}gC_0(1-f)$, the dependence of diversity on $a$ is given by equation 166. Note that this expression correctly captures the fold-change in diversity as a function of parameters, but the value itself is off by about a factor of 10.

$$a \approx \frac{4e\mu L\hat{\eta}(gC_0(1-f))^3}{(Bp_V-1)^2\alpha^2 p_V r} \tag{166}$$

Is this dependence quite different from what we get under neutral evolution without coevolution? Since we are measuring bacterial diversity in the coevolution model, the spacer acquisition probability $\eta$ is a good analog to bacterial mutation rate in the non-coevolving model. Let's look at the change in diversity for a given change in spacer acquisition probability.

In the coevolution model, a $k-$fold change in spacer acquisition probability gives approximately a $k^{1/3}$ change in diversity, while under the non-coevolution model, a $k-$fold change in mutation rate $u$ gives a $\frac{k(1-u)}{1-ku}(1+\frac{\ln k}{\ln u})-$fold change in diversity. For small $u$, this is approximately $k(1+\frac{\ln k}{\ln u})$. The change is dependent on the mutation rate, but if $k$ is not very large and $u << 1$, then a $k-$fold change in $u$ gives approximately a $k-$fold change in diversity. A similar correspondence can be reached with a different approximation. If $a$ is large, the leading order contribution to $m$ is $a^{1/3}$, but a better approximation is given by equation 167.

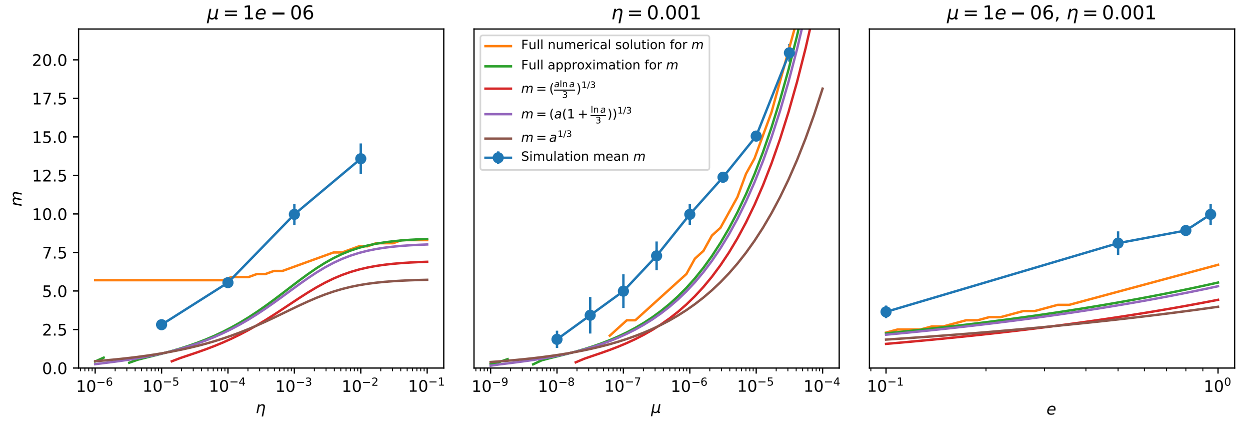

Figure 88: Diversity vs. $\eta$ (left), $\mu$ (centre) and $e$ (right) for $C_0 = 10^4$. The $\eta$ dependence of diversity is not very well predicted even by the full numerical solution, but for mutation rate and spacer effectiveness the approximate solutions do pretty well in this regime. The simulation data increase in diversity as a function of spacer acquisition probability actually goes more like $m \propto \ln \eta$.

$$m \approx \left( a(1 + \frac{\ln a}{3}) \right)^{\frac{1}{3}} \tag{167}$$

If $a$ is large, which it typically is, then the leading term is more accurately $\frac{a \ln a}{3}^{1/3}$. This is directly comparable to the $u \ln u$ dependence of the non-coevolving model: diversity in the coevolution model goes like $u \ln u^{1/3}$, and in the non-coevolving model it goes like $u \ln u$. Either way, there's a 1/3 power in the coevolution model that isn't there in the simple model.

Figure 89 compares the predicted diversity and fold-change in diversity in each model under the simple approximation assumptions here. Diversity increases more rapidly with mutation rate in the simple model than in the coevolution model.

### 6 Data analysis

#### 6.1 Theoretical considerations when calculating average immunity from data

In our simulations and model, we assume that each phage has a single protospacer and each bacterium has a single spacer. In reality, phages can have hundreds to thousands of possible protospacers, and bacteria can also acquire tens to hundreds of spacers. If we assume that average immunity plays out at the organism level; that is, if a bacterium that contains one or more spacers matching one or more phage protospacers is immune to that phage, then calculating average immunity in practice requires knowing something about the typical numbers of protospacers and spacers in organisms.

First, let's consider the effect of changing the number of protospacers while keeping spacer array length constant at 1. This is a reasonable model for experimental data at short timescales when most bacteria acquire only one new spacer. For a set $i$ of observed spacers and a set $j$ of observed protospacers with abundances $n_i$ and $n_j$, if each spacer and protospacer are assumed to belong to one organism only, then the average immunity is given by equation 168. We assume for simplicity that any matching spacer provides perfect immunity, i.e. $p_V(i,j) = p_V(1 - \delta_{ij})$.

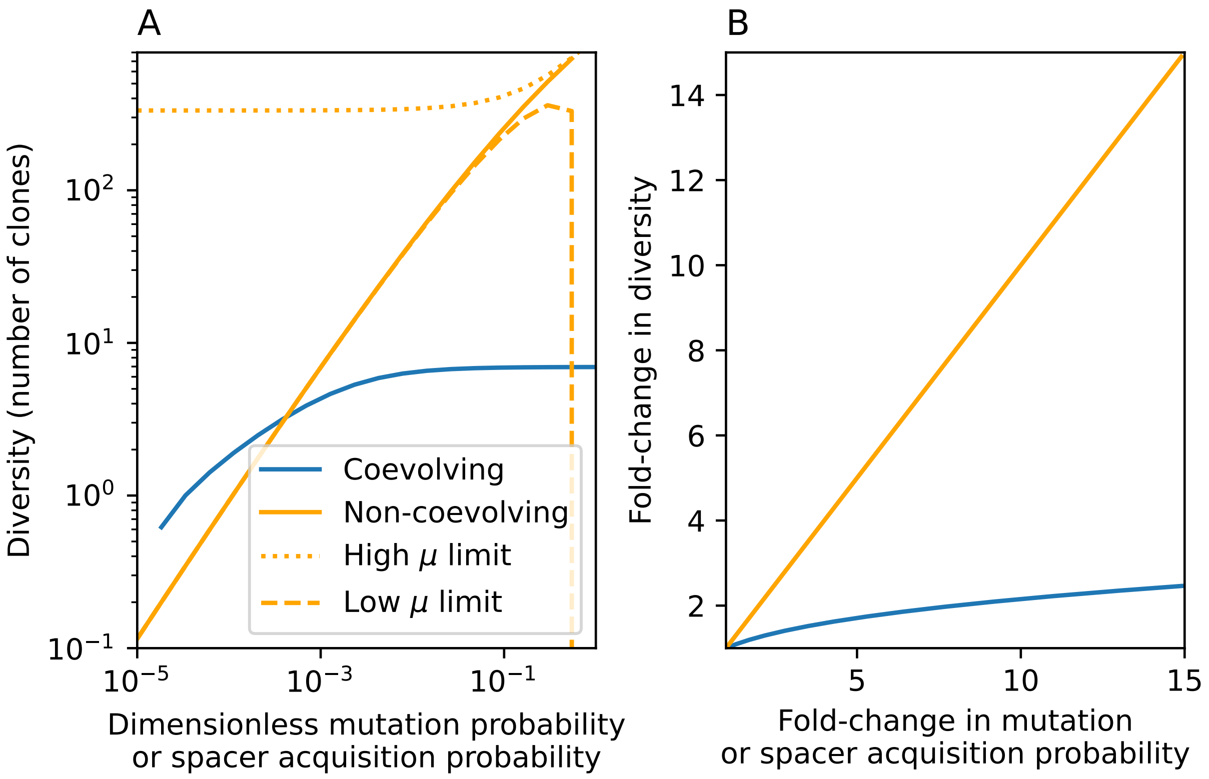

Figure 89: Approximate predictions for diversity in our coevolution model (blue) or a simple model with mutation but no coevolution (orange). (A) Predicted diversity as a function of spacer acquisition probability in our coevolution model as given by equation 165 for $C_0 = 10^4$, $e = 0.95$, and $\mu = 10^{-6}$ (blue). Predicted diversity in the non-coevolving model as given by $m = -N_b \frac{u}{1-u} \ln u$ with $N_b = 1000$ (orange). The low-$\mu$ limit is $m = -N_b u \ln u$. The high-$\mu$ limit is a series expansion in $\epsilon$ for $\mu = 1 - \epsilon$ giving $m \approx N_b(\frac{1}{3} + \frac{5\mu}{6} - \frac{\mu^2}{6})$. (B) Fold-change in diversity as a function of fold-change in mutation rate or spacer acquisition probability under the simple approximation that $m \sim \eta^{1/3}$ (blue) and $m \sim u$ (orange).

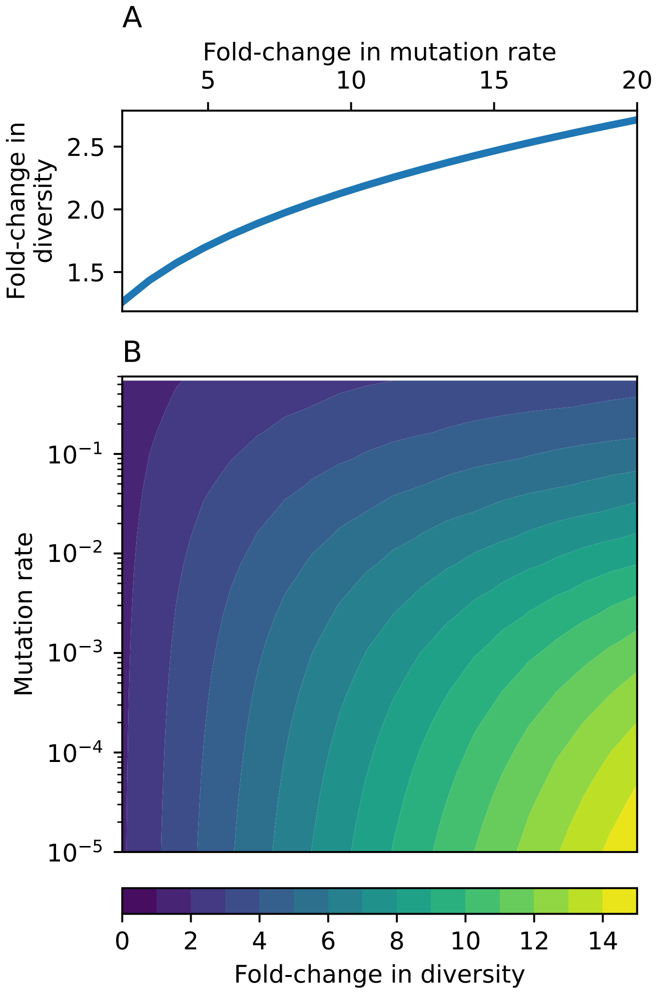

Figure 90: Fold-change in diversity (number of species) as a function of mutation rate and $k$, the fold-change in mutation rate. (A) Fold-change in diversity as a function of fold-change in mutation rate in the coevolution model: diversity increases by approximately a factor of $k^{\frac{1}{3}}$, independent of mutation rate (solid line). (B) Fold-change in diversity as a function of both mutation rate and fold-change in mutation rate in the model without coevolution.

$$1 - \frac{\sum_{i,j} n_i n_j p_V(i,j)}{p_V \sum_{i,j} n_i n_j} = 1 - \frac{p_V \sum_{i,j} n_i n_j (1-\delta_{ij})}{p_V \sum_{i,j} n_i n_j} = \frac{\sum_{i,j} n_i n_j \delta_{ij}}{N_i N_j} \tag{168}$$

where $N_i$ denotes $\sum_i n_i$.

If instead the same set of protospacers is divided among fewer phages so that each phage has $a$ protospacers on average, then the total number of phages is $N_j/a$. The numerator of average immunity remains the same since each bacterium still contains only a single spacer. Average immunity is exactly the same as before but multiplied by the average number of protospacers per phage (equation 169).

$$\frac{\sum_{i,j} n_i n_j \delta_{ij}}{N_i \frac{N_j}{a}} = \frac{a \sum_{i,j} n_i n_j \delta_{ij}}{N_i N_j} \tag{169}$$

This agrees with the intuition that more spacer targets per phage increases the immune potential of the CRISPR system.

In data from [38], we find quantitative agreement between the range of average immunity values in their data and the types of values in our simulations once we apply this simple transformation with $a = 696$, since there are 231 possible CRISPR1 protospacers and 465 possible CRISPR3 protospacers in the phage genome (determined by searching for the canonical PAM for each CRISPR locus).

If bacteria contain multiple spacers and phages contain multiple protospacers, the combinatorics of average immunity gets more interesting. Now, if a bacterium contains multiple spacers that target the same phage, we assume this does not increase its immunity (although in reality there may be a benefit to multiple matching spacers).

We simulated sets of spacers and protospacers, randomly divided them into arrays of different sizes, and calculated the organism-level average immunity in each case. Figure 91 shows the average immunity values for several array lengths with different assumptions for how the set of spacer sequences and array lengths are distributed. We chose a set of spacer 'sequences' where each sequence is a letter of the alphabet, then constructed phage protospacer arrays by randomly drawing 5 unique letters 50 times to create 50 phages. We then sampled letters from the alphabet either uniformly or from an exponential distribution (where 'A' is the number 1, etc.). The exponential sampling captures the fact that in practice spacer abundance distributions are highly non-uniform. To create bacteria spacer arrays, we sampled from the set of spacers without replacement, either creating arrays of constant size or arrays with the same mean size but with their length either normally or exponentially distributed.

We developed a simple theoretical model for the change in average immunity as array length increases. The baseline average immunity when bacteria are assumed to have single spacers we denote $a_1$, and define a constant $C = 1 - a_1$. Then as array length increases, the total number of bacteria decreases and the overlap increases, but by a smaller factor than the population size decrease: the remaining average immunity that could be gained is reduced with the same fraction as the original average immunity so that the average immunity for an array of length $n+1$ is $a_{n+1} = 1 - (1 - a_n)C$. Plotting $1 - a_n$ vs $n$ on a log plot gives a straight line. Plugging in $a_1 = 1 - C$ gives $a_n = 1 - C^n$, plotted as a black dashed line in Figure 91. The theory agrees well with the simulated results except for when array length is exponentially distributed; in this case we believe the presence of several very long and very short arrays means that short-array bacteria don't get the coverage of multiple spacers while the long-array bacteria have redundant spacers. This functional form for average immunity as a function of array length is very similar to a quantity derived by Iranzo *et al.* [24] — they calculated that the probability that a random bacterium is immune to a random phage, each with a random set of spacers and protospacers is $p_c = 1 - (1 - \alpha \frac{n}{N_t})^{N_s}$, where $n$ is the average bacterial array length, $N_t$ is the total number of unique protospacers, and $N_s$ is the number of protospacers per phage. The constant $\alpha$ is a scaling factor that measures the degree of correlation between matching spacer and protospacer

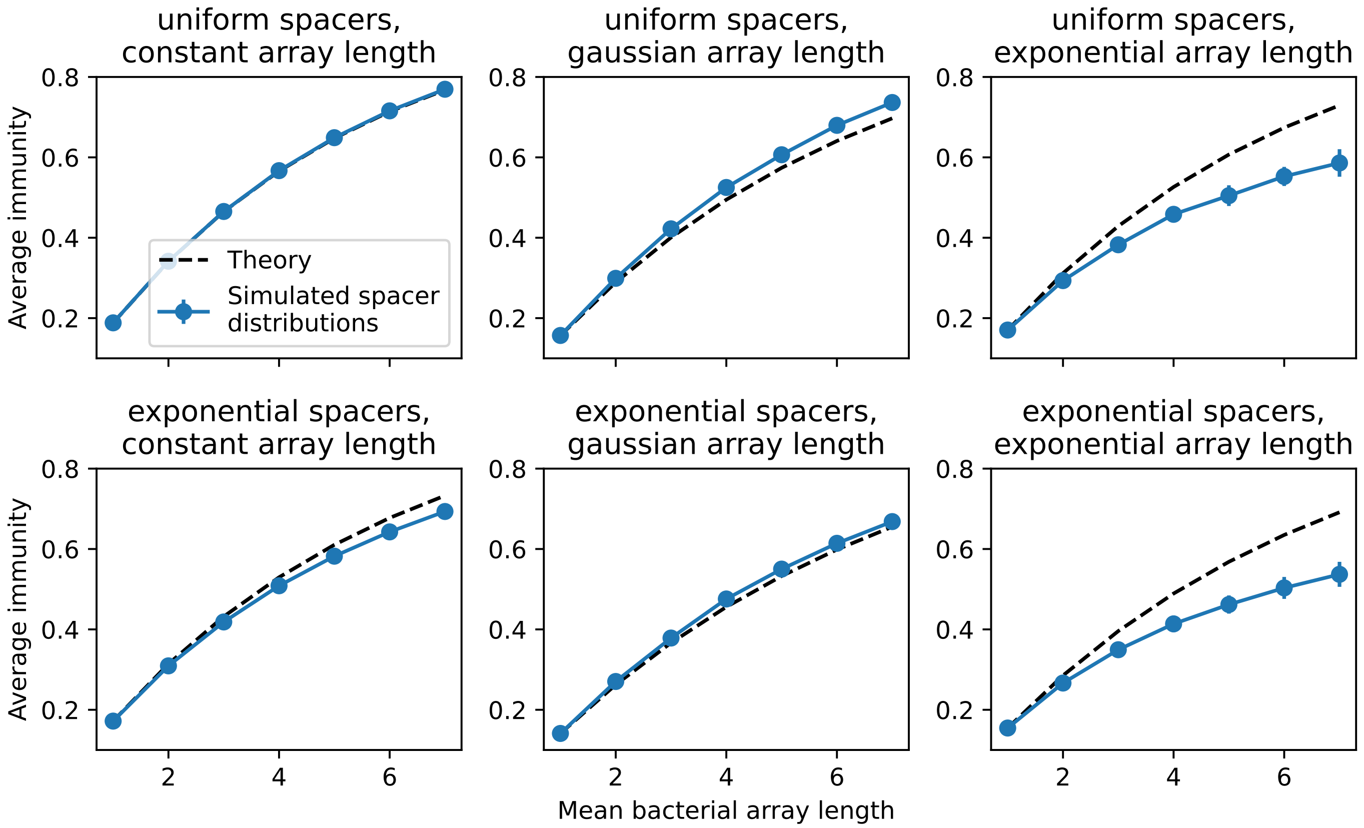

Figure 91: Average immunity vs. bacterial spacer array length for simulated distributions of protospacers and spacers. We simulate 50 phages, each with 5 protospacers represented by letters from the alphabet, uniformly sampled. We simulate 420 bacterial spacers, drawn from the alphabet either uniformly (top row) or following an exponential distribution with mean 6 (bottom row). We construct arrays by sampling from the set of 420 spacers without replacement, either creating arrays of constant length (left column), or of variable length with a gaussian distribution (middle column) or exponential distribution (right column). Average immunity is calculated as in equation 168 except the indices run over all arrays and not over individual sequences, and the presence of any matching pair gives perfect immunity. Blue points are average results over 50 simulations; error bars are standard deviation. The black dashed curve is given by $a_n = 1 - C^n = 1 - (1 - a_1)^n$.

abundances. Their scaling is as a function of the *phage* array length instead of bacterial, but the same intuition applies.

An interesting result of this analysis is that the relative immune benefit from gaining a spacer decreases as immunity increases: a CRISPR array twice as long does not provide double the immunity. A similar diminishing-returns result has also been observed in theoretical models of vertebrate immunity where the relative decrease in immune susceptibility and increase in immune memory both decrease as the number of infections increases over an organism's lifetime [39]. Long CRISPR arrays are also subject to a dilution effect: spacers compete to form complexes with limited *Cas* protein machinery, and if the number of spacers is high, there is a higher chance that the needed spacer is not available in high enough numbers during an infection [40, 41].

##### 6.1.1 Average immunity negatively correlates with diversity regardless of array size

In our simulations we observed an inverse proportionality between average immunity and bacterial diversity (Figure 58). We wondered if this trend would hold for array sizes and numbers of protospacers larger than 1. Our intuition is that this trend should hold for any arrangement of spacers and protospacers into arrays, provided the following two statements are true: 1) bacteria and phage diversity is coupled; that is, phage protospacer diversity matches bacterial spacer diversity, and 2) diversity is larger than CRISPR array length. We tested this by generating exponentially distributed random sets of 1000 spacer sequences, varying the total diversity indirectly by changing the size of the pool of available sequences and generating phage and bacterial arrays by sampling from this pool of sequences. We sampled uniformly without replacement to create 100 phages with $n$ unique protospacers each, and we sampled without replacement to create bacteria with either a Gaussian-distributed array size about a mean size $a$ (Figure 92) or a constant array size $a$ (Figure 93). We calculated average immunity and diversity (the number of unique bacterial clones) and repeated the array assortment process 50 times for each total spacer diversity and combination of array sizes.

We find that across all combinations of array sizes, average immunity is negatively correlated with diversity. Both conditions generally hold in this toy model: phage and bacterial diversity are coupled, and diversity is lower-bounded by the number of protospacers. We can conclude that for a relatively fixed CRISPR array size and number of protospacers per phage, we should expect that average immunity does not increase with diversity if phage and bacterial diversity are correlated.

The top-left panel of Figure 93) represents the same situation as our simulations: one spacer and one protospacer (but with a random distribution of sequences), and the trend matches our simulations exactly with average immunity inversely proportional to diversity. This strict inverse dependence changes shape as the number of spacers and protospacers per organism changes.

#### 6.2 Laboratory co-evolution experimental data

We analyzed experimental data from Paez-Espino *et al.* [38]. In this experiment, *S. thermophilus* bacteria were mixed with phage 2972 and allowed to co-evolve until phage extinction, up to 232 days in one replicate. Bacteria and phage whole-genome shotgun sequencing was performed at irregular intervals (13 time points for series MOI-2B). This data is publicly available in the NCBI Sequence Read Archive under the accession PRJNA275232.

We analyzed data from the MOI-2B series and detected spacers in the CRISPR1 and CRISPR3 loci by searching raw reads for matches to the *S. thermophilus* repeats (`GTTTTTGTACTCTCAAGATTTAAGTAACTGTACAAC` for CRISPR1 and `GTTTTAGAGCTGTGTTGTTTCGAATGGTTCCAAAAC` for CRISPR3) using BLAST.

The expected structure of the CRISPR locus is 36 nt repeats interspaced with 30 nt spacers. Reads are 100 nt long (Illumina), and so at most two complete spacers can be detected per read (one full repeat match near the centre of the read). To maximize the number of genuine spacers detected while removing low-quality matches, we kept only full-length alignments to the repeat (length 36), unless a shorter alignment was also present on the same read as a full-length alignment (for instance if the repeat is partially present at the start or end of a read). If a single repeat match was present on a read, we extracted 30 nt on either side and labelled these spacers. We set a minimum spacer length to be 26 nt ($\approx 0.85 \times 30$) and did not keep spacer sequences at the start or end of a read that were shorter than this. If two repeat matches were present, we extracted the spacer sequence between the matches.

To detect wild-type spacers, we searched for matches to the CRISPR repeats on the *S. thermophilus* DGCC7710 reference genome (accession NZ_CP025216) and extracted the sequences between repeat matches. We also searched raw reads from the day 1 data of the control replicate (no phages) for matches to the repeats and performed the same spacer extraction procedure.

We grouped all extracted spacers using the AgglomerativeClustering method from scipy with an

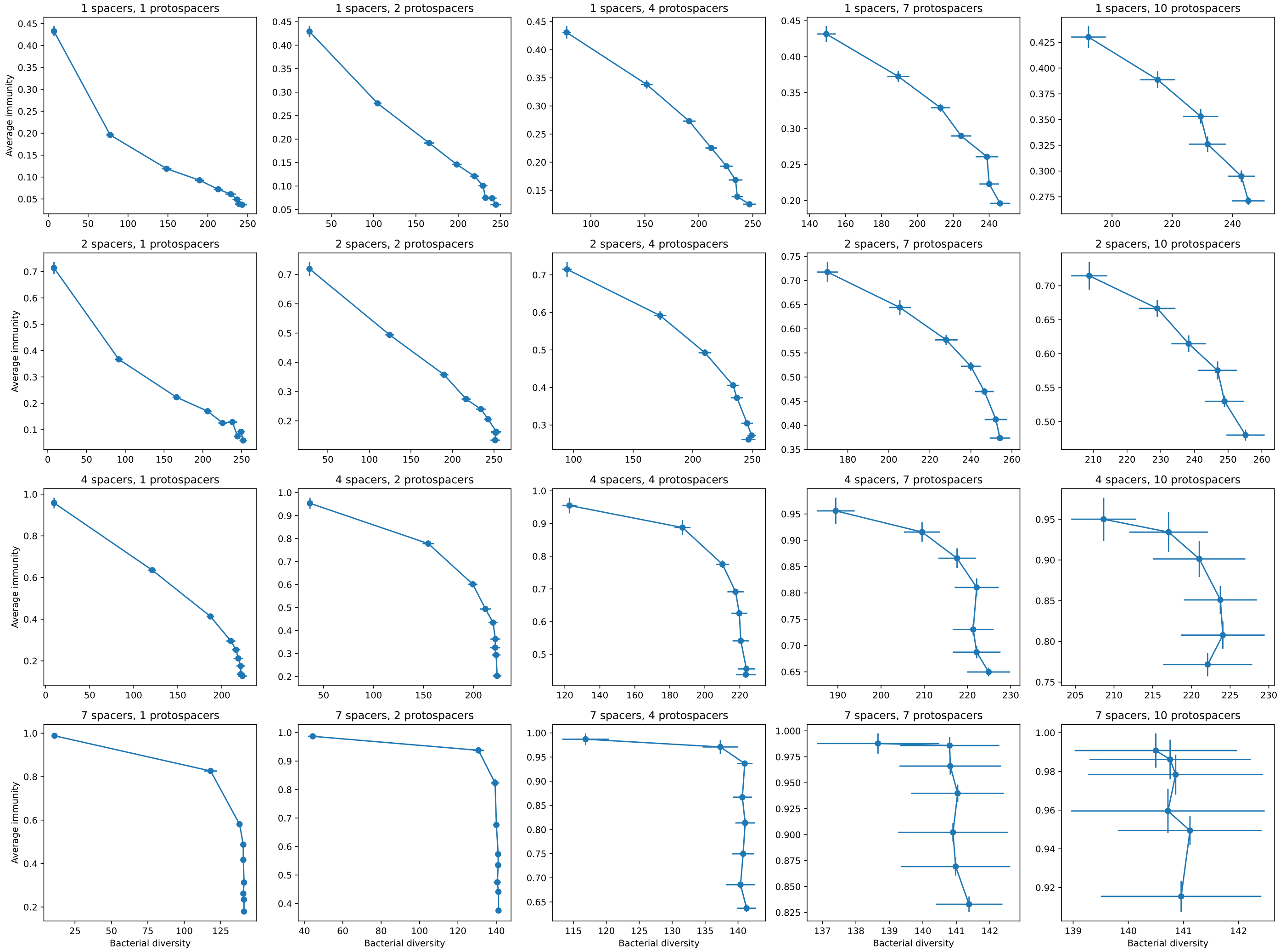

Figure 92: Average immunity vs. bacterial diversity for simulated distributions of spacers and protospacers with different bacterial array sizes (increasing top to bottom) and different numbers of protospacers per phage (increasing left to right). Bacterial array sizes are drawn from a Gaussian distribution with mean given by the array mean for each row and a standard deviation of 2. Points are averages across 50 independent runs.

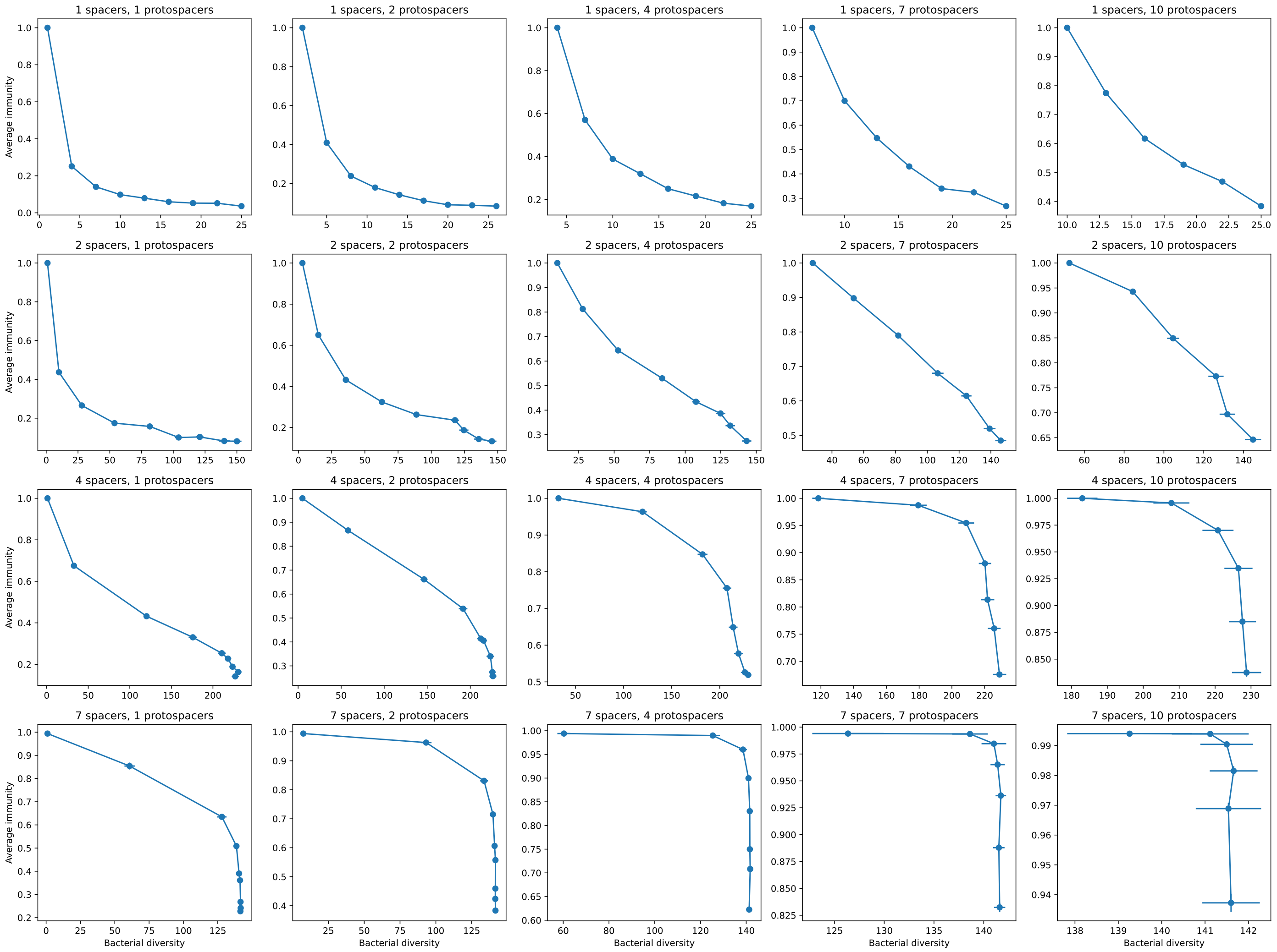

Figure 93: Average immunity vs. bacterial diversity for simulated distributions of spacers and protospacers with different bacterial array sizes (increasing top to bottom) and different numbers of protospacers per phage (increasing left to right). Bacterial array size is a constant at each row. Points are averages across 50 independent runs.

85% similarity threshold and the 'average' linkage criterion. We performed this grouping separately for CRISPR1 and CRISPR3 spacers. Each spacer sequence was then labelled with a group type identifier. Figure 94 shows our detected spacer counts after grouping by 85% average similarity and eliminating single sounts and matches to wild-type spacers, compared with the reported results from [38]; there is good agreement between our counts and the original authors' counts.

To detect protospacers, we first blasted all reads against the *S. thermophilus* DGCC7710 reference genome and the phage 2972 reference genome (accession NC_007019). We next blasted all unique detected spacer sequences from the previous step against all the reads, removing any query sequences that are a perfect subset of another sequence and any sequences that were more than 30% N nucleotide. This detects both potential protospacer sequences and the original spacers themselves. To isolate protospacers, we kept only results that were on reads that did not match the bacterial genome and that did not match the CRISPR1 repeat (CRISPR3 repeats were not checked, however they represent a very small number of additional reads, less than 0.1%). We kept only results that were 26 nt or larger. If there were multiple hits on a read from the same spacer type (but different sequences), we kept only the match with the lowest e-value. We extracted the matched sequence from the read and ten nucleotides after it to check for a PAM sequence (all spacers were stored oriented in the same direction relative to the repeat to facilitate comparison and PAM detection). Since reads are paired-end, some reads will overlap, and some spacers may be double-counted in the overlap. We decremented the total spacer or protospacer count for a sequence by 1 if a spacer sequence was present on both ends of a paired read. We also removed sequences that contained long strings of 11 or more of the same nucleotide, assuming these to be sequencing errors. This removed around 1% of sequences.

We grouped all spacer and protospacer sequences together (separated by CRISPR locus) by several different similarity thresholds between 85% and 99% to assign type labels based on each similarity threshold.

We checked the 10 nt region downstream from each potential protospacer and removed any sequences that did not have a perfect match to the PAM: AGAAW for CRISPR1 [42, 38, 43, 44] and GGNG for CRISPR3 [38, 43]. Since targeting is highly sensitive to PAM mutations [43], we assumed that any deviation from the perfect PAM meant that the protospacer would not be successfully targeted. There are cases where the PAM sequence is incomplete because of hitting the start or end of a read; these were also assumed to be imperfect PAMs (though some would be genuine). Changing this assumption to include partial PAMs that were subsets of perfect PAMs caused total average immunity numbers to increase and the slope of each curve to decrease. Results including incomplete PAMs were qualitatively similar to the results including all protospacer matches regardless of PAM, highlighting the importance of a perfect PAM match for functional immunity (Figures 101 and 102).

**Calculating average immunity**

To calculate average immunity between bacteria and phage, we interpolated spacer and protospacer counts between sequenced time points using the shortest interval between experimental time points as the sampling frequency from the interpolated data. We removed the first time point and last two time points from the data because the first time point has low bacterial spacer counts and a very large phage population size, and at the last two time points, phage counts are very low because phages are about to go extinct (Figures 98 and 95). Results including all time points are in Figure 103); they are qualitatively similar except for large changes at the first and last time points where only the first and last phage population are included in the average respectively.

We calculated the time-shifted average overlap between bacteria and phage spacer types by comparing bacteria and phage types separated by a time delay and averaging over all points with the same time delay (Figure 103). As described in section 6.1, we multiplied raw average immunity values by the total number of protospacers to account for multiple protospacers on each phage genome. The number

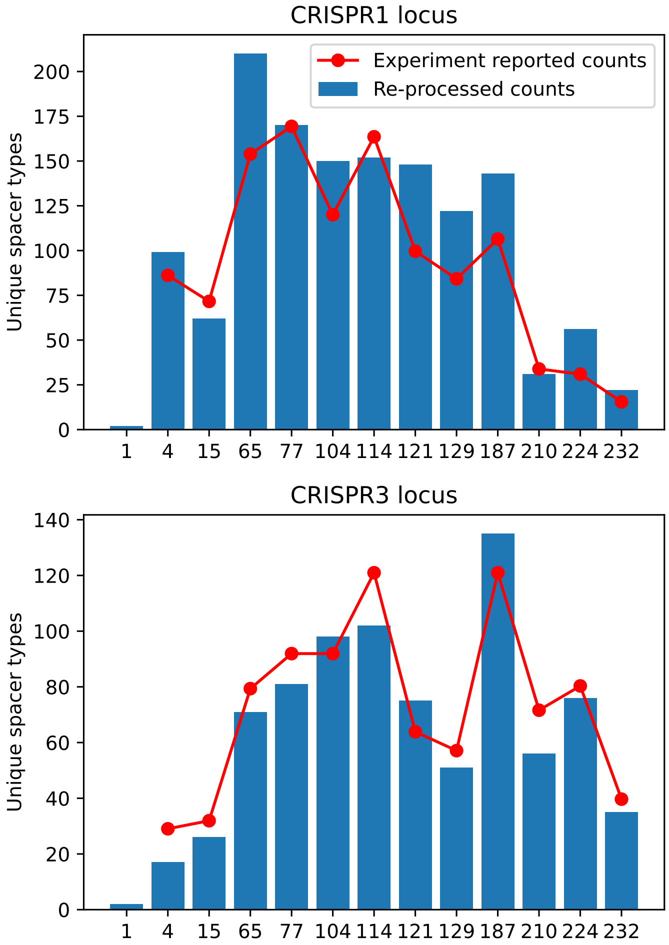

Figure 94: Unique spacer types detected in our analysis of data from [38] after grouping by 85% average similarity and removing single spacer counts (blue bars). Counts reported in [38] are red points.

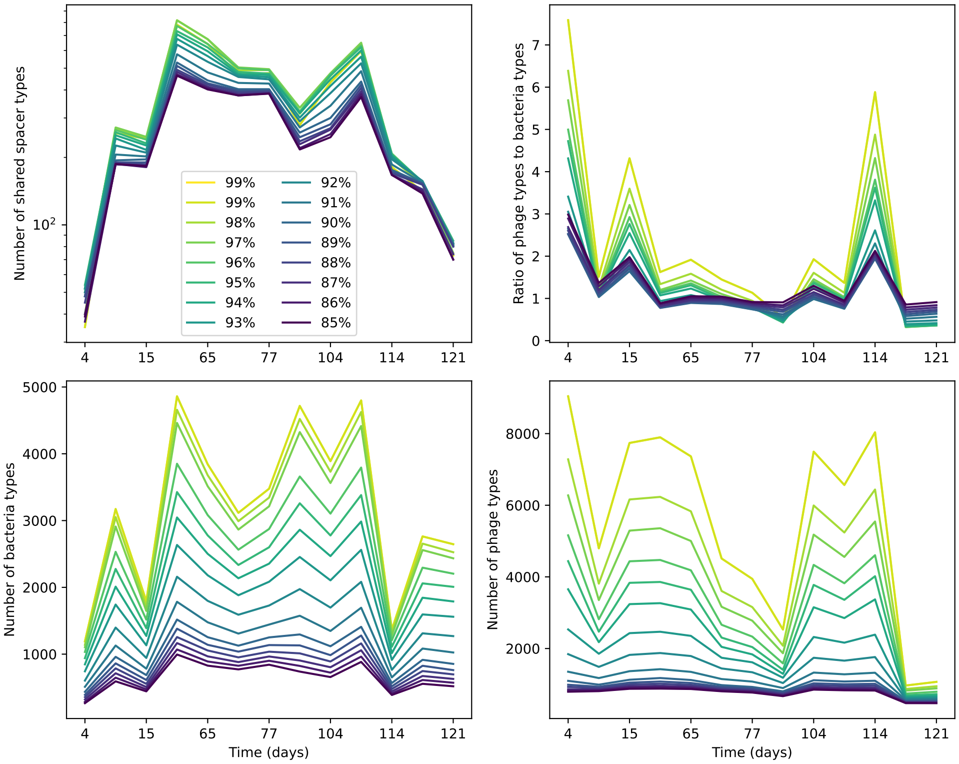

Figure 95: Number of shared spacer types between bacteria and phage (top left), ratio of the number of phage types to the number of bacteria types (top right), number of bacteria types (bottom left), and number of phage types (bottom right) as a function of the sampling date for data we analyzed from [38]. Colours indicate different similarity grouping thresholds. Spacer counts include wild-type spacers, and protospacers are included only if they possess a perfect PAM. Data is summed over CRISPR1 and CRISPR3. All sequences are included regardless of abundance.

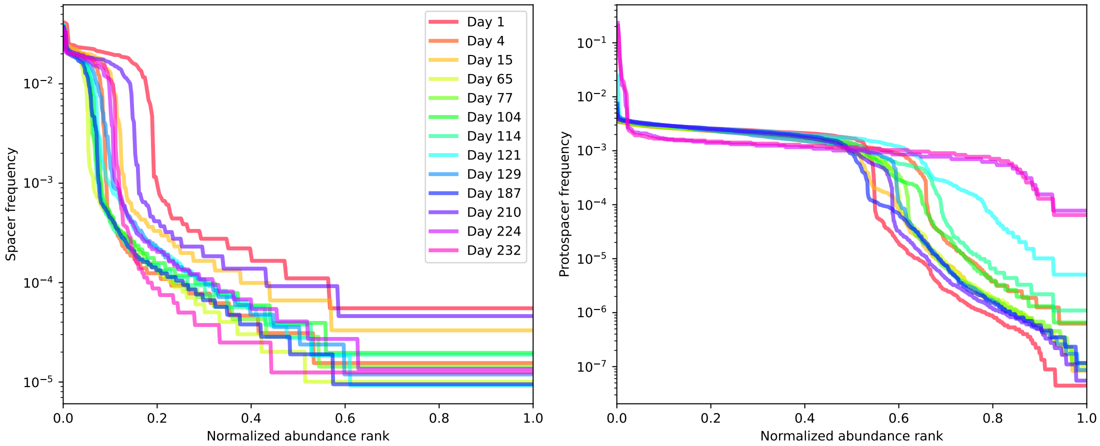

Figure 96: Rank abundance distribution for detected spacers (left) and protospacers (right) at each time point in data from [38]. Spacers are generally present at a range of abundances; protospacer abundances are more tightly peaked (a flatter rank-abundance curve) especially at high abundances. Both sets of distributions appear to have two regimes of abundances: a high-abundance region with several similar-abundance sequences, and a low-abundance regime with an exponential tail.

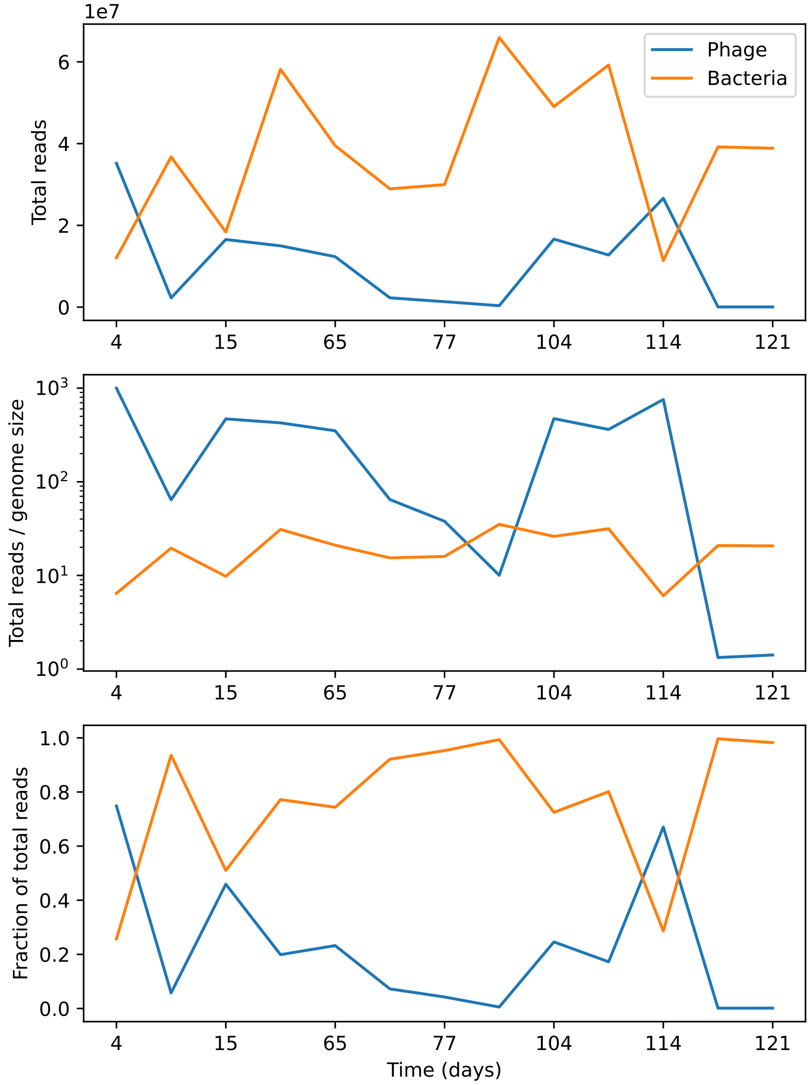

Figure 97: Total reads per time point that match phage (blue) or bacteria (orange). Top: total reads. Middle: total reads divided by phage and bacteria genome sizes. Bottom: fraction of total reads matching bacteria or phages.

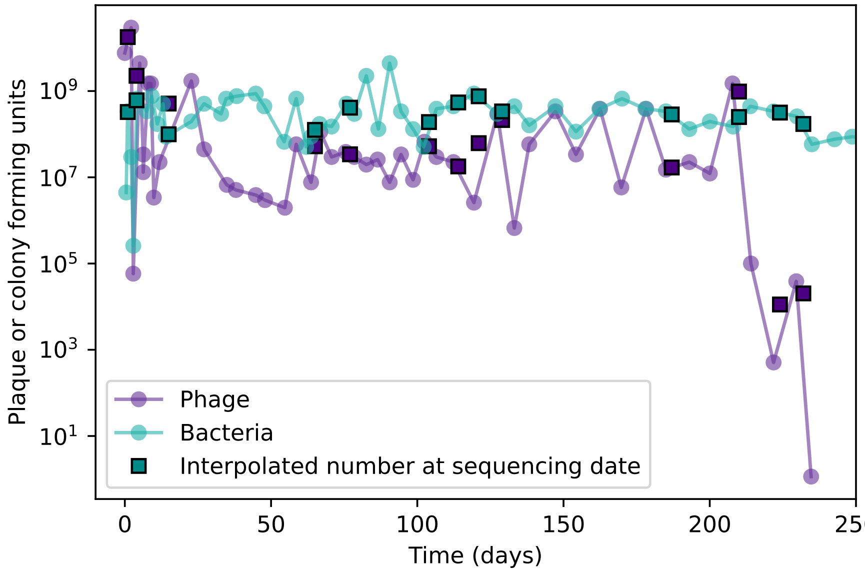

Figure 98: Total phage and bacteria population size for the MOI2B experiment in [38]. Circles are points digitized from Figure 1A in ref. [38]; squares are the population size interpolated to match the sequencing dates in the experiment.

of possible protospacers in phage 2972 for the CRISPR1 locus in *S. thermophilus* (AGAAW PAM) is 231 and the number for CRISPR3 is 465 (696 total). Note that because PAM mutations are common, the true number of protospacers may be less than this hypothetical amount, which would cause us to slightly overestimate average immunity. However, we are also assuming that each bacterium has one effective spacer, and this is likely an underestimate based on estimates of locus length from [38].

As the similarity threshold increases, the overall overlap goes down slightly (Figures 103 and following) because there are more total types (Figure 95). Interestingly, the number of shared types across the whole dataset is very stable regardless of the clustering threshold: there is a tradeoff between the total number of types (which increases as the similarity threshold increases) and the likelihood that types are shared (which increases as the similarity threshold decreases). At high similarity thresholds, we expect that many spurious types will be created from sequencing errors in spacers, while at low similarity thresholds, genuine escape mutants will be grouped with spacers meaning that the immunity information contained in the overlap is not accurate. There are about 700 unique protospacer sequences in the wild-type phage genome, so we expect the number of unique types to be on the order of 700; they are for bacteria, but the number of phage types in our analysis can be quite a bit higher. For all similarity thresholds, we see that bacterial immunity is higher to phages from the past than phages from the future.

We experimented with different trimming thresholds for the data, removing time points from the beginning and end of the data. The early and late parts of the experiment experienced large fluctuations in population sizes, so if we are interested in steady-state behaviour, it makes sense to remove some time from the beginning and end. But, how much to remove? We can remove the points that look more different from the others either in terms of population size, total reads, or number of types: this could lead us to remove the first three and last three time points. Looking just at population size alone, it makes sense to remove the first time point and last two time points. Figure 103 shows results with different sets of points removed to compare the resulting overlap.

There were no protospacer matches to the reference genome wild-type spacers for CR1 and only

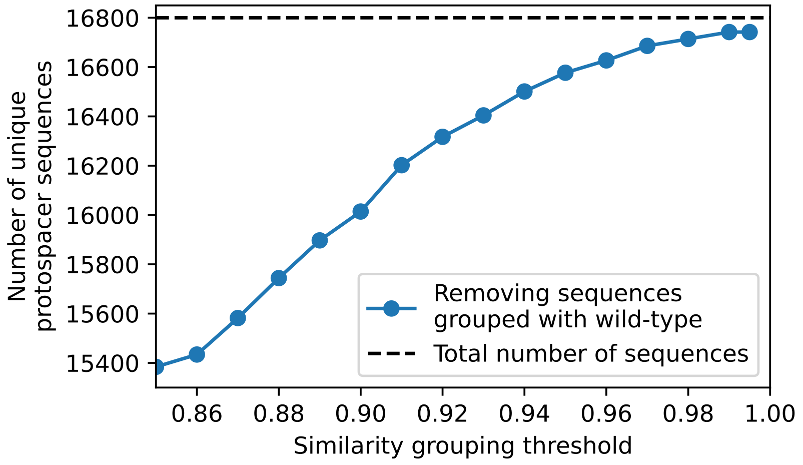

Figure 99: Total number of unique protospacer sequences after removing all sequences that cluster with wild-type sequences at different similarity thresholds. Only protospacers with a perfect PAM are included.

a single protospacer match to CR3, even with a lenient 85% similarity threshold. This means the phage has effectively escaped all the wild-type spacers and only the new spacers matter. However, our results differ when wild-type spacers are removed, since we also included spacer sequences from the control experiment as wild-type spacers. There are many sequence variants in the control experiment wild-type spacers that do cluster with other spacers, and there are protospacer sequences that cluster with supposed wild-type spacers at lenient grouping thresholds (Figure 99). This means that average immunity results with and without wild-type spacers are more similar at high similarity thresholds than at low thresholds.

Our model predicts that phage population sizes should decrease as average immunity increases and that bacterial population sizes should increase as average immunity increases. We calculated average immunity at each experimental time point and compared it to the reported phage and bacterial population size at each time point. Phage and bacteria population sizes were digitized from Figure 1A in [38]. We found that for all similarity groupings, the log-transformed phage population size was significantly negatively correlated with average immunity, while there was no significant correlation with bacterial population size (Figure 104). This suggests that the mechanism of phage extinction is increasing bacterial immunity over time, and indeed the average immunity just before phage extinction is very high (Figure 105).

#### 6.3 Experimental data from wastewater treatment plant

We analyzed metagenomic sequencing data from Guerrero *et al.* [45]. This is a longitudinal study of a natural community of bacteria and phages in a wastewater treatment plant. 60 samples were collected approximately biweekly over a 3-year period from a municipal sewage treatment plant, and whole-genome sequencing was performed on extracted DNA (Illumina platform, 250 bp paired-end reads). They focused on bacteria from the genus *Gordonia*, which is one of the most abundant taxa and was consistently present in samples. They detected CRISPR loci in *Gordonia* genome assemblies, then searched for matches to the two identified CRISPR repeats in all reads to detect other spacers. They constructed graphs of all the linked spacers based on spacer adjacency within reads. They searched for matches to spacers in the metagenomic data, then used matches to identify contigs that might be phages, resulting in two draft phage genomes (DC-56 and DS-92) that they followed over time. They

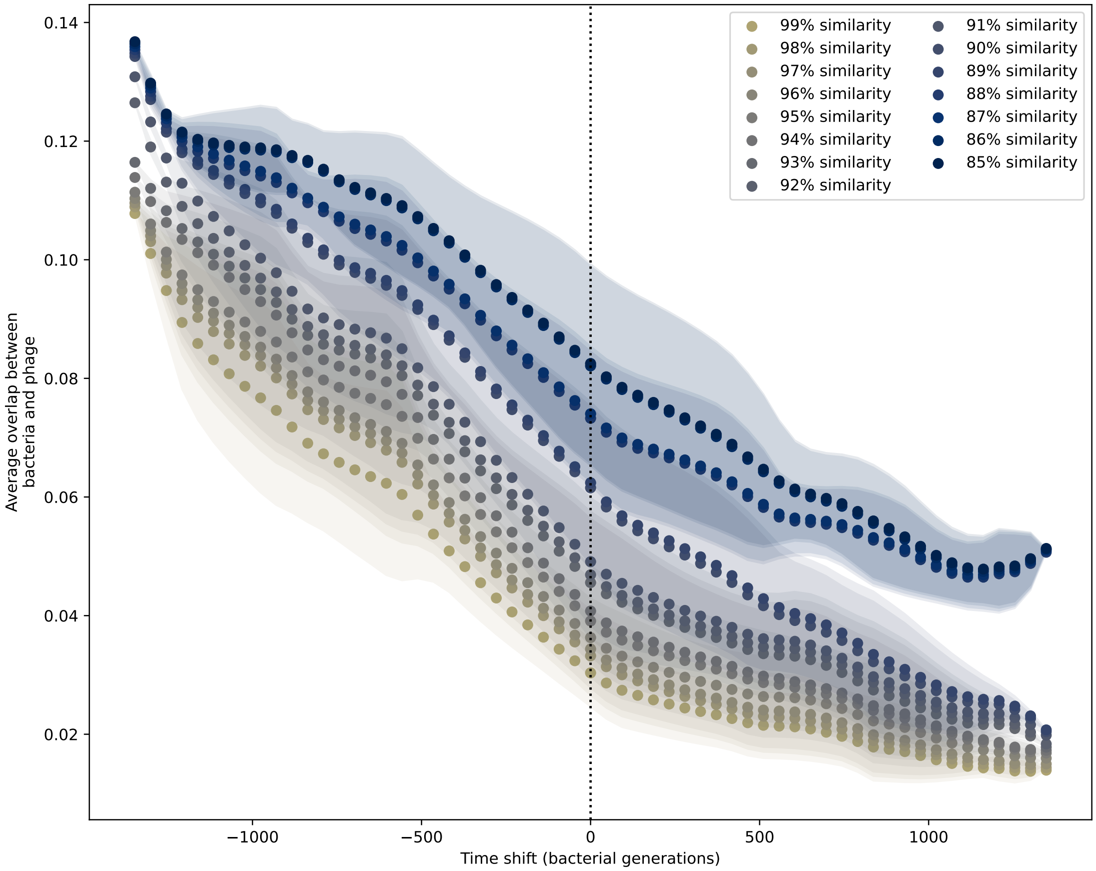

Figure 100: Time shifted overlap with the first time point and last two time points removed. Interpolation spacing is three days, the smallest interval between remaining time points. Time intervals in days were multiplied by 6.64, the estimated number of bacterial generations per day assuming exponential growth with 100:1 serial dilutions. Protospacers are included if they have a perfect PAM, and all wild-type spacers are included.

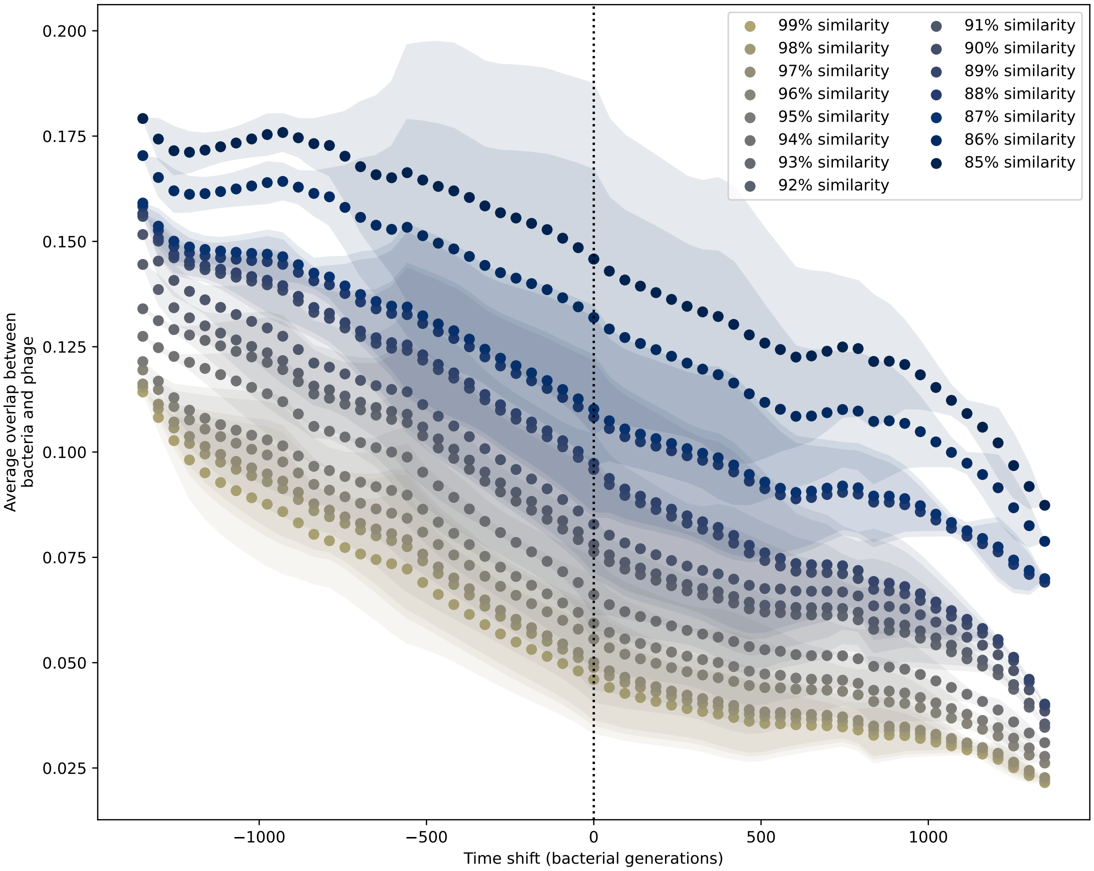

Figure 101: Time shifted overlap with the first time point and last two time points removed. Interpolation spacing is three days, the smallest interval between remaining time points. Time intervals in days were multiplied by 6.64, the estimated number of bacterial generations per day assuming exponential growth with 100:1 serial dilutions. Protospacers are included if they have a perfect PAM or partially-perfect PAM, and all wild-type spacers are included.

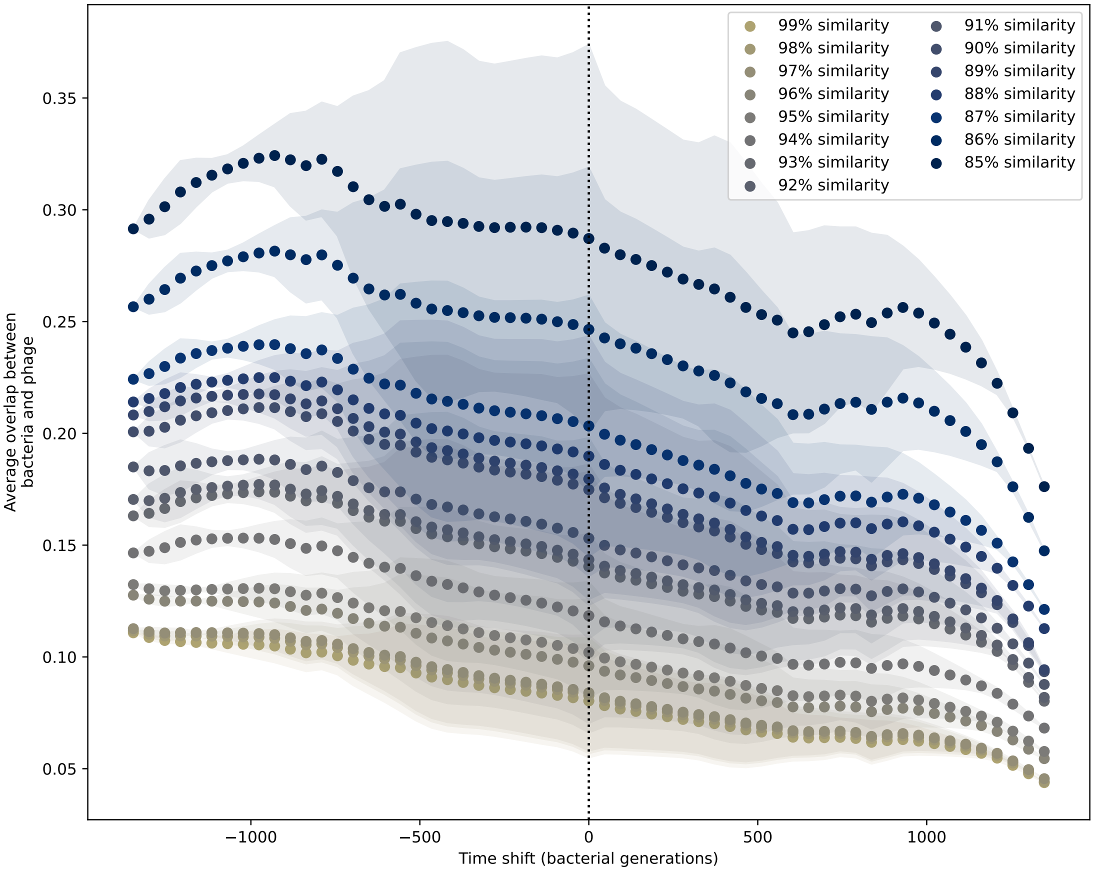

Figure 102: Time shifted overlap with the first time point and last two time points removed. Interpolation spacing is three days, the smallest interval between remaining time points. Time intervals in days were multiplied by 6.64, the estimated number of bacterial generations per day assuming exponential growth with 100:1 serial dilutions. All potential protospacers are included regardless of PAM sequence, and all wild-type spacers are included.

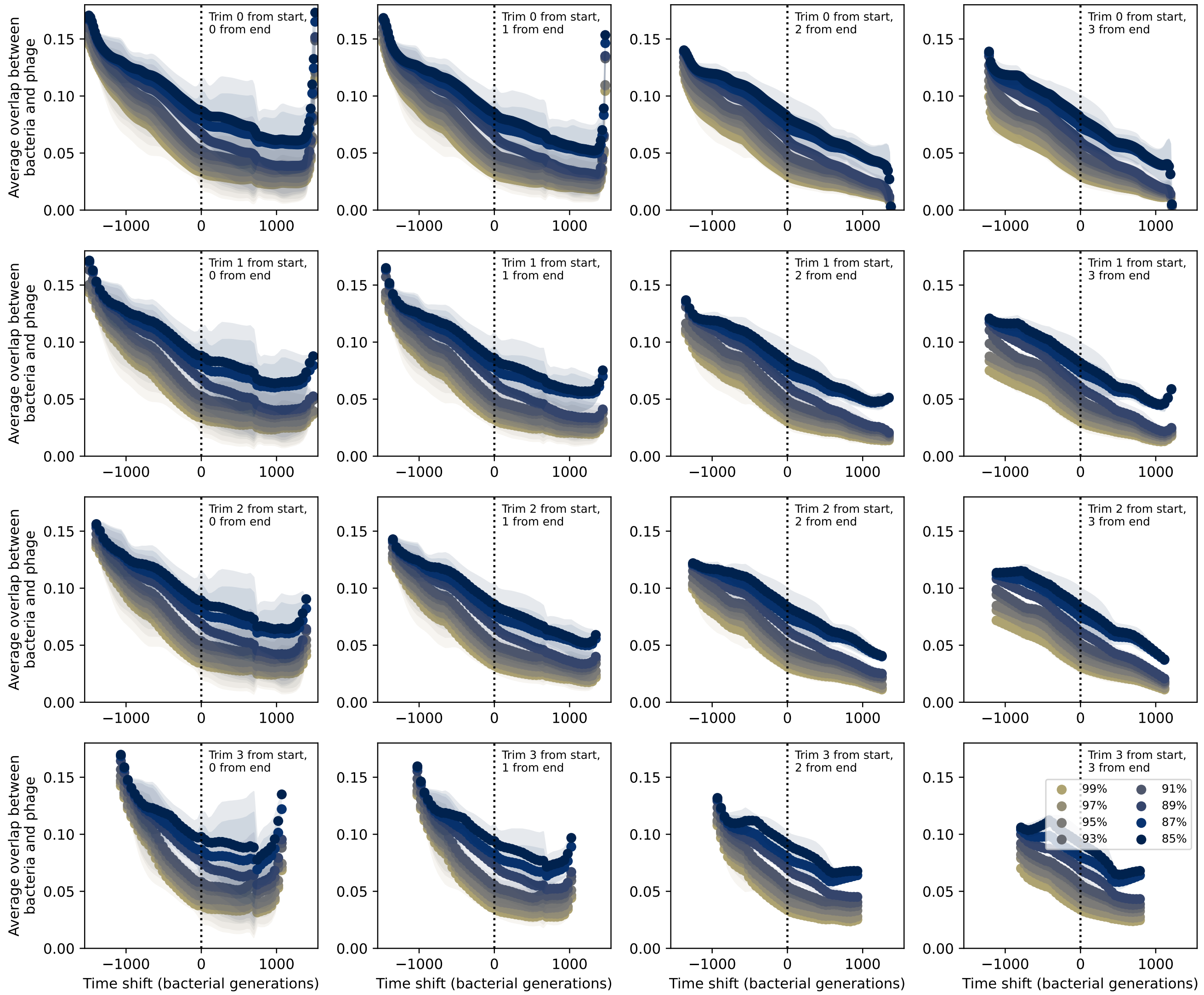

Figure 103: Time shifted overlap with increasing numbers of time points from the start and end removed. Interpolation spacing is the smallest interval between remaining time points. Only perfect-PAM protospacers are included; all wild-type spacers are included.

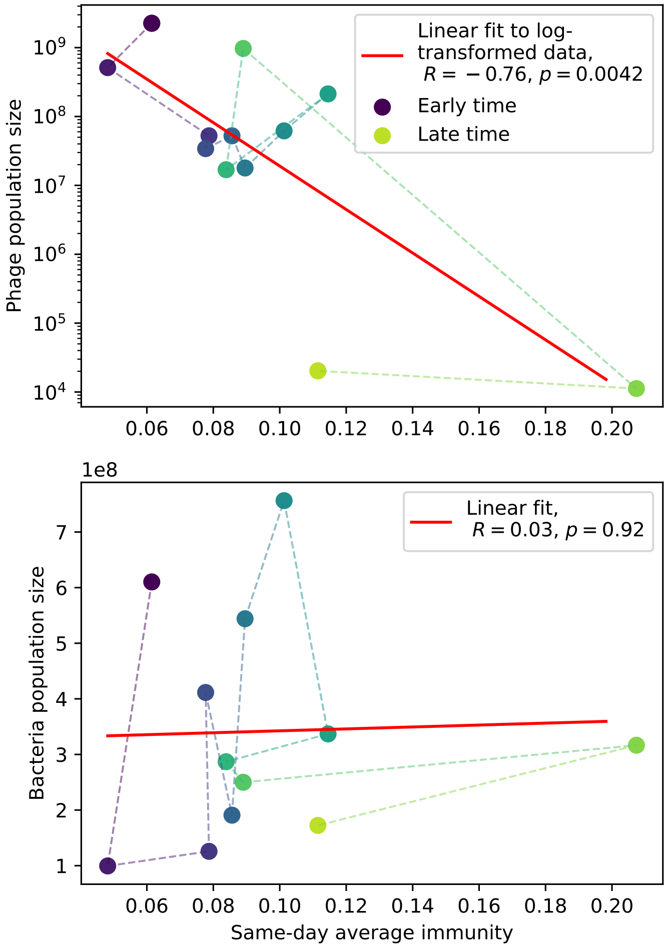

Figure 104: Phage (top) and bacteria (bottom) population size vs. average immunity calculated with data from [38]. Protospacers are included if they have a perfect PAM, and all wild-type spacers are included. Spacers and protospacers are grouped with an 85% similarity threshold. Colours from blue to yellow indicate increasing time.

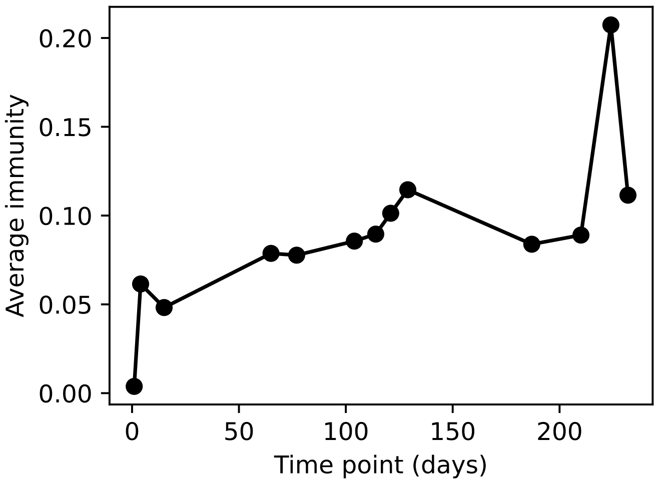

Figure 105: Average immunity vs time in the MOI-2B experiment. Protospacers are included if they have a perfect PAM, and all wild-type spacers are included. Spacers and protospacers are grouped with an 85% similarity threshold.

searched for protospacers within these phage genomes using blast. They did not find matches between the *Gordonia* spacers and any other contigs from the dataset, suggesting that those two phages are the only two from which *Gordonia* is acquiring spacers. They used Crass and metaCRT to detect other possible CRISPR arrays within the unassembled data and other contigs; they did not find and/or do not discuss any other detected CRISPR loci. They found a broad fan-like network of CRISPR arrays within *Gordonia* that reflect new spacer acquisition, many of which matched the phage genomes. Older, conserved spacers largely did not match the phage genomes. They found that peaks in phage genome coverage (a proxy for abundance) over time were highly correlated with bacterial coverage, suggesting that their population dynamics are tightly coupled. They identified a PAM sequence, GTT (5') for the CRISPR1 locus, but did not find that mutations occurred more frequently in the protospacer regions or PAM regions of the phage genomes than elsewhere in the genome.

Following the same analysis procedure we used for data from ref. [38] (section 6.2), we used the two reported *Gordonia* CRISPR repeats (`GTGCTCCCCGCGCGAGCGGGGATGATCCC` for CRISPR1 (Type-I) and `ATCAAGAGCCATGTCTCGCTGAACAGCGAATTGAAAC` for CRISPR2 (Type-III)) to search for spacers in the raw read data using BLAST.

To detect wild-type spacers, we searched for matches to the CRISPR repeats on the *Gordonia* metagenome-assembled genome (MAG) provided on the study's GitHub repository and extracted the sequences between repeat matches. We determined that spacers from the CRISPR1 locus had an average length of 32 nt and spacers from the CRISPR2 locus had an average length of 35 nt.

We detected spacers as sequences between or adjacent to repeat sequences. To maximize the number of genuine spacers detected while removing low-quality matches, we kept only full-length alignments to the repeat, unless a shorter alignment was also present on the same read as a full-length alignment (for instance if the repeat is partially present at the start or end of a read). If a single repeat match was present on a read, we extracted 32 (CRISPR1) or 35 (CRISPR2) nt on either side and labelled these spacers. We set a minimum spacer length of 85% of the expected spacer length and did not keep spacer sequences at the start or end of a read that were shorter than this. If multiple repeat matches were present, we extracted intervening sequences as spacers.

We grouped all extracted spacers using the AgglomerativeClustering method from scipy with an 85% similarity threshold and the 'average' linkage criterion. We performed this grouping separately for CRISPR1 and CRISPR2 spacers. Each spacer sequence was then labelled with a group type identifier.

To detect protospacers, we first blasted all reads against the *Gordonia* MAG and the two reported phage genomes (DC-56 and DS-92). We next blasted all unique detected spacer sequences from the previous step against all the reads, removing any query sequences that are a perfect subset of another sequence and any sequences that were more than 30% N nucleotide. This detects both potential protospacer sequences and the original spacers themselves. To isolate protospacers from spacers, we kept only results that were on reads that did not match the CRISPR1 or CRISPR2 repeat. Some reads matched both the bacterial genomes and the phage genomes; we removed matches for which the log-difference in e-value between phage genome match and bacteria genome match was less than 25, i.e. the phage genome e-value must be lower than the bacteria e-value by a factor of $10^{25}$ to be considered not a bacteria match. Figure 106 shows the e-value differences for reads that matched both phage and bacteria for one time point.

We kept only potential protospacers that were 85% of the expected spacer length or larger. If there were multiple hits on a read from the same spacer type (but different sequences), we kept only the match with the lowest e-value. We extracted the matched sequence from the read and ten nucleotides on either side to check for a PAM sequence (all spacers were stored oriented in the same direction relative to the repeat to facilitate comparison and PAM detection). Consistent with the PAM reported by Guerrero *et al.*, we identified a GTT PAM at the 5' end of protospacer sequences (Figure 107, reverse-complement).

Since reads are paired-end, some reads will overlap, and some spacers may be double-counted in the overlap. We decremented the total spacer or protospacer count for a sequence by 1 if a spacer sequence

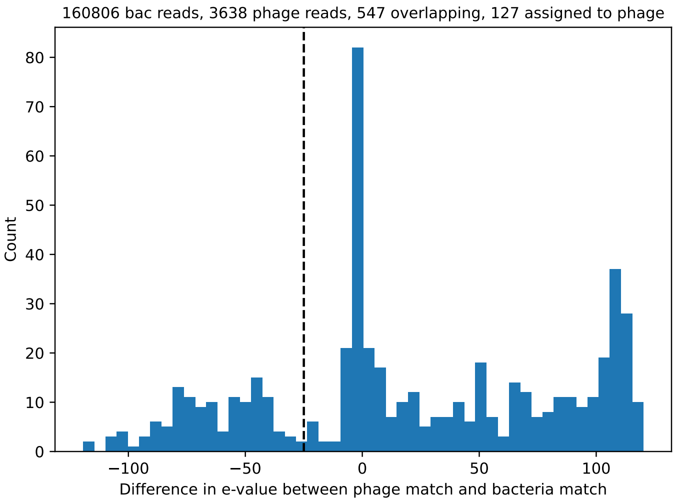

Figure 106: Histogram of reads that matched both the *Gordonia* MAG and either phage reference genome vs. the base-10 log difference in e-value between the matches for accession SRR9260993. A positive value means the bacteria match has a lower e-value than the phage match. The vertical dashed line indicates the −25 cutoff; matches to the left were considered phage matches, matches to the right were considered bacteria matches.

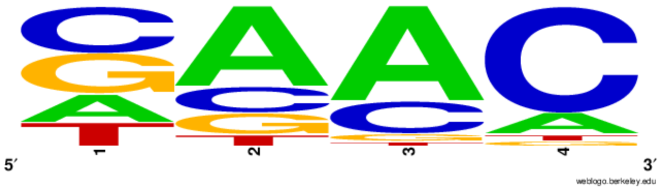

Figure 107: Probability logo for the 4 nucleotides at the 5' end of potential protospacer sequences, generated with WebLogo; the spacer sequence starts at the right edge of the logo meaning that the reverse-complement PAM is GTT.

was present on both ends of a paired read.

We grouped all spacer and protospacer sequences together (separated by CRISPR locus) by several different similarity thresholds between 85% and 99% to assign type labels based on each similarity threshold.

We checked the 10 nt region upstream from each potential protospacer and removed any sequences that did not have a perfect match to the GTT PAM. Since targeting is highly sensitive to PAM mutations [43], we assumed that any deviation from the perfect PAM meant that the protospacer would not be successfully targeted. There are cases where the PAM sequence is incomplete because of hitting the start or end of a read; these were also assumed to be imperfect PAMs (though some would be genuine). We detected 627555 potential protospacers for the CRISPR1 locus and only 61 potential protospacers for the CRISPR2 locus. Since the CRISPR2 locus is a Type III CRISPR system, we expect that there is no PAM for this system and so included all CRISPR2 protospacers; because there are so few, this choice does not affect results. After removing all CRISPR1 protospacers that did not have an adjacent match to the perfect PAM, we were left with 102761 protospacers in total representing 19292 unique sequences and 537 unique types after grouping with an 85% similarity threshold. The total number of unique spacers detected from the CRISPR1 locus over the experiment was 5289 (2078 unique types at 85% similarity), while the total number from the CRISPR2 locus was an order of magnitude lower at 972 (357 unique types at 85% similarity). In total, 28386 CRISPR1 spacers and 3353 CRISPR2 spacers

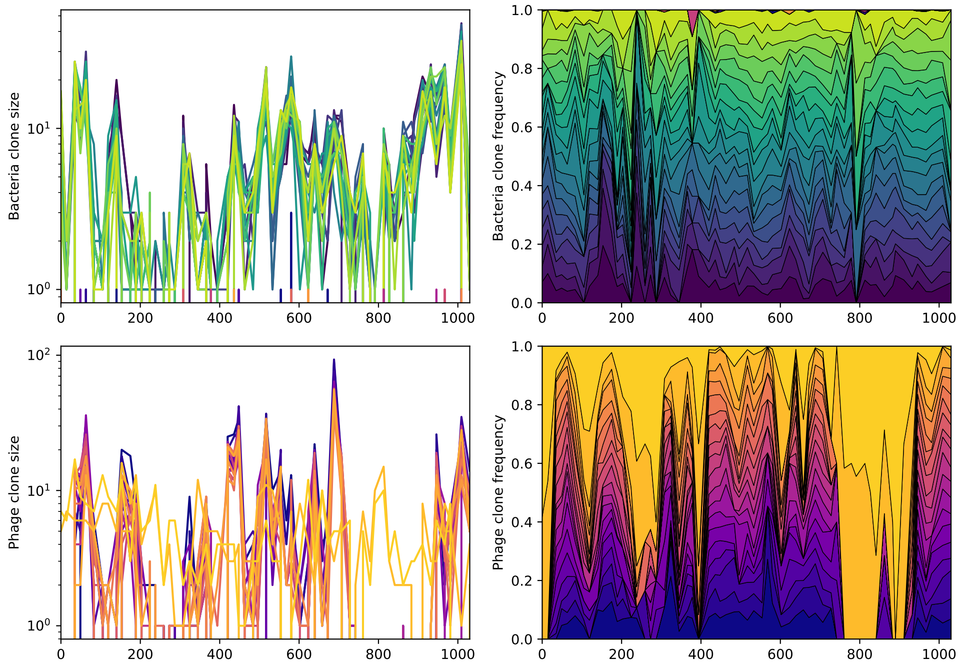

Figure 108: Abundance over time for the 20 largest bacteria clones (top) and phage clones (bottom) over time for data from [45]. The left panels show absolute counts, the right panels show fractional abundance for the included types. None of the top types are shared between phage and bacteria.

were detected.

Our detection of reads that matched the *Gordonia* MAG and phage genomes is closely correlated with the reported coverage from the study (Figure 109).

The number of unique protospacer and spacer types we detected fluctuated considerably over the course of the study (Figure 110). The number of shared types as well as the ratio of phage types to bacteria types also fluctuated considerably.

About 40% of spacer and protospacer types remained throughout the time series without going extinct at an 85% similarity threshold (Figure 112). In data from [38], about 20% of original types remained at the end of the experiment (main text Figure 6E), however the fraction of types remaining decreases steadily in data from [38] unlike in this dataset.

##### 6.3.1 Calculating average immunity

To calculate average immunity between bacteria and phage, we interpolated counts between sequenced time points using a 14-day spacing (the most common sampling interval in the data). We calculated the time-shifted average overlap between bacteria and phage spacer types by comparing bacteria and phage types separated by a time delay and averaging over all points with the same time delay (Figure 113). As described in section 6.1, we multiplied raw average immunity values by the average number of protospacers with the GTT PAM from the phage DC-56 and DS-92 genomes (1956 protospacers) to account for multiple protospacers on each phage genome. Note that because PAM mutations are common, the true number of protospacers may be less than this hypothetical amount, which would cause us to slightly overestimate average immunity. However, we are also assuming that each bacterium has one effective spacer, and this is likely an underestimate.

As the similarity threshold increases, the overall overlap goes down slightly because there are more total types. As in the data from [38], the number of shared types across the whole dataset is insensitive to the clustering threshold: there is a tradeoff between the total number of types (which increases as the

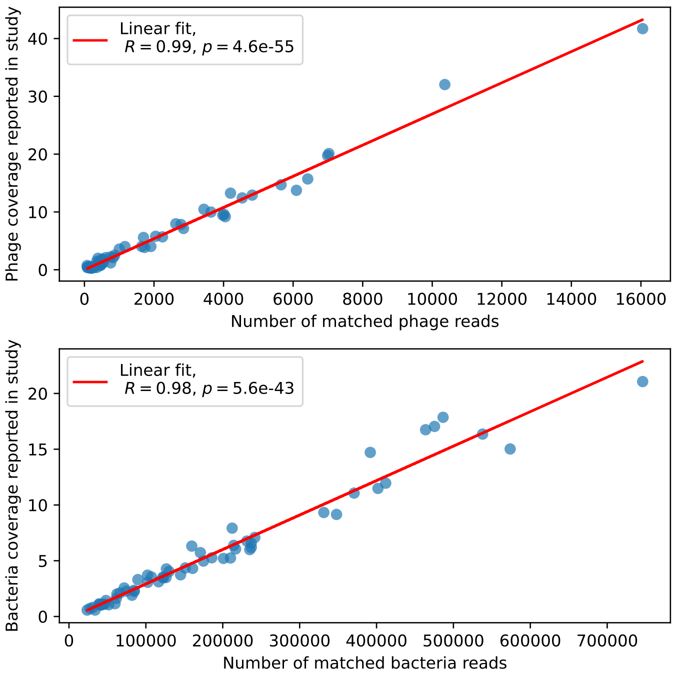

Figure 109: Phage genome (top) and bacteria genome (bottom) coverage, digitized from Figure 2D of ref. [45] vs. number of reads assigned to phage or bacteria from our analysis. Each marker is a separate time point.

similarity threshold increases) and the likelihood that types are shared (which increases as the similarity threshold decreases) (Figure 110).

We experimented with different trimming thresholds for the data, removing time points from the beginning and end of the data series before calculating time-shifted average immunity. For all similarity thresholds and most trimming choices, there is a peak in average immunity at zero time shift that decays in both the past and future directions (Figure 113). However, if enough data is trimmed from the beginning to remove the large spikes in average immunity between days 200 and 400 (Figure 114), the central peak is quite diminished or removed altogether (Figures 121 and 122). The standard deviation of average immunity is large near the zero time shift (Figure 119), which is because average immunity is highly variable across the time series: there is a large spike in average immunity near the middle of the time series (Figure 114).

Because the variability in average immunity between time points was very high, we performed a Wilcoxon signed-rank test between the average immunity at zero time delay and the average immunity at a time delay ±200 and ±500 (arbitrarily chosen). We paired each shifted time point with its corresponding average immunity at zero delay for each bacterial abundance. Using a paired test answers the following question for each time point: is the overlap between bacteria and past phages significantly lower than the overlap in the present, and is the overlap between bacteria and future phages significantly lower than the overlap in the present? We found that past immunity after 200 days is significantly lower than present ($Z = 1241, p = 0.008$), but that immunity after 500 days is not significantly lower than present ($Z = 392, p = 0.27$). When comparing all time points with present, only a small range of past immunity values were significantly lower than present (SI Figure 118). For both time delays, we found that future immunity is significantly lower than present ($Z = 580, p = 0.0012$ for 500 days and $Z = 1293, p = 0.003$ for 200 days). Interestingly, these significance values are not symmetric if we pose the question from the perspective of phage: the overlap between phage and future bacteria at 500 days is lower than the overlap for present phages ($Z = 507, p = 0.024$), and the overlap between

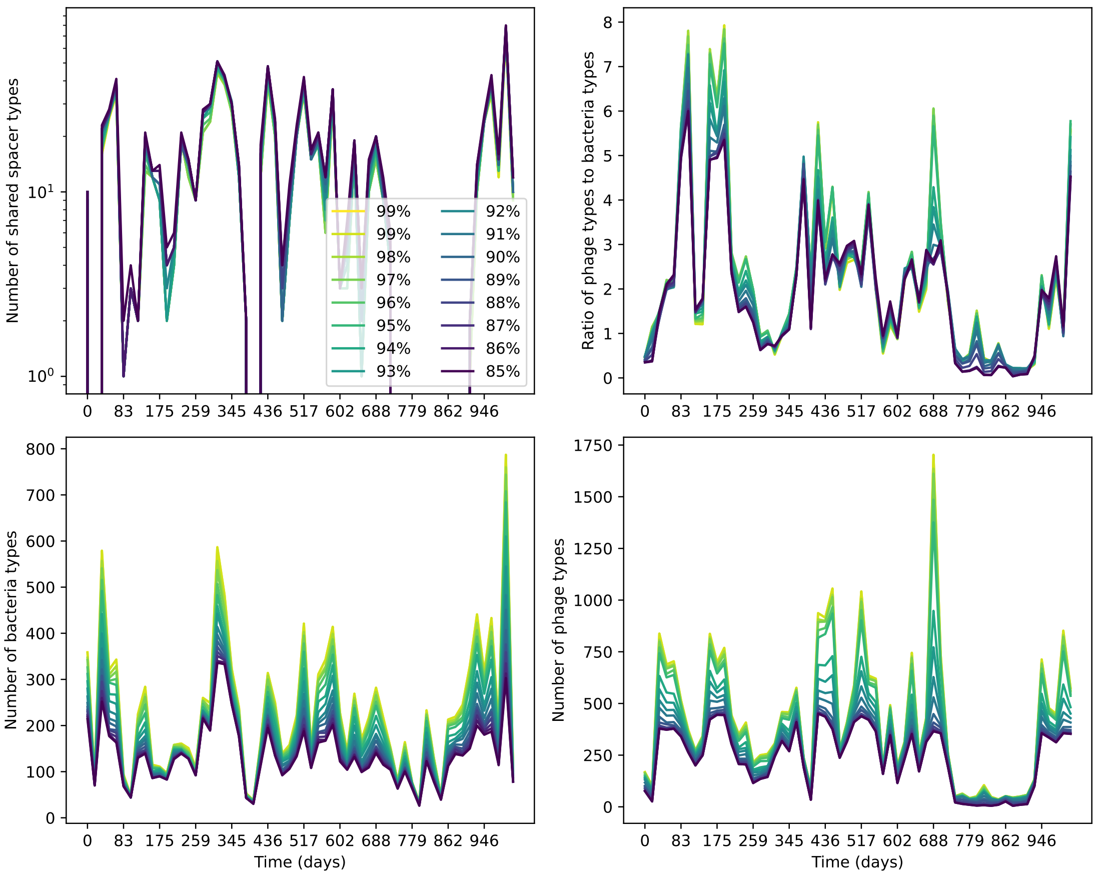

Figure 110: Number of shared spacer types between bacteria and phage (top left), ratio of the number of phage types to the number of bacteria types (top right), number of bacteria types (bottom left), and number of phage types (bottom right) as a function of the sampling date for data we analyzed from [45]. Colours indicate different similarity grouping thresholds. Spacer counts include wild-type spacers, and protospacers are included only if they possess a perfect PAM. Data is summed over CRISPR1 and CRISPR2.

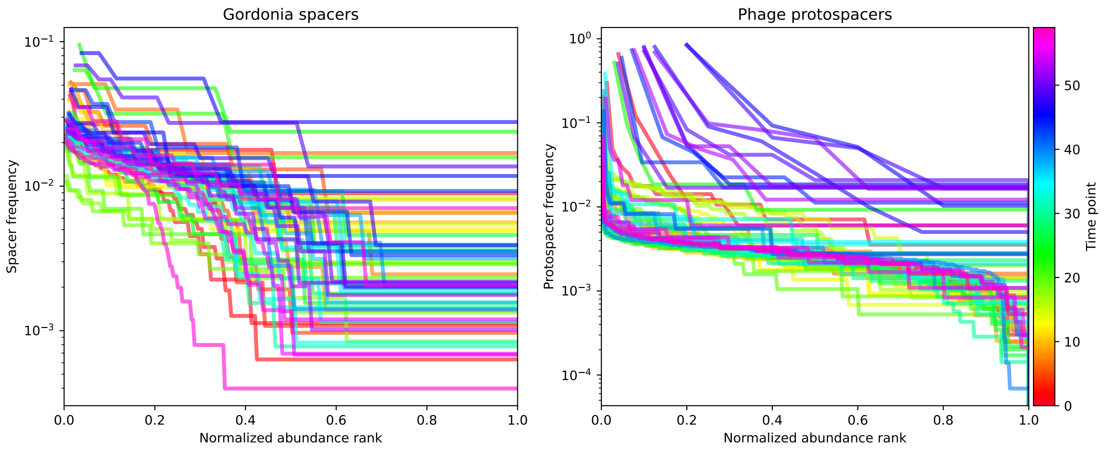

Figure 111: Rank abundance distribution for detected spacers (left) and protospacers (right) at each time point in data from [45]. Spacers are generally present at a range of abundances; protospacer abundances are more tightly peaked (a flatter rank-abundance curve).

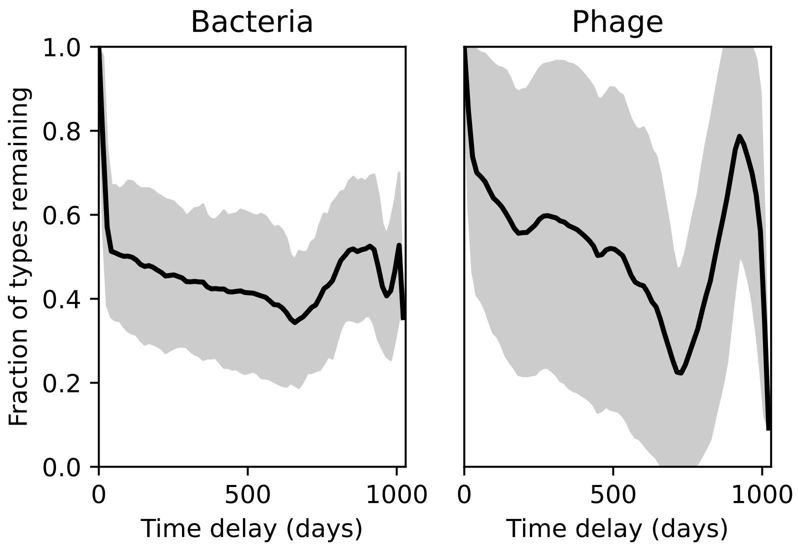

Figure 112: Fraction of spacer types (left) and protospacer types (right) remaining as a function of time delay, averaged over the entire time series from [45]. Types are grouped with an 85% similarity threshold.

phage and past bacteria is lower than the overlap at present($Z = 388, p = 0.027$.) This asymmetry, that bacteria are generally more immune to all past phages while phage are not more infective against all past bacteria, is qualitatively the same as that reported by Dewald-Wang *et al.* in an explicit time shift study of bacteria and phage immunity and infectivity in chestnut trees [46].

We confirmed that the past and future shifted average immunity becomes less correlated with the zero-shift average immunity as the time shift increases. Figure 123 shows the time-shifted average immunity as a function of the zero-shift average immunity for three time shift values, and we see that average immunity is highly correlated at small time shifts but becomes less correlated at larger shifts.

We randomly shuffled time point labels for bacteria and phages separately 115) and together 116) and calculated average immunity with the bootstrapped data. The average immunity trend as a function of time shift disappears in the bootstrapped data even if bacteria and phage clone abundances remain matched at each time point.

Unlike in the data from [38], we did not find a correlation between phage or bacteria coverage and average immunity, though ref. [45] did report that bacteria and phage coverage was correlated and that periods of high phage coverage coincided with higher abundance of *Gordonia* strains lacking matching spacers, suggesting that the expected negative correlation between phage abundance and bacterial immunity may be playing out at some level but may be confounded by generally higher detected average immunity when coverage is high.

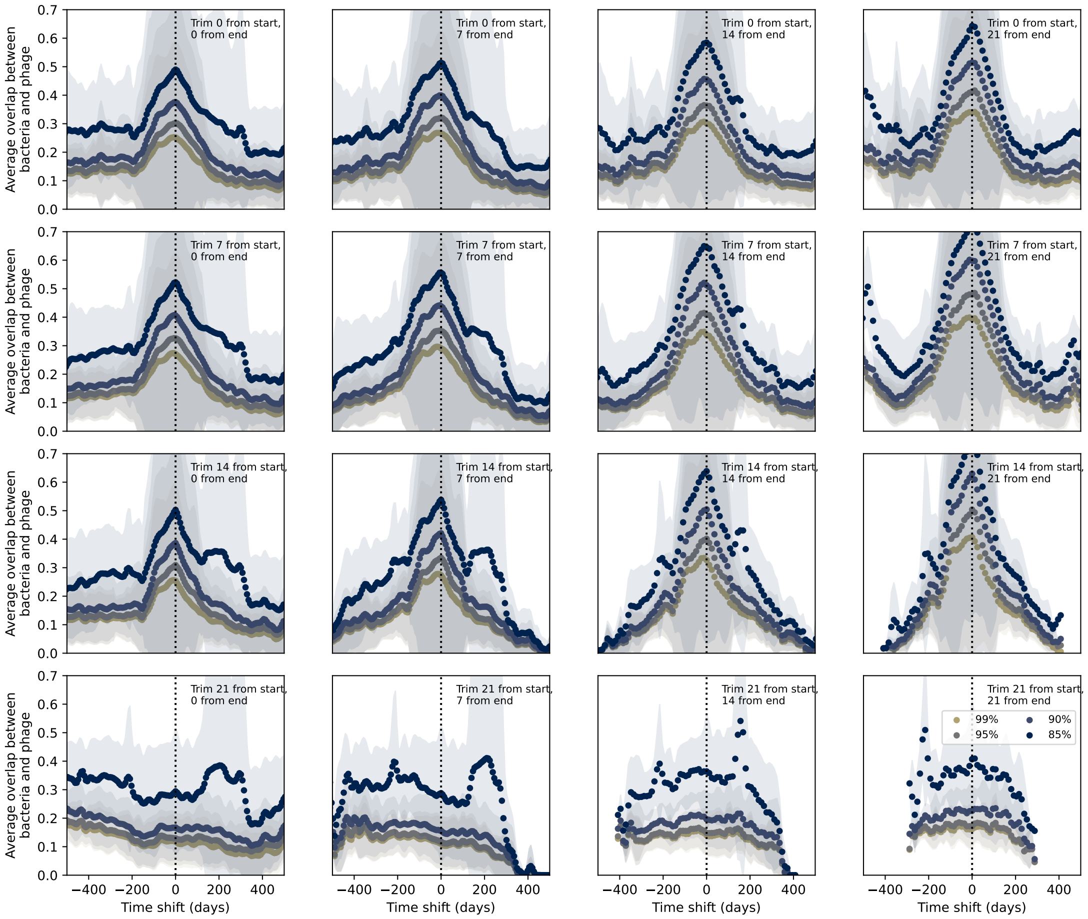

Figure 113: Time shifted average immunity for spacers and protospacers grouped at four different similarity thresholds. Points are trimmed from the start and end of the time series in each panel as described in the overlay text. Raw values are multiplied by 1956, the average number of protospacers with the GTT PAM from the phage DC-56 and DS-92 genomes. Only perfect-PAM protospacers are included; all wild-type spacers are included.

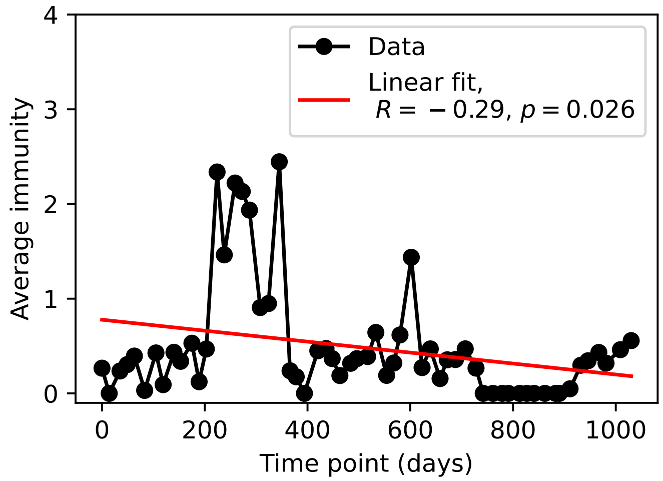

Figure 114: Average immunity at each time point in the time series. Raw values cannot be larger than one, but plotted values are multiplied by 1956, the average number of protospacers with the GTT PAM from the phage DC-56 and DS-92 genomes, yielding some values larger than one.

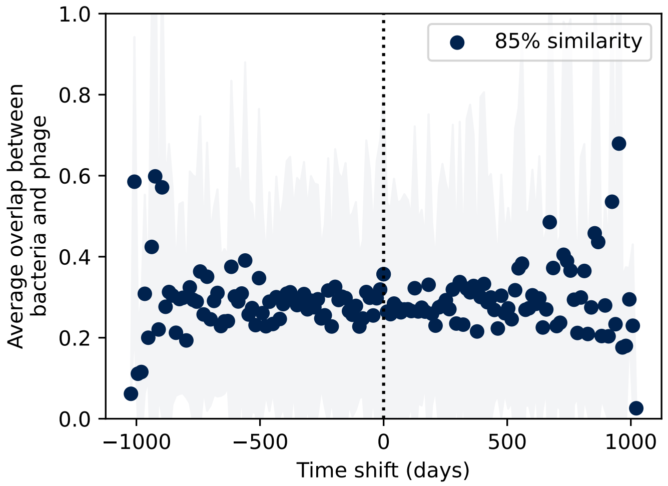

Figure 115: Bootstrapped control: time shifted average immunity after randomly shuffling the interpolated clone abundances separately for bacteria and phage.

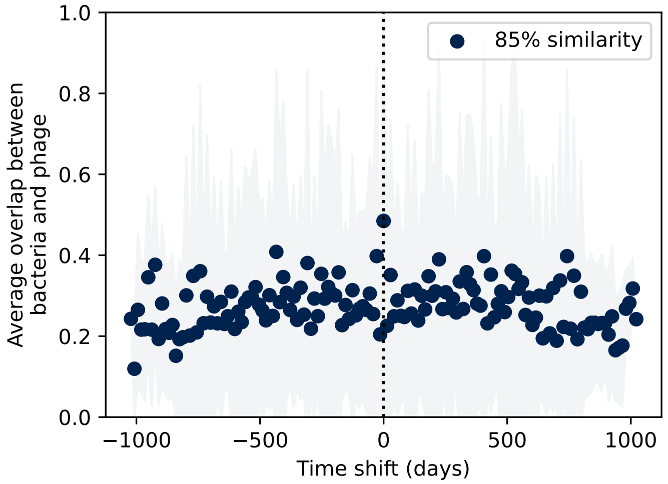

Figure 116: Bootstrapped control: time shifted average immunity after randomly shuffling pairs of interpolated bacteria and phage clone sizes such that time point matching is maintained between bacteria and phages.

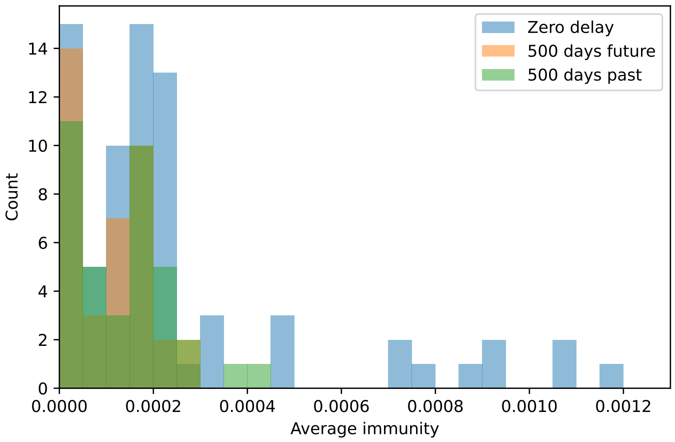

Figure 117: Distribution of average immunity values at zero time shift (blue) and at 500 days time shift for future phages (orange) and 500 days time shift for past phages (green).

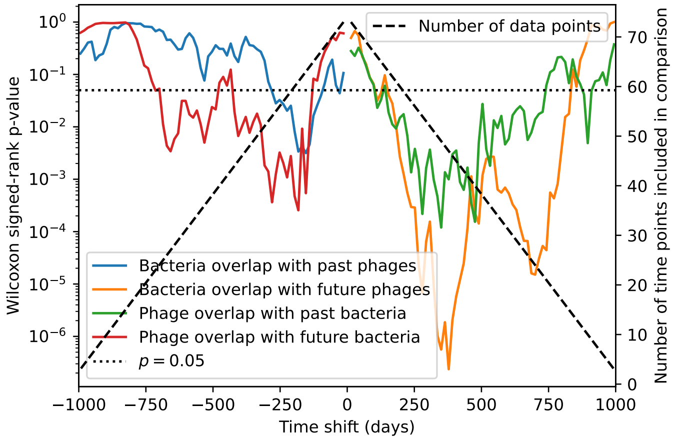

Figure 118: $p$-values for the Wilcoxon signed-rank test comparing the average immunity at zero time shift with all other time shifts. Bacteria immunity against past phages is generally not significantly lower than bacterial immunity against current phages (blue), while bacterial immunity against future phages is significantly lower than immunity against current phages for almost all time shifts (orange). The time shifts are not symmetric; bacterial overlap with past phages is not necessarily the same as phage overlap with future bacteria and vice versa. The number of points that are available to compare decreases as the time shifts get larger (black dashed line).

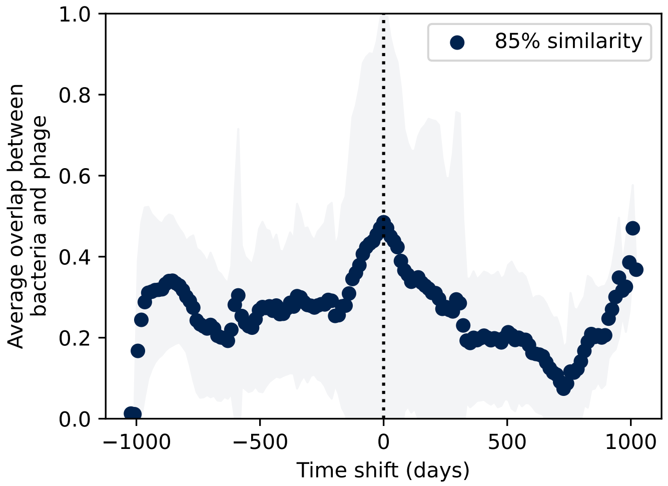

Figure 119: Time shifted average immunity for spacers and protospacers grouped at an 85% similarity threshold. All time points are included. Raw values are multiplied by 1956, the average number of protospacers with the GTT PAM from the phage DC-56 and DS-92 genomes. Only perfect-PAM protospacers are included; all wild-type spacers are included.

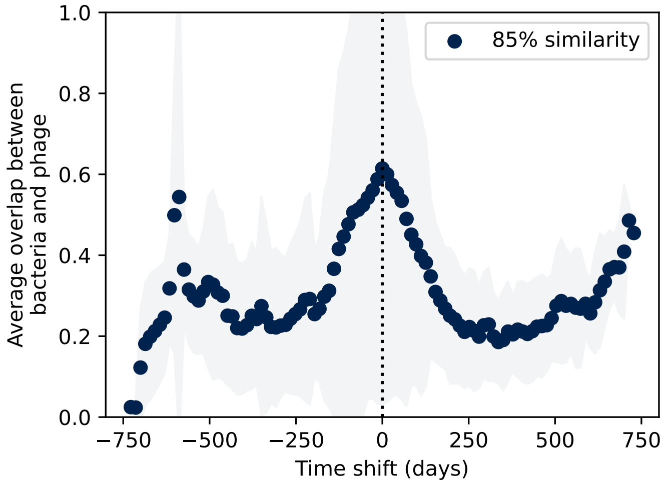

Figure 120: Time shifted average immunity for spacers and protospacers grouped at an 85% similarity threshold. 16 time points are removed from the end of the data series in order to remove the region with zero protospacer counts. Raw values are multiplied by 1956, the average number of protospacers with the GTT PAM from the phage DC-56 and DS-92 genomes. Only perfect-PAM protospacers are included; all wild-type spacers are included.

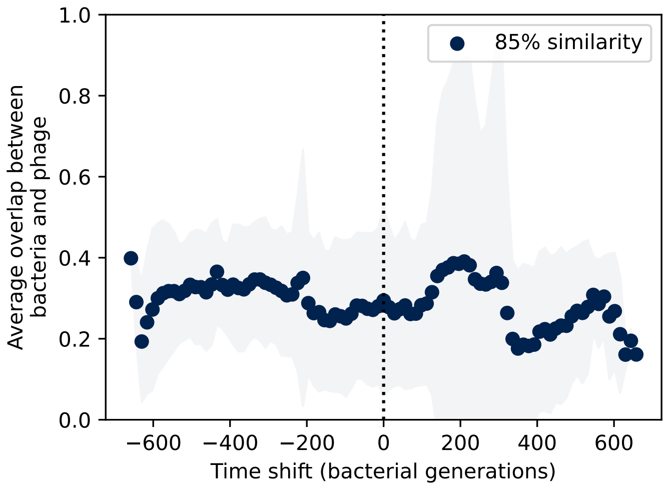

Figure 121: Time shifted average immunity for spacers and protospacers grouped at an 85% similarity threshold. 21 time points are removed from the beginning of the data series to remove the region with anomalously high average immunity. Raw values are multiplied by 1956, the average number of protospacers with the GTT PAM from the phage DC-56 and DS-92 genomes. Only perfect-PAM protospacers are included; all wild-type spacers are included.

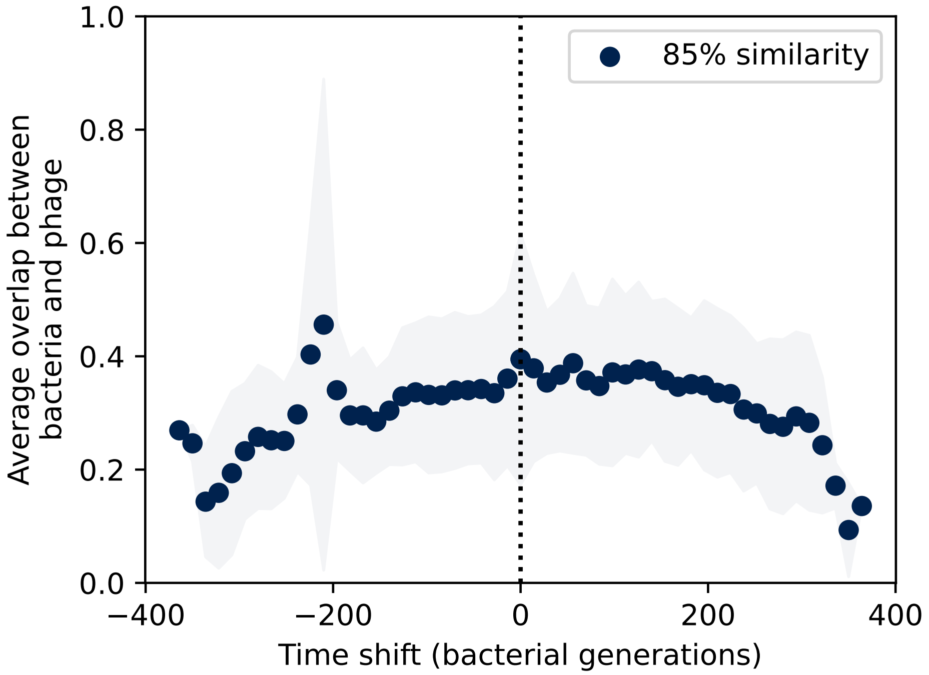

Figure 122: Time shifted average immunity for spacers and protospacers grouped at an 85% similarity threshold. 21 time points are removed from the beginning of the data series to remove the region with anomalously high average immunity, and 16 time points are removed from the end of the data series in order to remove the region with zero protospacer counts. Raw values are multiplied by 1956, the average number of protospacers with the GTT PAM from the phage DC-56 and DS-92 genomes. Only perfect-PAM protospacers are included; all wild-type spacers are included.

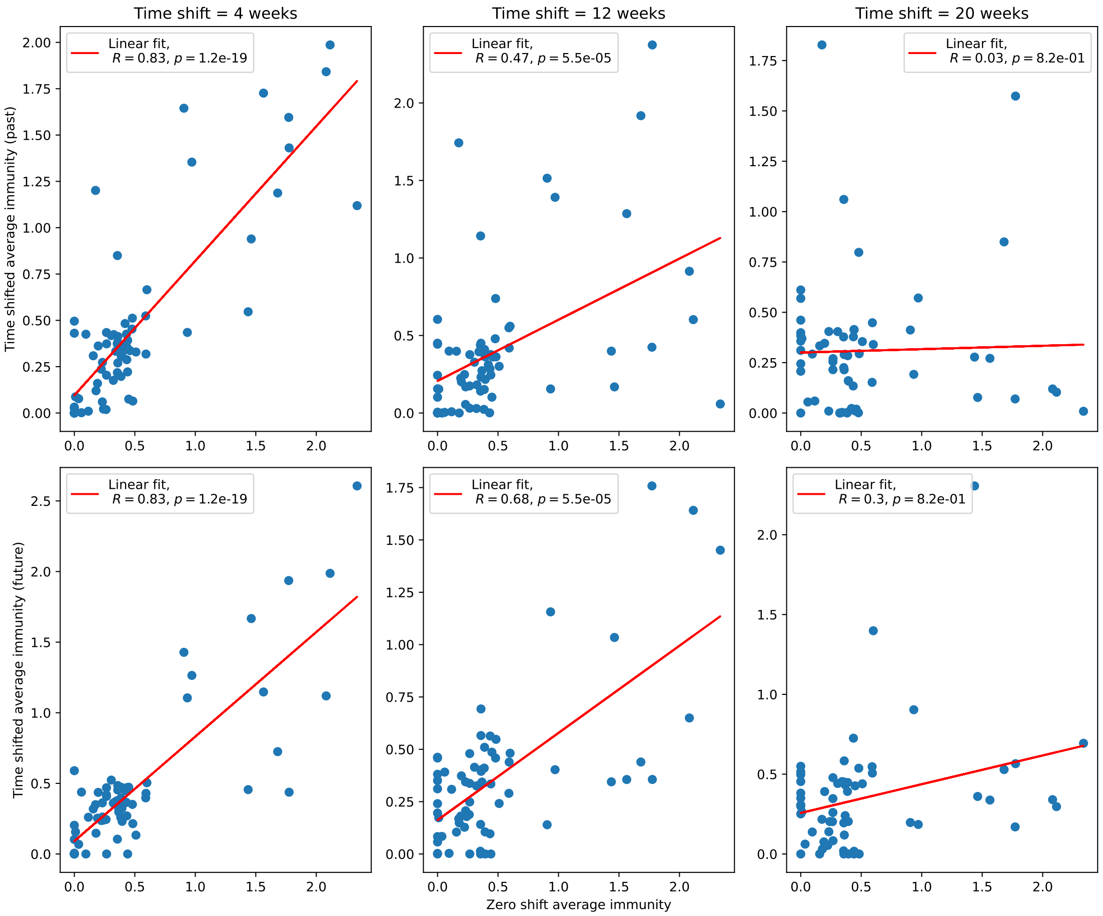

Figure 123: Time-shifted average immunity for each time point as a function of the zero-shift average immunity of that data point for shifts comparing past phages (top row) and future phages (bottom row).

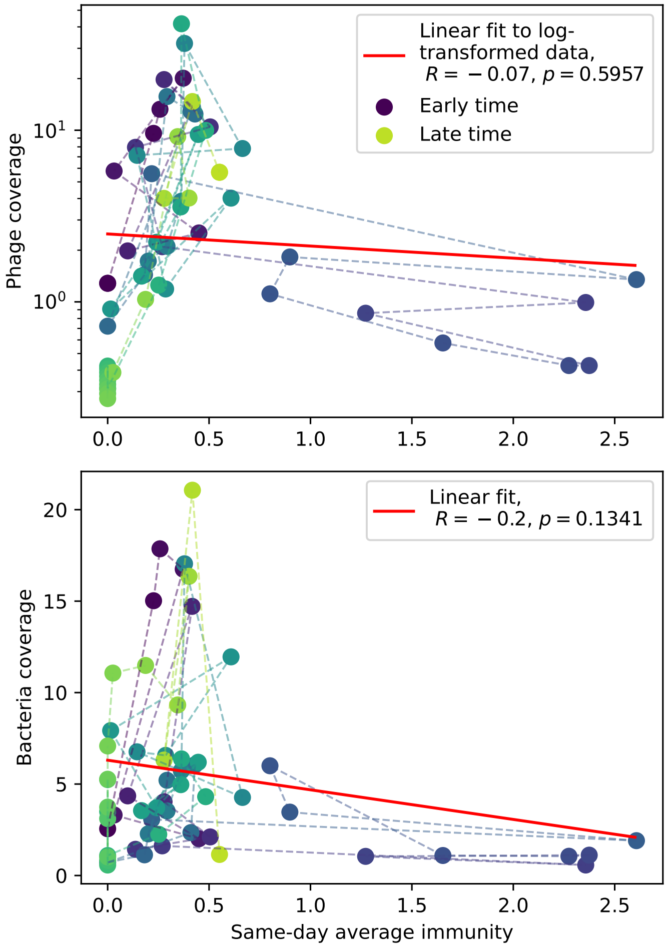

Figure 124: Phage (top) and bacteria (bottom) population size vs. average immunity calculated with data from [45]. Protospacers are included if they have a perfect PAM, and all wild-type spacers are included. Spacers and protospacers are grouped with an 85% similarity threshold. Colours from blue to yellow indicate increasing time.

[6] Marcelo Ponce, Ramses Van Zon, Scott Northrup, Daniel Gruner, Joseph Chen, Fatih Ertinaz, Alexey Fedoseev, Leslie Groer, Fei Mao, Bruno C. Mundim, Mike Nolta, Jaime Pinto, Marco Saldarriaga, Vladimir Slavnic, Erik Spence, Ching Hsing Yu, and W. Richard Peltier. Deploying a top-100 supercomputer for large parallel workloads: The Niagara supercomputer. *ACM International Conference Proceeding Series*, 2019.

[7] Yang Cao, Daniel T. Gillespie, and Linda R. Petzold. Efficient step size selection for the tau-leaping simulation method. *The Journal of chemical physics*, 124(4):044109, 2006.

[8] Pu Han, Liang Ren Niestemski, Jeffrey E. Barrick, and Michael W. Deem. Physical Model of the Immune Response of Bacteria Against Bacteriophage Through the Adaptive CRISPR-Cas Immune System. *Physical Biology*, 10(025004):025004, apr 2013.

[9] Pu Han and Michael W Deem. Non-classical phase diagram for virus bacterial coevolution mediated by clustered regularly interspaced short palindromic repeats. *Journal of The Royal Society Interface*, 14(127):20160905, 2017.

[10] Sacha Lucchini. *Genetic Diversity of Streptococcus thermophilusPhages and Development of.* PhD thesis, Swiss Federal Institute of Technology Zurich, 1999.

[21] Raymond H.J. Staals, Simon A. Jackson, Ambarish Biswas, Stan J.J. Brouns, Chris M. Brown, and Peter C. Fineran. Interference-driven spacer acquisition is dominant over naive and primed adaptation in a native CRISPR-Cas system. *Nature Communications*, 7, 2016.

[35] N. G. van Kampen. *Stochastic Processes in Physics and Chemistry.* Elsevier, third edition, 1981.

[36] M Vallade, B Houchmandzadeh, M Vallade, and B Houchmandzadeh Analytical. Analytical solution of a neutral model of biodiversity. *Physical Review E : Statistical, Nonlinear, and Soft Matter Physics*, 68, 2003.
